# Borrowing Transcriptional Kinases to Activate Apoptosis

**DOI:** 10.1101/2023.10.23.563687

**Authors:** Roman Sarott, Sai Gourisankar, Basel Karim, Sabin Nettles, Haopeng Yang, Brendan G. Dwyer, Juste M. Simanauskaite, Jason Tse, Hind Abuzaid, Andrey Krokhotin, Tinghu Zhang, Stephen M. Hinshaw, Michael R. Green, Gerald R. Crabtree, Nathanael S. Gray

## Abstract

Protein kinases are disease drivers whose therapeutic targeting traditionally centers on inhibition of enzymatic activity. Here chemically induced proximity is leveraged to convert kinase inhibitors into context-specific activators of therapeutic genes. Bivalent molecules that link ligands of the transcription factor B-cell lymphoma 6 (BCL6) to ATP-competitive inhibitors of cyclin-dependent kinases (CDKs) were developed to re-localize CDK to BCL6-bound loci on chromatin and direct phosphorylation of RNA Pol II. The resulting BCL6-target proapoptotic gene expression translated into killing of diffuse large B-cell lymphoma (DLBCL) cells at 72 h with EC50s of 0.9 – 10 nM and highly specific ablation of the BCL6-regulated germinal center response in mice. The molecules exhibited 10,000-fold lower cytotoxicity in normal lymphocytes and are well tolerated in mice. Genomic and proteomic evidence corroborated a gain-of-function mechanism where, instead of global enzyme inhibition, a fraction of total kinase activity is borrowed and re-localized to BCL6-bound loci. The strategy demonstrates how kinase inhibitors can be used to context-specifically activate transcription, accessing new therapeutic space.

## INTRODUCTION

Protein kinases play pivotal roles in cellular signaling and are among the most important drug targets(*1*). Many are key regulators of transcription, such as the cyclin-dependent kinases (CDKs) CDK9, CDK12, and CDK13 that function in concert with their cyclin binding partners to modulate RNA Polymerase II (Pol II) activity in the nucleus. CDK9 and one of cyclins T1, T2, or K form the <u>p</u>ositive transcription <u>e</u>longation factor <u>b</u> (P-TEFb) complex to enable the release of paused Pol II into elongation by phosphorylation of negative elongation factors and serine 2 (ser 2) of the Pol II C-terminal domain (CTD) (*2*). Since many cancers are addicted to the transcription of proto-oncogenes such as *MYC* (*3*), potent and specific ATP-competitive inhibitors have been developed to silence the activity of CDK9 and other transcriptional kinases to abolish oncogenic transcription (*4, 5*).

Approaches using chemically induced proximity have recently emerged as promising alternatives to small molecule inhibitors. Small molecule <u>c</u>hemical inducers of <u>p</u>roximity (CIPs) that induce molecular proximity between cellular proteins have been used to recapitulate diverse biological processes in living cells and organisms, including post-translational modification, signal transduction, and transcription (*6*). Among the advantageous features of CIPs is their event-driven mechanism of action. CIP induction of protein-protein proximity can be catalytic for a cellular process of interest (*7*). Bivalent small molecules called proteolysis-targeting chimeras (PROTACs) that target proteins for degradation by the ubiquitin-proteasome system represent one of the most established small molecule modalities relying on CIP and have enabled removal of disease-relevant proteins including kinases (*8, 9*). While mechanistically distinct, both inhibitors and protein degraders phenocopy genetic loss of protein function.

We hypothesized that CIP may be used to turn kinase inhibitors into activators of epigenetically silenced transcriptional states. To realize this concept, we focused on the induction of genes silenced by zinc-finger transcription factor BCL6 (<u>B</u>-<u>c</u>ell lymphoma <u>6</u>), which is de-regulated and overexpressed in 40-60% of cases of human diffuse large B-cell lymphoma (DLBCL), driving the progression of this cancer (*10, 11*). During germinal center B cell development, BCL6 ordinarily silences the transcription of tumor suppressor and programmed cell death (apoptotic) genes, including *PMAIP1/*NOXA*, TP53*, and *CDKN1B/*p27(*12*). Target gene silencing is mediated by epigenetic co-repressors such as BCOR, NCOR1, and SMRT that bind to the BCL6 N-terminal BTB domain (BCL6^BTB^) (*13, 14*). BCL6 acts as an oncogene in DLBCL by suppressing DNA damage response and cell death pathways and is deregulated directly by chromosomal translocations and mutations in regulatory regions (*11*) or indirectly via inactivating mutation of antagonistic factors(*15*). Multiple high-affinity inhibitors of BCL6^BTB^ and BCL6 degraders have been developed but exhibit only modest effects on tumor anti-proliferation (*16–18*). We previously reported a strategy to directly activate BCL6-target gene expression by CIP-mediated recruitment of the elongation factor BRD4 to BCL6-bound loci (*19*).

A recent genome-wide screen for proteins that induce transcription via CIP nominated several kinases ordinarily involved in elongation of RNA Pol II for CIP-based transcription induction (*20*). Owing to the myriad of clinical-stage, small molecule binders of kinases, we explored whether recruitment of transcription elongation complexes via their kinase subunits to DNA sequences bound by BCL6 can activate BCL6-repressed cell death. We devised a general CIP-based strategy (Fig. 1A) to recruit endogenous CDK9 and other elongation factor kinases to induce the transcription of BCL6-regulated death genes in lymphoma cells. Our results showed that ATP-competitive kinase inhibitors can be converted to nanomolar activators of transcription when linked to ligands of BCL6^BTB^. Such molecules, termed CDK-<u>T</u>ranscriptional/epigenetic <u>C</u>hemical Inducers of <u>P</u>roximity (CDK-TCIPs) operate by rapid re-localization of a fraction of CDK9, formation of ternary complexes at BCL6-bound loci, induction of Pol II ser 2 phosphorylation (Pol II ser 2 phos), and transcription of pro-apoptotic, BCL6-target genes. The cell-killing effect was specific to BCL6-driven cells with a 10,000-fold difference in cytotoxicity between DLBCL cells and primary human lymphocytes. We also illustrate a sequential array of targeted assays to predictably develop gain-of-function CDK-TCIPs to rewire seven different clinical-stage kinase inhibitors of CDK9, CDK12, and CDK13 into context-specific inducers of transcription. Collectively, our CDK-TCIP approach highlights a new strategy that redirects rather than inhibits kinase activity, accessing new therapeutic space.

**Figure 1.**
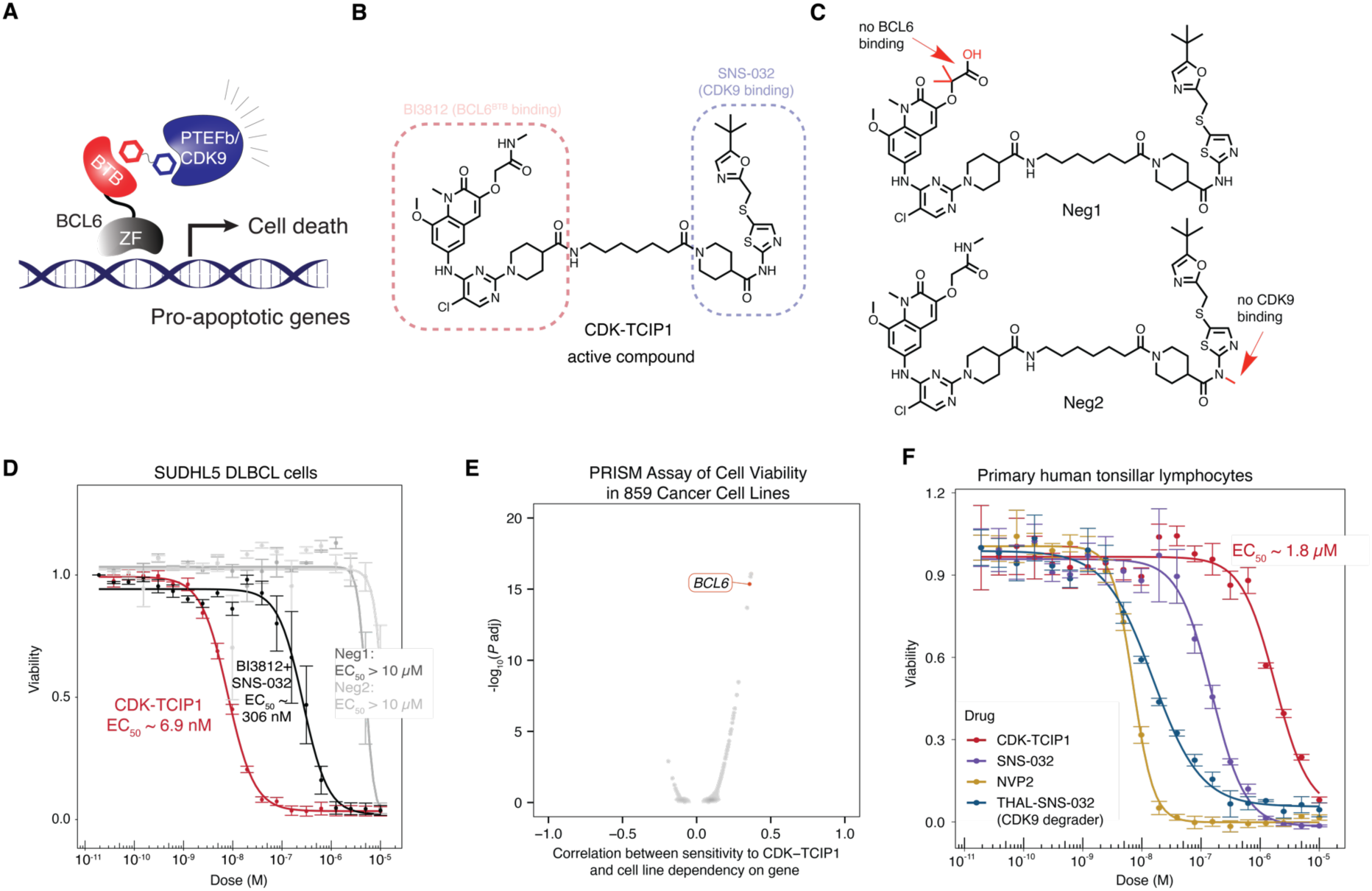
Development of Transcriptional Chemical Inducers of Proximity to recruit Transcriptional Kinases (CDK-TCIPs). **A.** Schematic of a CDK-TCIP targeting BCL6-regulated loci. **B.** Structure of lead compound **CDK-TCIP1**. **C.** Structures of negative controls containing small chemical modifications abolishing BCL6 binding (**Neg1**) or CDK9 binding (**Neg2**). **D.** Cell killing potencies of CDK-TCIP1 compared to negative controls and the additive effect of both BCL6 and CDK9 inhibitors combined at 72 h in SUDHL5 cells; mean±s.e., n=3 biological replicates. **E.** Correlation between genetic dependencies as measured by CRISPR knockout and sensitivity to CDK-TCIP1 among 859 cancer cell lines in PRISM (*56*). **F.** Toxicity of **CDK- TCIP1** compared to CDK9 inhibitors or degraders in primary human tonsillar lymphocytes at 72 hours; mean±s.d., n=2 biological repeats for **CDK-TCIP1** and NVP2, n=1 with 3 technical repeats for SNS-032 and THAL-SNS-032.

## RESULTS

### CDK-TCIPs Exhibit BCL6-Specific Cell Killing

Reasoning that the local proximity of the P-TEFb complex would be sufficient for activation of BCL6-target genes (Fig. 1A), we synthesized a library of bivalent molecules where inhibitors of CDK9 and ligands of the BCL6^BTB^ were connected via linkers of varying length and chemical composition (Supplemental Fig. 1A, Supplemental Table 1). These studies nominated a lead compound, **CDK-TCIP1**, constructed using CDK9 inhibitor SNS-032 (*8, 21*) and BCL6^BTB^- domain ligand BI3812 (*16*) (Fig. 1B). To evaluate the necessity for both binding moieties of **CDK-TCIP1** for biological activity, we synthesized two negative control compounds containing minor chemical modifications to **CDK-TCIP1** that abolish binding to one of either BCL6 (*16*) (**Neg1**) or CDK9 (**Neg2**) (Fig. 1C). **Neg1** and **Neg2** remain comparably strong cell-permeable binders of either CDK9 or BCL6, respectively, as measured by probe-displacement assays inside living cells using bioluminescence resonance energy transfer (nanoBRET(*22*)) between tagged, full-length CDK9/CycT and BCL6 constructs (Supplemental Fig. 2A,B).

In DLBCL cell lines dependent on high BCL6 expression, such as SUDHL5 (BCL6 transcripts/million = 301 (*23*)), **CDK-TCIP1** had a potent cell-killing effect with an EC_50_ of 6.9 nM in a 72-hour cell viability assay (Fig. 1D). The cytotoxicity of **CDK-TCIP1** was ∼50-fold greater than the additive effect of co-treatment of the CDK9 and BCL6 parental inhibitors and >10,000- fold greater than negative control co-treatment (Fig. 1D). In an analysis of 859 different cancer cell lines, **CDK-TCIP1** was uniquely potent in DLBCL lines dependent on high levels of BCL6 expression, indicating a requirement of BCL6 for **CDK-TCIP1** potency (Fig. 1E, Supplemental Fig. 3A,B). **CDK-TCIP1** was more specific to BCL6-expressing DLBCL cells than a previously reported BCL6-directed bivalent compound designed using inhibitors of BRD4 (Supplemental Fig. 3C) (*19*). In primary T and B lymphocytes taken from human tonsils, which have some of the highest levels of BCL6 of normal tissues, **CDK-TCIP1** was 10,000-fold less cytotoxic than optimized CDK9 degraders and inhibitors, which broadly inhibit transcription (Fig. 1F). Our results indicate that the CDK-TCIP strategy delivers cancer-cell-specific effects while overcoming the on-target toxicity associated with inhibition or degradation of essential kinases, potentially enabling a therapeutic window.

### CDK-TCIP1 Activity Depends on Ternary Complex Formation

Titration of BCL6^BTB^ inhibitor BI3812 or CDK9 inhibitor SNS-032 against constant, lethal CDK- TCIP1 doses impaired the cell killing effect, suggesting a requirement for concomitant engagement of CDK9 and BCL6 for the observed cytotoxicity (Supplemental Fig. 2C). We hence hypothesized that ternary complex formation between CDK9 and BCL6 is necessary for cell killing through induction of BCL6-target genes. A cascade of three critical assays was established to iteratively design and assess CDK-TCIPs by measuring (i) ternary complex formation using purified BCL6^BTB^ and full-length CDK9/CCNT1 in a time-resolved fluorescence resonance energy transfer (TR-FRET) experiment, (ii) induction of BCL6-controlled GFP reporter gene expression, and (iii) viability of high BCL6-expressing DLBCL cells (Supplemental Fig. 4A). Bivalent molecules synthesized based on CDK9 inhibitor SNS-032 exhibited the most consistent structure activity-relationship between ternary complex formation *in vitro* and cell killing potency (Fig. 2A). As expected, compounds that induced higher affinity ternary complexes *in vitro* and in cells exhibited greater induction of BCL6-controlled GFP and more potent cytotoxicity in SUDHL5 cells (Fig. 2A, Supplemental Table 1, Supplemental Figs. 4B,C).

**Figure 2.**
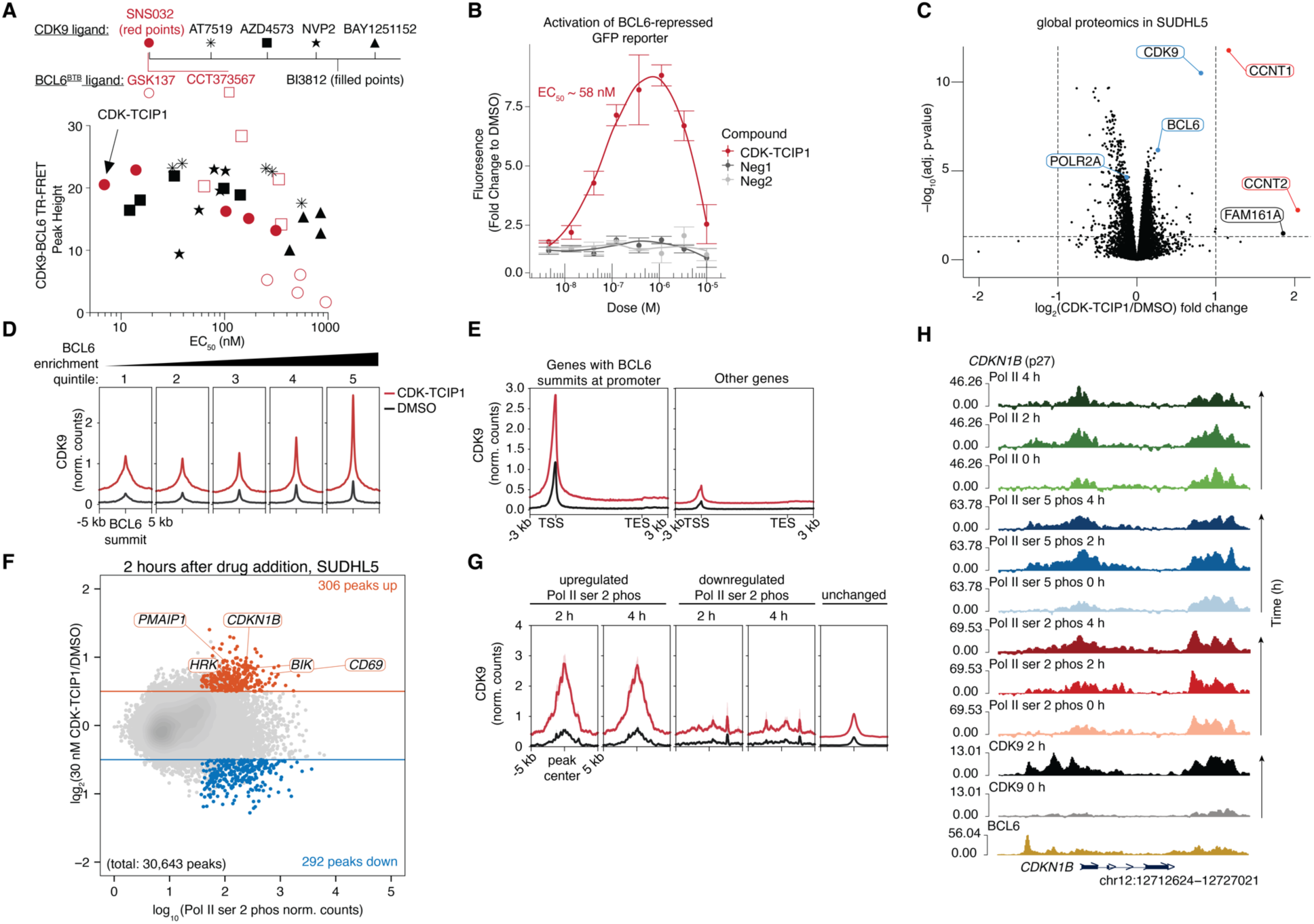
CDK-TCIP1 functions by ternary complex formation and relocalization of CDK9 activity to BCL6 on chromatin. **A.** Correlation of ternary complex formation with cell killing potency (72 h in SUDHL5 cells) for CDK-TCIPs constructed from diverse CDK9 and BCL6 inhibitors. Dots represent mean of n=3 technical replicates for TR-FRET. **B**. Activation of BCL6- repressed GFP reporter construct integrated into KARPAS422 cells after compound treatment for 24 h; mean±s.d., n=3 biological replicates. **C**. Whole-cell global proteomics in SUDHL5 cells after 30 nM **CDK-TCIP1** treatment for 2 h; 4 biological replicates, p-values computed by moderated t-test and adjusted by Benjamini-Hochberg, significance cutoffs at |log_2_(fold change)|≥1 and p_adj_≤0.05. **D.** Recruitment of CDK9 to BCL6 summits genome-wide after 30 nM **CDK-TCIP1** treatment for 2 h in SUDHL5 cells. **E.** Recruitment of CDK9 to genes with BCL6 summits at their promoters. **F.** Changes in local Pol II ser 2 phosphorylation as measured by ChIP-seq after 2 hours 30 nM **CDK-TCIP1** addition in SUDHL5 cells; colored: p_adj_≤0.05 and |log_2_(Drug/DMSO)|≥0.5), n=2 biological replicates; P-values computed by two-sided Wald test and adjusted for multiple comparisons by Benjamini-Hochberg. **G.** CDK9 recruitment correlates with induced Pol II ser 2 phos peaks. **E.** CDK9 is robustly recruited to the known BCL6 target, *CDKN1B*, concomitant with induction of Pol II ser 2 phos across the gene body. **H.** CDK9 is robustly recruited to the known BCL6 target *CDKN1B*, concomitant with induction of Pol II ser 2 phos across the gene body. In **D, E, G, H** metaprofiles and tracks for CDK9 measured with abcam ab239364 antibody shown with two biological replicates merged, spike-in-normalized, and input-subtracted. For Pol II ser 2 phos and Pol II ser 5 phos, two biological replicates merged, sequence-depth normalized and input-subtracted. For Pol II, three biological replicates merged, sequence-depth normalized, and input-subtracted. BCL6 track from(*57*).

The most potent molecule, **CDK-TCIP1**, formed a ternary complex between BCL6^BTB^ and CDK9 with an EC_50_ of 11 nM (Supplemental Fig. 5A) and activated BCL6-controlled GFP expression in DLBCL cells with a comparable EC_50_ of 58 nM (Fig. 2B). Inside HEK293T cells, which do not contain endogenous BCL6, **CDK-TCIP1** forms a ternary complex between overexpressed full- length CDK9 and full-length BCL6 constructs with a EC_50_ of 22 nM as measured by a nanoBRET assay (Supplemental Fig. 5B). In almost all assays we observed a hook effect characteristic of the saturation behavior of bivalent molecules at high concentrations (*6*). **Neg1** and **Neg2** had negligible effects in the transcriptional activation reporter and ternary complex formation assays at comparable doses, demonstrating the requirement for dual binding of CDK9 and BCL6. Collectively, these studies support the necessity of formation of a transcription- competent ternary complex that is critical for **CDK-TCIP1**-mediated DLBCL cell killing.

To evaluate the direct effect on endogenous proteins with CDK-TCIP1 in an unbiased manner, we performed global proteome profiling by treating the DLBCL cell line SUDHL5 with 30 nM **CDK-TCIP1**, **Neg1**, and **Neg2** for 2 h followed by digestion and LC-MS/MS analysis via diaPASEF (Methods), which quantified 169,000 peptides and 8,1000 unique proteins on average. Unlike for **Neg1** or **Neg2** treatment, **CDK-TCIP1** uniquely increased the abundance of cyclin T1 (CCNT1; >2-fold) and T2 (CCNT2; >2-fold), CDK9 (1.8-fold), and BCL6 (1.2-fold) with negligible changes to other proteins (>2-fold) above the statistical cutoff (adj. p-value of 0.05; Fig. 2C, Supplemental Fig. 6A,B,C, Supplemental Table 3). Given the short treatment time and unchanged mRNA abundance of these highlighted proteins (see RNA-sequencing in Fig. 3), these changes in abundance are unlikely to occur through induced transcription. Instead, the data supports thermodynamic stabilization or protection from proteasomal degradation of the BCL6-P-TEFb complex by **CDK-TCIP1**.

**Figure 3.**
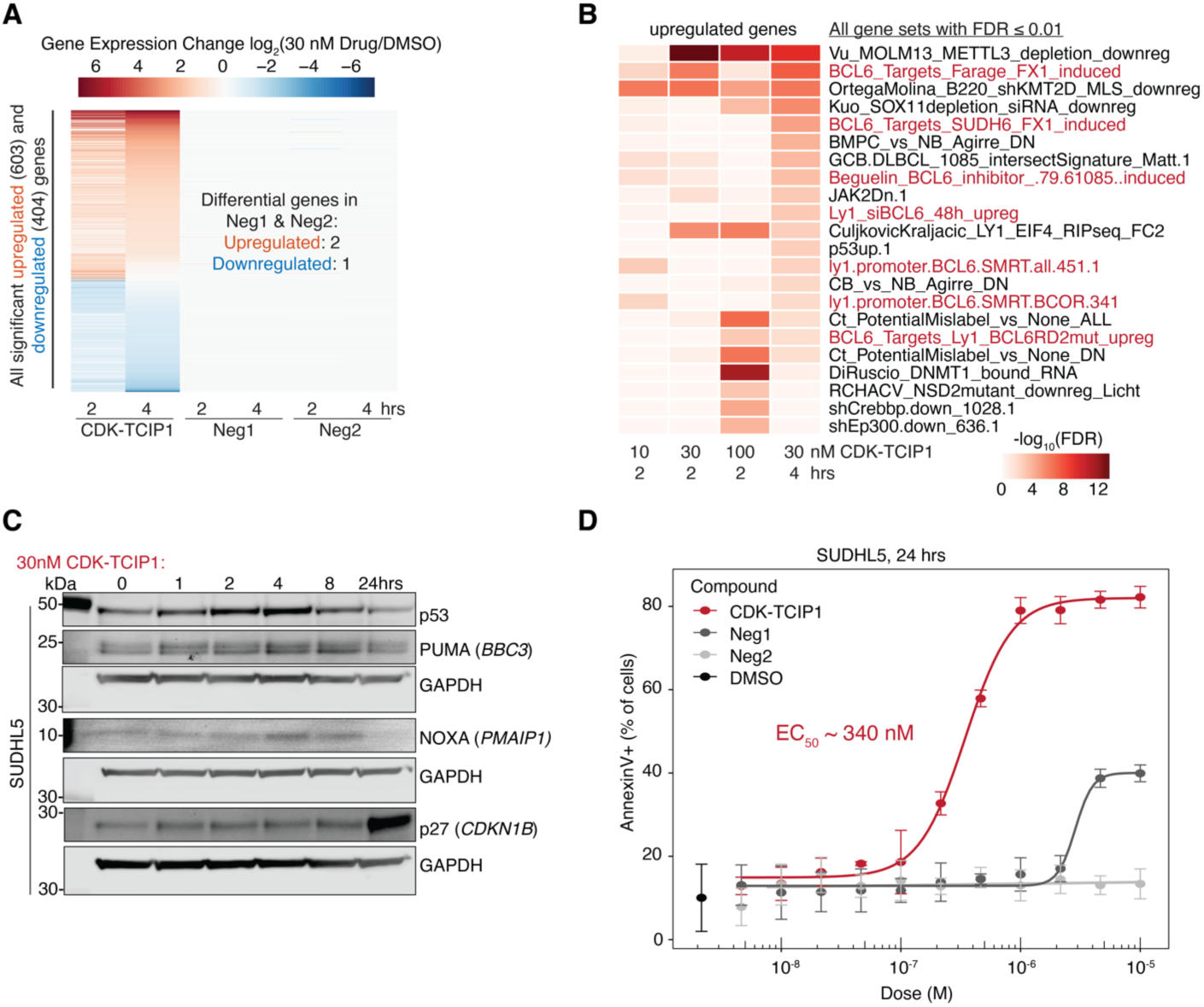
Activation of apoptotic signaling. **A.** Time-dependent changes in gene expression after 30 nM **CDK-TCIP1** addition in SUDHL5 cells compared to controls. Significance cutoffs p_adj_≤0.05 and |log_2_(fold change)|≥1), n=3-4 biological repeats; P-values computed by two-sided Wald test and adjusted for multiple comparisons by Benjamini-Hochberg. **B.** Gene set enrichment analysis of upregulated genes (p_adj_≤0.05, log_2_(fold change)≥0.58). P-values computed by hypergeometric test and adjusted for multiple comparisons by Benjamini- Hochberg. **C.** Time-dependent changes in apoptotic protein activation, p53 signaling, and downstream proteins after addition of 30 nM **CDK-TCIP1** in SUDHL5 cells, representative of 3 biological replicates. **D.** Dose-dependent increase in annexin V staining in SUDHL5 cells after 24 h of drug, n=3 biological replicates, mean±s.d.

### Induced Proximity of CDK9 is Sufficient to Induce Transcription

To assess the functional importance of recruited CDK9 catalytic activity, we constructed a synthetic CIP system by overexpressing FKBP^F36V^-tagged wild-type (WT) or mutant CDK9 in our BCL6-controlled-GFP reporter cell line (Supplemental Fig. 7A,C). We then synthesized a cell- permeable bivalent compound **RCS-03-207**, which linked the <u>s</u>ynthetic ligand of <u>F</u>KBP12 (SLF) with the BCL6^BTB^-domain binder BI3812 (Supplemental Fig. 7B,D). In this model system, **RCS- 03-207** treatment produced a dose-dependent increase in GFP for fused constructs containing WT but not for catalytically dead D167N (*24*) mutant CDK9 (Supplemental Fig. 7E). Overexpression of the WT CDK9-FKBP construct further potentiated GFP transcription following **CDK-TCIP1** treatment with a characteristic hook effect relative to endogenous CDK9 levels (Supplemental Fig. 7F).

Recruitment of functionally active CDK9 is thus likely required for CDK-TCIP-dependent transcription activation of BCL6 gene loci. Given a diffusion-limited rate constant of ∼ 10^8^ M^-1^s^-1^ and the ternary binding apparent EC_50_ of **CDK-TCIP1** of ∼22 nM inside the cell (Supplemental Fig. 5B), the off-rate of ternary complex formation is approximately 2.2 s^-1^, similar to rates of ADP release from kinase active sites (*25–27*). Hence the **CDK-TCIP1** ternary complex should fall apart frequently enough (once every ∼0.45 seconds) to permit catalytic activity at BCL6-bound loci. Therefore, we hypothesize a potential “catch-and-release” mechanism where **CDK- TCIP1** recruits and subsequently releases catalytically active CDK9 to BCL6 loci for transcriptional activation of BCL6-repressed genes (Supplemental Fig. 7G).

### Relocalization of CDK9 Activity to BCL6 Targets on Chromatin

To define the direct effects of **CDK-TCIP1** on CDK9 localization and activity on chromatin, we conducted chromatin immunoprecipitation followed by next-generation sequencing (ChIP-seq) of CDK9 in DLBCL cells treated with 30 nM **CDK-TCIP1** at an early time point (2 h) using spiked *D. melanogaster* chromatin to enable normalization and quantification of absolute changes in CDK9 levels. At this concentration, only a fraction of CDK9 or BCL6 is engaged, as evidenced by the nanoBRET probe displacement assays in Supplemental Fig. 2A,B. Approximately 10,000 CDK9 peaks were reconstructed in each cell line with a strong correlation between biological replicates and the majority of the variance driven by CDK-TCIP1 treatment (Supplemental Fig. 8A,B,C). CDK-TCIP caused rapid and robust recruitment of CDK to BCL6 binding sites (Methods) on chromatin in both SUDHL5 and KARPAS422 cells, as detected with two different anti-CDK9 antibodies (Fig. 2D, Supplemental Fig. 8D,E). The increase in local CDK9 levels correlated with increasing BCL6 enrichment, corroborating BCL6-dependent recruitment on chromatin. Genes with high-confidence BCL6 binding summits within 3 kilobases (kb) of their transcription start sites (TSS) at their promoters showed a 3-fold increase in CDK9 at both the promoter and across the gene body, while other genes had negligible changes over background (Fig. 2E, Supplemental Fig. 8F,G). CDK9 was recruited to both promoters and enhancers with BCL6 summits; regions without BCL6 showed modest increases or no changes in CDK9 occupancy (Supplemental Fig. 8H). Overall, five times as many peaks increased (604, 5.1% of all 11,716 peaks) in CDK9 binding as decreased (142, 1.2% of all 11,716 peaks) as calculated by differential peak analyses using relative log expression (RLE) normalization (Supplemental Fig. 8I), consistent with global proteomics results indicating stabilization of CDK9 protein in Fig. 2C. Taken together, the analysis indicates that CDK9 is rapidly and specifically recruited to BCL6-bound chromatin.

To define the immediate consequences of relocalizing CDK9 to BCL6-bound loci on chromatin we characterized changes in Pol II and its serine phosphorylations at 2 h and 4 h after **CDK- TCIP1** addition (30 nM, Supplemental Fig. 9A). A select number of Pol II ser 2 phos peaks were induced, primarily at promoters (306 peaks up, p_adj_ ≤ 0.05 and fold change ≥ 1.4, Fig. 2F, Supplemental Fig. 9B). Peaks that increased in Pol II ser 2 phos at 2 h and at 4 h were highly enriched for CDK9 binding, while those that decreased or remained unchanged had negligible changes in CDK9 levels (Fig. 2G). Among the most induced Pol II ser 2 phos sites were known BCL6-repressed cell death and tumor suppressor genes, such as *PMAIP1*/NOXA, *BIK*, *HRK/*HARAKIRI, and *CDKN2B*/p27 (Figs. 2F,H and Supplemental Fig. 8H). Induced peaks were significantly enriched at sites that had been previously identified as bound by components of the BCL6:Polycomb Repressive Complex 1 (PRC1) epigenetic repressor complex (BCL6, BCOR, KDM2B (*28, 29*)) as analyzed in public ChIP-seq datasets from over 6,500 blood cell lines (Supplemental Fig. 8C), consistent with the observation that CDK9 is recruited to BCL6 sites in Fig. 2D and E. A peak-to-gene analysis using GREAT (*30*) confirmed that these induced peaks occurred near genes related to pathways known to be repressed by BCL6, including programmed cell death and cell cycle arrest biological processes (FDR < 0.01) (Supplemental Fig. 9D). Loci that showed decreased Pol II ser 2 phosphorylation (292 peaks down, p_adj_ ≤ 0.05 and fold change ≤ -1.4 Fig. 2F) corresponded with some known BCL6-regulated pathways such as the humoral immune responses (Supplemental Fig. 9D), but had lower enrichment of BCL6 (Supplemental Fig. 9E) and less overlap with BCL6 peaks (Supplemental Fig. 9F), as compared to induced Pol II ser 2 peaks, indicating that decreased peaks may be off-target or indirect effects. These results provide evidence that CDK-TCIPs selectively recruits CDK9 and its activity to BCL6-repressed genetic loci.

Consistent with a mechanism of locus-specific activity, 4 h **CDK-TCIP1** treatment did not globally alter Pol II ser 2 phosphorylation or BCL6, CDK9, or Pol II protein levels by immunoblotting until 10-fold greater concentrations than that those used for these studies (Supplemental Fig. 10A), which was consistent with global proteome profiling (Fig. 1G) and nanoBRET displacement data (Supplemental Fig. 2A,B). There was little change in total Pol II at either genes with BCL6 sites at their transcription start sites or elsewhere (Supplemental Fig. 9G). Our data supports a gain-of-function mechanism where **CDK-TCIP1** relocalizes a fraction of cellular CDK9 to BCL6-bound loci.

### Activation of BCL6-target Gene Expression and Apoptotic Signaling

Transcriptome analysis by RNA-sequencing (RNA-seq) after short treatments (2 h and 4 h) in SUDHL5 cells of **CDK-TCIP1** (10, 30, or 100 nM) identified a set of 603 genes that were significantly induced (p_adj_ ≤ 0.05 and fold change ≥ 1) and fewer (404) that decreased in expression (Fig. 3A). Induced genes were enriched for multiple annotated BCL6-target gene sets (Fig. 3B). Gene expression changes and enrichment for BCL6 signaling were dose-dependent (Supplemental Fig. 11A).

**CDK-TCIP1** induced known pro-apoptotic, BCL6-target genes in mRNA and protein expression immediately after treatment, including *BBC3/*PUMA, *PMAIP1*/NOXA, and *CDKN1B*/p27 (Figs. 3C, Supplemental Fig. 11B). These genes also displayed recruitment of CDK9 and increased Pol II ser 2 phosphorylation as shown in Fig. 2F. PUMA/*BBC3* (<u>p</u>53 <u>u</u>p-regulated <u>m</u>odulator of <u>a</u>poptosis) is of special importance as it rapidly causes the release of apoptogenic mitochondrial proteins via interaction with BCL2 and BCL-XL(*31–33*). p53 signaling is known to be downstream of BCL6 as not only a direct gene target but also on multiple post-transcriptional levels (*13, 14, 34*). We observed an increase of p53 at the protein level as early as 1 h after compound treatment by immunoblotting (Fig. 3C). Intriguingly, in KARPAS422 DLBCL cells, which exhibit comparable levels of BCL6 but harbor biallelic truncations of *TP53* that ablate its tetramerization and stable DNA-binding ability (K319* (*35, 36*)), p53 protein levels also increased, yet PUMA was not induced and **CDK-TCIP1** exhibited 10-fold lower killing potency (Supplemental Fig. 11D,E, left). There was no difference in antiproliferative effects of combined BCL6 and CDK9 inhibition between *TP53^WT^*and *TP53^mut/mut^* mutant lines (Supplemental Fig. 11E, right), indicating that sensitivity to p53 status is an emergent feature of **CDK-TCIP1**. As expected, genes induced after 4 h of **CDK-TCIP1** treatment were enriched for p53 signaling (Fig. 3B, Supplemental Fig.11F) and p53 binding in their promoters (Supplemental Fig. 11G). No DNA damage response was observed as measured by γH2AX in CDK-TCIP1-treated SUDHL5 cells (Supplemental Fig. 11H).

Annexin V staining after 24 h showed that **CDK-TCIP1** induced dose-dependent apoptosis in SUDHL5 cells with an EC_50_ of 340 nM (Fig. 3D). In contrast, negative controls **Neg1** and **Neg2** exhibited 10- to 1,000-fold lower potency in this assay, consistent with their negligible effects on gene induction (Fig. 3D). Taken together our data shows that CDK-TCIP1 activates p53 signaling as well as other death pathways repressed by BCL6, consistent with the rational optimization of the compound to induce BCL6-repressed transcription in Figs. 1 and 2.

An investigation into transcripts decreasing after 30 nM **CDK-TCIP1** treatment revealed that these were enriched for germinal center (GC) B-cell programs, for which BCL6 is a master regulator (Supplemental Fig. 12A). *BCL6* itself, which contains a BCL6 binding site in its first intron and is auto-regulated(*37*), showed a modest decrease in Pol II ser 2 phosphorylation and increase in CDK9, Pol II, and Pol II ser 5 phosphorylation at its promoter, suggesting initiated but paused Pol II, (Supplemental Fig. 12B). *BCL6* expression decreased by ∼12% (p_adj_ = 0.003) at 4 hours after 30 nM CDK-TCIP1, although little change in BCL6 protein was observed (Fig. 2C and Supplemental Fig. 10A). Calculation of the pausing ratio of each gene in cells before **CDK-TCIP1** treatment, defined as the ratio between Pol II ser 5 phos at the transcription start site and Pol II ser 2 phos at transcription end site (Methods), indicated that genes decreasing in expression were more likely to be already paused (higher ratio) than induced or unchanged genes (Supplemental Fig. 12C). CDK-TCIPs may locally inhibit elongation and transcription at these loci, which are also BCL6-target genes.

### Rational Chemical Optimization of CDK-TCIPs

We used our design cascade (Supplemental Fig. 4A) in a medicinal chemistry campaign to access CDK-TCIPs with improved physicochemical properties and increased potency as compared to **CDK-TCIP1**, which exhibited poor pharmacokinetic properties that precluded further preclinical drug development (Supplemental Fig. 13A, B). Inspired by optimization studies of protein degrader drug candidates (*38, 39*), we focused our synthetic efforts on introducing rigid and amine-containing linkers to replace the flexible *n*-hexyl (C_6_) chain in **CDK- TCIP1** (Supplemental Fig. 14A). These studies nominated several compounds that exhibit enhanced ternary complex formation, activation of BCL6-controlled transcription, and cell killing (Supplemental Fig. 14A-F).

**CDK-TCIP2**, which incorporates a rigid 3,9-diazaspiro[5.5]undecyl linker motif, emerged as a new lead compound that killed SUDHL5 cells at sub-nanomolar concentrations (EC_50_ = 0.9 nM, Supplemental Fig. 14C). In mice, **CDK-TCIP2** exhibited improved metabolic stability and exposure levels over **CDK-TCIP1**, permissive for additional *in vivo* studies (AUC = 1.4 μM·hrs, Supplemental Fig. 15A,B). Drug accumulation was ∼100-fold above cellular EC_50_s predominantly in the liver, kidney, and spleen (Supplemental Fig. 15C). Our studies demonstrate that CDK-TCIPs are amenable to rational optimization of potency and physicochemical properties through medicinal chemistry and underline the utility of our novel TCIP design cascade.

### Specific Ablation of Germinal Center Response in Immunized Mice

In normal physiology, BCL6 is the master regulator of the germinal center (GC) reaction that is required for T-cell dependent antibody affinity maturation (*40, 41*). The BCL6 protein is highly expressed in the GC B cells and T follicular helper (Tfh) cells (*42*), and is down-regulated upon GC exit towards memory B-cells or plasma cells (*40*). The GC B cell is thus the counterpart to follicular lymphoma, Burkitt’s lymphoma, and the GC B-cell like (GCB) subtype of DLBCL which collectively make up more than 40% of B cell malignancies(*43*). Given the BCL6-targeted, specific effects on transcription and apoptosis in DLBCL cell lines, we hypothesized that **CDK- TCIP2** would specifically suppress the proliferation of GC B cells in a mouse immunization model.

Two days following immunization of C57Bl/6 mice with the T-cell dependent antigen sheep red blood cells (SRBCs) to stimulate the GC reaction, we began administering **CDK-TCIP2** at 5 mg/kg once daily, 10 mg/kg once daily, 5 mg/kg twice daily, or with vehicle twice daily by intraperitoneal injection (i.p.). Ten days post-immunization, at the peak of the GC reaction, the mice were euthanized, and the spleens harvested for flow cytometry of GC B-cell populations. GCs were robustly induced in vehicle treated mice and the percentage of splenic GC B cells (B220^+^Fas^+^GL7^+^) was significantly reduced in **CDK-TCIP2**-treated mice (Fig. 4B). The reduction was dose-dependent and highest in the 5 mg/kg twice-daily (BID) dosing regimen likely due to the half-life (T_1/2_ ∼ 3.2 h, Supplemental Fig. 15A,B) of the molecule. The total frequency of B220+ B-cells was only modestly reduced, commensurate with the loss GC B-cells, supporting on-target specificity for the BCL6+ GC B-cell compartment (Fig. 4C). A modest and statistically insignificant increase in memory B-cells was noted, likely reflecting the shunting of B-cells out of the GC reaction following its establishment in the first 2 days post-immunization before treatment (Fig. 4D). The compound was well-tolerated with no changes in body weight or other adverse effects noticed over the 8-day dosing period (Fig. 4E). This data is consistent with studies that showed that although *BCL6^-/-^* mice die within weeks of birth from a lethal inflammatory reaction(*44*), mice carrying mutations in the regions of the BCL6 protein that bind its co-repressors have normal lifespans in controlled conditions but exhibit a defect in their ability to form GCs (*45, 46*). Taken together, the results indicate that **CDK-TCIP2** specifically inhibits immunological processes ordinarily facilitated by the repressive function of BCL6. These experiments also suggest that CDK-TCIPs may be useful in rare autoimmune conditions such as systemic lupus erythematosus (SLE) characterized by elevated autoantibodies acquired during an elevated germinal center response (*47*).

**Figure 4.**
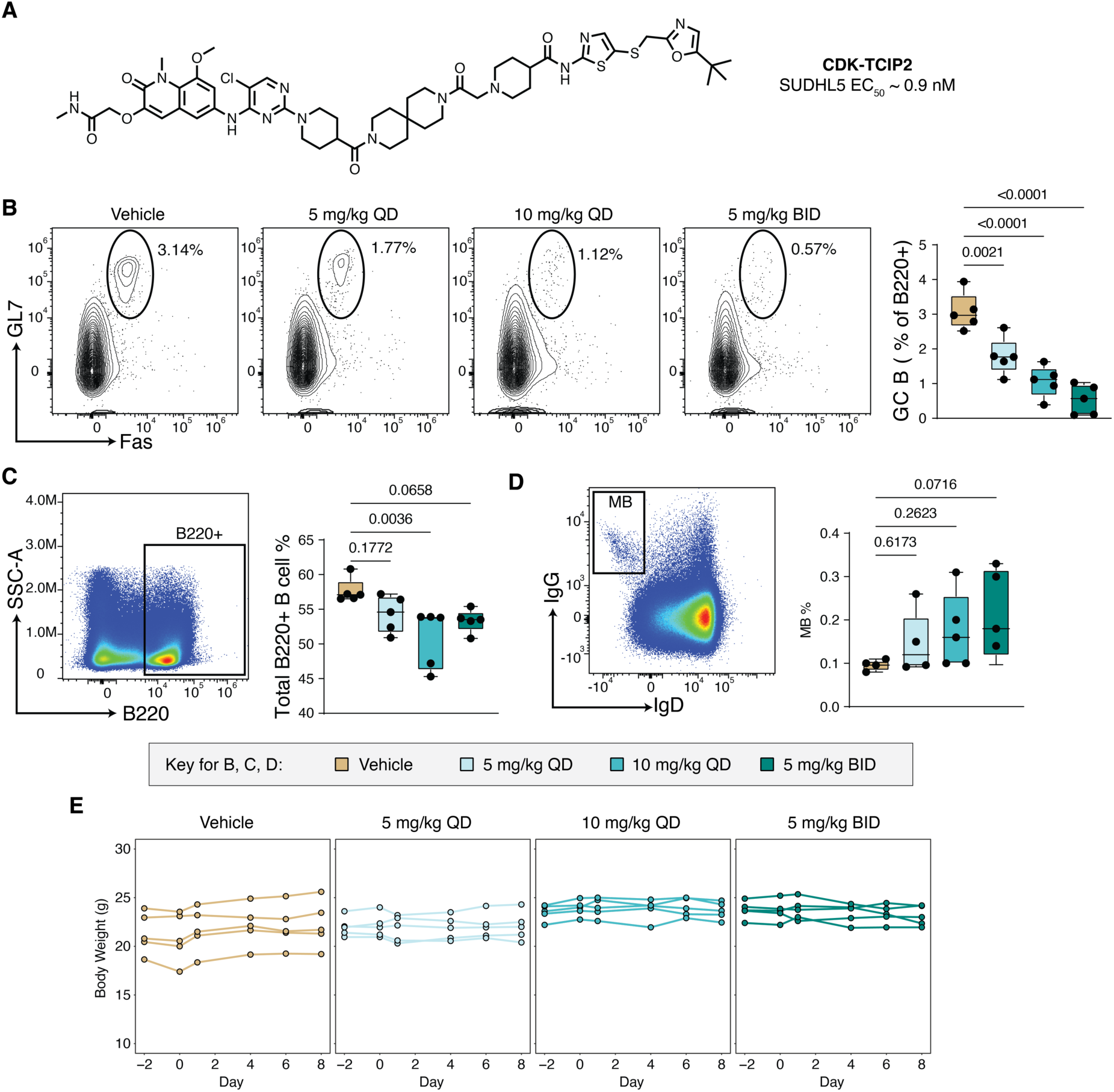
Ablation of germinal center (GC) B cells in immunized mice. **A.** Structure of **CDK- TCIP2**. **B.** Dose-dependent change in percentage of splenic GC (B220^+^Fas^+^GL7^+^) B cells. **C.** Change in total frequency of splenic B220^+^ B cells. **D.** Change in total frequency of splenic memory B cells (MB). In **B,C,D:** representative flow cytometry on left and quantification on right from 5 biological replicates (5 different mice) for each condition, median and range shown; p- values computed by unpaired Students’ t-test. QD: once daily; BID: twice daily. Vehicle was given twice daily. **E.** Body weight changes over time.

### Generalization of the CDK-TCIP Concept to CDK12/13

We explored the predictability of our strategy to redirect kinase activity by developing molecules that recruit the transcriptional kinases CDK12 and CDK13 (Fig. 5A). CDK12 and CDK13, similar to CDK9, phosphorylate Pol II ser 2 and contribute to transcriptional elongation (*48, 49*). In some cancers, CDK12 and CDK13 are deregulated by amplification or mutations (*50, 51*). We synthesized **CDK-TCIP3**, a molecule that links a potent and selective CDK12/13 inhibitor (*52*) with the BCL6^BTB^ ligand BI3812 (Fig. 5B). **CDK-TCIP3** retained 5-fold binding selectivity for CD12/13 over CDK9 (CDK13/CycK IC_50_ ∼ 65 nM, CDK9/CycT1 IC_50_ ∼ 261 nM) and potently induced BCL6-repressed reporter GFP expression, exhibiting the hook effect that is characteristic of CIP function (Fig. 5C). **CDK-TCIP3** exhibited a comparably potent cell killing effect with an EC_50_ of ∼ 609 nM in SUDHL5 cells, more than 10-fold greater than the effect of the CDK12/13 inhibitor and the combination of the CDK12/13 inhibitor and the BCL6 inhibitor BI3812 in these cells (Fig. 5D). These studies suggest that inhibitors of transcriptional kinases beyond CDK9 can be rationally rewired into activators of cell death.

**Figure 5.**
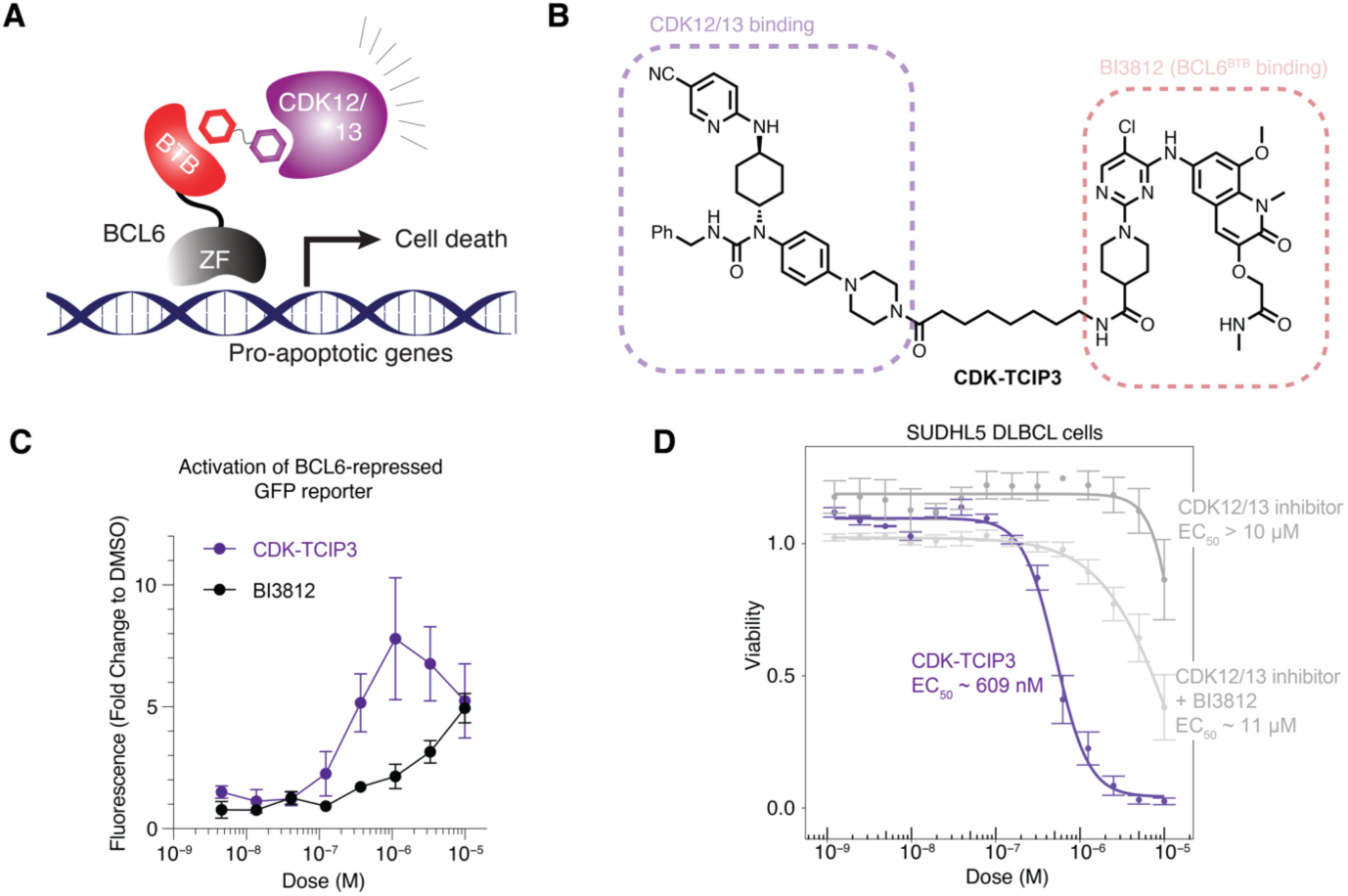
Generalization of the CDK-TCIP concept to CDK12/13. **A.** Schematic of recruiting CDK12/13 to BCL6-bound loci. **B.** Structure of CDK-TCIP3. **C.** Activation of BCL6-repressed GFP reporter constructs after 24 hours compound addition to lymphoma cells; mean±s.d., n=4 biological replicates. **D.** Comparison of cell killing potency in DLBCL cells at 72 hours between CDK-TCIP3 and additive effect of both BCL6 and CDK12/13 inhibitors; n=3 biological replicates, mean±s.e.

## DISCUSSION

Kinases control and amplify signals from the cell membrane to the nucleus and are commonly deregulated in disease. Oncogenic kinases have been subject to extensive drug discovery efforts, with many potent and specific inhibitors of kinase catalytic activity developed. Almost 10% of the human kinome has been targeted with potent and specific inhibitors, the vast majority of which are ATP-competitive, reversible small-molecules (*1*).

As transcriptional deregulation is a hallmark of malignant transformation, many inhibitors of transcriptional kinases (and other nuclear transcriptional regulators) have been optimized for potency, specificity, and physiochemical properties. However, clinical applications of the inhibitors are constrained by on-target, mechanism-based toxicities, largely due to the need for near-complete inhibition of kinase activity (*53*). Recent efforts have also sought to convert inhibitors into selective protein degraders by chemical linking to E3 ligase-binding moieties (*8*). However, both inhibitors and protein degraders fundamentally aim to phenocopy genetic loss of aberrant protein function.

Here we developed CDK-TCIPs, a chemically induced proximity-based gain-of-function strategy to activate apoptosis by re-localizing transcriptional kinase activity to pro-apoptotic gene loci bound by an overexpressed, oncogenic transcription factor, BCL6. These bivalent molecules function by borrowing a fraction of elongation kinase activity, as evidenced by the locus-specific recruitment of CDK9 and induction of Pol II ser 2 phosphorylation on chromatin. CDK-TCIPs induce tumor suppressors and pro-apoptotic proteins, such as CDKN1B/p27, NOXA, and p53 signaling, at low nanomolar drug concentrations that do not inhibit global kinase activity. The ensuing killing effect is exquisitely lineage-specific to only those cells driven by BCL6 in both malignant cell lines and mouse models of B cell development. This contrasts with CDK9 inhibitors and degraders that are more broadly cytotoxic to both normal and malignant cells. Our approach thus offers an alternative to conventional chemotherapeutic approaches that phenocopy genetic or enzymatic loss-of-function effects. We note that bivalent molecules have previously been developed to redirect the catalytic activity of kinases to new substrates, revealing the roles of proximity and orientation in kinase activation (*54, 55*). However, to the best of our knowledge, our work represents the first example of rewiring endogenous kinase activity to drive a therapeutically relevant phenotype *in vivo*.

The assay cascade employed for rational improvement of CDK-TCIP physicochemical properties further served to extend the strategy beyond CDK9 to other kinases such as CDK12/13. Our work indicates that the many targeted and highly optimized kinase inhibitors can be leveraged for the design of therapeutic CIPs that borrow and re-localize kinase activity.

These molecules would be designed to turn on new cellular signaling pathways rather than to silence aberrant catalytic activity. The potential of gain-of-function kinase CIPs extends beyond the induction of programmed cell death pathways to, for example, the activation of differentiation or pluripotency for applications in regenerative medicine or the attenuation of inflammatory signaling.

## ONE SENTENCE SUMMARY

Development of chemical inducers of proximity that rewire kinase inhibitors into activators of programmed cell death.

## METHODS

### Cell Culture

Lymphoma and leukemia cells were cultured in RPMI-1640 (ATCC 30-2001) + 10% FBS with antibiotics (100X PenStrep GIBCO #15140122). KARPAS422 cells were obtained from Sigma (#06101702) and SUDHL5 were obtained from the American Tissue Culture Collection (ATCC). Primary human tonsillar lymphocytes were a kind gift from M. M. Davis. Cells were routinely checked for mycoplasma and immediately checked upon suspicion. No cultures tested positive.

### Cell Viability Measurements

30,000 cells were plated in 100 µL media per well of a 96 well plate and treated with drug for indicated times and doses. A resazurin-based indicator of cell health (PrestoBlue, ThermoFisher #P50200) was added for 1.5 hours after which the fluorescence ratio at 560/590nm was recorded. The background fluorescence was subtracted and the signal was normalized to DMSO-treated cells. Fit of dose-response curves to data and statistical analysis was performed using drc package in R using the four-parameter log-logistic function.

### PRISM Cell Proliferation Assay

The PRISM cell proliferation assay was carried out as previously described(*58*). Briefly, up to 859 barcoded cell lines in pools of 20-25 were thawed and plated into 384-well plates (1250 cells/well for adherent cells, 2000 cells/well for suspension or mixed suspension/adherent pools). Cells were treated with an 8-point dose curve starting at 10 µM with threefold dilutions in triplicate and incubated for 120 hours, then lysed. Each cell’s barcode was read out by mRNA-based Luminex detection as described previously(*56*) and input to a standardized R pipeline (https://github.com/broadinstitute/prism_data_processing) to generate viability estimates relative to vehicle treatment and fit dose-response curves. The area under the dose-response-curve (AUC), which is correlated with drug potency, was used as a metric of drug potency in a cell line, and correlated with *BCL6* transcripts/million as annotated in the Cancer Cell Line Encyclopedia (CCLE)(*23*).

### Chemical Synthesis

Additional details are provided in the Supplemental Methods section.

### Protein Expression and Purification

Biotinylated BCL6^BTB^-AviTag protein used for TR-FRET assays included BCL6 amino acids 5- 129 with the following mutations: C8Q, C67R, C84N (*59*). These enhance stability but do not affect the affinity for BI3812 or for SMRT. Preparation of this protein is described elsewhere (*16*, *60*). The baculovirus expression vector for CDK9-Cyclin T1 (renamed pNSG162) was a generous gift from Dr. Stefan Knopf and was created as follows: codon-optimized coding sequences for CDK9 (residues 2-230; Uniprot P50750-1) and Cyclin T1 (residues 2-259; Uniprot O60563-1) were synthesized and cloned into pFastBac dual (Sigma). Each open reading frame codes for N-terminal 6-His tags and TEV cleavage sites. Bacmid was generated according to the manufacturer’s instructions, and corresponding baculovirus was propagated for three passages in adherent Sf9 cells (Sigma) grown in Sf900 medium (Thermo Fisher).

Passage 3 virus was used to infect a shaking culture of Sf9 cells, and protein expression was carried out for 72 hours before pelleting the cells by centrifugation. Cell pellets were resuspended in ∼10 ml per liter buffer B50 (20 mM HEPES, pH 7.5; 50 mM NaCl; 10 mM imidazole, pH 8.0; 2 mM beta-mercaptoethanol; 10 % glycerol (*v:v*)), supplemented with protease inhibitors (1 mM PMSF, 1 mM benzamidine, ∼20 μg/ml pepstatin, aprotinin, and leupeptin), and stored at -80 ***°***C.

To purify CDK9-Cyclin T1, cell pellets were thawed in warm water before adjustment of the NaCl concentration to 800 mM. All subsequent steps were carried out at 4 ***°***C or on ice. Cells were lysed by sonication before centrifugation at 16,233 rcf for 1 hr. Clarified lysate was mixed with ∼0.5 ml/L of culture cobalt resin (TaKaRa) for one hour. Beads were washed by centrifugation and subsequently by gravity flow with ∼25 column volumes of buffer D800 (B50 with 800 mM NaCl) followed by ∼10 column volumes of buffer B50. Protein was eluted with 50 ml C50 (B50 with 400 mM imidazole), and the eluate was applied to a 5 ml anion exchange column (Q HP, Cytiva) and eluted via salt gradient (8 CV; B50 to D800). Peak fractions were concentrated by ultrafiltration before application to a 24 ml gel filtration column (S200 increase, Cytiva) charged with GF150 buffer (20 mM Tris-HCl, pH 8.5, 150 mM NaCl, 1 mM TCEP). Peak fractions were again concentrated by ultrafiltration, supplemented with 5% glycerol by volume (final), and aliquoted and frozen at -80 ***°***C. Individual aliquots were thawed and stored for no more than 24 h at 4 ***°***C or on ice before use or disposal.

### HiBiT Tag Quantification

For HiBiT Tag Quantification, 1,000,000 cells were resuspended in 1 mL of RPMI media (10% FBS, 1% PS) as a stock solution. The Nano-Glo HiBiT Lytic Reagent was prepared according to established protocols. 40 µL of each stock solution of cells were plated in a 384 well plate. 20 uL of Nano-Glo HiBiT Lytic Reagent was then added to each well. The plate was shaken on an orbital shake for 10-15 mins at room temperature before luminescence reading on a a PHERAstar FS plate reader (BMG Labtech).

### TR-FRET

Each reaction contained 10 nM CDK9/CCNT1, 100 nM BCL6-BTB-Avi-Biot, 20 nM Streptavidin-FITC (Thermo #SA1001), and 1:400 anti-6xHis terbium antibody (PerkinElmer #61HI2TLF) in 10 uL of buffer containing 20 mM HEPES, 150mM NaCl, 0.1% BSA, 0.1% NP-40, and 1 mM TCEP in a 384well plate. Protein was incubated with drug digitally dispensed (Tecan D300e) for 1 h in the dark room at room temperature before excitation at 337 nm and measurement of emission at 520 nm (FITC) and 490 nm (terbium) with a PHERAstar FS plate reader (BMG Labtech). The ratio of signal at 520 nm to 490 nm was calculated and normalized to DMSO-treated conditions and plotted.

### nanoBRET

For BCL6-SMRT: HEK293T cells were plated at a density of 4×10^5^ cells/mL in 2 mL of DMEM/well in a tissue culture treated, 6-well plate and were allowed to incubate overnight at 37 °C. The next day, each well of cells was transfected with 0.2 μg NanoLuc-BCL6 and 2 μg SMRT(aa1292-1500)-HaloTag using Lipofectamine2000 for a ratio of 1:10 NanoLuc to HaloTag plasmids. The transfected cells were allowed to incubate overnight at 37 °C. The following day, the transfected cells were trypsinized and brought to a concentration of 2×10^5^ cells/mL in Fluorobrite DMEM containing 10% FBS. 40 μL of cells per well were plated in a 384-well plate and treated with 100 nM HaloTag618 Ligand via a Tecan D300e Digital drug dispenser.

Designated wells were not treated with the HaloTag618 Ligand and normalized with DMSO to be used as no acceptor controls for data analysis. The cells were allowed to incubate overnight at 37 °C. The next day, the cells were treated with compounds for testing using a Tecan D300e and incubated at 37 °C for 2 h. After 2 h, 10 μL of 5x NanoBRET NanoGlo Substrate was added to each well and mixed on an orbital shaker at 350RPM for 30 s. Data was then obtained using a PheraStar plate reader measuring luminescence with 450-BP and 610-LP filters.

For CDK9 Kinase Engagement: HEK293T cells were trypsinized and brought to a concentration of 2×10^5^ cells/mL in DMEM. They were transfected with 1.5 μg NanoLuc-CDK9 fusion vector (Promega #NV2871) and 13.5 μg CCNT1 expression vector (Promega #NV2681) using FuGENE 4K transfection reagent (Promega #E5911) at a volumetric ratio of 1:20, DNA/FuGENE mix:Cell stock (2×10^5^ cells/mL). 30 mL of cells were transfected and plated in a 15 cm, tissue culture treated dish. The transfected cells were allowed to incubate for 24 h at 37 °C. Following the 24 h incubation, the transfected HEK293T cells were harvested and brought to a concentration of 2.5×10^5^ cells/mL in Fluorobrite DMEM containing 10% FBS. The cells were then plated in 384-well, white, tissue culture treated, opaque bottom plates with 34 μL plated per well for a total of 8500 cells per well. The Promega Kinase Tracer K-8 solution was then prepared by diluting the stock K-8 tracer with DMSO to make a 100X stock. The 100X K-8 stock was then further diluted using Promega kinase dilution buffer to prepare a 20X stock. The 20X stock was further diluted using Fluorobrite DMEM containing 10% FBS to make a 6.67X stock. 6 μL of the 6.67X K-8 stock was dispensed into each well for a final concentration of 63 nM K-8 tracer in cells. 6 μL of FluorobriteDMEM containing 10% FBS were added to designated wells in place of the K-8 tracer solution to act as no acceptor controls for data analysis. The plate was then treated with compounds using a Tecan D300e digital dispenser and allowed to incubate for 2 h at 37 °C. Following the 2 h incubation, a 3X NanoBRET NanoGlo substrate with extracellular NanoLuc inhibitor solution was made, and 20 μL was dispensed into each well. The plate was mixed for 30 s at 350 RPM on an orbital shaker. Data was then obtained using a PheraStar plate reader measuring luminescence with 450-BP and 610-LP filters.

For all nanoBRET assays, a dose-response curve with 3 technical replicates for each CDK- TCIP carried out, and corrected BRET ratios were calculated according to manufacturer assay protocol (Promega #TM439). Data was fit using a standard four parameter log-logistic function using the R package drc or using GraphPad Prism.

### Flow Cytometry

For annexin V assays, 100,000 cells plated at 1M/mL treated with drug for indicated timepoints and doses were harvested on ice and washed twice in 2.5% FBS/PBS. 2.5 µL of 7-AAD and 2.5 µL of FITC-Annexin V (Biolegend #640922) were added. Cells were incubated for 15 minutes at RT, then immediately measured on a BD Accuri. Gates were drawn based off single-stain and no-stain controls.

### BCL6 Reporter Assay

KARPAS422 cells were lentivirally transduced with a construct containing the reporter. After selection, cells were plated and treated with indicated amount of TCIP1 for 8 hours. Cells were washed in 2.5% FBS/PBS, 1:250 v/v of 7-AAD was added to distinguish live from dead cells, and harvested for flow cytometry on a BD Accuri. Given the polyclonal population after transduction, the area under the curve of the histogram representing FITC signal across all live cells was calculated as an integrative measure of total GFP signal. A GFP-positive gate was drawn off non-transduced cells and the area past the threshold for each sample was calculated and normalized to cells treated with DMSO.

### RNA Extraction, qPCR, and Sequencing Library Preparation

Cells were plated at 1M/mL and harvested in TRIsure (Bioline #38033). RNA was extracted using Direct-zol RNA MicroPrep columns (Zymo #R2062) treated with DNAseI. For sequencing library preparation, polyA-containing transcripts were enriched, prepared into paired-end libraries, and sequenced on an Illumina NovaSeq (Novogene).

### Western Blots

Cells were plated at 1M/mL and treated with drug at indicated timepoints and doses. 2M Cells were harvested on ice in RIPA buffer (50mM Tris-HCl pH 8, 150mM NaCl, 1% NP-40, 0.1% DOC, 1% SDS, protease inhibitor cocktail (homemade), 1mM DTT) and 1:200 benzonase (Sigma #E1014) was added and incubated for 20 mins. After 10 min centrifugation at 14,000g and 4 ^O^C, the supernatant was collected and protein concentration was measured by Bradford. Antibodies used for immunoblots are: BCL6 (Cell Signaling D65C10), CDK9 (Cell Signaling C12F7), Pol II (Santa Cruz 8WG16), TP53 (Santa Cruz DO-1), c-MYC (Cell Signaling D84C12), PUMA (Cell Signaling E2P7G), NOXA (Cell Signaling D8L7U), p27 (Cell Signaling D69C12), CASP8 (Cell Signaling 1C12), GAPDH (Santa Cruz 6C5), Pol II ser 2 phos (Active Motif C1984), yH2AX (abcam ab11174). All antibodies were used at 1:1000 v/v dilutions except GAPDH (1:2000).ImageStudio (Licor) was used for blot imaging.

### Global Proteomic Profiling via LC-MS/MS

#### Sample Preparation

SUDHL5 cells (40M cells in 40 mL RPMI-1640 media supplemented with 10% FBS and 1x pen-strep) were treated with 0.1% DMSO or 30 nM CDK-TCIP1, Neg1, or Neg2 (1000x stocks in DMSO for final 0.1% DMSO concentration) for 2 h before washing cells twice with cold serum-free media and then twice with TBS (50 mM Tris, pH 8.5, and 150 mM NaCl). The cell pellets were then thermally denatured in residual TBS for 5 min at 95 °C and then cooled to room temperature. Fresh Lysis Buffer (8 M urea, 150 mM NaCl, and 100 mM HEPES, pH 8.0, in MS-grade water) was then added and the lysates were homogenized using needle ultrasonication. Samples containing 236 µg protein lysate in final 96 µL of Lysis Buffer using a Bradford protein concentration assay (Bio-Rad 5000006) were then prepared for subsequent reduction and alkylation for 25 min at room temperature by the simultaneous addition of 10X tris(2-carboxyethyl)phosphine hydrochloride (TCEP; Sigma Aldrich C4706) and chloroacetamide (CAM; Sigma Aldrich C0267) prepared in 50 mM HEPES, pH 8.0, in MS-grade water for final concentrations of 10 mM and 40 mM, respectively. The samples were then diluted by the addition of CaCl_2_ (Sigma Aldrich C4901) in 50 mM HEPES, pH 8.0, for final concentrations of 0.9 M urea and 1 mM CaCl_2_. Digestion using 1:100 (w/w) protease to protein lysate with MS-grade Trypsin/Lys-C Protease Mix (Thermo A40009) was then performed by mixing overnight at 37 °C.

The samples were then acidified by adding 3x Sample Buffer for final concentration of 0.5% (v/v) trifluoroacetic acid (TFA) and 5% (v/v) acetonitrile (ACN) resulting in sample pH ≤ 3, as confirmed by pH paper. The samples were then desalted using self-packed C18 spin columns containing ∼80 ± 10 mg of Sigma Supelclean™ LC-18 SPE Bulk Packing (57202) using all MS-grade chemicals and centrifugation at 1,500 x g for 1 min. The columns were activated using 400 µL of 50% (v/v) methanol in water and then equilibrated using 400 µL twice of Wash Buffer (0.5% (v/v) TFA and 5% (v/v) ACN in water). Samples were then loaded onto the equilibrated column by passing the sample twice through the column into the sample microcentrifuge tube. The samples were then washed four times using 400 µL of Wash Buffer. The desalted peptides were then eluted with two washes of 100 µL of 0.1% formic acid (FA) and 70% ACN in water and then immediately dried on a centrifugal vacuum concentrator (Labconoco at maximum 45°C. The peptides were then reconstituted in MS-grade 0.1% FA in water before determining their concentration using a Pierce Quantitative Fluorometric Peptide Assay (Thermo Scientific 23290) following the manufacturer’s protocol.

### LC-MS/MS analysis

The reconstituted desalted peptides were then analyzed using a nanoElute 2 UHPLC (Bruker Daltonics, Bremen, Germany) coupled to a timsTOF HT (Bruker Daltonics, Bremen, Germany) via a CaptiveSpray nano-electrospray source. The peptides were separated in the UHPLC using an Aurora Ultimate nanoflow UHPLC column with CSI fitting (25 cm x 75 µm ID, 1.7 µm C18; IonOptics AUR3-25075C18-CSI) over a gradient from 4% to 26% (v/v) of Mobile Phase B (MPB; 0.1% FA in ACN) in Mobile Phase A (MPA; 0.1% FA in water) over 60 min, then 26% to 32% MPB for 5 min, and finally 95% MPB for up to total 70 min with a flow rate of 350 nL/min with column temperature maintained at 50 °C.

The TIMS elution voltages were calibrated linearly with three points (Agilent ESI-L Tuning Mix Ions; 622, 922, 1,222 m/z) to determine the reduced ion mobility coefficients (1/K_0_). diaPASEF was performed using the MS settings 100 m/z for Scan Begin and 1700 m/z for Scan End in positive mode, the TIMS settings 0.70 V⋅s/cm^2^ for 1/K_0_ start, 1.30 V⋅s/cm^2^ for 1/K_0_ end, ramp time of 120.0 ms, 100% duty cycle, ramp rate of 7.93 Hz, and the capillary voltage set to 1600 V. diaPASEF windows from mass range 226.8 Da to 1226.8 Da and mobility range 0.70 1/K_0_ to 1.30 1/K_0_ were designed to provide 25 Da windows covering doubly and triply charged peptides as confirmed by DDA-PASEF scans, whereas singly charged peptides were excluded from the acquisition due to their position in the m/z-ion mobility plane.

#### Processing and statistical analysis

The raw diaPASEF files were processed using library-free analysis in DIA-NN 1.8.1(*61*) to identify and quantify peptides and their false discovery rate through in silico digestion of the human proteome (Uniprot reviewed human proteome UP000005640 containing reverse sequence decoys and common contaminants; accessed April 12, 2023) using deep learning-based predictions to generate a spectral library from the DIA precursor data and MS spectra, which is then used for targeted DIA analysis by spectral matching after determining MS accuracy. This search followed default settings with the following modifications: “FASTA digest” and “Deep-learning-based spectra” settings enabled, “trypsin/P” (includes cleavage after P) for the protease with up to 2 missed cleavages, carbamidomethylation of cysteine, oxidation of methionine and precursor Q-value (FDR) cut-off of 0.01, MBR enabled, and “Robust LC (high accuracy)” for precursor quantification strategy.

The identified and quantified peptides from the DIA-NN analysis were further analyzed in R. Peptides were filtered by removing (i) reverse and contaminant peptides, (ii) peptides with Global.Q.Value (FDR for peptide across all samples) and/or PG.Q.Value (FDR for quantified protein for peptide) greater than 0.0105, and (iii) at least two observed intensities values in at least one condition. Protein intensities were then re-calculated using the MaxLFQ method provided in the DIA-NN R package(*61*) and then differential statistics was performed using the DEqMS (a modified LIMMA) R package(*62*) to determine p-value, fold change, and the Benjamini-Hochberg adjusted p-value.

### RNA-seq Analysis

Raw reads were checked for quality using fastqc (https://www.bioinformatics.babraham.ac.uk/projects/fastqc/) and trimmed from adapters using cutadapt(*63*) using parameters cutadapt -a AGATCGGAAGAGCACACGTCTGAACTCCAGTCA-b AGATCGGAAGAGCGTCGTGTAGGGAAAGAGTGT --nextseq-trim=20 --minimum-length 1. Transcripts were quantified using kallisto(*64*) against the human Gencode v33 indexed transcriptome and annotations. Differential gene analysis was performed using DESeq2(*65*) using apeglm(*66*) to shrink log_2_FoldChanges and pathway and enrichment analyses using Enrichr(*67*) and ChIP-Atlas(*68*). GSEA analysis was performed on an in-house dataset consisting of LymphDB (L. Staudt, NIH, https://lymphochip.nih.gov/signaturedb/), MSigDB pathways, and an in-house dataset generated at MD Anderson of lymphoma-specific signaling.

### ChIP-seq Experiment and Library Preparation

25-30 million cells were treated with CDK-TCIP1 or DMSO for indicated timepoints. Cells were washed in PBS containing CDK-TCIP1 and crosslinked for 12 min in CiA Fix Buffer (50 mM HEPES pH 8.0, 1 mM EDTA, 0.5 mM EGTA, 100 mM NaCl) with addition of formaldehyde to a final concentration of 1%. The crosslinking reaction was quenched by glycine added at 0.125 M final concentration. Crosslinked cells were centrifuged at 1,000 x g for 5 min. Nuclei were prepared by 10 min incubation of resuspended pellet in CiA NP-Rinse 1 buffer (50 mM HEPES pH 8.0, 140 mM NaCl, 1 mM EDTA, 10% glycerol, 0.5% IPEGAL CA-630, 0.25% Triton ×100) followed by wash in CiA NP-Rinse 2 buffer (10 mM Tris pH 8.0, 1 mM EDTA, 0.5 mM EGTA, 200 mM NaCl). The pellet was resuspended in CiA Covaris Shearing Buffer (0.1% SDS,1 mM EDTA pH 8.0, 10 mM Tris HCl pH 8.0) with 1000x protease inhibitors (Roche) and sonicated for 40 min with Covaris E220 sonicator (Peak Power 140, Duty Factor 5.0, Cycles/Burst 200). The distribution of fragments was confirmed with D1000 Tapestation. 50 µg of chromatin per ChIP was used with anti-CDK9 antibodies (Cell Signaling CST2361S, abcam ab239364), with 40 ng Drosophila chromatin (53083, ActiveMotif) spiked in. 50 µg of chromatin was used with anti-Pol II ser 2 phos (2 µg, Abcam ab5095) and anti-Pol II ser 5 phos (2 µg, Abcam ab5131) antibodies, and anti-Pol II (5 µg, Santa Cruz 8WG16) antibodies. After overnight incubation at 4 ^O^C in IP buffer (50 mM HEPES pH 7.5, 300mM NaCl, 1mM EDTA, 1% Triton ×100, 0.1% sodium deoxycholic acid salt (DOC), 0.1% SDS), IPs were washed twice with IP buffer, once with DOC buffer (10 mM Tris pH 8, 0.25 M LiCl, 0.5% IPEGAL CA-630, 0.5% sodium deoxycholic acid salt (DOC), 1mM EDTA), and once with 10 mM Tris/1 mM EDTA buffer (TE) pH 8. IPs and inputs were reverse-crosslinked in TE/0.5% SDS/0.5 µg/µL proteinase K for 55 ^°^C/3 hrs then 65 ^O^C/18 hrs, then DNA was purified using a PCR cleanup spin column (Takara #74609). The sequencing library preparation was performed using NEBNext Ultra II DNA kit (#E7645S). Libraries were sequenced on an Illumina NovaSeq (Novogene).

### ChIP-seq Analysis

The data quality was checked using fastqc. The raw reads were trimmed from adapters with trim_galore (parameters: --paired –illumina) and raw reads were aligned to hg38 human genome assembly and the dm6 fly genome assembly using bowtie2 (parameters: --local -- maxins 1000). Low quality reads, duplicated reads and reads with multiple alignments were removed using samtools(*69*) and Picard (https://broadinstitute.github.io/picard/). macs2(*70*) was used to map position of peaks with FDR cutoff of 0.05. Bedtools(*71*) was used to find consensus set of peaks by merging peaks across multiple conditions (bedtools merge), count number of reads in peaks (bedtools intersect -c) and generate genome coverage (bedtools genomecov - bga). deepTools(*72*) was used to generate coverage densities across multiple experimental conditions (deeptools computeMatrix and deeptools plotProfile) and to generate bigwig files (deeptools bamCoverage), where reads mapping to ENCODE blacklist regions were excluded(*73*). For CDK9, normalization was performed as suggested by the manufacturer protocol (61686 and 53083, ActiveMotif) in which the human genome-mapped unique reads in each ChIP were downsampled proportional to a normalization factor calculated by: (1) counting the unique reads in each sample that align to the fly genome; (2) identifying the sample containing the least amount of mapped fly genome reads; and (3) computing the normalization factor for each sample as (reads mapping to the fly genome in the sample with minimum mapped fly reads)/(reads mapping to the fly genome in the current sample). This procedure was carried out on a per-antbody and per-cell basis. All browser tracks and metaprofiles shown were calculated with spike-in-normalized and input-subtracted data. All Pol II, Pol II ser 2 phos, and Pol II ser 5 phos browser tracks and metaprofiles shown were calculated with sequence-depth normalized and input-subtracted data. The peak differential analysis and PCA analysis was performed using DESeq2(*65*). GREAT(*30*) analysis was performed using a background set of all detected peaks and an association rule of “single nearest gene”, since Pol II serine phosphorylation is closely linked to transcription. Pausing ratio was calculated as the ratio of integrated Pol II ser 5 phos at the transcription start site (TSS) - 100 basepairs (bp) to TSS + 200 bp, and the integrated Pol II ser 2 phos at the transcription end site (TES) + 2 kbp, as described in(*28*). The SRX4609168 BCL6 ChIP-seq dataset from SUDHL4 cells was used to call peaks and extract positions of BCL6 summits and loading, defined as enrichment fold change over input. For annotation of high-confidence BCL6-regulated promoters and enhancers, only summits with fold change ≥ 10 were included. Data from Bal et al(*11*) was used to annotate enhancers using ROSE(*74*) by stitching together H3K27ac peaks in untreated cells within 12.5 kb but excluding regions within 2 kb of a transcription start site unless within a larger H3K27ac domain..

### Mouse Tolerability and Pharmacokinetic Study

The PK and tolerability study was performed in the Drug Metabolism and Pharmacokinetics (DMPK) Core facility at Scripps Florida (https://www.scripps.edu/science-and-medicine/cores-and-services/dmpk-core/index.html). The mice are housed in individually ventilated cages (IVC) in JAG 75 cages with micro-isolator lids. HEPA filtered air is supplied into each cage at a rate of 60 air exchanges per hour for the mice. The dark/light cycle is set for 8:00pm on-8:00pm off.

The temperature is set for 72 ^O^F and are maintained +/-2 ^O^F. The humidity is low/Hi of 30-70%. There is a computerized system in place to control and or monitor the temperatures within the Animal Holding Room. Each animal room is equipped with a thermos-hygrometer that is monitored and recorded daily on room log. 5 mg/kg of CDK-TCIPs were injected intraperitoneally into C57BL/6 male mice (n=3 in treatment and n=3 in vehicle conditions) using a 25-29 gauge needle to deliver 10 µL/g body weight of a formulation of 0.5 mg/mL CDK-TCIPs in 5% DMSO, 5% Tween-80, and 90% Saline. Vehicle was the same formulation (5/5/90 DMSO/Tween-80/Saline). The formulation was checked to be a clear solution and after administration, the animal was put back in its cage. For PK properties, plasma levels were measured at 0, 5, 15, 30, 60, 120, 240, 360, and 480 mins after drug administration. For tolerability work, bodyweights and observation of animal health were recorded each day through 5 days of dosing once daily. After 5 days, tissues were collected 8 hours after the last drug administration and taken for measurement of drug levels where molar concentrations were recorded with the assumption of 1g tissue ∼ 1mL. Samples were processed for analysis by precipitation using acetonitrile and analyzed with LC/MS/MS. PK parameters were calculated using the noncompartmental analysis tool of WinNonlin Enterprise software (version 6.3). All procedures were approved by the Scripps Florida Institutional Animal Care and Use Committee (IACUC), and the Scripps Vivarium is fully accredited by the Association for Assessment and Accreditation of Laboratory Animal Care International.

### Mouse Immunization and Germinal Center Response Assessment

To induce germinal center (GC) formation, 8-week-old male C57BL/6J mice were immunized with 0.5 mL of a 2% sheep red blood cells (SRBC) suspension (Cocalico Biologicals) in PBS intraperitoneally. To evaluate the effect of CDK-TCIP2 on GC formation, mice were randomly grouped as vehicle (8% DMSO, 5% Tween-80 and 87% saline), 5 mg/kg CDK-TCIP2 once a day, 10 mg/kg CDK-TCIP2 once a day, and 5 mg/kg CDK9-TCIP2 twice a day. The treatment was initiated 2 days after SRBC immunization by intraperitoneal injection. Mice were monitored daily and weighted every other day. Mice were euthanized after 8 days of treatment. Spleens were homogenized by filtering through a 40 µm cell strainer followed by red blood cell lysis using ACK Lysing Buffer (A1049201, Thermo Fisher Scientific). Cells were resuspended in PBS with 0.5% BSA, performed with Fc blocking (422302, BioLegend), and labelled with anti-CD138 (142508, BioLegend), anti-GL7 (144614, BioLegend), anti-IgD (405723, BioLegend), anti-CD86 (105037, BioLegend), anti-IgM (406539, BioLegend), anti-B220 (103246, BioLegend), anti-Fas (557653, BD Biosciences), anti-CD184 (742171, BD Biosciences), anti-IgG (103246, Cytek), and anti- CD38 (557653, Cytek) conjugated with fluorochromes for 30 minutes at room temperature. Samples were acquired on the Cytek Aurora flow cytometer and data was analyzed using FlowJo software.

## DATA AVAILABILITY

Uncropped blots of Westerns and flow gating strategy are available in Supplemental Information. Pharmacology data on CDK-TCIPs in DLBCL cells is available in Supplemental Table 1. Abbreviations used in PRISM analysis of cancer cell lines are provided in Supplemental Table 2. Global proteomics data is available in Supplemental Table 3. Genomic sequencing data has been deposited to GSE245600.

### ACKNOWLEDGEMENTS

The studies described in this manuscript were funded from a grant from HHMI to G.R.C., NIH grants CA276167, CA163915 and MH126720-01 to G.R.C. Funding was also provided by a grant from the Mary Kay Foundation. Funding was provided to S.G. from NIH grant 5F31HD103339-03. Research was financially supported by Stanford’s SPARK Translational Research Program. This research was financially supported by Stanford Bio-X. R.C.S. acknowledges the Swiss National Science Foundation for a postdoctoral fellowship (SNF Mobility grant P500PN_206898). B.A.K. acknowledges support from the Molecular Pharmacology Training Program (Stanford University). Funding was provided to N.S.G. from departmental funds from Chemical and Systems Biology and the Stanford Cancer Institute. The Gray lab also receives or has received research funding from Novartis, Takeda, Astellas, Taiho, Jansen, Kinogen, Arbella, Deerfield, Springworks, Interline and Sanofi. Funding for pharmacokinetic studies was provided by NIH grant number 1 S10OD030332-01. M.R.G. is supported by a Leukemia and Lymphoma Society Scholar award. S.G. would like to thank T. Reindl for helpful advice in biochemical studies. All authors would like to thank members of the Crabtree and Gray laboratories for constructive comments.

## AUTHOR CONTRIBUTIONS

G.R.C., N.S.G., S.G., R.C.S. conceived the project. S.G. conducted cell biological, biochemical, and genomic studies. R.C.S. and B.A.K. designed and conducted chemical syntheses. S.A.N. conducted ChIP-seq studies with S.G. J.M.S., J.T., and H.A. carried out experiments designed by G.R.C., S.G., N.S.G., R.C.S., and S.M.H. S.G., B.A.K and S.M.H. conducted studies with FKBP constructs. J.T. and S.M.H. purified CDK9 protein. B.G.D. performed (phospho)proteomic experiments. B.A.K. conceived the CDK12/13 synthesis and together with S.G., conducted cell biological studies on those compounds. H.Y. and M.R.G. contributed GSEA analyses relevant to DLBCL and studies of the GC response in mice. A.K. and T.Z. contributed to CDK-TCIP biological application and chemical synthesis, respectively. The manuscript was written by G.R.C, R.C.S., S.G., N.S.G. with input from all authors.

## SUPPLEMENTAL FIGURES

**Supplemental Figure 1.**
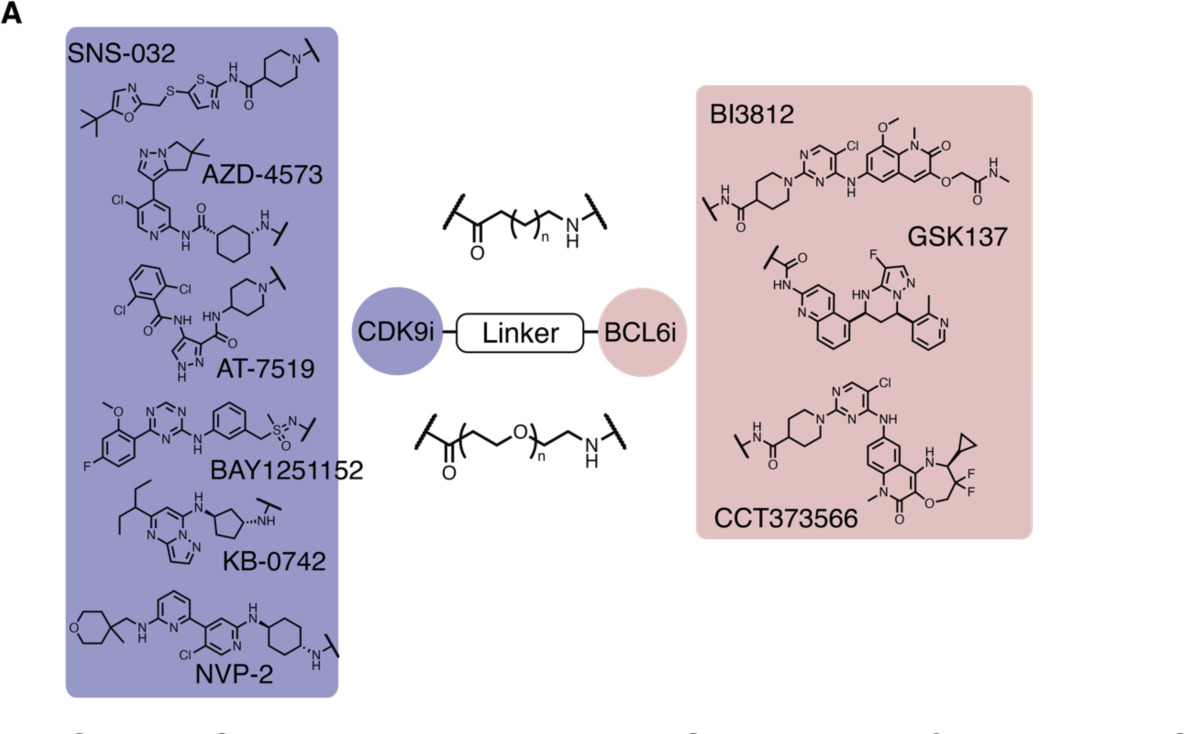
CDK-TCIP library design. **A. Structures of employed CDK9 inhibitors and BCL6^BTB^ binders.**

**Supplemental Figure 2.**
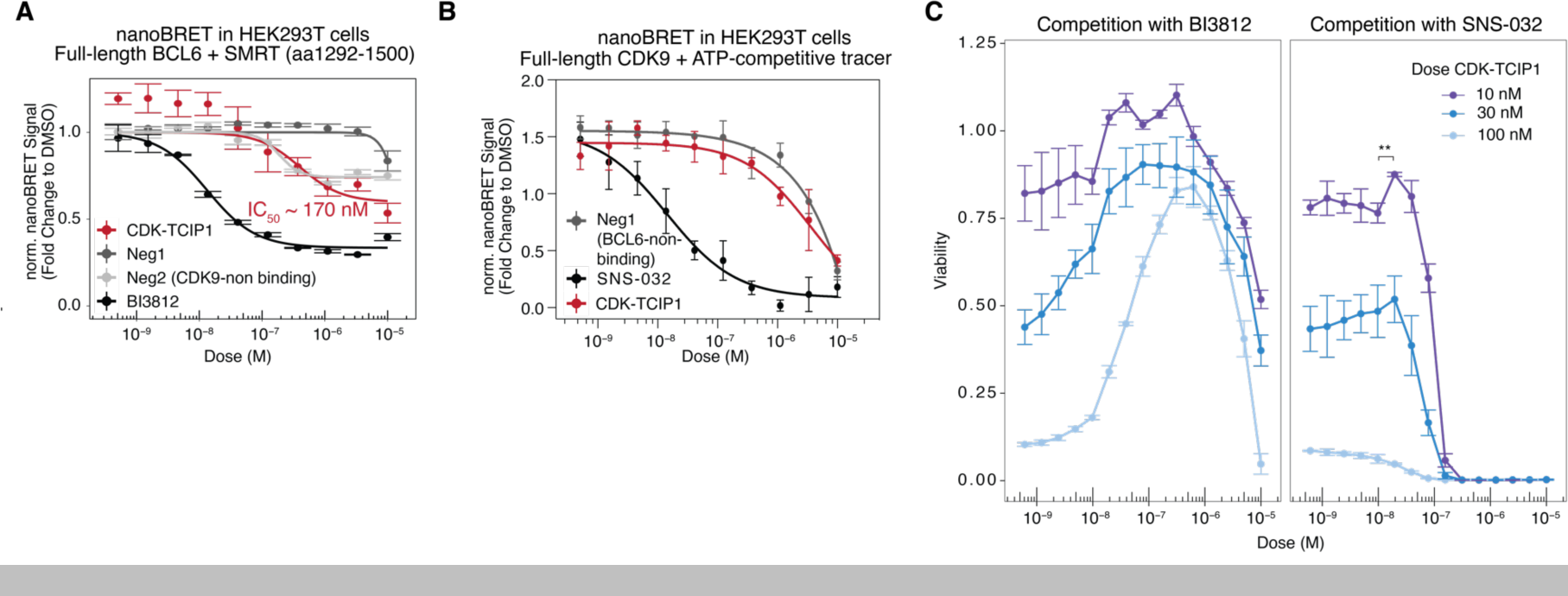
Characterization of negative controls and viability rescue experiments. **A.** Displacement of SMRT/BCL6 in HEK293T cells by **CDK-TCIP1**. **B.** Displacement of ATP-competitive tracer probe to CDK9 in HEK293T cells by **CDK-TCIP1**. For **A, B:** mean±s.d., n=3 technical replicates. **C.** Competitive titration of constant **CDK-TCIP1** with a BCL6^BTB^ inhibitor (BI3812) or a CDK9 inhibitor (SNS-032) to reverse the cell death phenotype; mean±s.d., n=3 biological replicates.

**Supplemental Figure 3.**
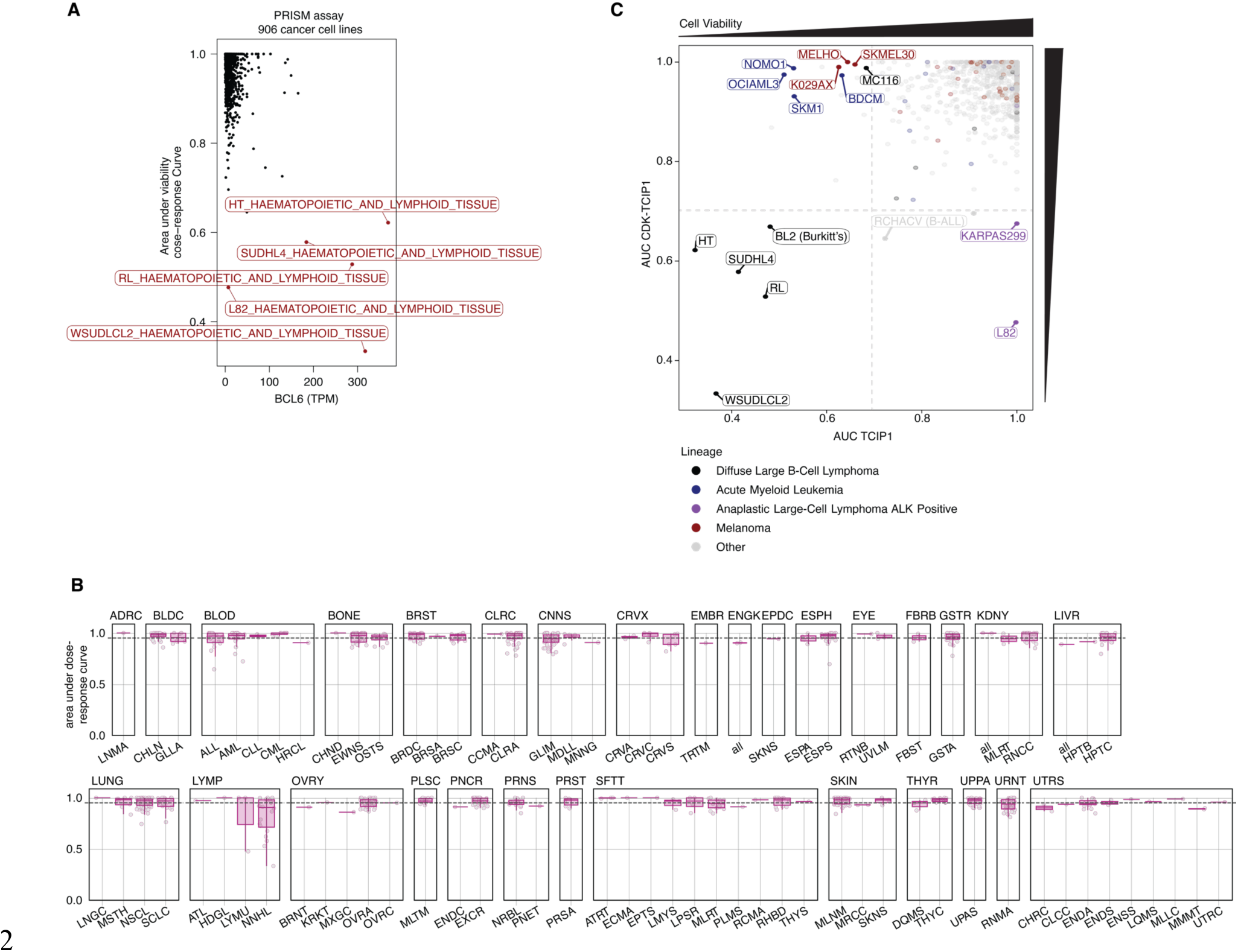
Analysis of CDK-TCIP1 potency in 859 cancer cell lines. **A.** Correlation of sensitive cell lines as indicated by area-under-dose-response curve and *BCL6* transcripts/million (TPM). **B.** Area-under-dose-response curve for each cancer cell line treated with CDK-TCIP1 for 120 h categorized by lineage subtype. See Supplemental Table 2 for list of abbreviations. **C.** Correlation of sensitivity to CDK-TCIP1 with sensitivity to TCIP1 (*19*).

**Supplemental Figure 4.**
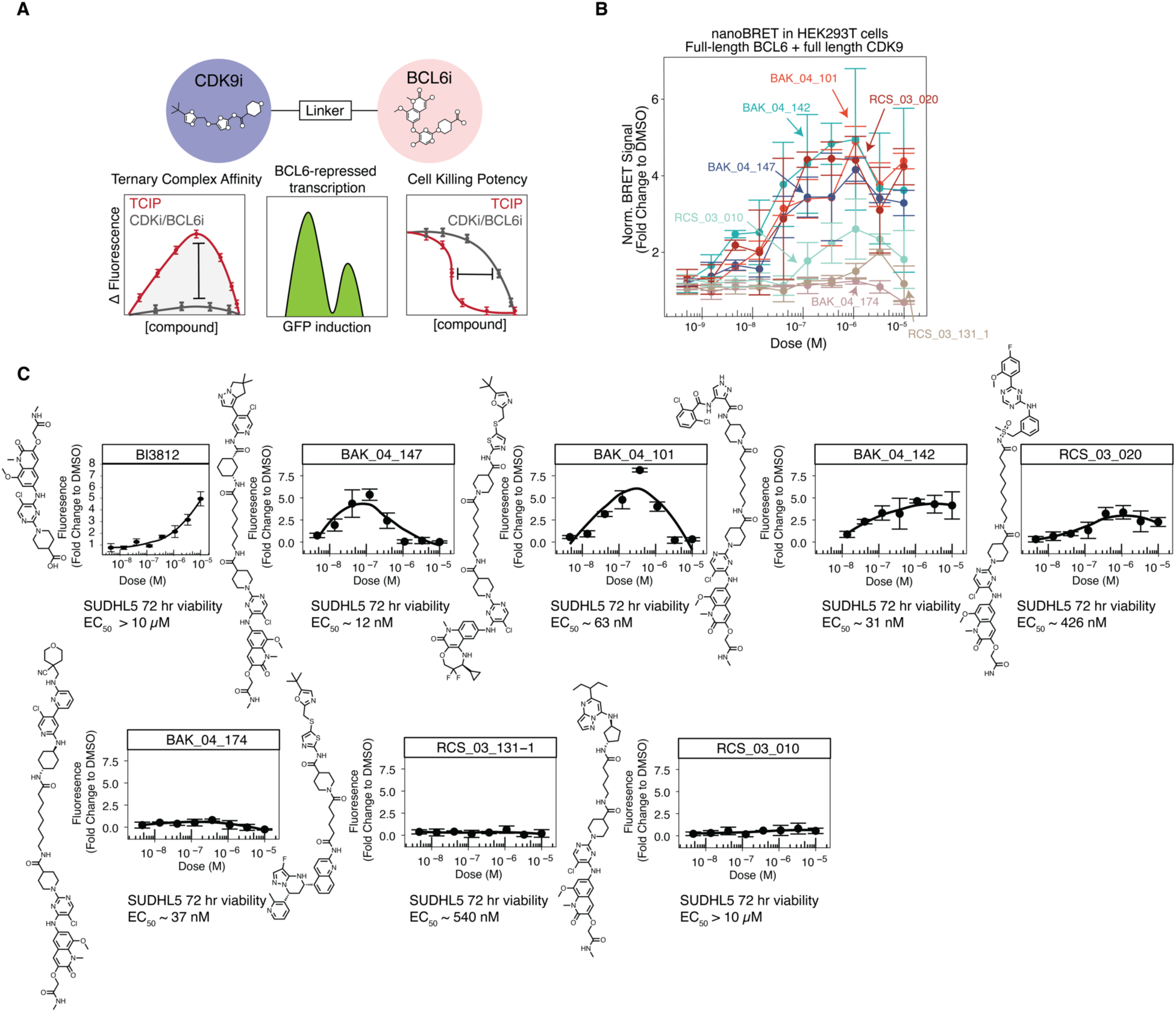
Assay cascade to rationally design CDK-TCIPs. **A.** Biochemical design cascade implemented. **B.** Ternary complex formation inside cells between full-length BCL6 and CDK9 overexpressed in HEK293T cells after CDK-TCIP addition. **C.** BCL6 reporter activation to top CDK-TCIPs from each inhibitor class with structures and 72-hour cell killing potencies (EC_50_ calculated from mean of 3 technical repeats) in SUDHL5 cells shown; for reporter, mean±s.d., n=3 technical replicates.

**Supplemental Figure 5.**
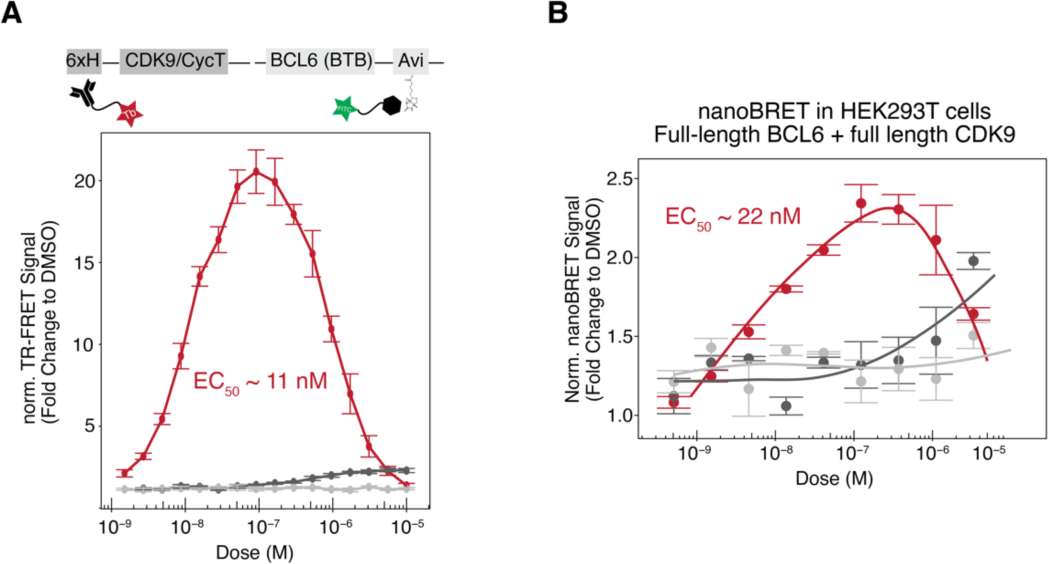
Characterization of CDK-TCIP ternary complex formation. **A.** Ternary complex formation as measured by TR-FRET (Methods) between purified CDK9/CycT1 and BCL6^BTB^. **B.** Ternary complex formation inside cells between full-length BCL6 and CDK9 overexpressed in HEK293T cells after CDK-TCIP addition. For **A, B:** mean±s.d., n=3 technical replicates.

**Supplemental Figure 6.**
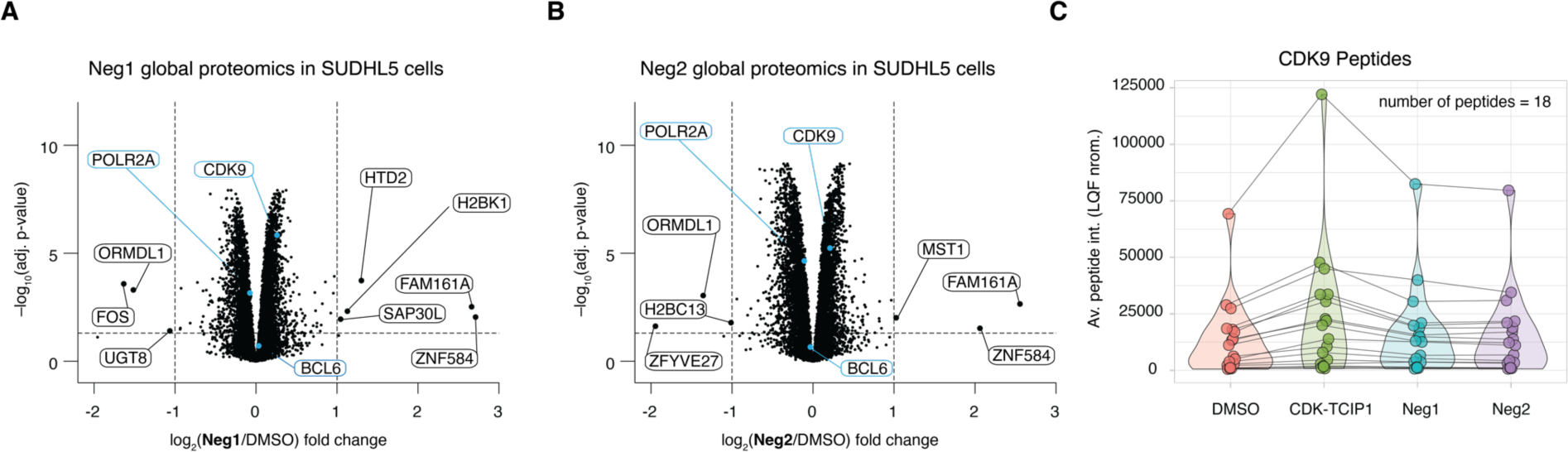
Global proteomic characterization of negative controls in SUDHL5 cells. **A.** Whole cell global proteomics of SUDHL5 cells treated with 30 nM **Neg1** or (**B**) **Neg2** for 2 h. **C.** Paired violin plot of quantified CDK9 peptides in each condition from global proteomic analysis, where each dot represents the average MaxLFQ normalized intensity for a given peptide with paired lines indicating that peptide across conditions. For **A & B:** 4 biological replicates, p-values computed by moderated t-test and adjusted by Benjamini-Hochberg, significance cutoffs at |log_2_(fold change)≥1| and p_adj_≤0.05.

**Supplemental Figure 7.**
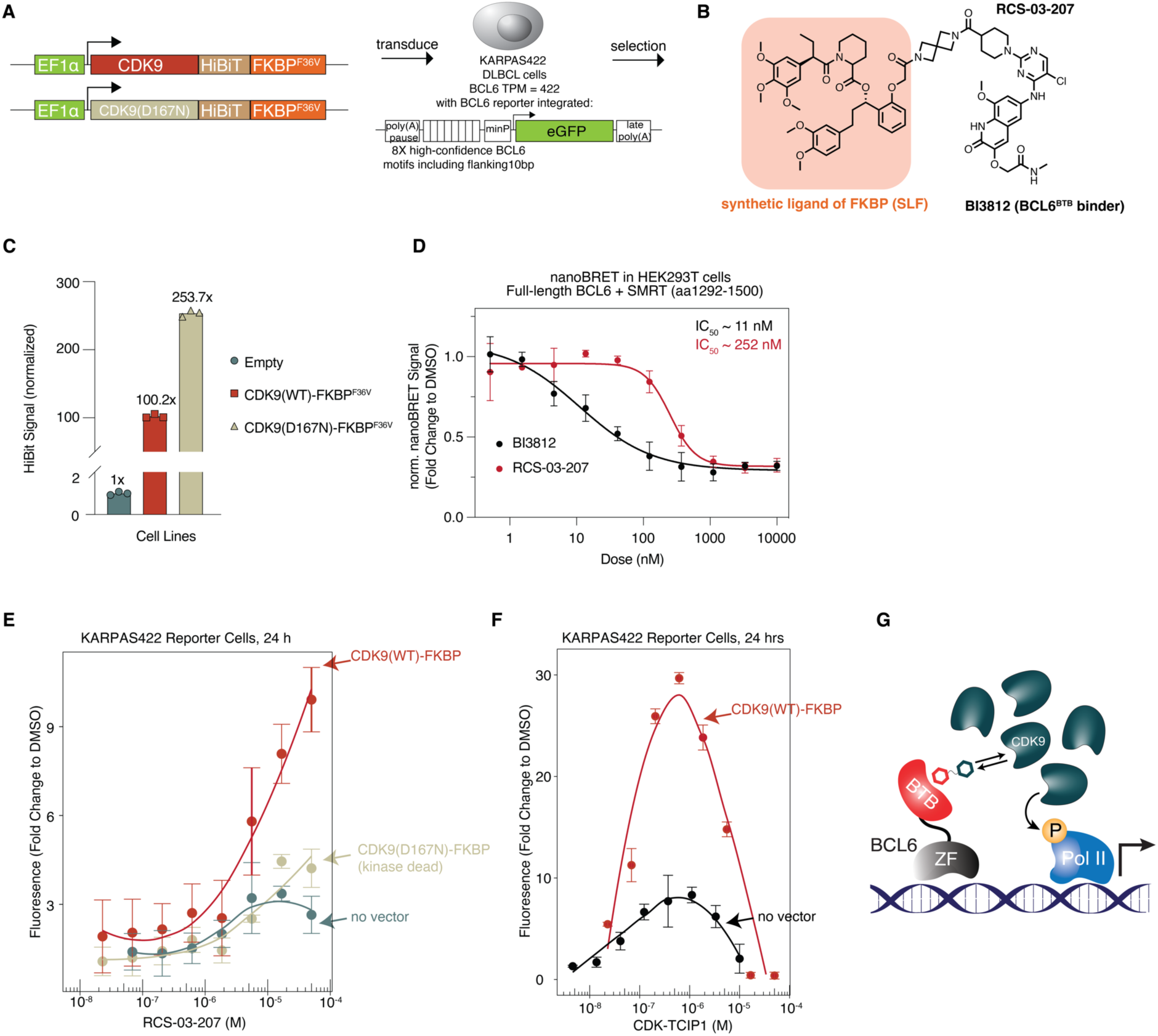
CDK9 recruitment by synthetic chemical inducers of proximity. **A.** Construction of FKBP^F36V^-tagged CDK9 wild-type and mutant overexpression system in BCL6 reporter cells. **B.** Structure of **RCS-03-207**, a bivalent molecule designed to dimerize FKBP and BCL6. **C.** Expression levels of overexpressed construct as measured by HiBit tag; mean±s.d., n=3 technical replicates. **D.** Comparison of cell-penetrability and displacement of SMRT/BCL6 in HEK293T cells by BI3812 and RCS-03-207; mean±s.d., n=3 technical repeats. **E.** Change in GFP expression in reporter cell lines 24 h after drug addition. n=2-3 biological replicates representative of two different reporter cell lines generated, mean±s.d. **F.** Response of reporter to **CDK-TCIP1** after overexpression of wild-type CDK9-FKBP^F36V^; mean±s.d., n=3 technical replicates. **G.** Working model of CDK-TCIP function.

**Supplemental Figure 8.**
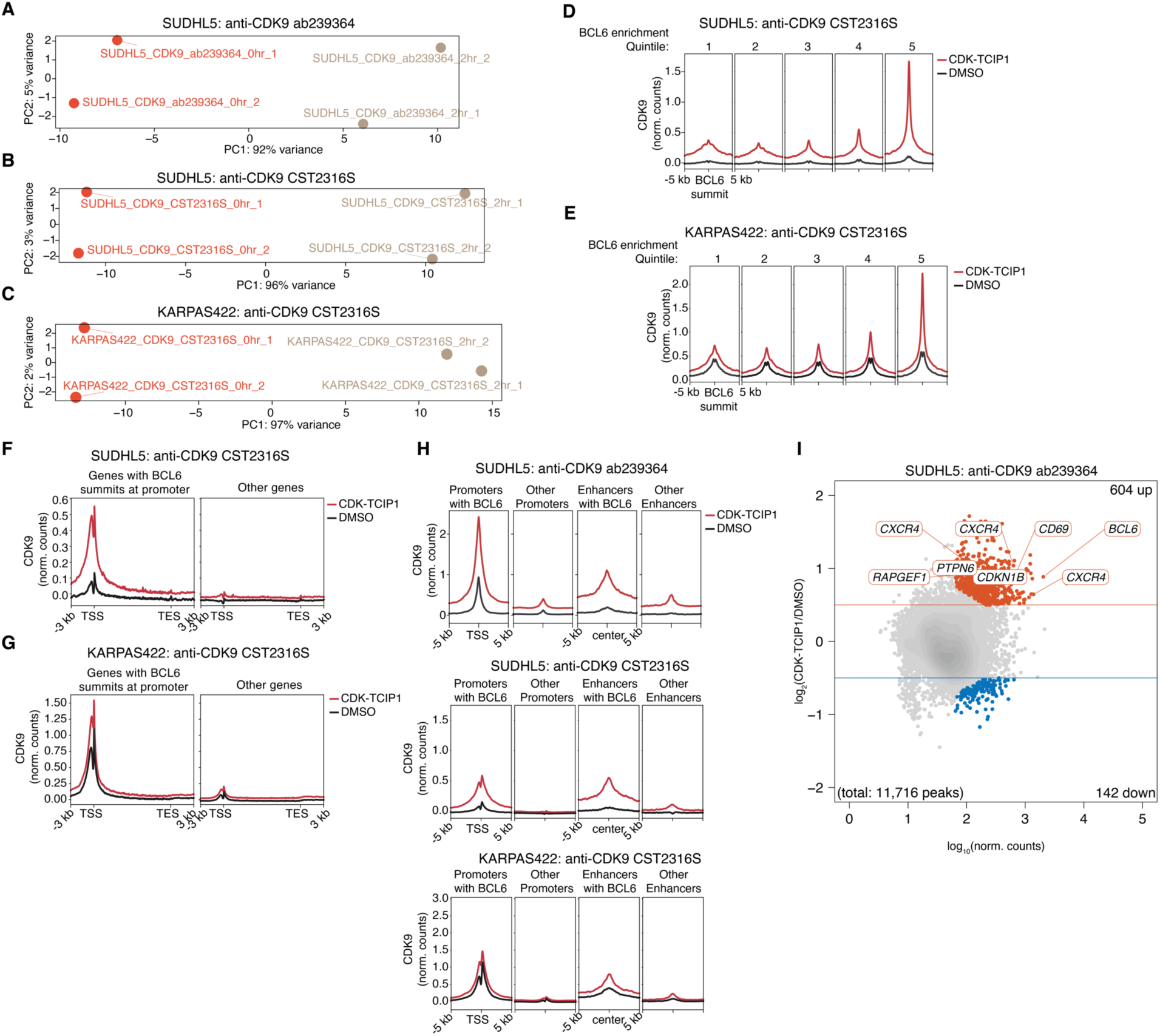
CDK9 recruitment to BCL6-bound chromatin in lymphoma cells. Principal component analyses for CDK9 ChIP-seq as performed with **A.** anti-CDK9 ab239364 in SUDHL5 cells, **B.** anti-CDK9 CST2316S in SUDHL5 cells, and **C.** anti-CDK9 CST2316S in KARPAS422 cells. Recruitment of CDK9 to BCL6 summits as detected with anti-CDK9 CST2316S in **D.** SUDHL5 cells and **E.** KARPAS422 cells. Increase in CDK9 at genes with BCL6 summits at their promoter as detected with anti-CDK9 CST2316S in **F.** SUDHL5 cells and **G.** KARPAS422 cells. Increase in CDK9 at promoters and enhancers with BCL6 summits in **H.** as detected with (top) anti-CDK9 ab239364 in SUDHL5 cells, (middle) anti-CDK9 CST2316S in SUDHL5 cells, and (bottom) with anti-CDK9 CST2316S in KARPAS422 cells. In **D-H,** CDK9 metaprofiles were merged from two biological replicates, spike-in normalized, and input-subtracted. **I.** Differential CDK9 peaks as detected with anti-CDK9 ab239364 in SUDHL5 cells; significance cutoffs were p_adj_≤0.05 and |log_2_(fold change)|≥0.5), n=2 biological replicates; p- values computed by two-sided Wald test and adjusted for multiple comparisons by Benjamini-Hochberg. In all panels **A-I, CDK-TCIP1** was added at 30 nM for 2 h.

**Supplemental Figure 9.**
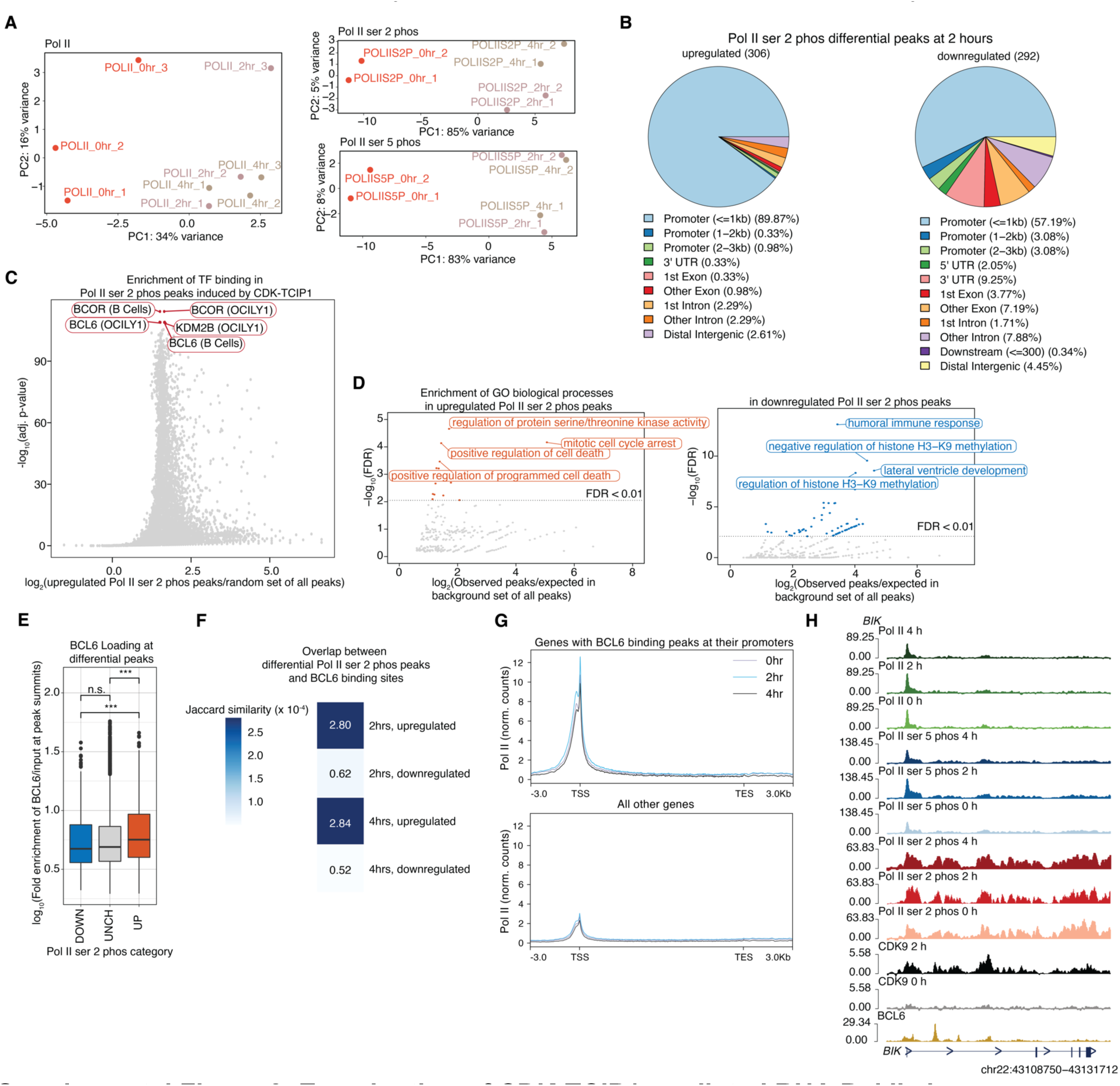
Examination of CDK-TCIP1-mediated RNA Pol II changes on chromatin. **A.** Principal component analysis of biological replicates for Pol II (n=3), Pol II ser 2 phos (n=2), and Pol II ser 5 phos (n=2) ChIP-seq experiments. **B.** Annotation of differential peaks in Pol II ser 2 phos after 2 hours 30 nM CDK-TCIP1 addition. **C.** Analysis of transcription factor binding at upregulated Pol II ser 2 phos peaks at 2 h in ≥6,500 public ChIP-seq datasets in blood-lineage cells. **D.** Enrichment of GO Biological Processes in upregulated (left) or downregulated (right) Pol II ser 2 phos peaks at 2 h by GREAT (Methods). For **C, D:** P-values computed by two-sided Fisher’s exact test and adjusted for multiple comparisons by Benjamini- Hochberg. **E.** BCL6 loading at peaks overlapping with differential Pol II ser 2 phos peaks. Students’ two-tailed, unpaired t-test, adjusted for multiple comparisons, ***: p<0.001. **F.** Overlap between differential Pol II ser 2 phos peaks and BCL6 peaks as computed by Jaccard similarity. **G.** Pol II at gene bodies after 30 nM **CDK-TCIP1** addition for the indicated timepoints. Average of n=3 biological replicates, sequence-depth normalized. **H.** ChIP-seq tracks at known cell death gene *BIK* regulated by BCL6 of CDK9, Pol II, Pol II ser 2 phos, and Pol II ser 5 phos after 30 nM **CDK-TCIP1** addition for the indicated timepoints; CDK9 was merged from two spike-in-normalized and input-subtracted biological replicates; Pol II ser 2 phos, and Pol II ser 5 phos merged from two sequence-depth-normalized and input-subtracted biological replicates and Pol II from three biological replicates; BCL6 track from(*57*).

**Supplemental Figure 10.**
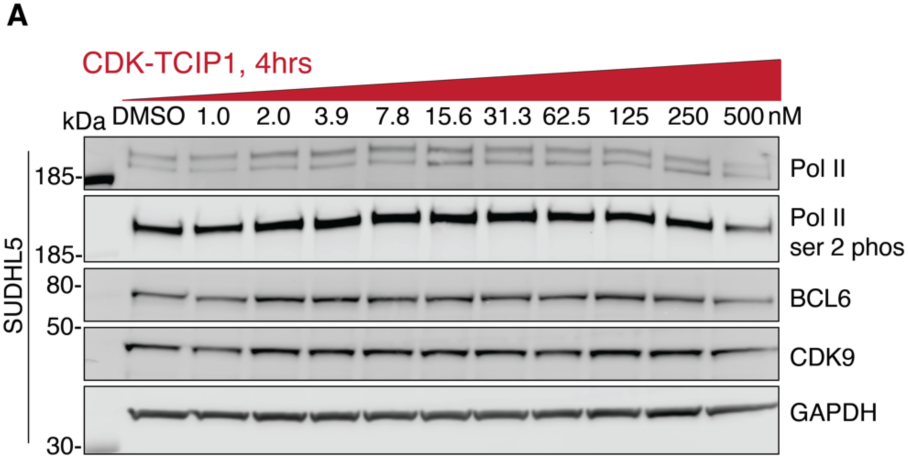
CDK-TCIP1 effect on global levels of Pol II, BCL6, and CDK9. **A.** Effect of 30 nM CDK-TCIP1 on protein and phosphorylation levels in SUDHL5 cells after 4 h of treatment.

**Supplemental Figure 11.**
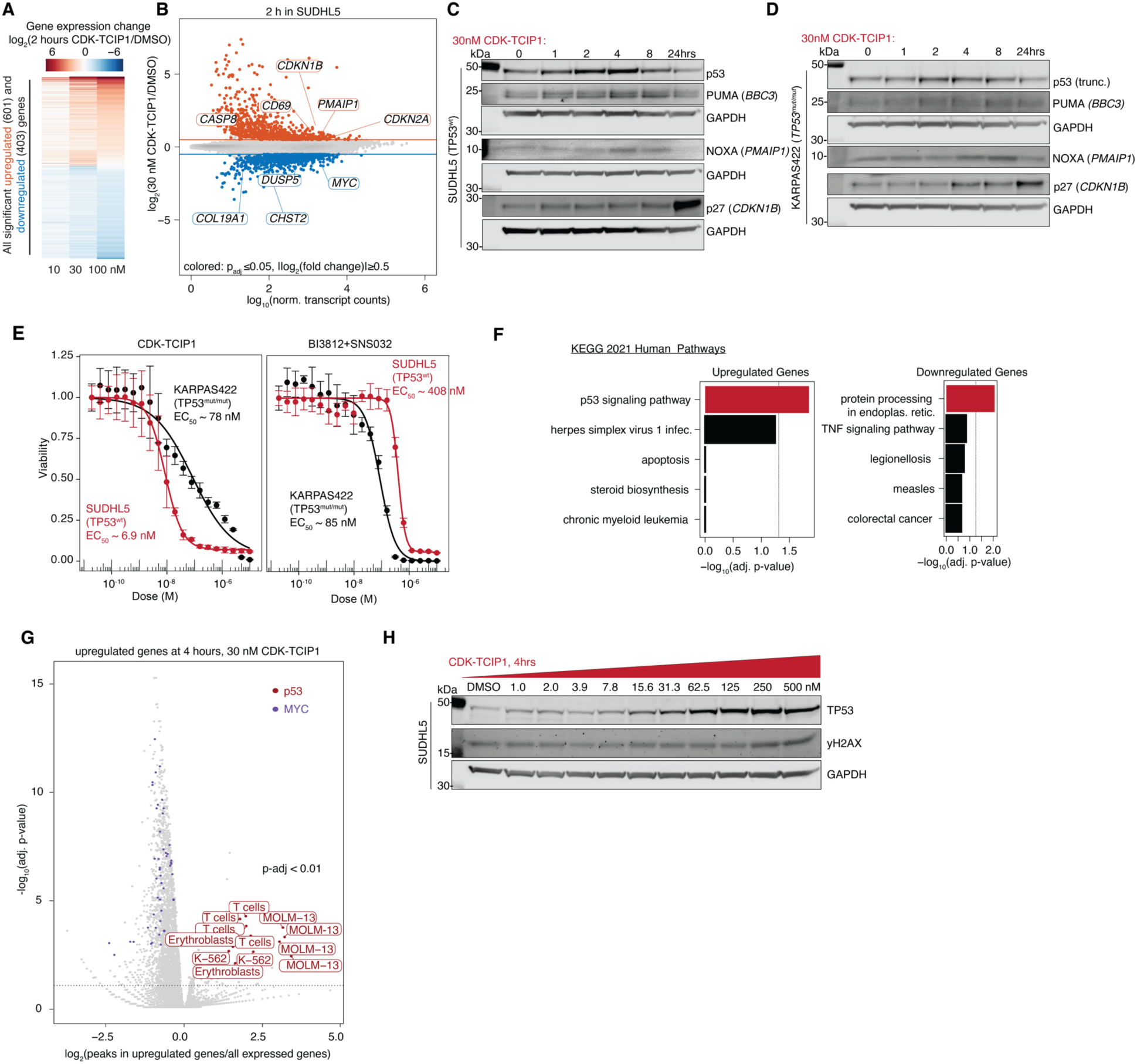
Transcriptomic characterization of the effects of CDK-TCIP1 in cells. **A.** Dose-dependent changes in gene expression by 30 nM **CDK-TCIP1** addition to SUDHL5 cells for 2 h; n=3-4 biological repeats, p-values computed by two-sided Wald test and adjusted for multiple comparisons by Benjamini-Hochberg, significance cutoffs were (p_adj_≤0.05, log_2_(fold change)≥1). **B.** Changes in gene expression after 30 nM **CDK-TCIP1** addition for 2 hours. n=3-4 biological repeats; p-values computed by two-sided Wald test and adjusted for multiple comparisons by Benjamini-Hochberg. **C.** (reproduced from Fig. 3C for comparison)**, D.** Time-dependent changes in apoptotic protein activation, p53 signaling, and downstream proteins after addition of 30 nM CDK-TCIP1. Representative of n=3 biological replicates for SUDHL5, n=1 for KARPAS422. **E.** Sensitivity of *TP53^WT^* (SUDHL5) and *TP53^mut/mut^*(KARPAS422) DLBCL cell lines to CDK-TCIP1 and the additive effect of BCL6 (BI3812) and CDK9 (SNS-032) inhibitors; mean±s.d., n=3 biological repeats for SUDHL5, n=2 biological repeats for KARPAS422. **F.** KEGG Pathway analysis among upregulated (left) or downregulated (right) genes (p_adj_≤0.05, |log_2_(Drug/DMSO)|≥1) after 4 hours 30 nM **CDK-TCIP1** drug addition. p-values computed by two-sided Fisher’s exact test and adjusted for multiple comparisons by Benjamini-Hochberg. **G.** Analysis of transcription factor binding at promoters (±5kb) of upregulated genes after 4 hours 30 nM CDK-TCIP1 addition (p_adj_≤0.05, log_2_(Drug/DMSO)≥1) in ≥6,500 public ChIP-seq datasets in blood-lineage cells. P-values computed by two-sided Fisher’s exact test and adjusted for multiple comparisons by Benjamini- Hochberg. **H.** Dose-dependent changes in p53 and γH2AX in SUDHL5 cells treated with **CDK- TCIP1** for 4 h; representative of n=2 biological replicates.

**Supplemental Figure 12.**
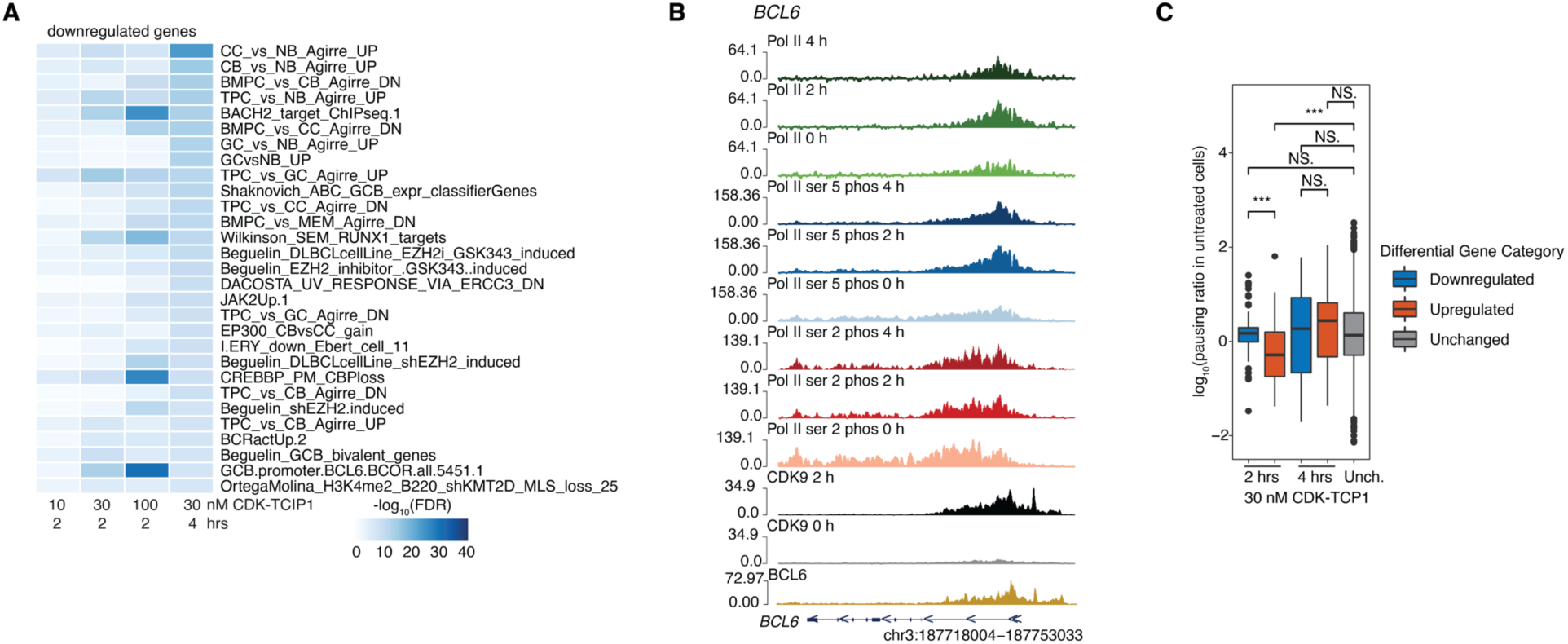
BCL6 target genes decreasing in expression are more paused. **A.** Gene set enrichment analysis of downregulated genes (p_adj_≤0.05, log_2_(fold change)≥0.58); p-values computed by hypergeometric test and adjusted for multiple comparisons by Benjamini- Hochberg. CB, centroblasts. CC, centrocytes. NB, naïve B-cells. **B.** ChIP-seq tracks of CDK9, Pol II, Pol II ser 2 phos, and Pol II ser 5 phos at *BCL6* after 30 nM CDK-TCIP1 addition for the indicated timepoints; CDK9 was merged from two spike-in-normalized and input-subtracted biological replicates; Pol II ser 2 phos, and Pol II ser 5 phos merged from two sequence-depth- normalized and input-subtracted biological replicates, Pol II from three biological replicates, BCL6 track from(*57*). **C.** Pausing ratio in untreated cells (Methods) for genes upregulated, down-regulated, and unchanged after CDK-TCIP1 treatment in SUDHL5 cells for indicated time points.

**Supplemental Figure 13.**
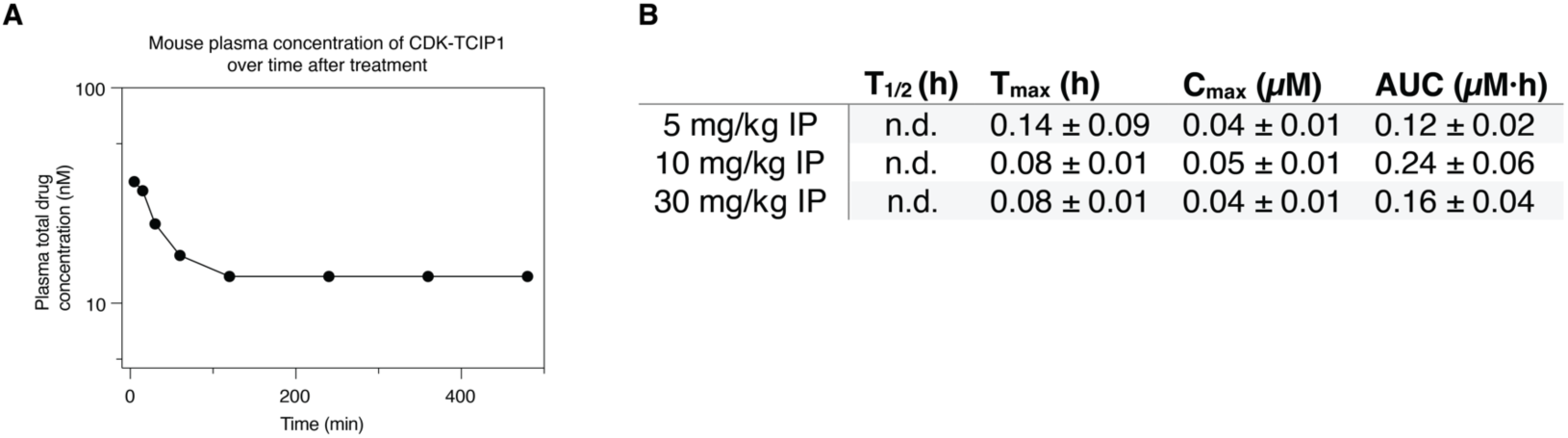
CDK-TCIP1 exhibits poor physicochemical properties. **A.** Mouse blood plasma concentration of **CDK-TCIP1** as function of time after treatment at 5 mg/kg intraperitoneal (IP) dosing. **B.** Pharmacokinetic parameters of **CDK-TCIP1**. t_1/2_, half-life; T_max_, time to maximum serum concentration; C_max_, maximum serum concentration; AUC, area under the curve from dosing to last measured concentration; mean±s.d, n=3 mice.

**Supplemental Figure 14.**
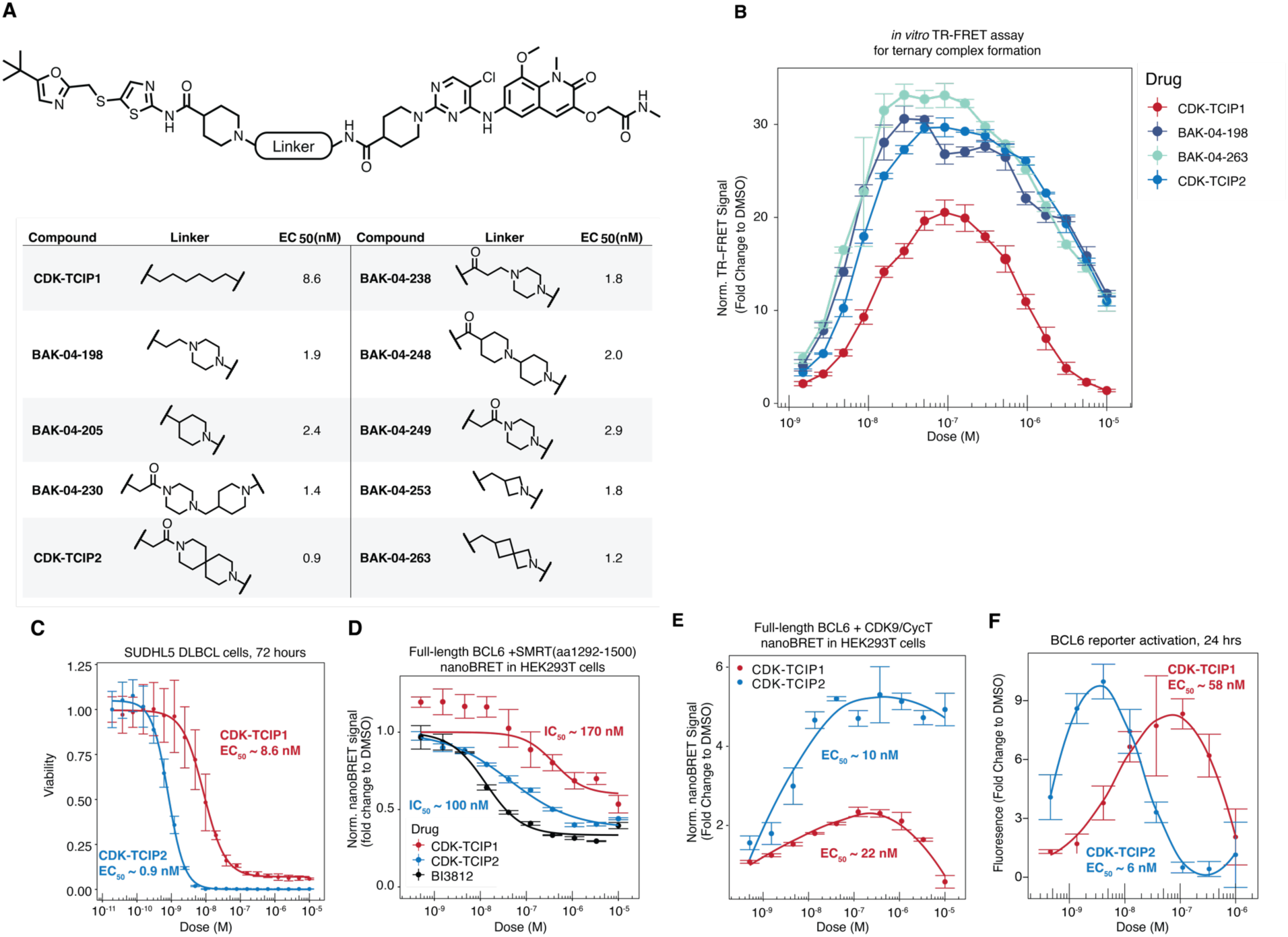
Rational chemical optimization od CDK-TCIPs. **A.** Structure of rigidified CDK-TCIP and cell killing potencies in SUDHL5 cells combined at 72 hours; mean±s.d, n=3 technical repeats. **B.** Ternary complex formation as measured by TR-FRET (Methods) between purified CDK9/CycT1 and BCL6^BTB^. **C.** SUDHL5 cell killing comparison between CDK- TCIP1 and CDK-TCIP2; mean±s.d, n=3 biological replicates. **D.** Comparison of cell-penetrability and displacement of SMRT/BCL6 in HEK293T cells between **CDK-TCIP1** and **CDK-TCIP2**; mean±s.d, n=3 technical repeats. **E.** Ternary complex formation between full-length BCL6 and CDK9 overexpressed in HEK293T cells after addition of **CDK-TCIP1** or **CDK-TCIP2**; mean±s.d, n=3 technical repeats. **F.** Transactivation of BCL6-repressed reporter constructs integrated into lymphoma cells; mean±s.d., n=3 technical replicates.

**Supplemental Figure 15.**
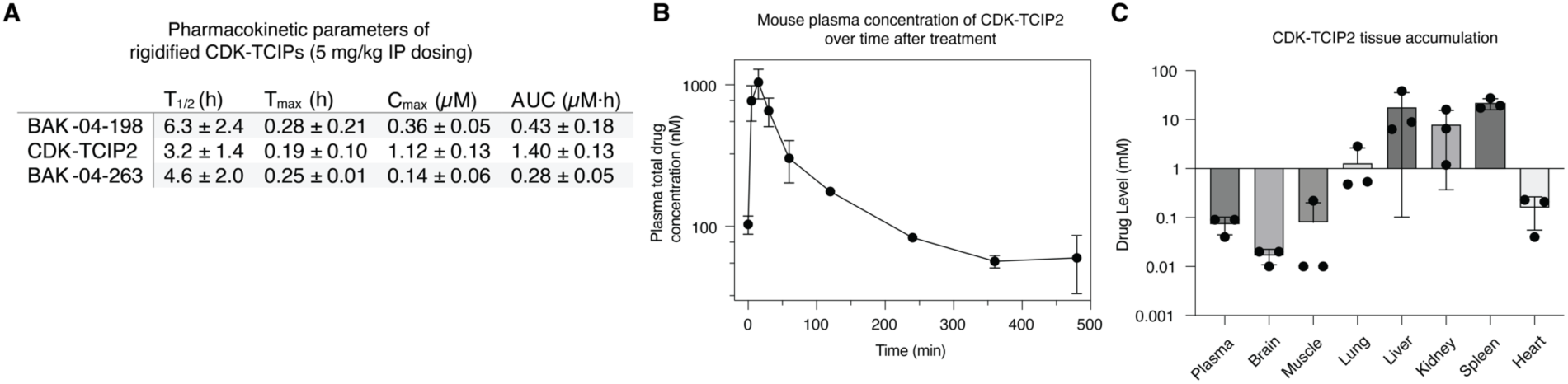
Rigidified CDK-TCIPs exhibit enhanced pharmacokinetic properties. **A.** Pharmacokinetic parameters of rigidified CDK-TCIPs. t_1/2_, half-life; T_max_, time to maximum serum concentration; C_max_, maximum serum concentration; AUC, area under the curve from dosing to last measured concentration; mean±s.d, n=3 mice per compound. **B.** Mouse blood plasma concentration of CDK-TCIPs as function of time after treatment at 5 mg/kg IP dosing. **C.** Tissue drug accumulation after 5-day daily dosing at 5 mg/kg.

