## Supplemental Information for "Borrowing Transcriptional Kinases to Activate Apoptosis"

[#]co-corresponding

Inquiries should be addressed to Gerald R. Crabtree:, Nathanael S. Gray:, or Michael R. Green:

TABLE OF CONTENTS

1. Chemical Synthesis
2. Full Scans of Western Blots
3. Flow Cytometry Gating Strategy
4. Supplemental Table 1: Pharmacology of CDK-TCIPs in DLBCL Cell Lines
5. Supplemental Table 2: List of Abbreviation for Lineage Subtypes in PRISM Assay
6. Supplemental Table 3: Global Proteomics Data

### 1. Chemical Synthesis

GENERAL SYNTHETIC METHODS

Unless otherwise noted, all reagents were purchased from commercial suppliers and used without further purification unless otherwise noted. Reactions were monitored using a Waters Acquity UPLC/MS system (Waters PDA eλ Detector, QDa Detector, Sample manager - FL, Binary Solvent Manager) using Acquity UPLC® BEH C18 column (2.1 x 50 mm, 1.7 µm particle size): solvent gradient = 85% A at 0 min, 1% A at 1.7 min; solvent A = 0.1% formic acid in water; solvent B = 0.1% formic acid in Acetonitrile; flow rate: 0.6 mL/min. Analytical thin layer chromatography (TLC) was performed on Merck silica gel 60 F254 TLC glass plates and analytes were visualized by fluorescence quenching (using 254 nm light) and stained by potassium permanganate solution (KMnO4 (3 g), $K_2CO_3$ (20 g), 5% aq. NaOH (5 mL), water (300 mL)) followed by heating. Purification of reaction products was carried out by flash column chromatography using CombiFlash®Rf with Teledyne Isco RediSep® normal-phase silica flash columns (4 g, 12 g, 24 g, 40 g or 80 g) or preparative RP-HPLC using Waters SunFireTM Prep C18 column (19 x 100 mm, 5 µm particle size) using a gradient of 10-90% methanol in water containing 0.05% trifluoroacetic acid (TFA) over 40 min (45 min run time) at a flow of 40 mL/min. Assayed compounds were isolated and tested as TFA salts and purities of assayed compounds were in all cases greater than 95%, as determined by reverse-phase UPLC analysis. NMR spectra were acquired on a 500 MHz Bruker Avance III spectrometer, operating at the denoted spectrometer frequency given in MHz for the specified nucleus. All experiments were acquired at 298.0 K with a calibrated Bruker Variable Temperature Controller unless otherwise noted. The chemical shifts are reported in parts per million (ppm) and coupling constants (*J*) are given in Hertz (Hz). $^1$H NMR spectra are reported with the solvent resonance as the reference unless noted otherwise ($CDCl_3$ at 7.26 ppm, $CD_3OD$ at 3.31 ppm, DMSO-$d_6$ at 2.50 ppm). Peaks are reported as (s = singlet, d = doublet, t = triplet, q = quartet, m = multiplet or unresolved, br = broad signal, coupling constant(s) in Hz, integration).

SYNTHESIS OF CDK TCIPS

**General Procedure A**

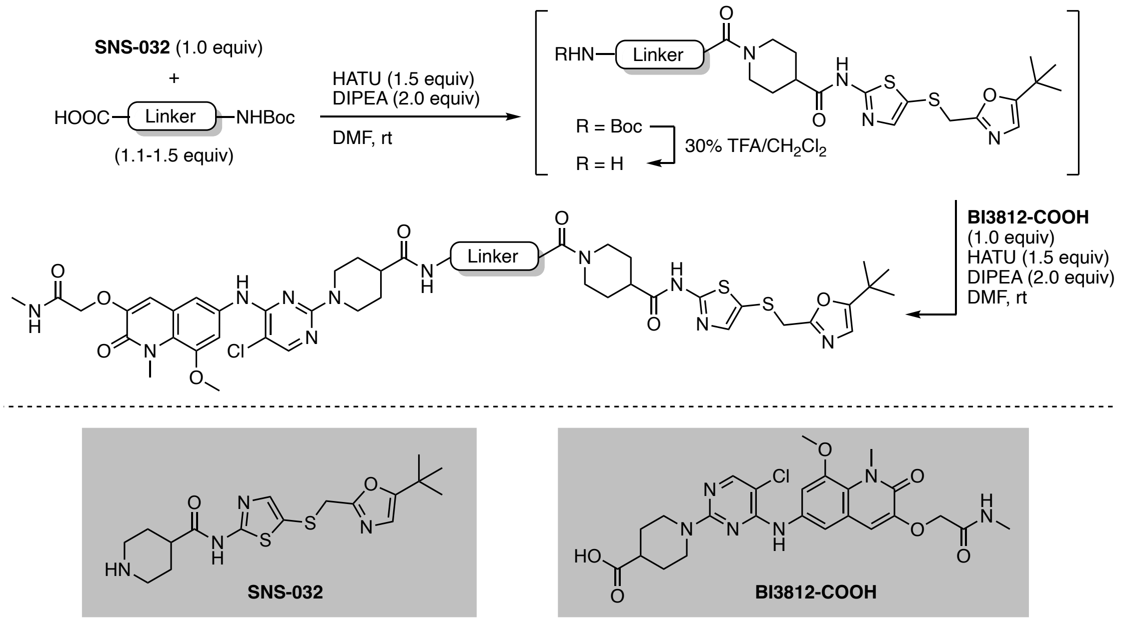


To a solution of **SNS-032**[1] (1.0 equiv) and linker (1.1-1.5 equiv) in DMF (0.2 M) was added HATU (1.5 equiv) and DIPEA (2.0 equiv). The resulting yellow solution was stirred at ambient temperature. Upon full consumption of starting material, as judged by LC-MS and/or TLC analysis, the reaction mixture was diluted with water and EtOAc. After separation of the organic layer, the aqueous phase was extracted with EtOAc (3x) and the combined organic extracts were washed with brine, dried over $Na_2SO_4$, and concentrated under reduced pressure. The resulting intermediate was taken up in 30% v/v TFA in $CH_2Cl_2$ and the resulting solution was stirred at ambient temperature until LC-MS showed quantitative formation of free amine intermediate. All volatiles were removed under reduced pressure. The intermediate was used without further purification, dissolved in DMF (0.2 M), and added to a solution of **BI3812-COOH**[2] (1.0 equiv), HATU (1.5 equiv), and DIPEA (2.0 equiv). The reaction mixture was purified by reverse-phase HPLC to afford the product upon lyophilization.

Synthesis of **BAK-04-014**

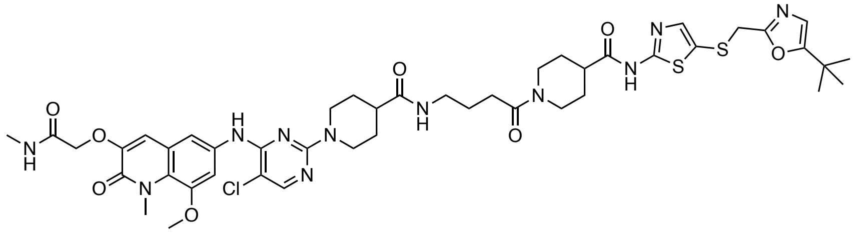

The corresponding compound was prepared following general procedure A using CDK9 inhibitor **SNS-032** (3.4 mg, 9.0 μmol, 1.0 equiv) and 4-((*tert*-butoxycarbonyl)amino)butanoic acid as linker (2.8 mg, 0.014 mmol, 1.5 equiv). Purification by reverse-phase HPLC (15-90% MeOH in water) afforded the product (5.0 mg, 60% yield).

**$^1$H NMR** (500 MHz, DMSO-$d_6$) δ = 12.29 (s, 1H), 9.20 (s, 1H), 8.13 (s, 1H), 7.95 (d, *J* = 4.9 Hz, 1H), 7.83 (t, *J* = 5.6 Hz, 1H), 7.50 (s, 2H), 7.38 (d, *J* = 1.8 Hz, 1H), 7.02 (s, 1H), 6.70 (s, 1H), 4.55 (s, 2H), 4.44 (d, *J* = 13.1 Hz, 2H), 4.37 (d, *J* = 13.0 Hz, 1H), 4.04 (s, 2H), 3.86 (7H, assigned by HSQC), 3.05 (q, *J* = 6.6 Hz, 2H), 2.97 (3H, assigned by HSQC), 2.76 – 2.68 (m, 1H), 2.65 (d, *J* = 4.6 Hz, 3H), 2.58 (t, *J* = 12.8 Hz, 1H), 2.45 – 2.36 (m, 1H), 2.30 (q, *J* = 7.4 Hz, 2H), 1.85 – 1.68 (m, 4H), 1.61 (p, *J* = 7.1 Hz, 2H), 1.58 – 1.47 (m, 2H), 1.40 (2H, assigned by HSQC), 1.17 (d, *J* = 3.6 Hz, 9H). **LC-MS:** m/z 978.48 [M+1]$^+$.

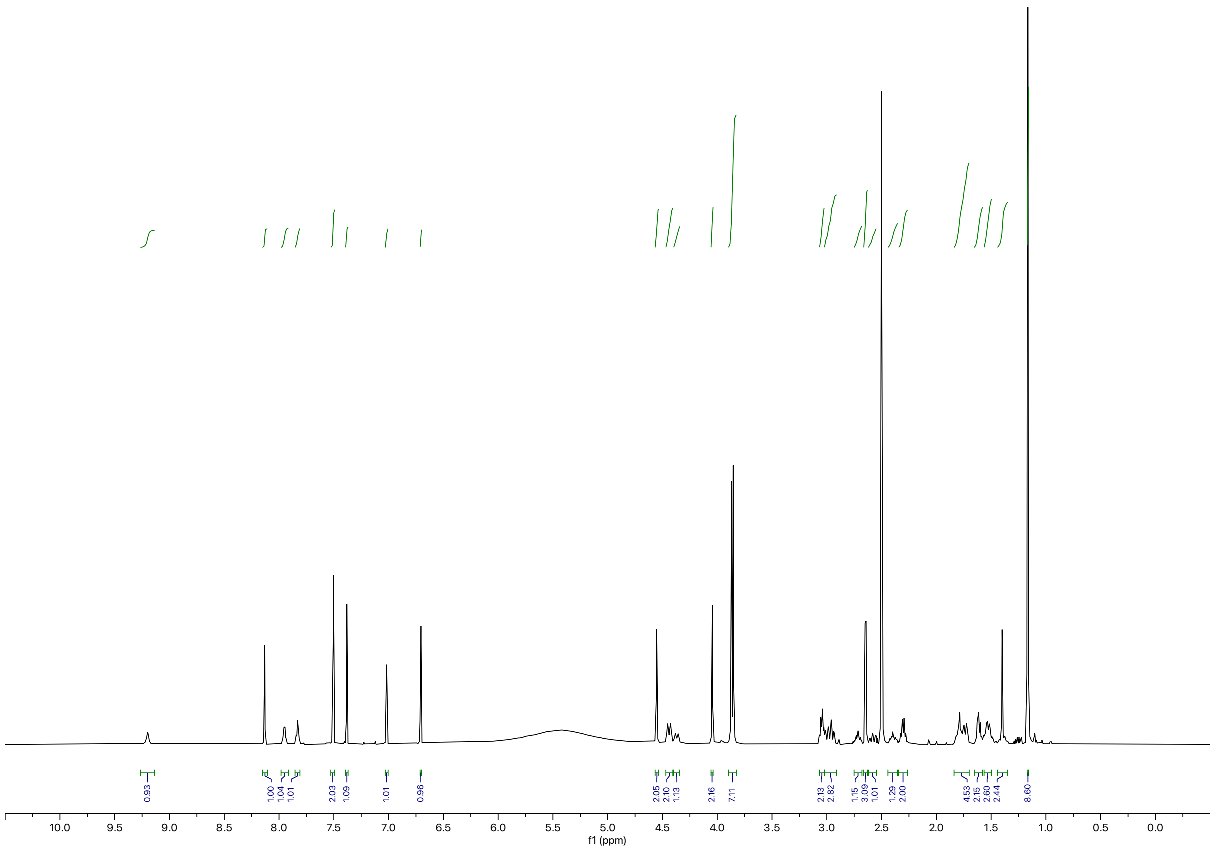

Synthesis of **BAK-04-015**

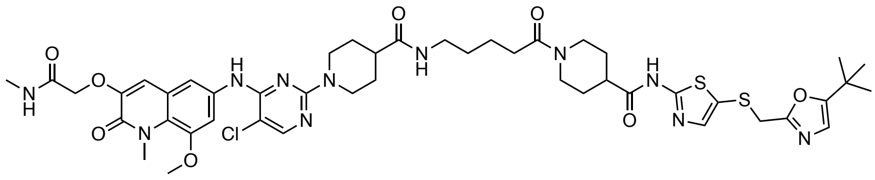

The corresponding compound was prepared following general procedure A using CDK9 inhibitor **SNS-032** (3.0 mg, 8.0 μmol, 1.0 equiv) and 5-((*tert*-butoxycarbonyl)amino)pentanoic acid as linker (2.6 mg, 0.012 mmol, 1.5 equiv). Purification by preparative reverse-phase HPLC (15-90% MeOH in water) afforded the product (4.0 mg, 50% yield).

**$^1$H NMR** (500 MHz, DMSO-d$_6$) δ = 12.30 (s, 1H), 8.99 (s, 1H), 8.10 (s, 1H), 7.96 (d, *J* = 4.8 Hz, 1H), 7.81 (t, *J* = 5.7 Hz, 1H), 7.55 – 7.51 (m, 2H), 7.39 (s, 1H), 7.01 (s, 1H), 6.71 (s, 1H), 4.56 (s, 2H), 4.49 (d, *J* = 13.1 Hz, 2H), 4.38 (d, *J* = 13.2 Hz, 1H), 4.05 (1H, assigned by HSQC), 3.87 (d, *J* = 7.4 Hz, 6H), 3.04 (q, *J* = 7.0 Hz, 3H), 2.93 (t, *J* = 12.6 Hz, 2H), 2.72 (ddt, *J* = 11.4, 7.7, 3.8 Hz, 1H), 2.66 (d, *J* = 4.6 Hz, 3H), 2.58 (t, *J* = 12.5 Hz, 1H), 2.43 – 2.36 (m, 1H), 2.31 (q, *J* = 7.1 Hz, 2H), 1.85 – 1.76 (m, 2H), 1.72 (d, *J* = 12.7 Hz, 2H), 1.57 – 1.44 (6H, assigned by HSQC), 1.44 – 1.36 (m, 2H), 1.17 (s, 9H). **LC-MS:** m/z 992.49 [M+1]$^+$.

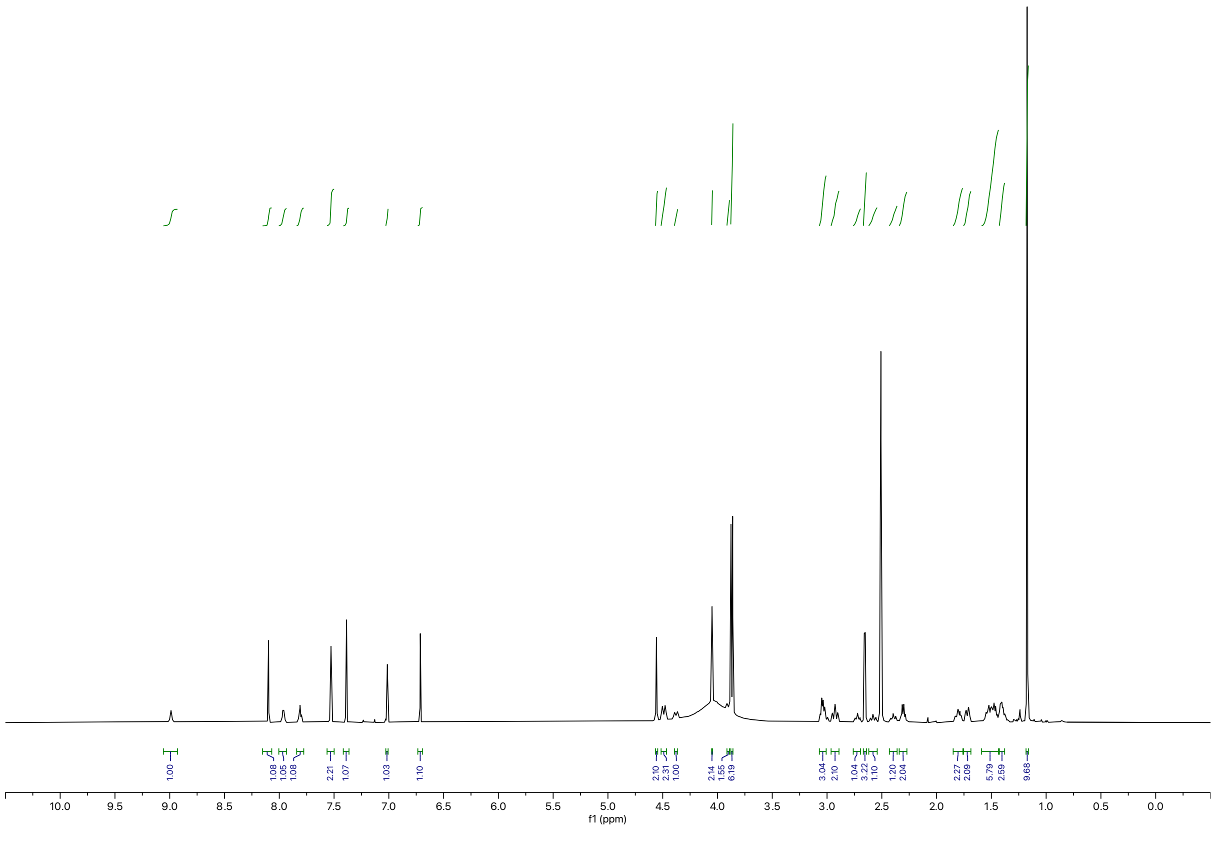

Synthesis of **BAK-04-016**

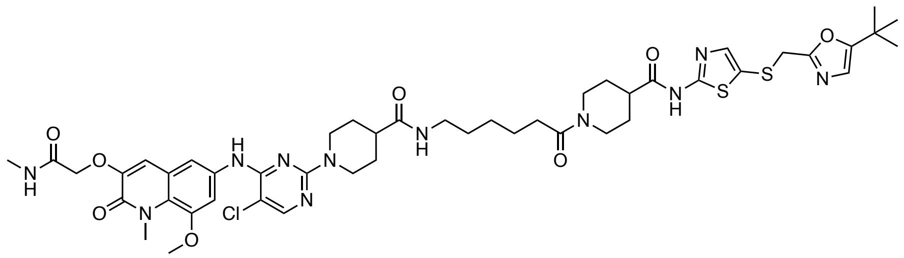

The corresponding compound was prepared following general procedure A using CDK9 inhibitor **SNS-032** (3.0 mg, 8.0 μmol, 1.0 equiv) and 6-((*tert*-butoxycarbonyl)amino)hexanoic acid as linker (2.8 mg, 0.012 mmol, 1.5 equiv). Purification by preparative reverse-phase HPLC (15-90% MeOH in water) afforded the product (3.2 mg, 40% yield).

**$^1$H NMR** (500 MHz, DMSO-d$_6$) δ = 12.29 (s, 1H), 9.00 (s, 1H), 8.09 (d, *J* = 1.4 Hz, 1H), 7.96 (d, *J* = 5.0 Hz, 1H), 7.78 (t, *J* = 5.6 Hz, 1H), 7.52 (d, *J* = 1.2 Hz, 2H), 7.38 (d, *J* = 1.3 Hz, 1H), 7.00 (d, *J* = 1.4 Hz, 1H), 6.70 (d, *J* = 1.4 Hz, 1H), 4.55 (s, 2H), 4.48 (d, *J* = 13.1 Hz, 2H), 4.36 (d, *J* = 13.2 Hz, 1H), 4.04 (d, *J* = 1.4 Hz, 2H), 3.86 (7H, assigned by HSQC), 3.01 (q, *J* = 7.1 Hz, 3H), 2.92 (t, *J* = 12.7 Hz, 2H), 2.71 (t, *J* = 11.1 Hz, 1H), 2.65 (dd, *J* = 4.6, 1.3 Hz, 3H), 2.56 (t, *J* = 12.0 Hz, 1H), 2.38 (t, *J* = 11.5 Hz, 1H), 2.31 – 2.25 (m, 2H), 1.84 – 1.75 (m, 2H), 1.71 (d, *J* = 12.5 Hz, 2H), 1.48 (dd, *J* = 18.5, 10.3 Hz, 4H), 1.38 (4H, assigned by HSQC), 1.26 (dt, *J* = 15.0, 6.8 Hz, 2H), 1.16 (d, *J* = 1.3 Hz, 9H). LC-MS: m/z 1006.59 [M+1]$^+$.

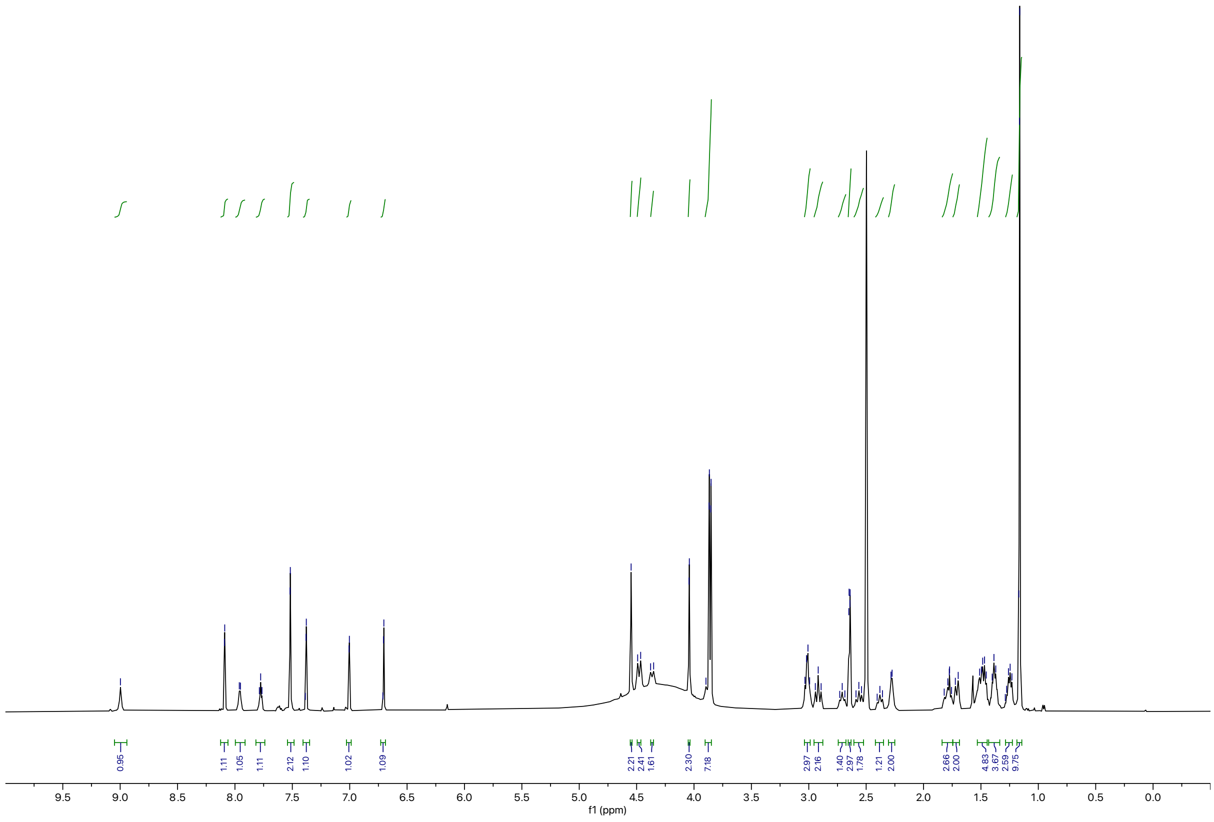

Synthesis of **BAK-04-021 (CDK-TCIP1)**

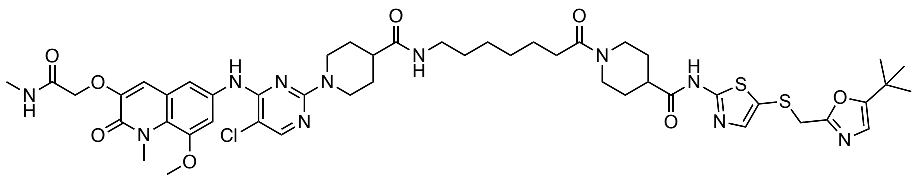

The corresponding compound was prepared following general procedure A using CDK9 inhibitor **SNS-032** (7.6 mg, 0.020 mmol) and 7-((*tert*-butoxycarbonyl)amino)heptanoic acid as linker (5.9 mg, 0.024 mmol, 1.2 equiv). Purification by preparative reverse-phase HPLC (15-90% MeOH in water) afforded the product (4.6 mg, 30% yield).

**$^1$H NMR** (500 MHz, DMSO-$d_6$) δ =12.29 (s, 1H), 8.92 (s, 1H), 8.08 (s, 1H), 7.97 (d, *J* = 4.9 Hz, 1H), 7.77 (t, *J* = 5.6 Hz, 1H), 7.52 (s, 2H), 7.38 (s, 1H), 7.00 (s, 1H), 6.71 (s, 1H), 4.55 (s, 2H), 4.49 (d, *J* = 13.1 Hz, 2H), 4.37 (d, *J* = 13.2 Hz, 1H), 4.04 (s, 2H), 3.86 (d, *J* = 7.3 Hz, 7H), 3.01 (q, *J* = 7.3 Hz, 3H), 2.90 (t, *J* = 12.6 Hz, 2H), 2.75 – 2.67 (m, 1H), 2.65 (d, *J* = 4.6 Hz, 3H), 2.55 (d, *J* = 15.5 Hz, 1H), 2.43 – 2.34 (m, 1H), 2.28 (q, *J* = 7.0 Hz, 2H), 1.80 (t, *J* = 11.4 Hz, 2H), 1.70 (d, *J* = 12.6 Hz, 2H), 1.60 – 1.42 (m, 4H), 1.37 (t, *J* = 7.0 Hz, 4H), 1.30 – 1.22 (m, 4H), 1.16 (s, 9H). **LC-MS:** m/z 1022.44 [M+1]$^+$.

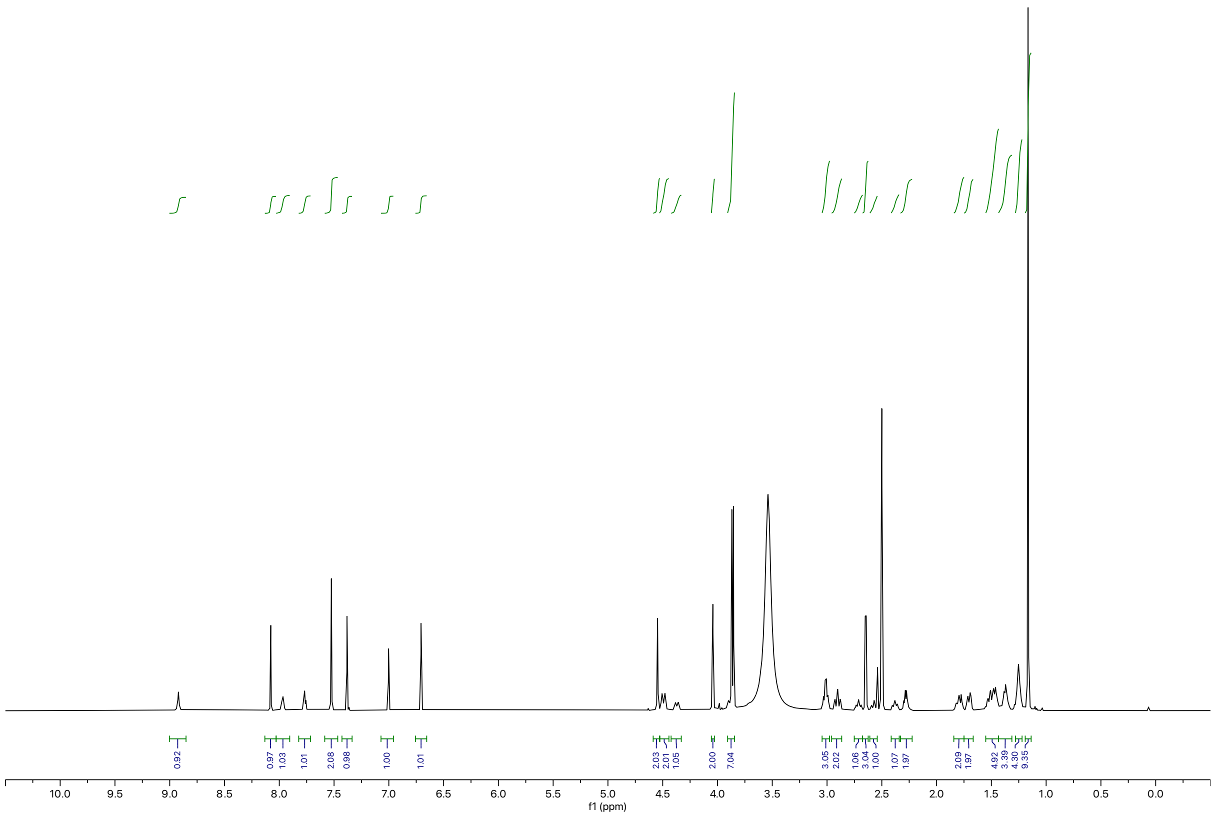

Synthesis of **BAK-04-022**

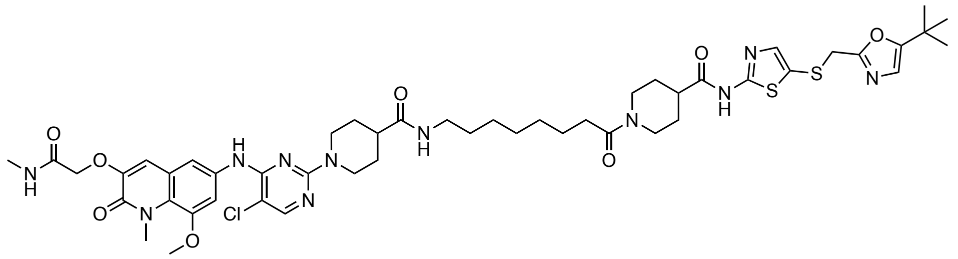

The corresponding compound was prepared following general procedure A using CDK9 inhibitor **SNS-032** (3.8 mg, 0.10 mmol, 1.0 equiv) and 8-((*tert*-butoxycarbonyl)amino)octanoic acid as linker (3.1 mg, 0.012 mmol, 1.2 equiv). Purification by reverse-phase HPLC (15-90% MeOH in water) afforded the product (2.8 mg, 20% yield).

**$^1$H NMR** (500 MHz, DMSO-d$_6$) δ = 12.29 (s, 1H), 8.92 (s, 1H), 8.08 (s, 1H), 7.97 (d, *J* = 4.9 Hz, 1H), 7.77 (t, *J* = 5.6 Hz, 1H), 7.52 (s, 2H), 7.38 (s, 1H), 7.00 (s, 1H), 6.71 (s, 1H), 4.55 (s, 2H), 4.49 (d, *J* = 13.1 Hz, 2H), 4.37 (d, *J* = 13.2 Hz, 1H), 4.04 (s, 2H), 3.86 (d, *J* = 7.3 Hz, 7H), 3.01 (q, *J* = 7.3 Hz, 3H), 2.90 (t, *J* = 12.6 Hz, 2H), 2.75 – 2.67 (m, 1H), 2.65 (d, *J* = 4.6 Hz, 3H), 2.55 (d, *J* = 15.5 Hz, 1H), 2.43 – 2.34 (m, 1H), 2.28 (q, *J* = 7.0 Hz, 2H), 1.80 (t, *J* = 11.4 Hz, 2H), 1.70 (d, *J* = 12.6 Hz, 2H), 1.60 – 1.42 (m, 4H), 1.37 (t, *J* = 7.0 Hz, 4H), 1.30 – 1.22 (m, 6H), 1.16 (s, 9H). **LC-MS:** m/z 1034.55 [M+1]$^+$.

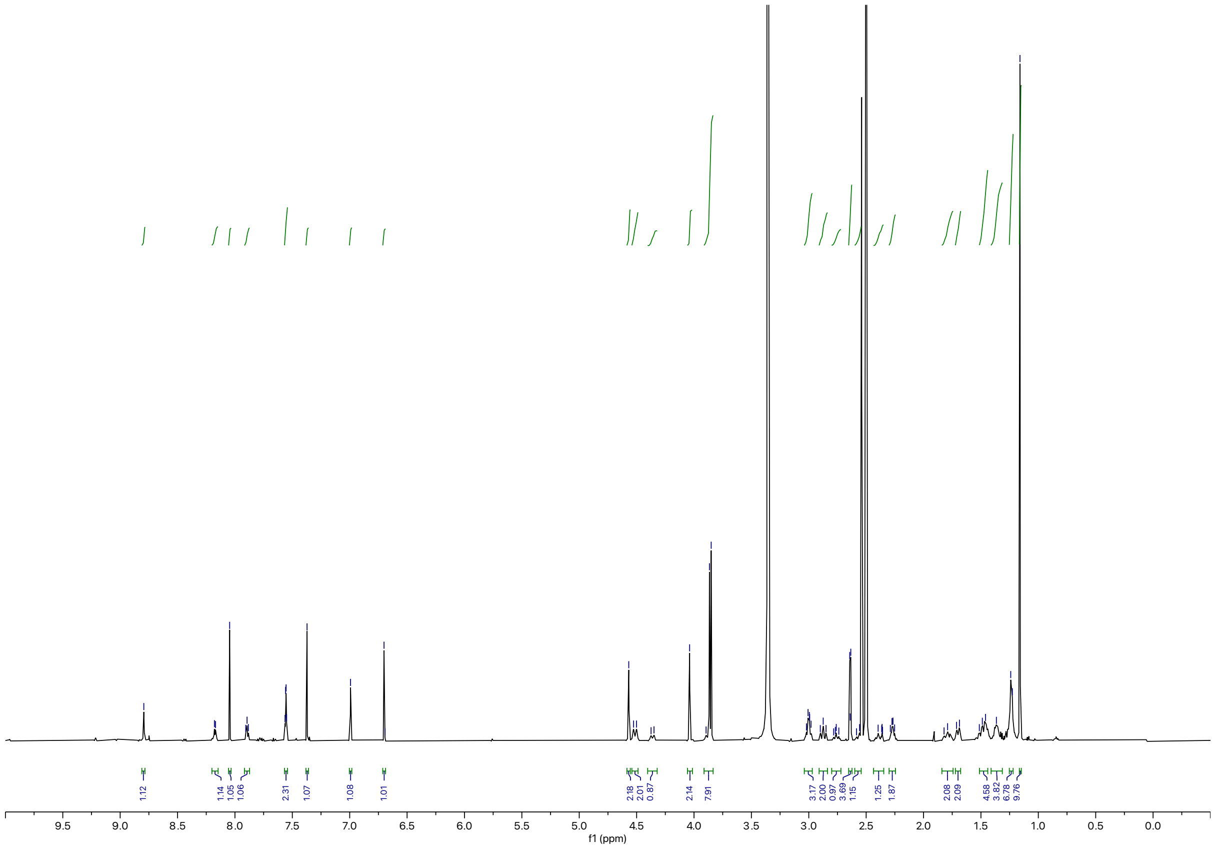

Synthesis of **BAK-04-023**

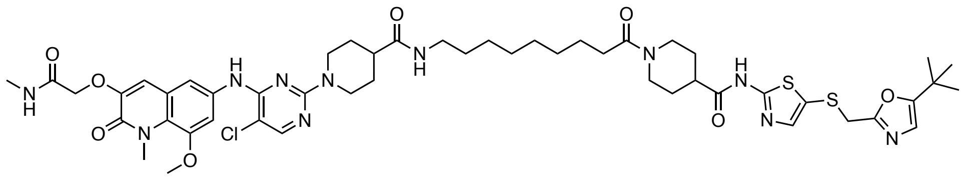

The corresponding compound was prepared following general procedure A using CDK9 inhibitor **SNS-032** (7.6 mg, 0.020 mmol) and 9-((*tert*-butoxycarbonyl)amino)octanoic acid as linker (6.6 mg, 0.024 mmol, 1.2 equiv). Purification by preparative reverse-phase HPLC (15-90% MeOH in water) afforded the product (4.1 mg, 30% yield).

**$^1$H NMR** (500 MHz, DMSO-d$_6$) δ = 12.30 (s, 1H), 9.07 (s, 1H), 8.11 (s, 1H), 7.97 (d, *J* = 4.8 Hz, 1H), 7.77 (t, *J* = 5.6 Hz, 1H), 7.52 (s, 2H), 7.39 (s, 1H), 7.02 (s, 1H), 6.72 (s, 1H), 4.56 (s, 2H), 4.47 (d, *J* = 13.1 Hz, 2H), 4.38 (d, *J* = 13.1 Hz, 1H), 4.05 (s, 2H), 3.91 (d, *J* = 2.0 Hz, 1H), 3.87 (d, *J* = 6.7 Hz, 6H), 3.02 (q, *J* = 7.0 Hz, 3H), 2.94 (t, *J* = 12.5 Hz, 2H), 2.72 (ddt, *J* = 11.4, 7.7, 3.7 Hz, 1H), 2.65 (d, *J* = 4.6 Hz, 3H), 2.62 – 2.53 (m, 1H), 2.43 – 2.36 (m, 1H), 2.28 (p, *J* = 7.6 Hz, 2H), 1.85 – 1.76 (m, 2H), 1.72 (d, *J* = 12.7 Hz, 2H), 1.58 – 1.44 (m, 4H), 1.38 (d, *J* = 7.5 Hz, 4H), 1.25 (s, 8H), 1.17 (s, 9H). **LC-MS:** m/z 1048.47 [M+1]$^+$.

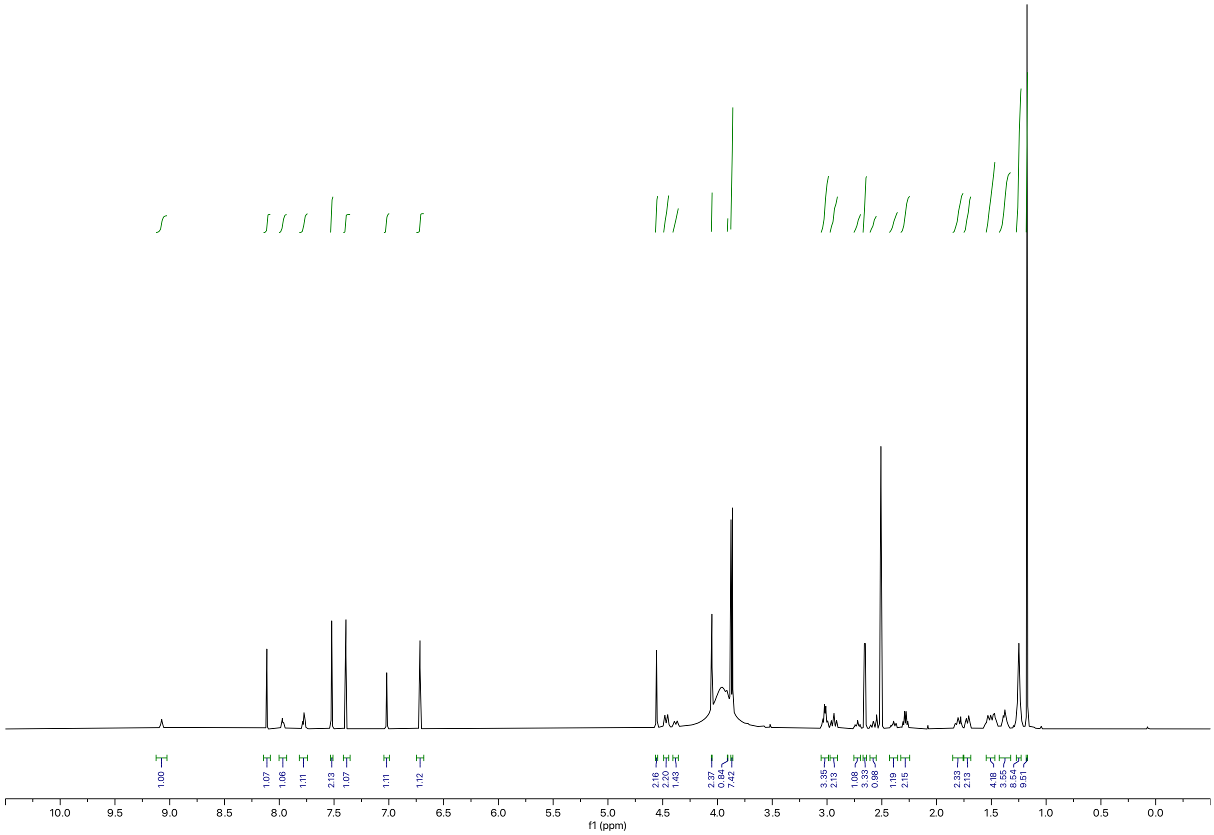

Synthesis of **BAK-04-028**

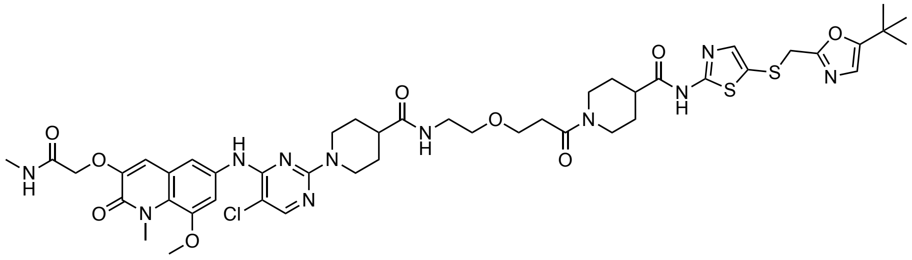

The corresponding compound was prepared following general procedure A using CDK9 inhibitor **SNS-032** (7.2 mg, 0.019 mmol, 1.0 equiv) and *N*-Boc-PEG1-acid as linker (5.3 mg, 0.023 mmol, 1.2 equiv). Purification by preparative reverse-phase HPLC (15-90% MeOH in water) afforded the product (2.7 mg, 14% yield).

**$^1$H NMR** (500 MHz, DMSO-d$_6$) δ = 12.29 (s, 1H), 8.99 (s, 1H), 8.09 (s, 1H), 7.94 (d, *J* = 4.8 Hz, 1H), 7.83 (t, *J* = 5.6 Hz, 1H), 7.52 (s, 2H), 7.38 (s, 1H), 7.00 (s, 1H), 6.70 (s, 1H), 4.55 (s, 2H), 4.47 (d, *J* = 13.1 Hz, 2H), 4.37 (d, *J* = 13.1 Hz, 1H), 4.04 (s, 2H), 3.91 (d, *J* = 13.7 Hz, 1H), 3.86 (d, *J* = 7.6 Hz, 6H), 3.61 (t, *J* = 6.5 Hz, 2H), 3.38 (t, *J* = 5.9 Hz, 2H), 3.18 (q, *J* = 6.0 Hz, 2H), 3.02 (t, *J* = 12.6 Hz, 1H), 2.96 – 2.88 (m, 2H), 2.71 (ddt, *J* = 11.3, 7.7, 3.9 Hz, 1H), 2.65 (d, *J* = 4.7 Hz, 3H), 2.62 – 2.52 (m, 3H), 2.45 – 2.38 (m, 1H), 1.84 – 1.67 (m, 4H), 1.51 (qd, *J* = 13.0, 5.5 Hz, 3H), 1.43 – 1.33 (m, 1H), 1.16 (s, 9H). **LC-MS:** m/z 1008.52 [M+1]$^+$.

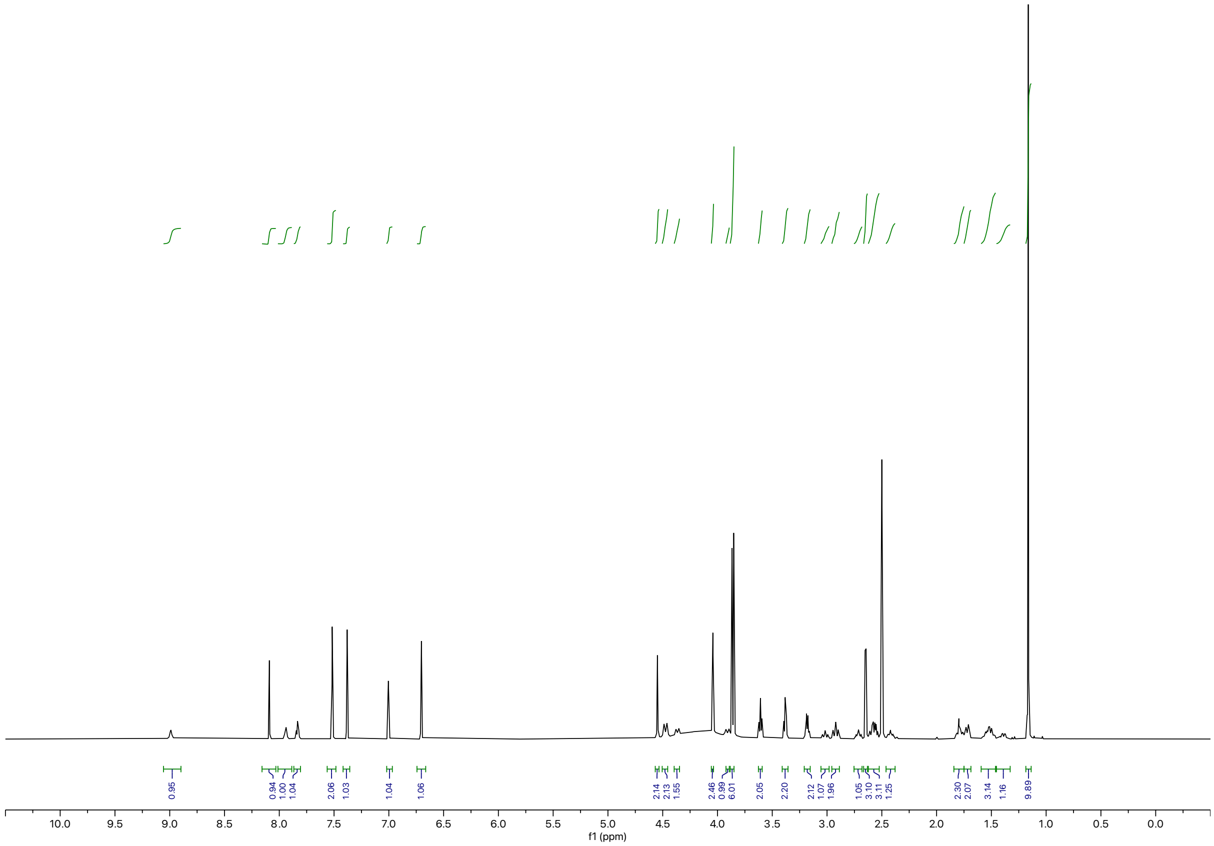

Synthesis of **BAK-04-029**

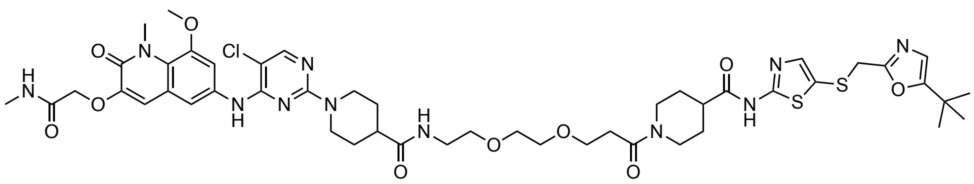

The corresponding compound was prepared following general procedure A using CDK9 inhibitor **SNS-032** (7.7 mg, 0.020 mmol, 1.0 equiv) and *N*-Boc-PEG2-acid as linker (6.7 mg, 0.024 mmol, 1.2 equiv). Purification by preparative reverse-phase HPLC (15-90% MeOH in water) afforded the product (6.5 mg, 40% yield).

**$^1$H NMR** (500 MHz, DMSO-d$_6$) δ = 12.28 (s, 1H), 8.95 (s, 1H), 8.08 (s, 1H), 7.95 (d, *J* = 4.8 Hz, 1H), 7.86 (t, *J* = 5.7 Hz, 1H), 7.52 (s, 2H), 7.38 (s, 1H), 7.00 (s, 1H), 6.70 (s, 1H), 4.55 (s, 2H), 4.48 (d, *J* = 13.1 Hz, 2H), 4.36 (d, *J* = 13.1 Hz, 1H), 4.04 (s, 2H), 3.91 (d, *J* = 13.8 Hz, 1H), 3.86 (d, *J* = 7.4 Hz, 6H), 3.61 (t, *J* = 6.7 Hz, 2H), 3.49 (s, 4H), 3.39 (t, *J* = 5.9 Hz, 2H), 3.18 (q, *J* = 5.9 Hz, 2H), 3.01 (t, *J* = 12.6 Hz, 1H), 2.91 (t, *J* = 12.5 Hz, 2H), 2.71 (tt, *J* = 11.4, 3.8 Hz, 1H), 2.65 (d, *J* = 4.7 Hz, 3H), 2.62 – 2.52 (m, 3H), 2.46 – 2.37 (m, 1H), 1.79 (m, 2H), 1.71 (d, *J* = 12.8 Hz, 3H), 1.62 – 1.45 (m, 2H), 1.44 – 1.34 (m, 1H), 1.16 (s, 9H). **LC-MS:** m/z 1052.58 [M+1]$^+$.

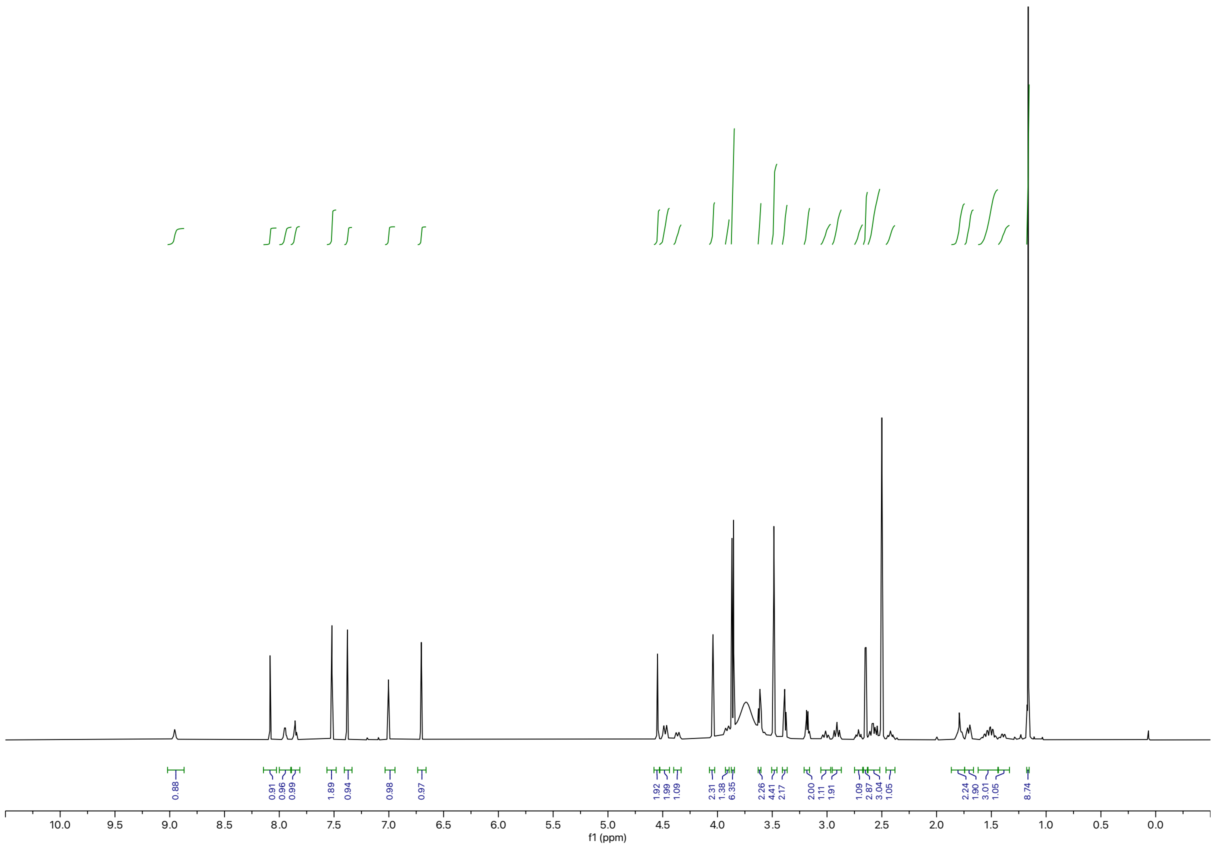

Synthesis of **BAK-04-030**

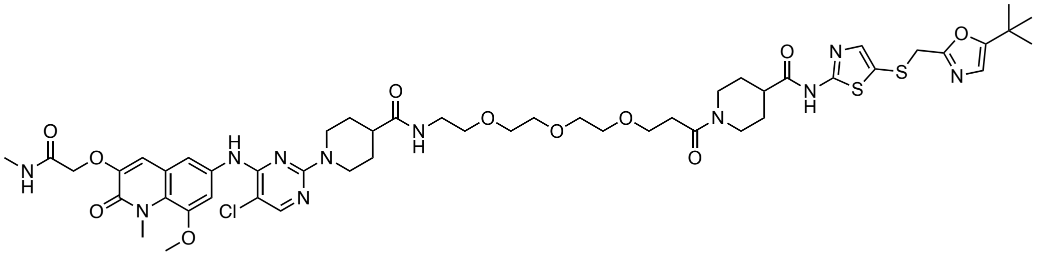

The corresponding compound was prepared following general procedure A using CDK9 inhibitor **SNS-032** (7.6 mg, 0.020 mmol, 1.0 equiv) and *N*-Boc-PEG3-acid as linker (7.7 mg, 0.024 mmol, 1.2 equiv). Purification by preparative reverse-phase HPLC (15-90% MeOH in water) afforded the product (3.6 mg, 20% yield).

**$^1$H NMR** (500 MHz, DMSO-d$_6$) δ = 12.28 (s, 1H), 9.21 (s, 1H), 8.13 (s, 1H), 7.95 (d, *J* = 4.8 Hz, 1H), 7.87 (t, *J* = 5.7 Hz, 1H), 7.50 (s, 2H), 7.38 (s, 1H), 7.02 (s, 1H), 6.70 (s, 1H), 4.55 (s, 2H), 4.42 (d, *J* = 13.2 Hz, 2H), 4.36 (d, *J* = 13.5 Hz, 1H), 4.04 (s, 2H), 3.92 (d, *J* = 13.9 Hz, 1H), 3.86 (d, *J* = 5.8 Hz, 6H), 3.60 (t, *J* = 6.7 Hz, 2H), 3.53 – 3.44 (m, 8H), 3.39 (t, *J* = 5.9 Hz, 2H), 3.18 (q, *J* = 5.9 Hz, 2H), 2.99 (dt, *J* = 25.9, 12.8 Hz, 3H), 2.71 (ddt, *J* = 11.4, 7.6, 3.7 Hz, 1H), 2.65 (d, *J* = 4.7 Hz, 3H), 2.62 – 2.52 (m, 3H), 2.46 – 2.39 (m, 1H), 1.79 (s, 2H), 1.72 (d, *J* = 12.8 Hz, 2H), 1.60 – 1.46 (m, 3H), 1.39 (q, *J* = 11.6 Hz, 1H), 1.16 (s, 9H). **LC-MS:** m/z 1096.55 [M+1]$^+$.

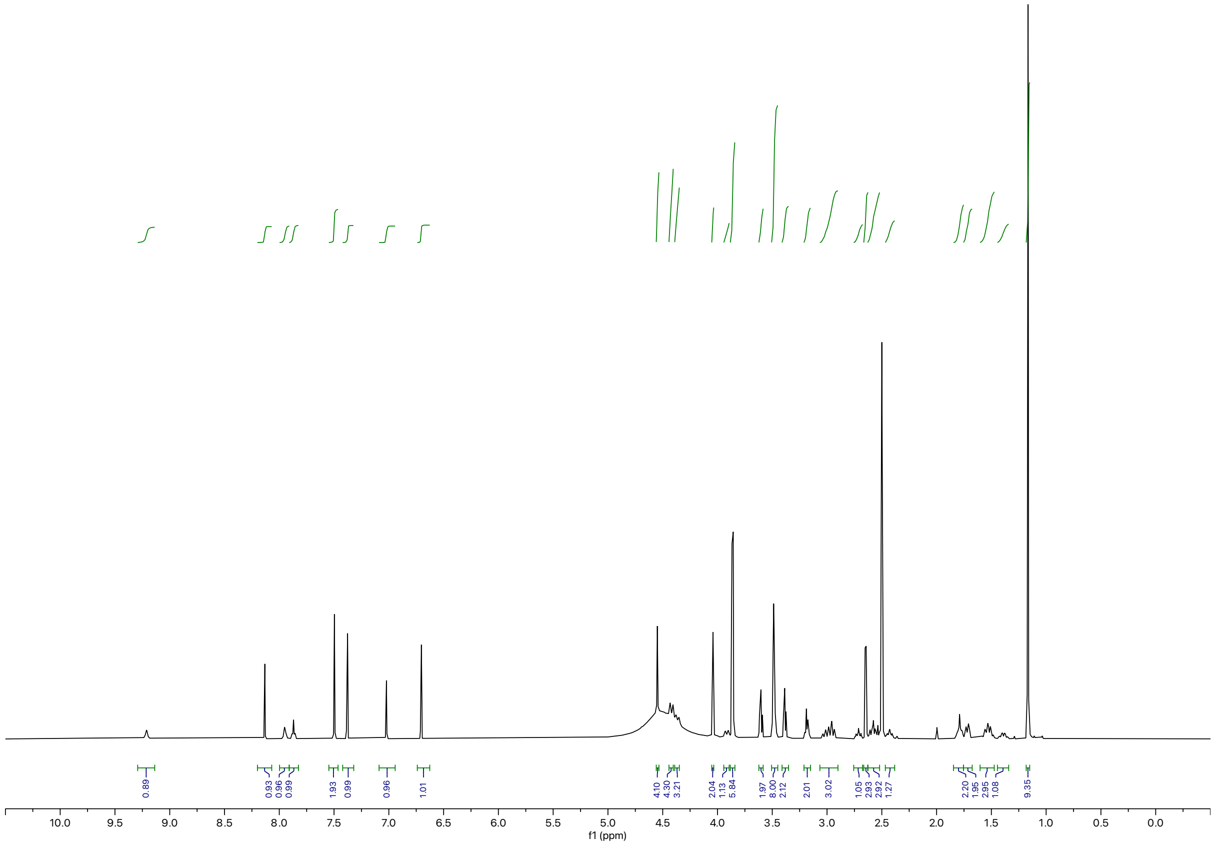

**Negative Controls**

Synthesis of **BAK-04-054 (Neg1)**

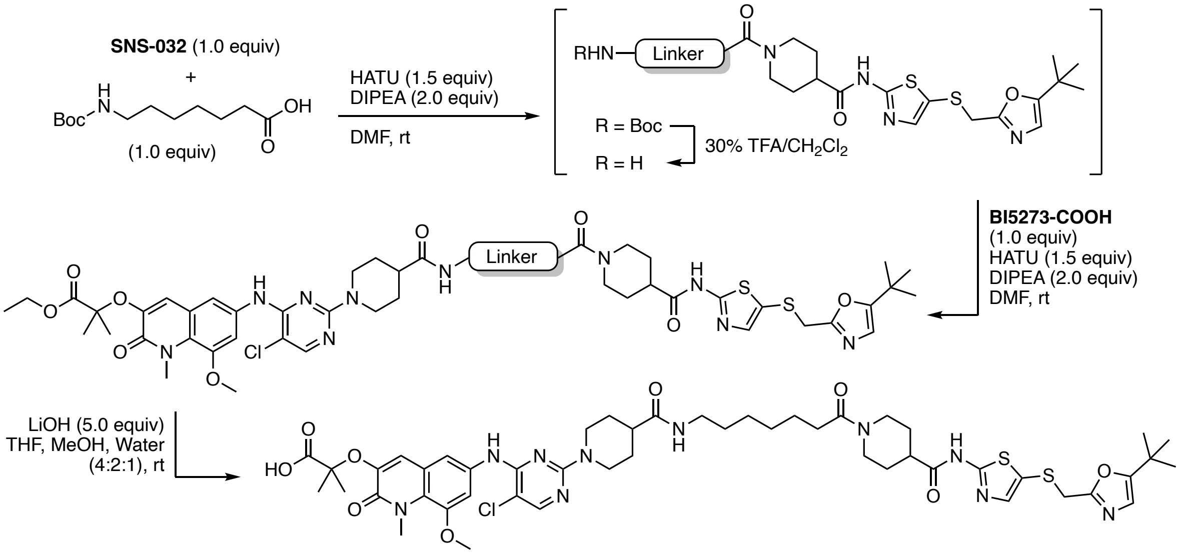


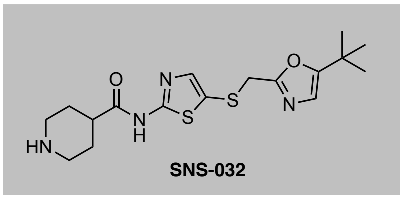


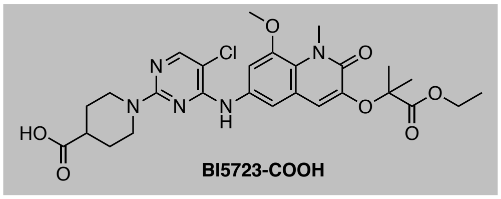


To a solution of **SNS-032** (3.6 mg, 9.4 μmol 1.0 equiv) and 7-((*tert*-butoxycarbonyl)amino)heptanoic acid (2.3 mg, 9.4 μmol 1.0 equiv) in DMF (0.2 M) was added HATU (5.4 mg, 0.014 mmol, 1.5 equiv) and DIPEA (3.3 μL, 0.019 mmol, 2.0 equiv). The resulting yellow solution was stirred at ambient temperature for 2 h. Upon full consumption of starting material, as judged by LC-MS and/or TLC analysis, the reaction mixture was diluted with water and EtOAc. After separation of the organic layer, the aqueous phase was extracted with EtOAc (3x) and the combined organic extracts were washed with brine, dried over $Na_2SO_4$, and concentrated under reduced pressure. The resulting intermediate was taken up in 30% v/v TFA in $CH_2Cl_2$ and the resulting solution was stirred at ambient temperature until LC-MS showed quantitative formation of free amine intermediate. All volatiles were removed under reduced pressure. The intermediate was used without further purification, dissolved in DMF (0.2 M), and added to a solution of **BI5723-COOH**[2] (5.4 mg, 9.4 μmol, 1.0 equiv), HATU (5.4 mg, 0.014 mmol, 1.5 equiv), and DIPEA (3.3 μL, 0.019 mmol, 2.0 equiv). After aqueous workup and concentration of the organic extracts under reduced pressure, the residue was taken up in THF/methanol/water (4:2:1 ratio) (0.2 M) and LiOH (1.0 mg, 0.05 mmol, 5.0 equiv) was added. The reaction was stirred at ambient temperature until LC-MS showed quantitative formation of the carboxylic acid. Purification by preparative reverse-phase HPLC (15-90% MeOH in water) afforded the product as off-white solid (5.4 mg, 55% yield).

**$^1$H NMR** (500 MHz, DMSO-d$_6$) δ =12.29 (s, 1H), 8.92 (s, 1H), 8.08 (s, 1H), 7.97 (d, *J* = 4.9 Hz, 1H), 7.77 (t, *J* = 5.6 Hz, 1H), 7.52 (s, 2H), 7.38 (s, 1H), 7.00 (s, 1H), 6.71 (s, 1H), 4.55 (s, 2H), 4.49 (d, *J* = 13.1 Hz, 2H), 4.37 (d, *J* = 13.2 Hz, 1H), 4.04 (s, 2H), 3.86 (d, *J* = 7.3 Hz, 7H), 3.01 (q, *J* = 7.3 Hz, 3H), 2.90 (t, *J* = 12.6 Hz, 2H), 2.75 – 2.67 (m, 1H), 2.65 (d, *J* = 4.6 Hz, 3H), 2.55 (d, *J* = 15.5 Hz, 1H), 2.43 – 2.34 (m, 1H), 2.28 (q, *J* = 7.0 Hz, 2H), 1.80 (t, *J* = 11.4 Hz, 2H), 1.70 (d, *J* = 12.6 Hz, 2H), 1.60 – 1.42 (m, 4H), 1.37 (t, *J* = 7.0 Hz, 4H), 1.30 – 1.22 (m, 4H), 1.16 (s, 9H). **LC-MS:** m/z 1035.53 [M+1]$^+$.

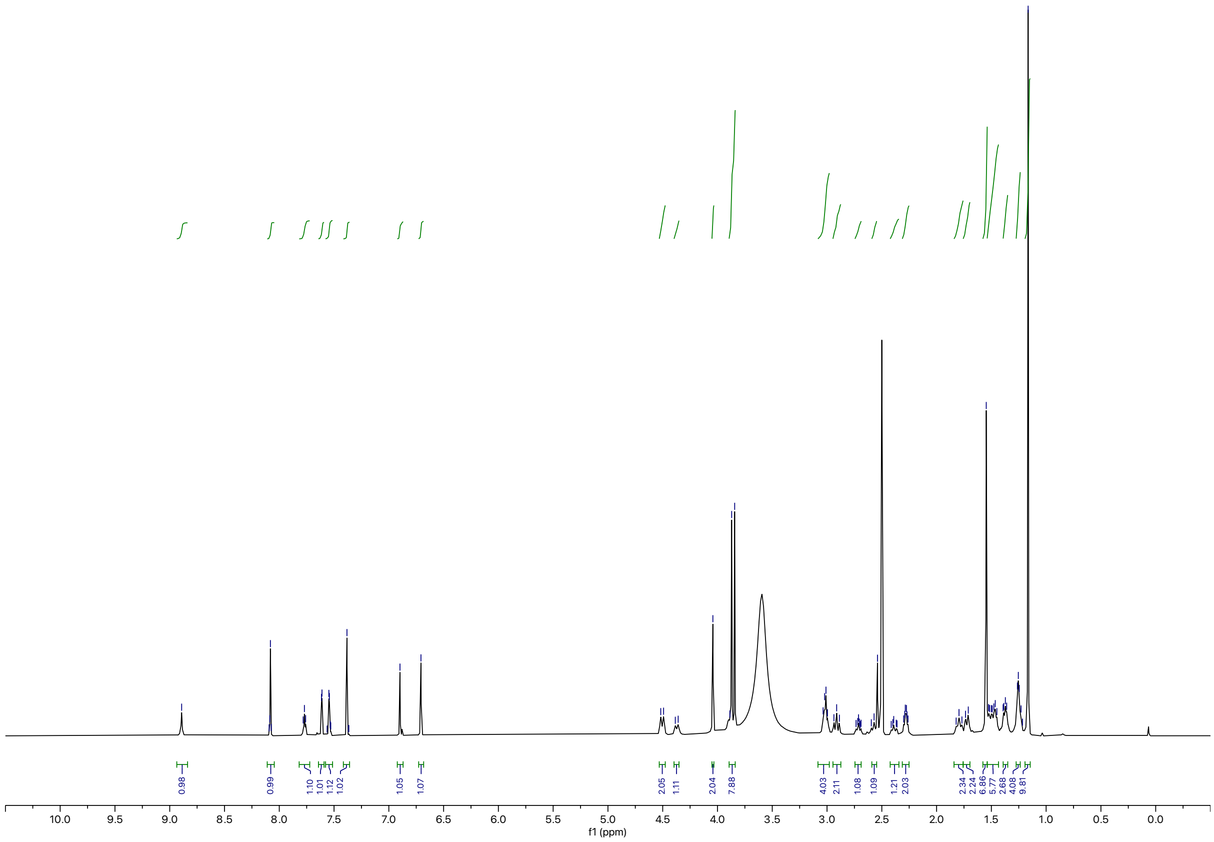

Synthesis of **SNS-032-Me-Boc** Intermediate

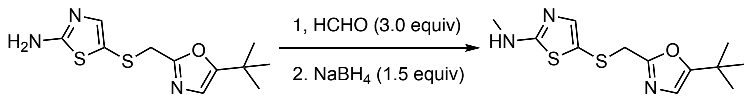


5-(((5-(*tert*-butyl)oxazol-2-yl)methyl)thio)-*N*-methylthiazol-2-amine: To a solution of 5-(((5-(*tert*-butyl)oxazol-2-yl)methyl)thio)thiazol-2-amine[3] (0.065 g, 0.24 mmol, 1.0 equiv) in MeOH (0.5 M), formaldehyde (0.060 mL, 37% wt, 0.72 mmol, 3.0 equiv) was added. The reaction was sealed and heated to 40 °C until quantitative formation of the imine intermediate was observed. The reaction mixture was then concentrated under reduced pressure. The residue was redissolved in MeOH (0.2 M) and $NaBH_4$ (0.014 g, 0.36 mmol, 1.5 equiv) was added. Upon reaction completion as judged by LC-MS, the mixture was concentrated under reduced pressure and directly purified by flash column chromatography on silica (0-20% $CH_2Cl_2$ in MeOH). To afford product as a orange solid (0.055 g, 80% yield). $^1$**H NMR** (500 MHz, MeOD) δ = 6.92 (s, 1H), 6.68 (s, 1H), 3.92 (s, 2H), 2.87 (s, 3H), 1.27 (s, 9H). **LC-MS:** m/z 284.15 $[M+1]^+$.

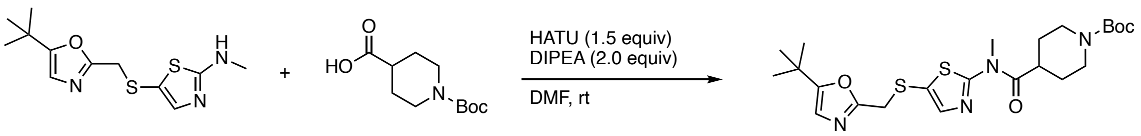


*tert*-butyl-4-((5-(((5-(*tert*-butyl)oxazol-2-yl)methyl)thio)thiazol-2-yl)(methyl)carbamoyl)piperidine-1-carboxylate (SNS-032-Me-Boc): To a solution of 1-(*tert*-butoxycarbonyl)piperidine-4-carboxylic acid (0.044 g, 0.19 mmol, 1.0 equiv), HATU (0.11 g, 0.29 mmol, 1.5 equiv), and DIPEA (0.070 mL, 0.38 mmol, 2.0 equiv) in DMF (0.2 M), 5-(((5-(*tert*-butyl)oxazol-2-yl)methyl)thio)-*N*-methylthiazol-2-amine (0.055 g, 0.19 mmol, 1.0 equiv) was added. The resulting yellow solution was stirred at ambient temperature for 2 h. Upon reaction completion as judged by LC-MS, the reaction was diluted with water (5 mL) and the aqueous layer was extracted with EtOAc (3x5 mL). The combined organic extracts were washed with water (2x3 mL) and brine (2x3 mL), concentrated under reduced pressured and the resulting crude material was purified by flash column chromatography on silica (0-100% EtOAc in hexanes). The product was obtained as a yellow oil (0.085 mg, 89% yield). $^1$**H NMR** (500 MHz, $CDCl_3$) δ = 7.36 (s, 1H), 6.56 (s, 1H), 4.17 (s, 1H), 3.91 (s, 2H), 3.69 (s, 3H), 2.95 – 2.74 (m, 4H), 1.78 (td, *J* = 10.1, 4.2 Hz, 4H), 1.45 (s, 9H), 1.23 (s, 9H). **LC-MS:** m/z 495.30 $[M+1]^+$.

Synthesis of **BAK-04-066 (Neg2)**

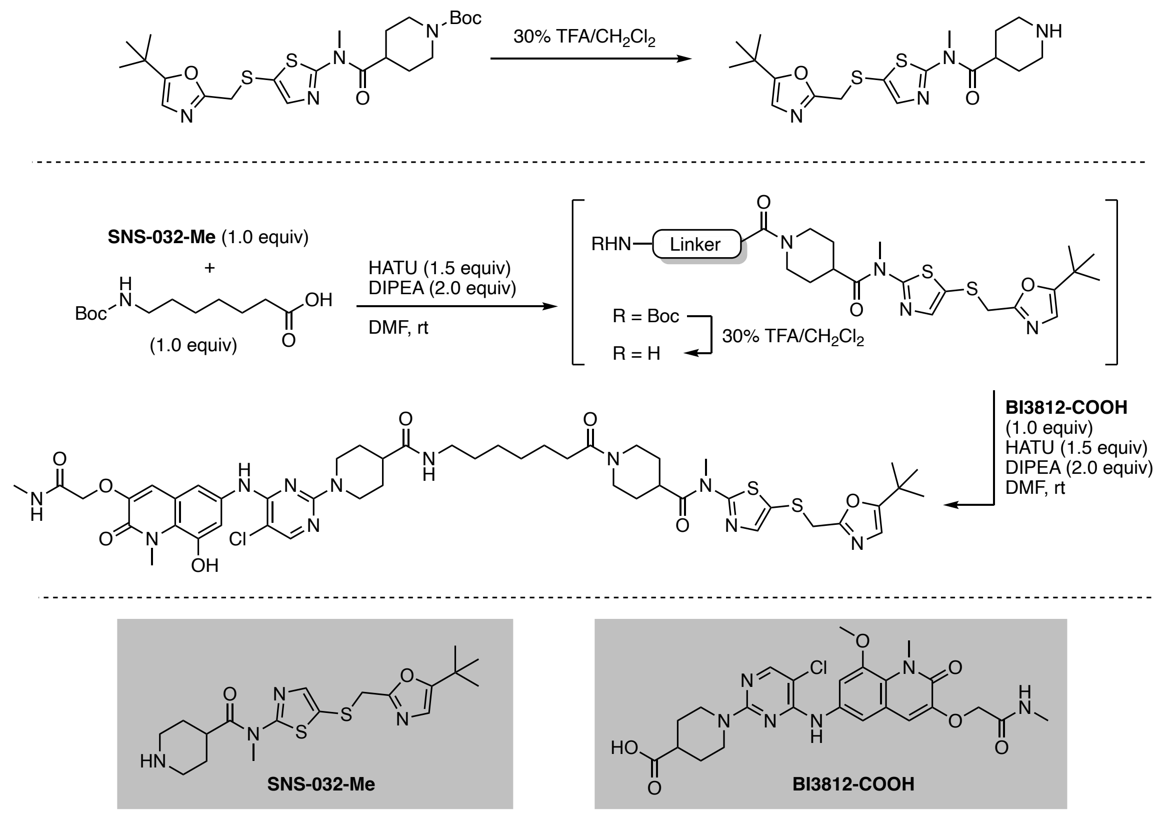


**SNS-032-Me-Boc** (0.016 g, 0.033 mmol 1.0 equiv) was dissolved in 30% v/v TFA in $CH_2Cl_2$. Upon quantitative formation of the free amine intermediate as indicated by LC-MS, all volatiles were removed under reduced pressure and the and residue was dissolved in DMF (0.2 M). 7-((*tert*-butoxycarbonyl)amino)heptanoic acid (7.4 mg, 0.033 mmol, 1.0 equiv), HATU (0.019 g, 0.050 mmol, 1.5 equiv), and DIPEA (0.012 mL, 0.068 mmol, 2.0 equiv) were added. The resulting yellow solution was stirred at ambient temperature for 2 h. Upon full consumption of starting material, as judged by LC-MS and TLC analysis, the reaction mixture was diluted with water and EtOAc. After separation of the organic layer, the aqueous phase was extracted with EtOAc (3x) and the combined organic extracts were washed with brine, dried over $Na_2SO_4$, and concentrated under reduced pressure. The resulting intermediate was taken up in 30% v/v TFA in $CH_2Cl_2$ and the resulting solution was stirred at ambient temperature until LC-MS showed quantitative formation of free amine intermediate. All volatiles were removed under reduced pressure. The intermediate was used without further purification, dissolved in DMF (0.2 M), and added to a solution of **BI3812-COOH** (0.018 g, 0.033 mmol, 1.0 equiv), HATU (0.019 g, 0.050 mmol, 1.5 equiv), and DIPEA (0.012 mL, 0.066 mmol, 2.0 equiv) in DMF (0.2 mL). Purification by preparative reverse-phase HPLC (15-90% MeOH in water) afforded the product as a white solid (5.0 mg, 10% yield).

**$^1$H NMR** (500 MHz, DMSO-$d_6$) δ = 8.86 (s, 1H), 8.06 (s, 1H), 8.01 (d, *J* = 5.0 Hz, 1H), 7.77 (t, *J* = 5.6 Hz, 1H), 7.53 (s, 2H), 7.45 (s, 1H), 7.00 (s, 1H), 6.70 (s, 1H), 4.54 (s, 2H), 4.51 (d, *J* = 13.3 Hz, 2H), 4.39 (d, *J*

= 13.1 Hz, 1H), 4.04 (s, 2H), 3.86 (d, *J* = 7.2 Hz, 7H), 3.67 (s, 3H), 3.27 – 3.19 (m, 1H), 3.11 (t, *J* = 12.8 Hz, 1H), 3.01 (m, 2H), 2.93 – 2.85 (m, 2H), 2.65 (m, 4H), 2.42 – 2.35 (m, 1H), 2.29 (q, *J* = 7.3 Hz, 2H), 1.82 (br, 2H), 1.73 – 1.66 (m, 2H), 1.48 (ddt, *J* = 19.2, 14.4, 8.5 Hz, 4H), 1.37 (t, *J* = 7.5 Hz, 4H), 1.25 (dq, *J* = 10.8, 3.7 Hz, 4H), 1.16 (s, 9H). **LC-MS:** m/z 1034.63 $[M+1]^+$.

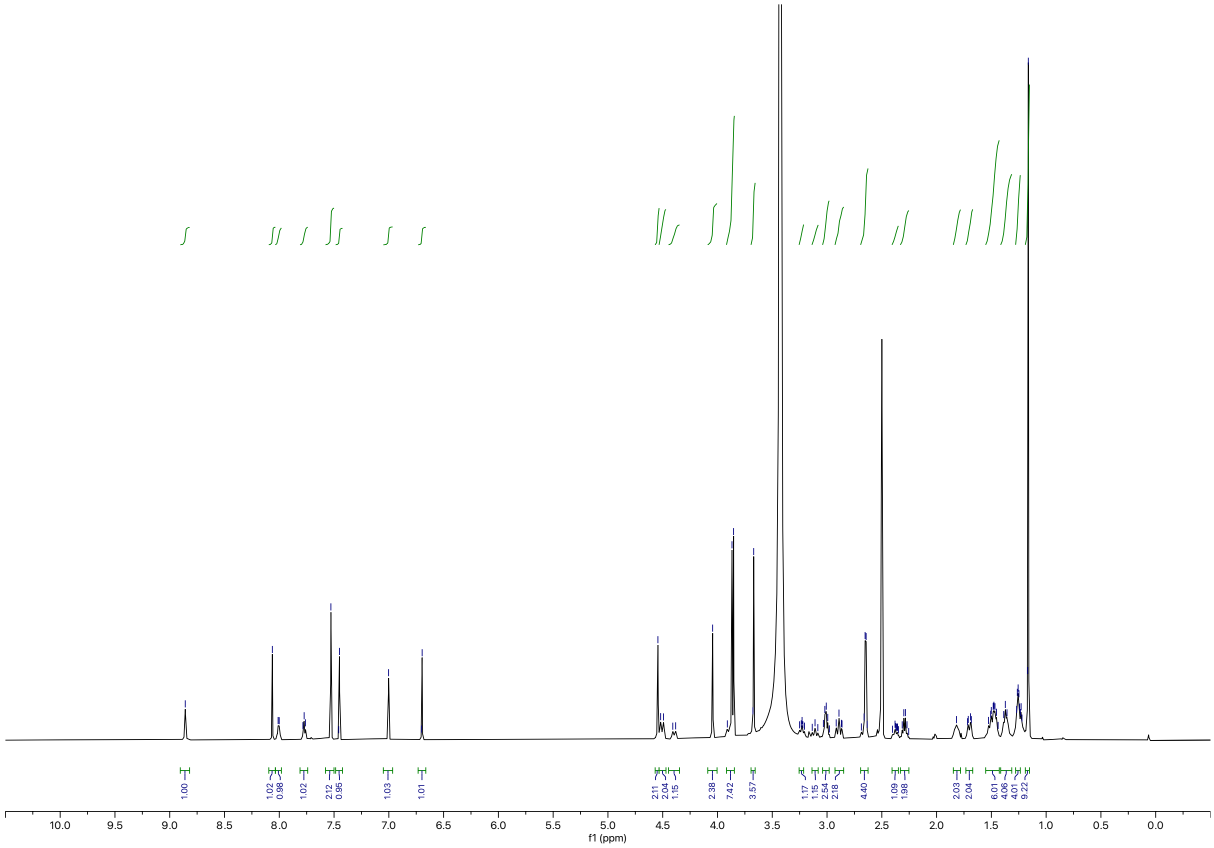

Synthesis of **RCS-03-036**

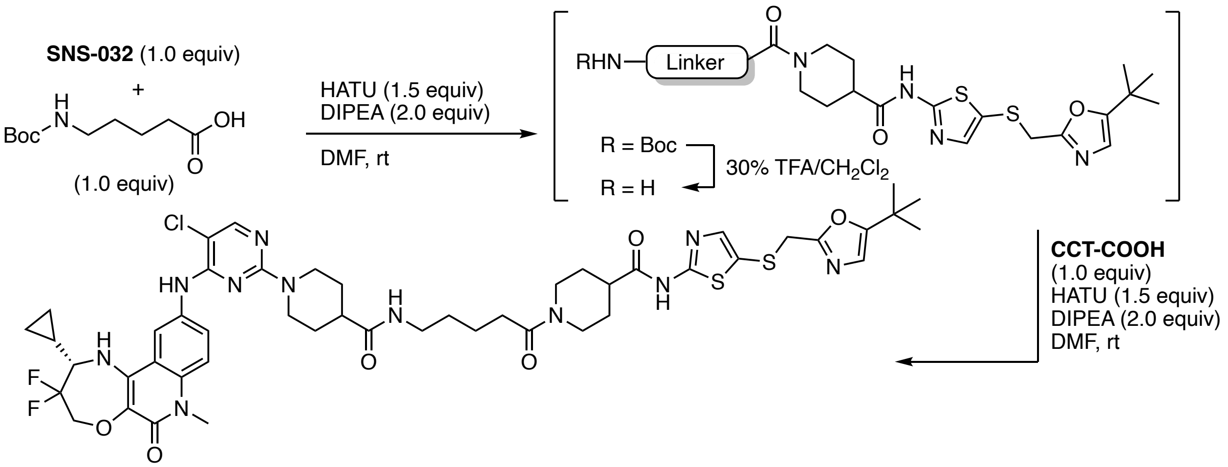


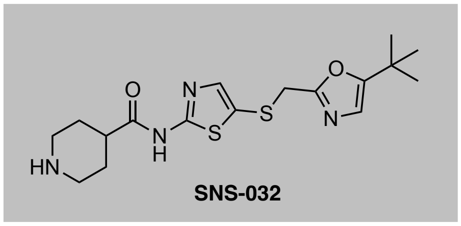


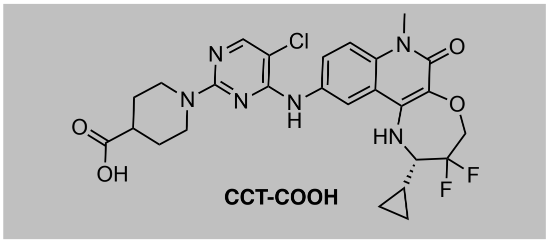


To a solution of **SNS-032** (5.0 mg, 0.013 mmol, 1.0 equiv) and 5-((*tert*-butoxycarbonyl)amino)pentanoic acid (2.9 mg, 0.013 mmol, 1.0 equiv) in DMF (0.2 M) was added HATU (7.5 mg, 0.020 mmol, 1.5 equiv) and DIPEA (4.6 μL, 0.026 mmol, 2.0 equiv). The resulting yellow solution was stirred at ambient temperature for 2 h. Upon full consumption of starting material, as judged by LC-MS and TLC analysis, the reaction mixture was diluted with water and EtOAc. After separation of the organic layer, the aqueous phase was extracted with EtOAc (3x) and the combined organic extracts were washed with brine, dried over $Na_2SO_4$, and concentrated under reduced pressure. The resulting intermediate was taken up in 30% v/v TFA in $CH_2Cl_2$ and the resulting solution was stirred at ambient temperature until LC-MS showed quantitative formation of free amine intermediate. All volatiles were removed under reduced pressure. The intermediate was used without further purification, dissolved in DMF (0.1 mL), and added to a solution of **CCT-COOH** (7.4 mg, 0.013 mmol, 1.0 equiv), HATU (7.5 mg, 0.020 mmol, 1.5 equiv), and DIPEA (4.6 μL, 0.026 mmol, 2.0 equiv) in DMF (0.2 mL). After stirring at ambient temperature for 30 min, purification by reverse-phase HPLC (15-90% MeOH in water) afforded the product as white solid (6.2 mg, 46% yield).

**$^1$H NMR** (500 MHz, DMSO-$d_6$) δ = 12.29 (s, 1H), 9.40 (s, 1H), 8.17 (d, *J* = 2.3 Hz, 1H), 8.14 (s, 1H), 7.78 (t, *J* = 5.7 Hz, 1H), 7.70 (dd, *J* = 9.1, 2.2 Hz, 1H), 7.46 (d, *J* = 9.1 Hz, 1H), 7.38 (s, 1H), 6.71 (s, 1H), 6.22 (s, 1H), 4.39 (dtd, *J* = 49.0, 15.4, 8.6 Hz, 5H), 4.04 (2H, assigned by HSQC), 3.86 (1H, assigned by HSQC), 3.56 (s, 3H), 3.22 (dd, *J* = 11.2, 6.1 Hz, 1H), 3.02 (q, *J* = 6.6 Hz, 3H), 2.89 (t, *J* = 12.4 Hz, 2H), 2.75 – 2.67 (m, 1H), 2.57 (t, *J* = 12.2 Hz, 1H), 2.42 – 2.24 (m, 3H), 1.80 (t, *J* = 12.8 Hz, 2H), 1.66 (d, *J* = 12.9 Hz, 2H), 1.59 – 1.42 (m, 6H), 1.42 – 1.27 (m, 2H), 1.34 – 1.27 (m, 1H), 1.17 (s, 9H), 0.75 – 0.65 (m, 1H), 0.55 – 0.46 (m, 2H), 0.33 (q, *J* = 5.9 Hz, 1H). **LC-MS:** m/z 1022.37 $[M+1]^+$.

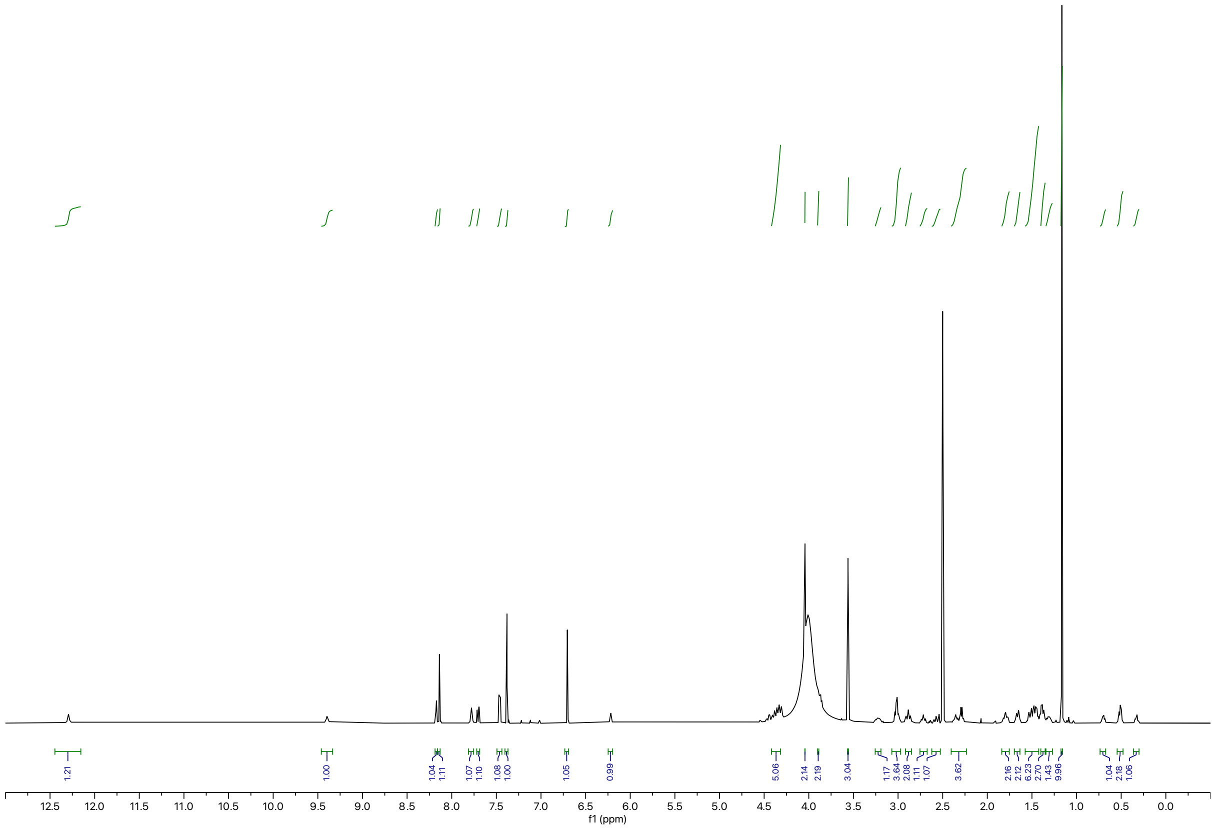
f1 (ppm)

Synthesis of **RCS-03-037**

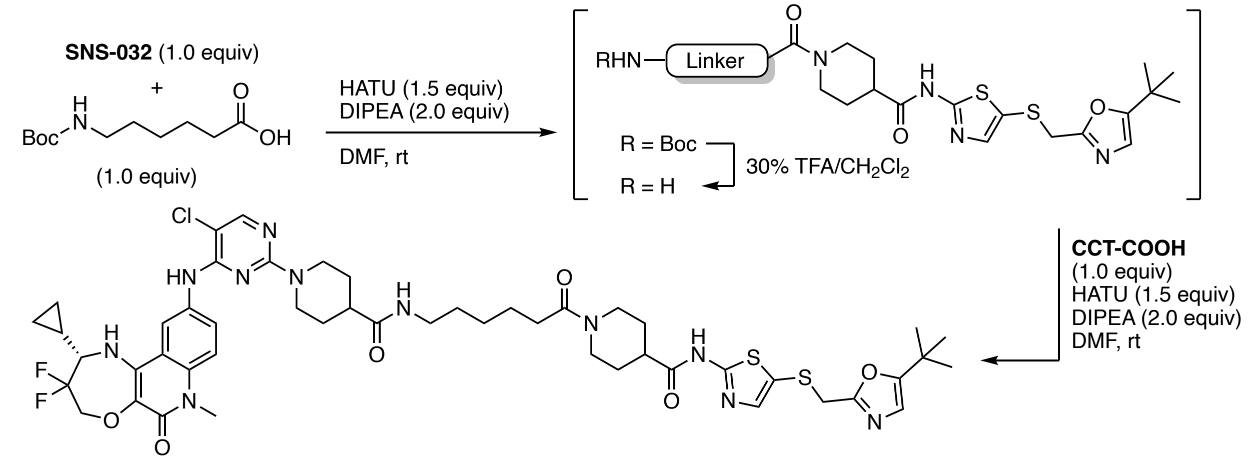


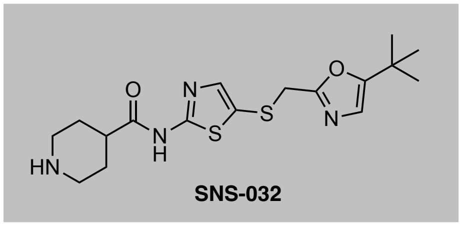


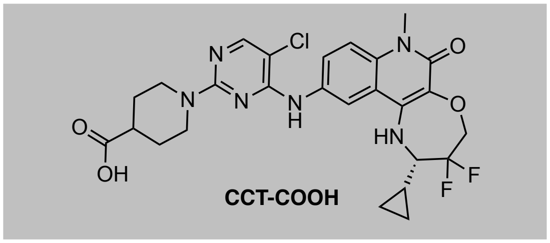


To a solution of **SNS-032** (5.0 mg, 0.013 mmol, 1.0 equiv) and 6-((*tert*-butoxycarbonyl)amino)hexanoic acid (3.0 mg, 0.013 mmol, 1.0 equiv) in DMF (0.2 M) was added HATU (7.5 mg, 0.020 mmol, 1.5 equiv) and DIPEA (4.6 μL, 0.026 mmol, 2.0 equiv). The resulting yellow solution was stirred at ambient temperature for 2 h. Upon full consumption of starting material, as judged by LC-MS and TLC analysis, the reaction mixture was diluted with water and EtOAc. After separation of the organic layer, the aqueous phase was extracted with EtOAc (3x) and the combined organic extracts were washed with brine, dried over $Na_2SO_4$, and concentrated under reduced pressure. The resulting intermediate was taken up in 30% v/v TFA in $CH_2Cl_2$ and the resulting solution was stirred at ambient temperature until LC-MS showed quantitative formation of free amine intermediate. All volatiles were removed under reduced pressure. The intermediate was used without further purification, dissolved in DMF (0.1 mL), and added to a solution of **CCT-COOH** (0.013 g, 0.023 mmol, 1.0 equiv), (7.5 mg, 0.020 mmol, 1.5 equiv), and DIPEA (4.6 μL, 0.026 mmol, 2.0 equiv) in DMF (0.2 mL). After stirring at ambient temperature for 30 min, purification by reverse-phase HPLC (15-90% MeOH in water) afforded the product as white solid (5.3 mg, 39% yield).

**$^1$H NMR** (500 MHz, DMSO-$d_6$) δ = 12.29 (s, 1H), 9.22 (s, 1H), 8.17 (d, *J* = 2.3 Hz, 1H), 8.11 (s, 1H), 7.80 – 7.62 (m, 2H), 7.45 (d, *J* = 9.1 Hz, 1H), 7.38 (s, 1H), 6.71 (s, 1H), 6.20 (s, 1H), 4.50 – 4.29 (m, 5H), 4.04 (s, 2H), 3.87 (1H, assigned by HSQC), 3.56 (s, 3H), 3.27 – 3.15 (m, 1H), 3.00 (q, *J* = 6.4 Hz, 3H), 2.85 (t, *J* = 12.6 Hz, 2H), 2.71 (ddd, *J* = 11.5, 7.5, 3.8 Hz, 1H), 2.60 – 2.53 (m, 1H), 2.39 – 2.24 (m, 3H), 1.79 (t, *J* = 15.2 Hz, 2H), 1.65 (d, *J* = 12.9 Hz, 2H), 1.47 (dd, *J* = 14.3, 7.7 Hz, 6H), 1.40 – 1.29 (m, 3H), 1.28 –.20 (m, 2H), 1.17 (s, 9H), 0.70 (td, *J* = 6.2, 4.3 Hz, 1H), 0.55 – 0.47 (m, 2H), 0.33 (q, *J* = 5.9 Hz, 1H). **LC-MS:** m/z 1036.37 $[M+1]^+$.

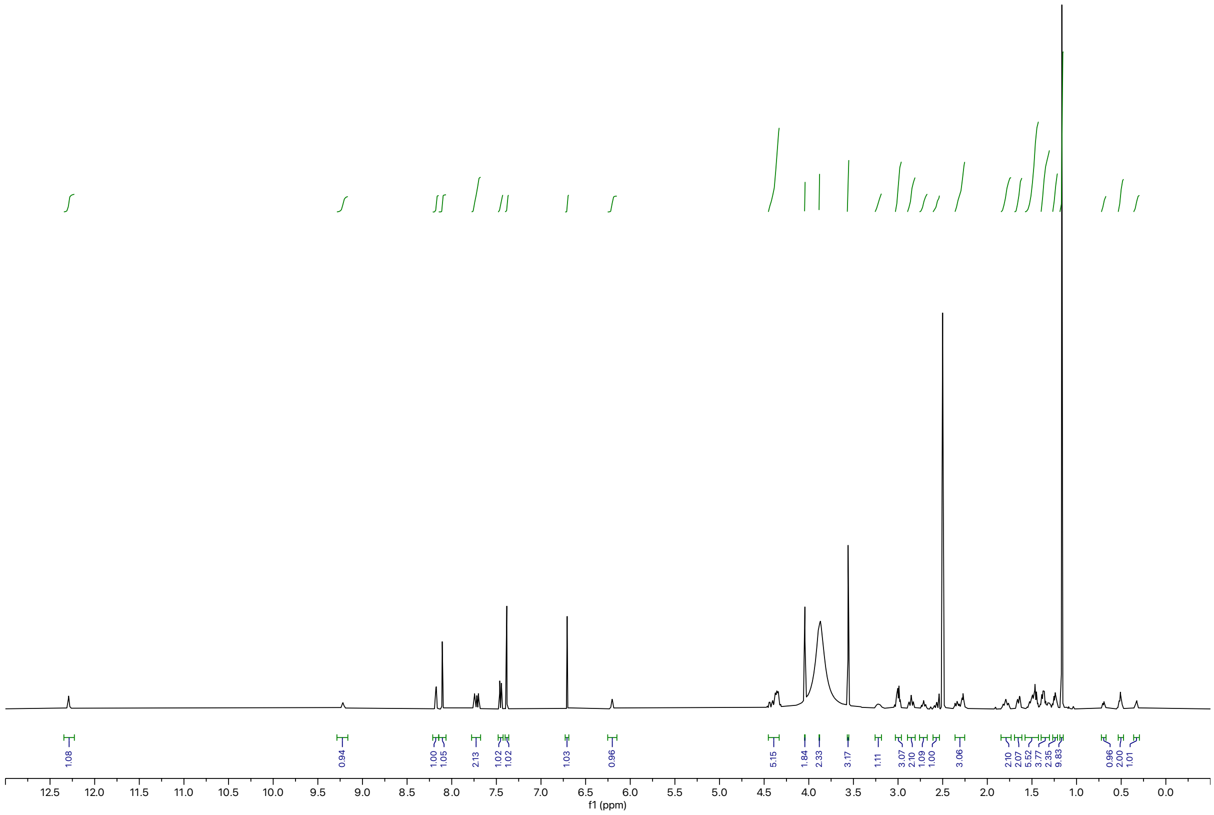

1.08
0.94
1.00
1.05
2.13
1.02
1.02
1.03
0.96
5.15
1.84
2.33
3.17
1.11
3.07
2.10
1.09
1.00
3.06
2.10
2.07
5.52
3.77
2.35
9.83
0.96
2.00
1.01
12.5 12.0 11.5 11.0 10.5 10.0 9.5 9.0 8.5 8.0 7.5 7.0 6.5 6.0 5.5 5.0 4.5 4.0 3.5 3.0 2.5 2.0 1.5 1.0 0.5 0.0
f1 (ppm)

Synthesis of **BAK-04-101**

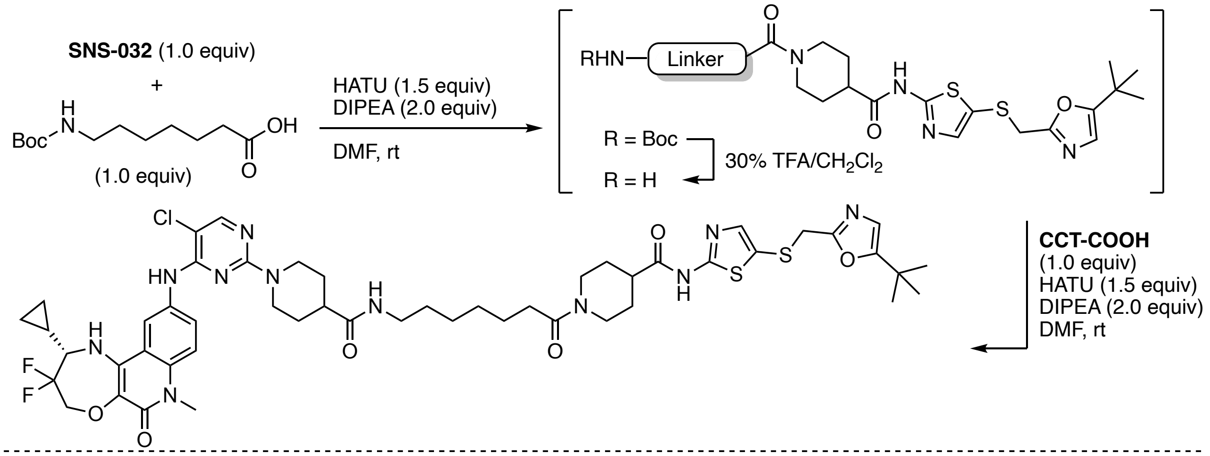


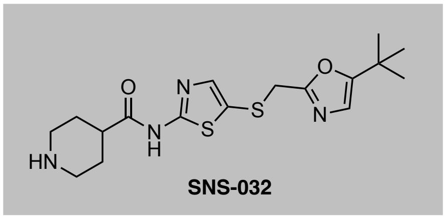


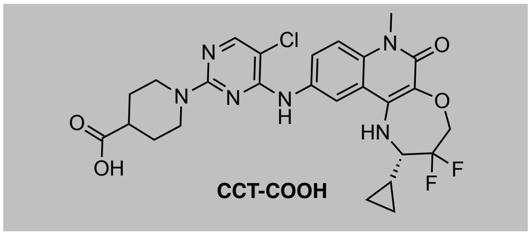


To a solution of **SNS-032** (3.8 mg, 0.010 mmol, 1.0 equiv) and 7-((*tert*-butoxycarbonyl)amino)heptanoic acid as linker (2.4 mg, 0.010 mmol, 1.0 equiv) in DMF (0.2 M) was added HATU (5.7 mg, 0.015 mmol, 1.5 equiv) and DIPEA (3.5 μL, 0.020 mmol, 2.0 equiv). The resulting yellow solution was stirred at ambient temperature for 2 h. Upon full consumption of starting material, as judged by LC-MS and TLC analysis, the reaction mixture was diluted with water and EtOAc. After separation of the organic layer, the aqueous phase was extracted with EtOAc (3x) and the combined organic extracts were washed with brine, dried over $Na_2SO_4$, and concentrated under reduced pressure. The resulting intermediate was taken up in 30% v/v TFA in $CH_2Cl_2$ and the resulting solution was stirred at ambient temperature until LC-MS showed quantitative formation of free amine intermediate. All volatiles were removed under reduced pressure. The intermediate was used without further purification, dissolved in DMF (0.2 M), and added to a solution of **CCT-COOH** (5.6 mg, 0.010 mmol, 1.0 equiv), HATU (5.7 mg, 0.015 mmol, 1.5 equiv), and DIPEA (3.5 μL, 0.020 mmol, 2.0 equiv) in DMF (0.2 mL). After stirring at ambient temperature for 30 min, purification by reverse-phase HPLC (15-90% MeOH in water) afforded the product as yellowish solid (6.6 mg, 40% yield). **$^1$H NMR** (500 MHz, DMSO-d$_6$) δ = 12.29 (s, 1H), 9.12 (s, 1H), 8.17 (d, *J* = 2.3 Hz, 1H), 8.09 (s, 1H), 7.76 – 7.68 (m, 2H), 7.45 (d, *J* = 9.2 Hz, 1H), 7.38 (s, 1H), 6.71 (s, 1H), 6.19 (s, 1H), 4.48 – 4.30 (m, 5H), 4.04 (s, 2H), 3.89 (d, *J* = 13.4 Hz, 1H), 3.56 (3H, assigned by HSQC), 3.22 (tt, *J* = 16.3, 7.7 Hz, 1H), 2.99 (q, *J* = 6.7 Hz, 3H), 2.83 (t, *J* = 12.6 Hz, 2H), 2.71 (tt, *J* = 11.3, 3.9 Hz, 1H), 2.61 – 2.52 (m, 1H), 2.40 – 2.24 (m, 3H), 1.80 (t, *J* = 13.3 Hz, 2H), 1.64 (d, *J* = 12.8 Hz, 2H), 1.55 – 1.42 (m, 6H), 1.36 (q, *J* = 7.4 Hz, 3H), 1.24 (4H, assigned by HSQC), 1.17 (s, 9H), 0.70 (ddd, *J* = 8.7, 6.3, 3.0 Hz, 1H), 0.51 (t, *J* = 6.1 Hz, 2H), 0.33 (q, *J* = 5.9 Hz, 1H). **$^{19}$F NMR** (471 MHz, DMSO-d$_6$) δ = -74.45. **LC-MS:** m/z 1050.60 $[M+1]^+$.

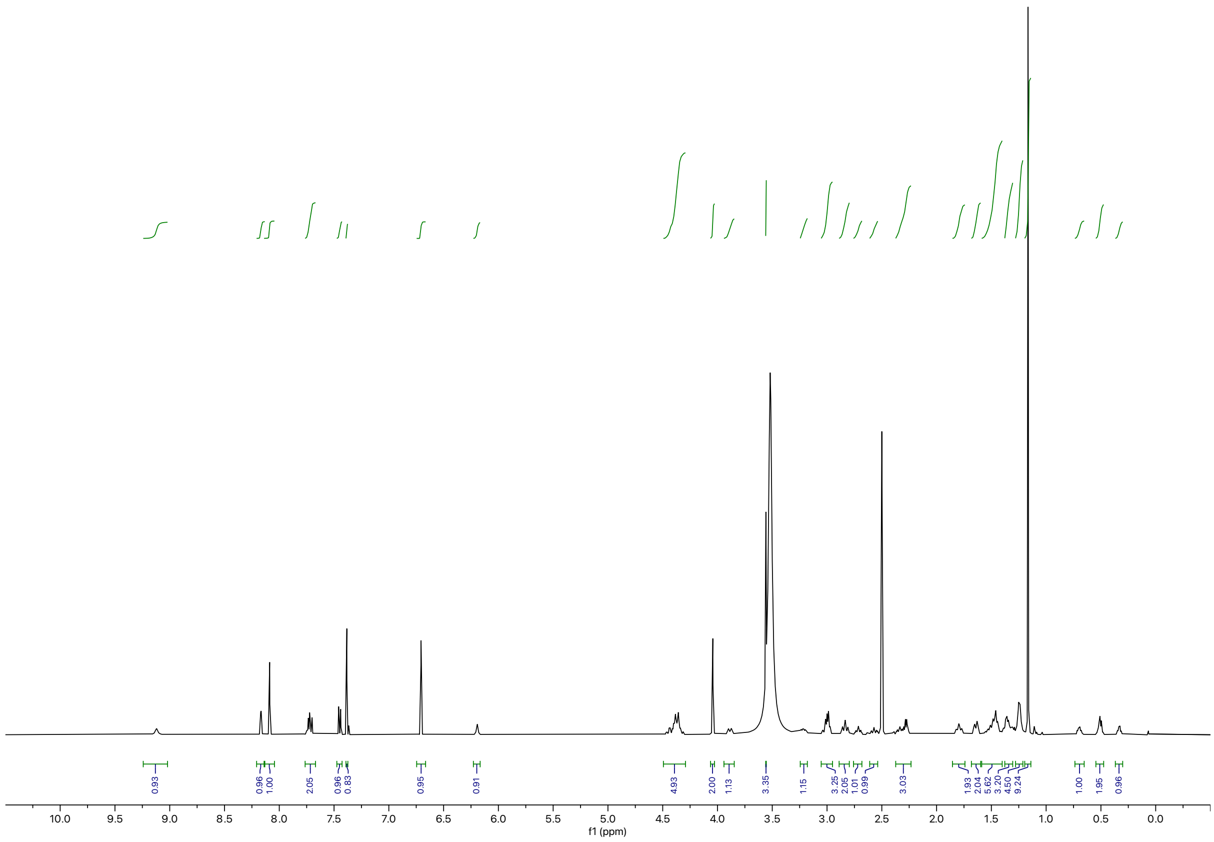

0.93
0.96
1.00
2.05
0.96
0.83
0.95
0.91
4.93
2.00
1.13
3.35
1.15
3.25
2.05
1.01
0.99
3.03
1.93
2.04
5.62
3.20
4.50
9.24
1.00
1.95
0.96
10.0 9.5 9.0 8.5 8.0 7.5 7.0 6.5 6.0 5.5 5.0 4.5 4.0 3.5 3.0 2.5 2.0 1.5 1.0 0.5 0.0
f1 (ppm)

Synthesis of **RCS-03-038**

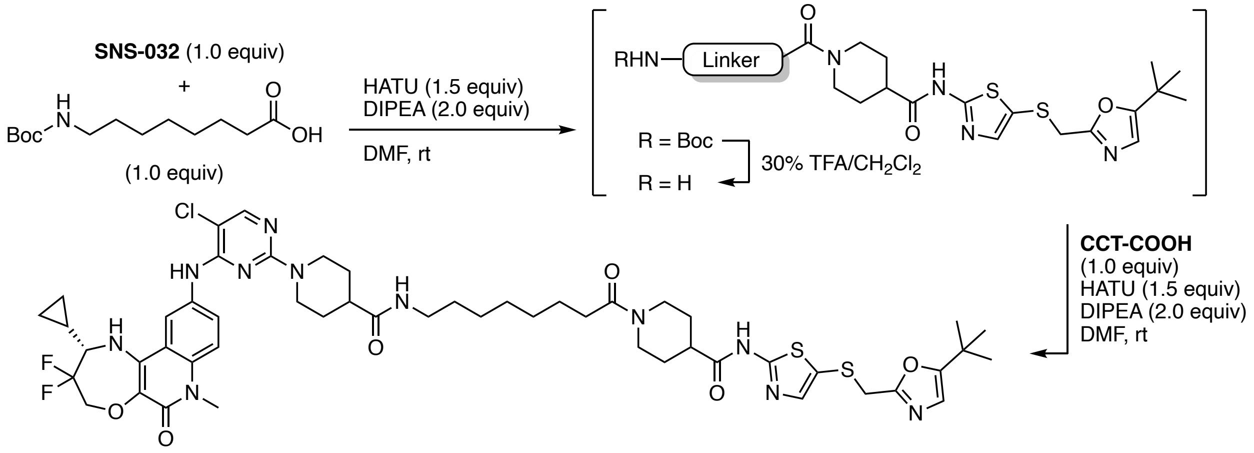


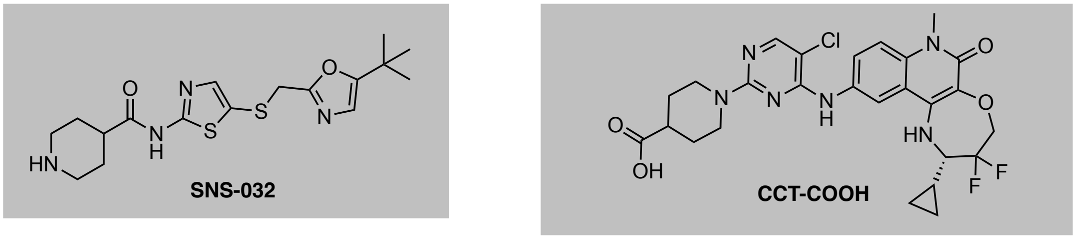


To a solution of **SNS-032** (5.0 mg, 0.013 mmol, 1.0 equiv) and 8-((*tert*-butoxycarbonyl)amino)octanoic acid (3.4 mg, 0.013 mmol, 1.0 equiv) in DMF (0.2 M) was added HATU (7.5 mg, 0.020 mmol, 1.5 equiv) and DIPEA (4.6 μL, 0.026 mmol, 2.0 equiv). The resulting yellow solution was stirred at ambient temperature for 2 h. Upon full consumption of starting material, as judged by LC-MS and TLC analysis, the reaction mixture was diluted with water and EtOAc. After separation of the organic layer, the aqueous phase was extracted with EtOAc (3x) and the combined organic extracts were washed with brine, dried over $Na_2SO_4$, and concentrated under reduced pressure. The resulting intermediate was taken up in 30% v/v TFA in $CH_2Cl_2$ and the resulting solution was stirred at ambient temperature until LC-MS showed quantitative formation of free amine intermediate. All volatiles were removed under reduced pressure. The intermediate was used without further purification, dissolved in DMF (0.1 mL), and added to a solution of **CCT-COOH** (7.4 mg, 0.013 mmol, 1.0 equiv), HATU (7.5 mg, 0.020 mmol, 1.5 equiv), and DIPEA (4.6 μL, 0.026 mmol, 2.0 equiv) in DMF (0.2 mL). After stirring at ambient temperature for 30 min, purification by reverse-phase HPLC (15-90% MeOH in water) afforded the product as off-white solid (4.5 mg, 32% yield).

**$^1$H NMR** (500 MHz, DMSO-$d_6$) δ = 12.29 (s, 1H), 9.12 (s, 1H), 8.17 (d, *J* = 2.3 Hz, 1H), 8.09 (s, 1H), 7.76 – 7.68 (m, 2H), 7.45 (d, *J* = 9.2 Hz, 1H), 7.38 (s, 1H), 6.71 (s, 1H), 6.19 (s, 1H), 4.48 – 4.30 (m, 5H), 4.04 (2H, assigned by HSQC), 3.89 (1H, assigned by HSQC), 3.56 (s, 3H), 3.22 (tt, *J* = 16.3, 7.7 Hz, 1H), 2.99 (q, *J* = 6.7 Hz, 3H), 2.83 (t, *J* = 12.6 Hz, 2H), 2.71 (tt, *J* = 11.3, 3.9 Hz, 1H), 2.61 – 2.52 (m, 1H), 2.40 – 2.24 (m, 3H), 1.80 (t, *J* = 13.3 Hz, 2H), 1.64 (d, *J* = 12.8 Hz, 2H), 1.55 – 1.42 (m, 6H), 1.36 (q, *J* = 7.4 Hz, 3H), 1.24 (4H, assigned by HSQC), 1.17 (s, 9H), 0.70 (ddd, *J* = 8.7, 6.3, 3.0 Hz, 1H), 0.51 (t, *J* = 6.1 Hz, 2H), 0.33 (q, *J* = 5.9 Hz, 1H). **LC-MS:** m/z 1064.43 $[M+1]^+$.

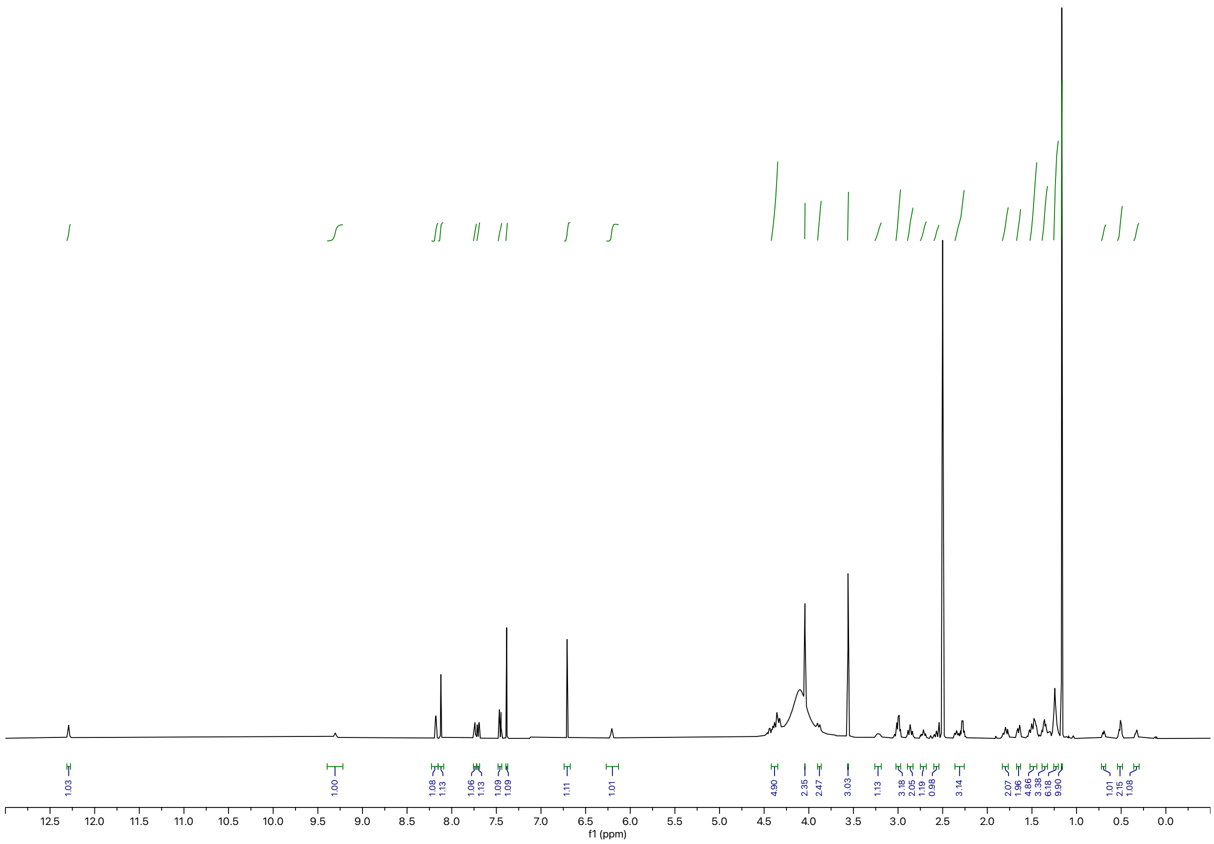

1.03
1.00
1.08
1.13
1.06
1.13
1.09
1.09
1.11
1.01
4.90
2.35
2.47
3.03
1.13
3.18
2.05
1.19
0.98
3.14
2.07
1.96
4.86
3.38
6.18
9.90
1.01
2.15
1.08
12.5 12.0 11.5 11.0 10.5 10.0 9.5 9.0 8.5 8.0 7.5 7.0 6.5 6.0 5.5 5.0 4.5 4.0 3.5 3.0 2.5 2.0 1.5 1.0 0.5 0.0
f1 (ppm)

Synthesis of **CCT-COOH**

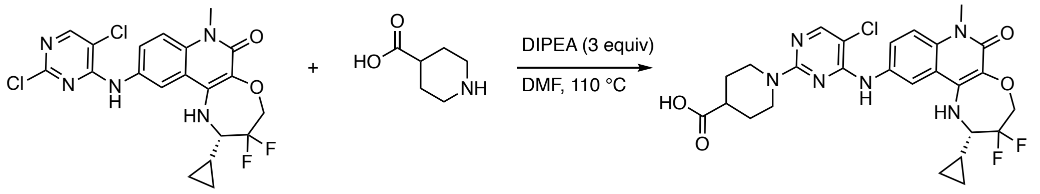


(*S*)-1-(5-chloro-4-((2-cyclopropyl-3,3-difluoro-7-methyl-6-oxo-1,2,3,4,6,7-hexahydro-[1,4]oxazepino[2,3-c]quinolin-10-yl)amino)pyrimidin-2-yl)piperidine-4-carboxylic acid (CCT-COOH): To a solution of (*S*)-2-cyclopropyl-10-((2,5-dichloropyrimidin-4-yl)amino)-3,3-difluoro-7-methyl-1,2,3,4-tetrahydro-[1,4]oxazepino[2,3-c]quinolin-6(7H)-one[4] (0.050 g, 0.11 mmol, 1.0 equiv) in NMP (0.3 M) were added piperidine-4-carboxylic acid (0.015 g, 0.12 mmol, 1.1 equiv) and DIPEA (0.056 mL, 0.33 mmol, 3.0 equiv) was added. The solution was heated to 110 °C. Upon quantitative conversion as judged by LC-MS, the reaction was diluted with DMSO and purified by preparative reverse-phase HPLC (80-25% water/MeOH) to afford the product as orange solid after lyophilization (0.043 g, 72% yield).

**$^1$H NMR** (500 MHz, DMSO-$d_6$) δ = 8.89 (br, 1H), 8.18 (d, *J* = 2.3 Hz, 1H), 8.05 (s, 1H), 7.72 (dd, *J* = 9.1, 2.3 Hz, 1H), 7.44 (d, *J* = 9.1 Hz, 1H), 6.19 (s, 1H), 4.49 – 4.34 (m, 2H), 4.34 – 4.24 (m, 2H), 3.56 (s, 3H), 3.26 – 3.17 (m, 1H), 2.94 (t, *J* = 12.2 Hz, 2H), 2.45 (dt, *J* = 11.0, 3.9 Hz, 1H), 1.84 – 1.73 (m, 2H), 1.49 – 1.40 (m, 2H), 1.32 (td, *J* = 8.2, 4.1 Hz, 1H), 0.70 (q, *J* = 5.8 Hz, 1H), 0.53 – 0.48 (m, 2H), 0.34 (q, *J* = 6.0 Hz, 1H). **$^{19}$F NMR** (471 MHz, DMSO-$d_6$) δ = -73.94. **LC-MS:** m/z 561.23 $[M+1]^+$.

Synthesis of **RCS-03-131-1**

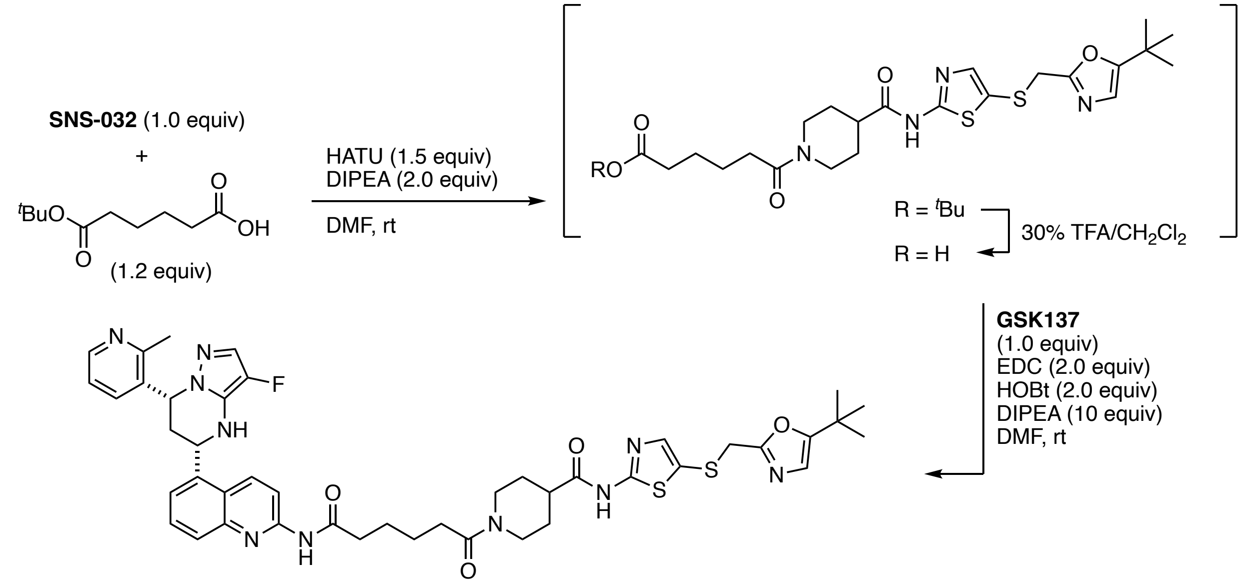


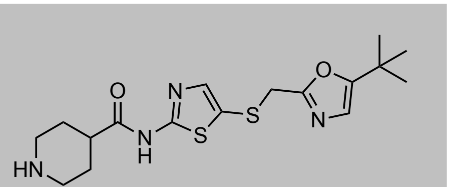

**SNS-032**

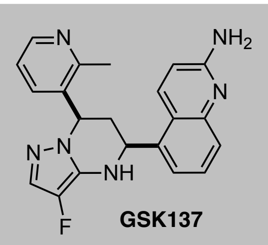

**GSK137**

To a solution of **SNS-032** (9.9 mg, 0.026 mmol, 1.0 equiv) and 6-(*tert*-butoxy)-6-oxohexanoic acid as linker (6.3 mg, 0.031 mmol, 1.2 equiv) in DMF (0.2 mL) was added HATU (0.015 g, 0.039 mmol, 1.5 equiv) and DIPEA (9.1 μL, 0.052 mmol, 2.0 equiv). The resulting yellow solution was stirred at ambient temperature for 2 h. Upon full consumption of starting material, as judged by LC-MS and TLC analysis, the reaction mixture was diluted with water and EtOAc. After separation of the organic layer, the aqueous phase was extracted with EtOAc (3x) and the combined organic extracts were washed with brine, dried over $Na_2SO_4$, and concentrated under reduced pressure. The resulting intermediate was taken up in 30% v/v TFA in $CH_2Cl_2$ and the resulting solution was stirred at 35 °C until LC-MS showed quantitative formation of acid intermediate. All volatiles were removed under reduced pressure. The intermediate was used without further purification, dissolved in DMF (0.2 M), and added to a solution of **GSK137**[5] (9.7 mg, 0.026 mmol, 1.0 equiv), EDC (8.1 mg, 0.052 mmol, 2.0 equiv), HOBt (7.0 mg, 0.052 mmol, 2.0 equiv), and DIPEA (0.045 mL, 0.26 mmol, 10 equiv) in DMF (0.2 mL). The mixture was heated to 50°C and stirred for 3 h. Purification by reverse-phase HPLC (15-90% MeOH in water) afforded the product as white solid (4.6 mg, 20% yield).

**$^1$H NMR** (500 MHz, DMSO-d$_6$) δ = 12.29 (s, 1H), 10.83 (s, 1H), 8.84 (d, *J* = 9.4 Hz, 1H), 8.70 (dd, *J* = 5.6, 1.5 Hz, 1H), 8.37 (d, *J* = 9.3 Hz, 1H), 8.19 (d, *J* = 7.9 Hz, 1H), 7.82 – 7.70 (m, 4H), 7.39 (s, 1H), 7.35 (d, *J* = 4.2 Hz, 1H), z6.85 (s, 1H), 6.70 (s, 1H), 6.00 (dd, *J* = 11.3, 4.8 Hz, 1H), 5.46 (d, *J* = 11.1 Hz, 1H), 4.38

(d, $J$ = 13.1 Hz, 1H), 4.05 (s, 2H), 3.92 (d, $J$ = 13.7 Hz, 1H), 3.03 (t, $J$ = 12.7 Hz, 1H), 2.76 – 2.65 (m, 4H), 2.60 (q, $J$ = 12.3 Hz, 2H), j2.48 – 2.46 (m, 2H), 2.36 (q, $J$ = 7.2 Hz, 2H), 2.23 (q, $J$ = 12.0 Hz, 1H), 1.81 (t, $J$ = 13.9 Hz, 2H), 1.64 (p, $J$ = 7.2 Hz, 2H), 1.54 (p, $J$ = 7.6 Hz, 3H), 1.40 (t, $J$ = 11.6 Hz, 1H), 1.17 (s, 9H). **$^{19}$F NMR** (471 MHz, DMSO-$d_6$) δ =-74.72. **LC-MS:** m/z 865.42 $[M+1]^+$.

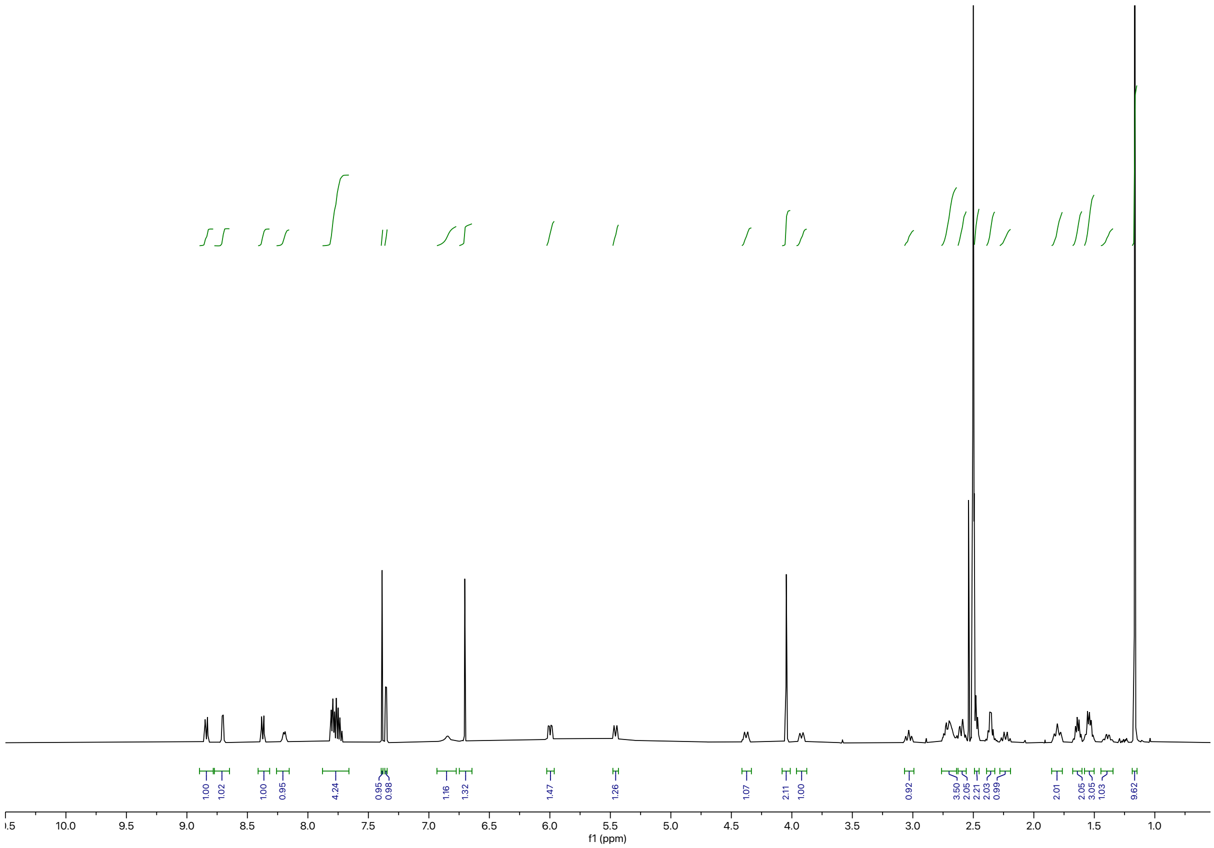

Synthesis of **RCS-03-131-2**

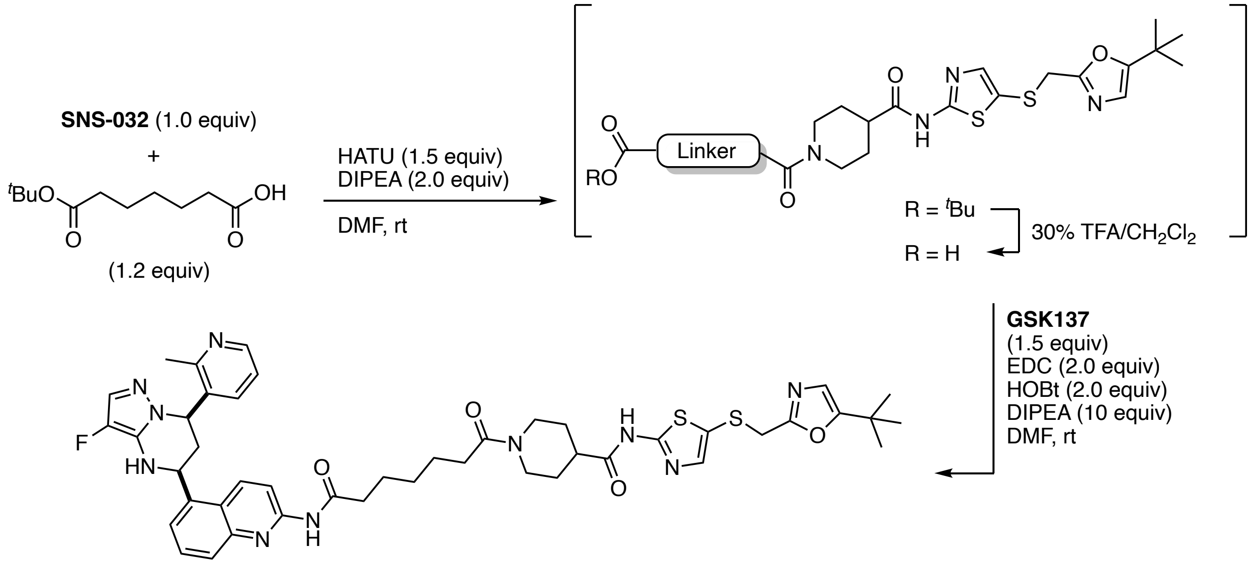


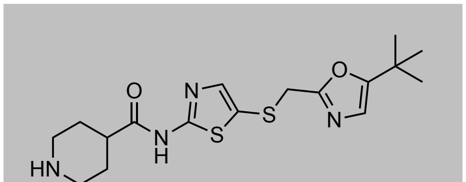

**SNS-032**

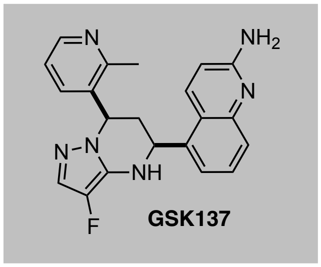

**GSK137**

To a solution of **SNS-032** (0.010 g, 0.026 mmol, 1.0 equiv) and 7-(*tert*-butoxy)-7-oxoheptanoic acid (6.8 mg, 0.032 mmol, 1.2 equiv) in DMF (0.2 mL) was added HATU (0.015 g, 0.039 mmol, 1.5 equiv) and DIPEA (9.2 μL, 0.052 mmol, 2.0 equiv). The resulting yellow solution was stirred at ambient temperature for 2 h. Upon full consumption of starting material, as judged by LC-MS and TLC analysis, the reaction mixture was diluted with water and EtOAc. After separation of the organic layer, the aqueous phase was extracted with EtOAc (3x) and the combined organic extracts were washed with brine, dried over $Na_2SO_4$, and concentrated under reduced pressure. The resulting intermediate was taken up in 30% v/v TFA in $CH_2Cl_2$ and the resulting solution was stirred at 35 °C until LC-MS showed quantitative formation of acid intermediate. All volatiles were removed under reduced pressure. The intermediate was used without further purification, dissolved in DMF (0.2 M), and added to a solution of **GSK137**[5] (0.015 mg, 0.039 mmol, 1.5 equiv), EDC (8.2 mg, 0.053 mmol, 2.0 equiv), HOBt (7.1 mg, 0.052 mmol, 2.0 equiv), and DIPEA (0.046 mL, 0.26 mmol, 10 equiv) in DMF (0.2 mL). The mixture was heated to 50°C and stirred for 3 h. Purification by reverse-phase HPLC (15-90% MeOH in water) afforded the product as off-white solid (5.8 mg, 25% yield).

**$^1$H NMR** (500 MHz, DMSO-d$_6$) δ = 12.30 (s, 1H), 10.82 (s, 1H), 8.84 (d, *J* = 9.5 Hz, 1H), 8.69 (dd, *J* = 5.6, 1.6 Hz, 1H), 8.37 (d, *J* = 9.3 Hz, 1H), 8.17 (d, *J* = 8.1 Hz, 1H), 7.85 – 7.69 (m, 4H), 7.38 (s, 1H), 7.35 (d, *J* = 4.1 Hz, 1H), 6.83 (s, 1H), 6.70 (s, 1H), 5.99 (dd, *J* = 11.2, 4.8 Hz, 1H), 5.46 (d, *J* = 11.1 Hz, 1H), 4.38

(d, *J* = 13.4 Hz, 1H), 4.04 (s, 2H), 3.90 (d, *J* = 13.6 Hz, 1H), 3.02 (t, *J* = 12.7 Hz, 1H), 2.76 – 2.64 (m, 4H), 2.64 – 2.55 (m, 2H), 2.45 (t, *J* = 7.4 Hz, 2H), 2.30 (td, *J* = 7.3, 4.6 Hz, 2H), 2.23 (q, *J* = 11.5 Hz, 1H), 1.80 (t, *J* = 13.2 Hz, 2H), 1.61 (t, *J* = 7.2 Hz, 2H), 1.50 (q, *J* = 9.7 Hz, 3H), 1.35 – 1.29 (m, 3H), 1.16 (s, 9H).
**LC-MS:** m/z 879.46 $[M+1]^+$.

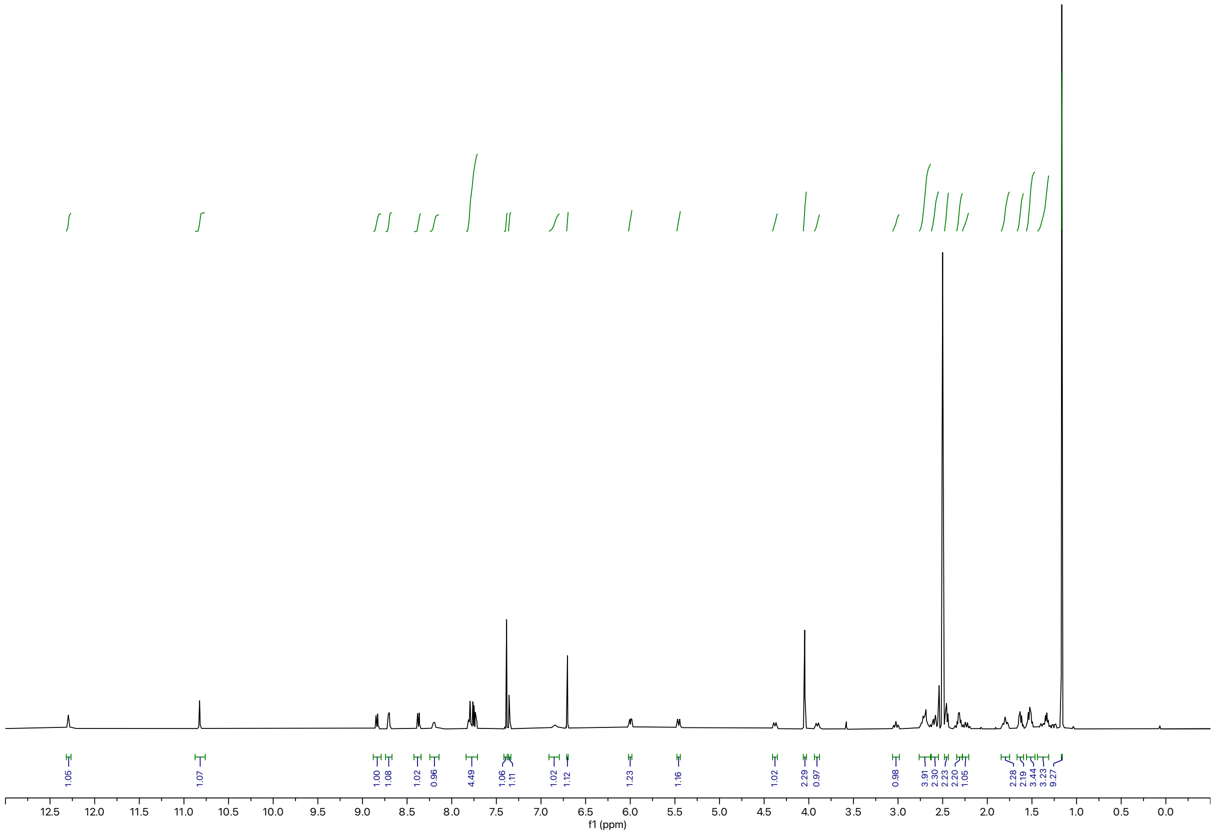

Synthesis of **RCS-03-131-3**

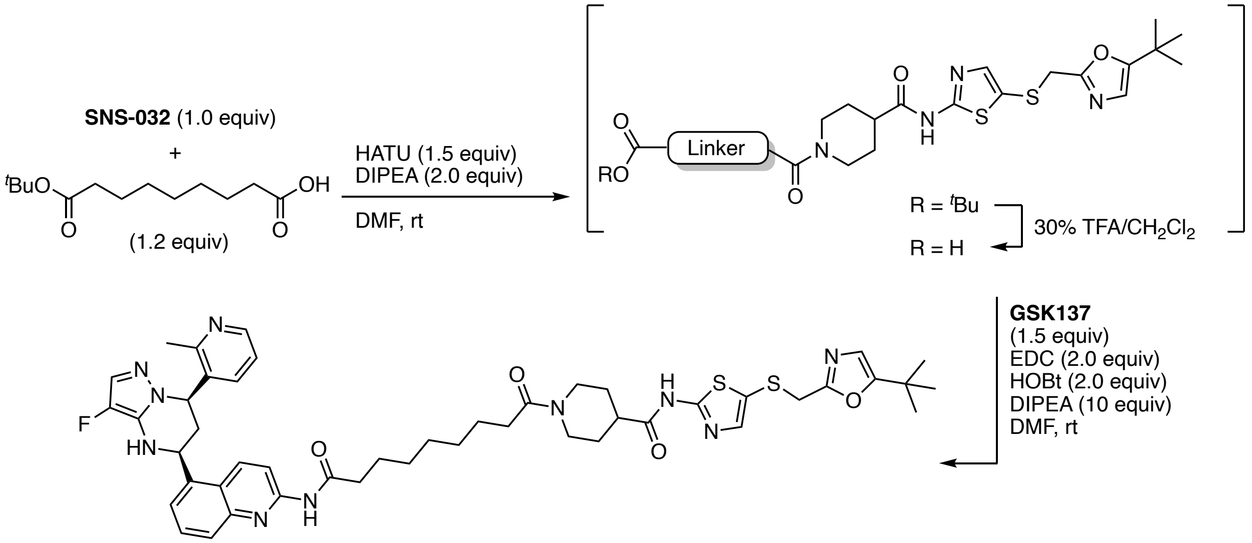


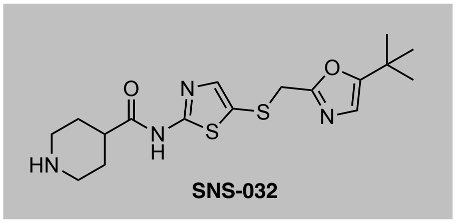


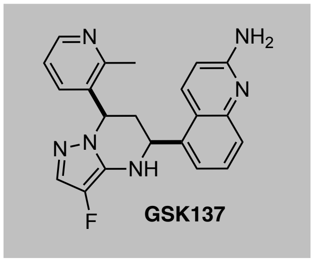


To a solution of **SNS-032** (0.010 g, 0.026 mmol, 1.0 equiv) and 9-(*tert*-butoxy)-9-oxononanoic acid (7.7 mg, 0.032 mmol, 1.2 equiv) in DMF (0.2 mL) was added HATU (0.015 g, 0.039 mmol, 1.5 equiv) and DIPEA (9.2 μL, 0.052 mmol, 2.0 equiv). The resulting yellow solution was stirred at ambient temperature for 2 h. Upon full consumption of starting material, as judged by LC-MS and TLC analysis, the reaction mixture was diluted with water and EtOAc. After separation of the organic layer, the aqueous phase was extracted with EtOAc (3x) and the combined organic extracts were washed with brine, dried over $Na_2SO_4$, and concentrated under reduced pressure. The resulting intermediate was taken up in 30% v/v TFA in $CH_2Cl_2$ and the resulting solution was stirred at 35 °C until LC-MS showed quantitative formation of acid intermediate. All volatiles were removed under reduced pressure. The intermediate was used without further purification, dissolved in DMF (0.2 M), and added to a solution of **GSK137**[5] (0.015 mg, 0.039 mmol, 1.5 equiv), EDC (8.2 mg, 0.053 mmol, 2.0 equiv), HOBt (7.1 mg, 0.052 mmol, 2.0 equiv), and DIPEA (0.046 mL, 0.26 mmol, 10 equiv) in DMF (0.2 mL). The mixture was heated to 50°C and stirred for 3 h. Purification by reverse-phase HPLC (15-90% MeOH in water) afforded the product as off-white solid (4.3 mg, 18% yield).

**$^1$H NMR** (500 MHz, DMSO-d$_6$) δ = 12.29 (s, 1H), 10.81 (s, 1H), 8.84 (d, *J* = 9.5 Hz, 1H), 8.70 (dd, *J* = 5.7, 1.5 Hz, 1H), 8.37 (d, *J* = 9.3 Hz, 1H), 8.19 (d, *J* = 8.1 Hz, 1H), 7.82 – 7.70 (m, 4H), 7.39 (s, 1H), 7.35 (d, *J* = 4.2 Hz, 1H), 6.84 (s, 1H), 6.70 (s, 1H), 6.00 (dd, *J* = 11.2, 4.8 Hz, 1H), 5.46 (d, *J* = 11.1 Hz, 1H), 4.37

(d, $J$ = 13.1 Hz, 1H), 4.04 (s, 2H), 3.89 (d, $J$ = 13.5 Hz, 1H), 3.02 (t, $J$ = 12.7 Hz, 1H), 2.78 – 2.65 (m, 4H), 2.58 (d, $J$ = 8.3 Hz, 2H), 2.45 (t, $J$ = 7.4 Hz, 2H), 2.29 (q, $J$ = 7.1 Hz, 2H), 2.26 – 2.20 (m, 1H), 1.80 (t, $J$ = 13.4 Hz, 2H), 1.61 (t, $J$ = 7.2 Hz, 2H), 1.57 – 1.44 (m, 3H), 1.38 (d, $J$ = 10.7 Hz, 1H), 1.31 (s, 6H), 1.17 (s, 9H). **LC-MS:** m/z 907.45 $[M+1]^+$.

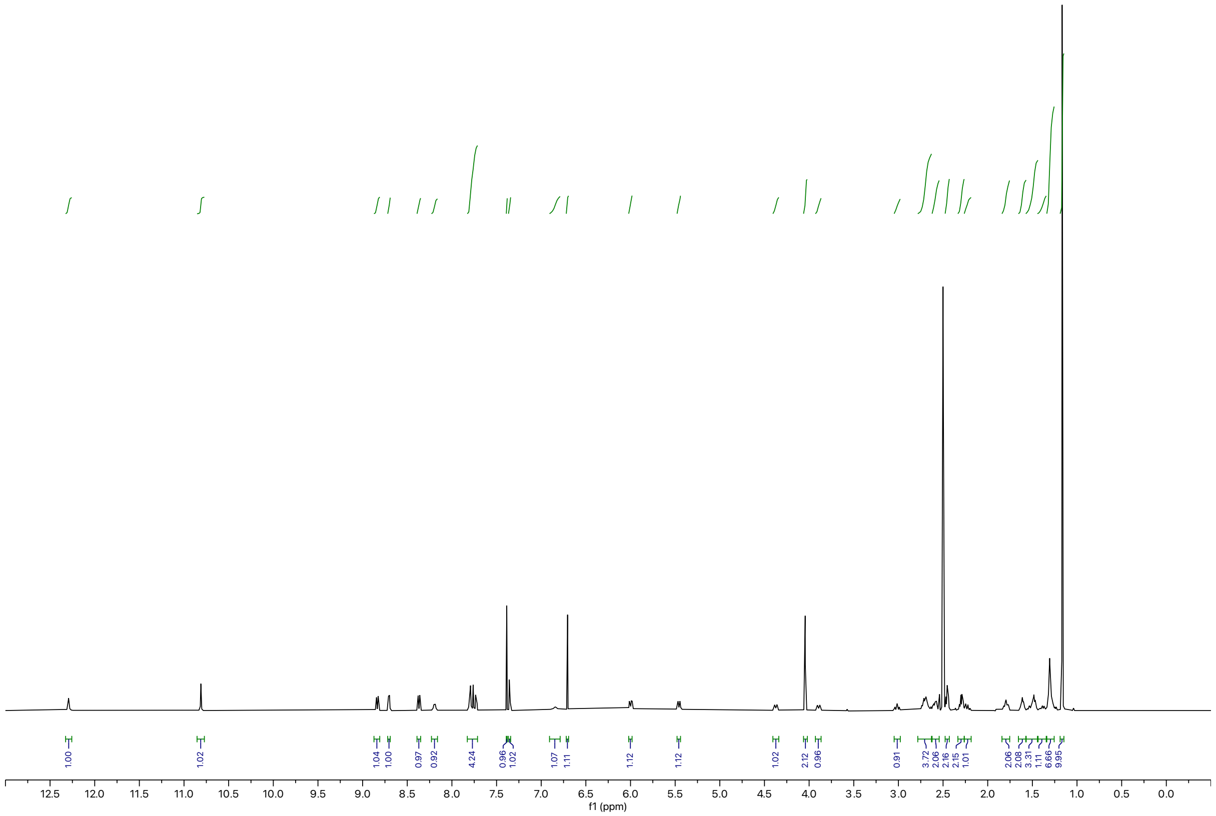

Synthesis of **RCS-03-131-4**

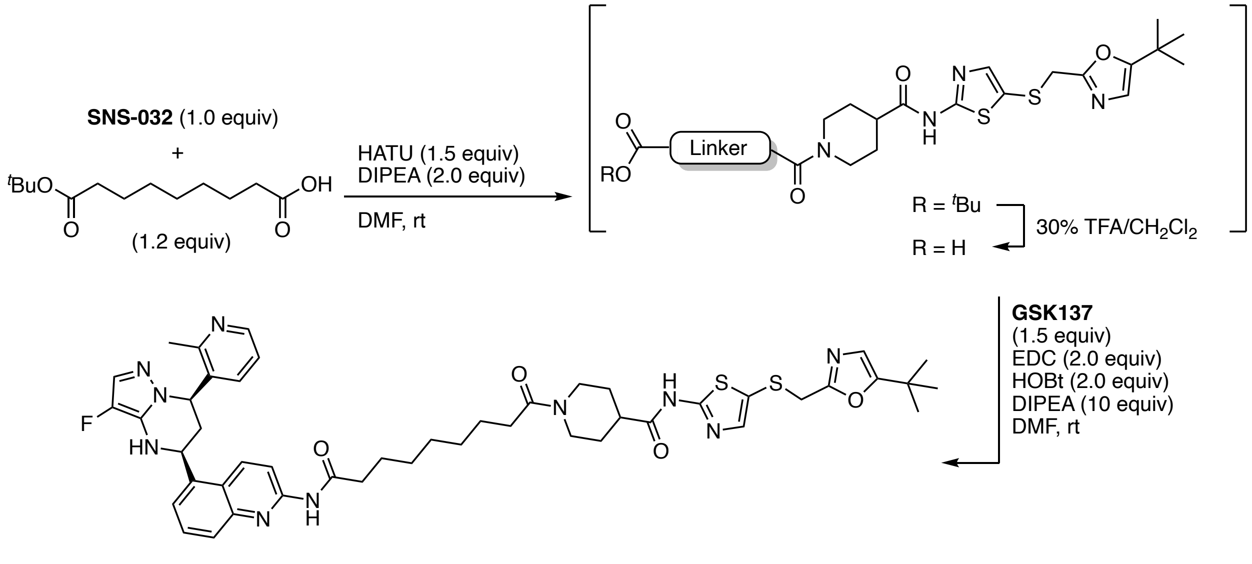


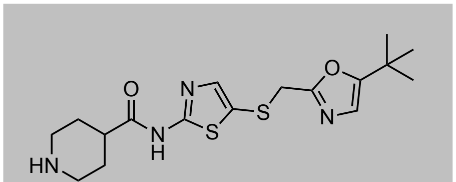
SNS-032

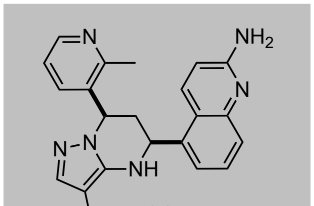
GSK137

To a solution of **SNS-032** (0.010 g, 0.026 mmol, 1.0 equiv) and 10-(*tert*-butoxy)-10-oxodecanoic acid (8.1 mg, 0.032 mmol, 1.2 equiv) in DMF (0.2 mL) was added HATU (0.015 g, 0.039 mmol, 1.5 equiv) and DIPEA (9.2 μL, 0.052 mmol, 2.0 equiv). The resulting yellow solution was stirred at ambient temperature for 2 h. Upon full consumption of starting material, as judged by LC-MS and TLC analysis, the reaction mixture was diluted with water and EtOAc. After separation of the organic layer, the aqueous phase was extracted with EtOAc (3x) and the combined organic extracts were washed with brine, dried over $Na_2SO_4$, and concentrated under reduced pressure. The resulting intermediate was taken up in 30% v/v TFA in $CH_2Cl_2$ and the resulting solution was stirred at 35 °C until LC-MS showed quantitative formation of acid intermediate. All volatiles were removed under reduced pressure. The intermediate was used without further purification, dissolved in DMF (0.2 M), and added to a solution of **GSK137**[5] (0.015 mg, 0.039 mmol, 1.5 equiv), EDC (8.2 mg, 0.053 mmol, 2.0 equiv), HOBt (7.1 mg, 0.052 mmol, 2.0 equiv), and DIPEA (0.046 mL, 0.26 mmol, 10 equiv) in DMF (0.2 mL). The mixture was heated to 50°C and stirred for 3 h. Purification by reverse-phase HPLC (15-90% MeOH in water) afforded the product as off-white solid (2.9 mg, 12% yield).

**$^1$H NMR** (500 MHz, DMSO-d$_6$) δ = 12.30 (s, 1H), 10.82 (s, 1H), 8.84 (d, *J* = 9.5 Hz, 1H), 8.69 (dd, *J* = 5.6, 1.6 Hz, 1H), 8.37 (d, *J* = 9.3 Hz, 1H), 8.17 (d, *J* = 8.1 Hz, 1H), 7.85 – 7.69 (m, 4H), 7.38 (s, 1H), 7.35 (d, *J*

= 4.1 Hz, 1H), 6.83 (s, 1H), 6.70 (s, 1H), 5.99 (dd, *J* = 11.2, 4.8 Hz, 1H), 5.46 (d, *J* = 11.1 Hz, 1H), 4.38 (d, *J* = 13.4 Hz, 1H), 4.04 (s, 2H), 3.90 (d, *J* = 13.6 Hz, 1H), 3.02 (t, *J* = 12.7 Hz, 1H), 2.76 – 2.64 (m, 4H), 2.64 – 2.55 (m, 2H), 2.45 (t, *J* = 7.4 Hz, 2H), 2.30 (td, *J* = 7.3, 4.6 Hz, 3H), 2.23 (q, *J* = 11.5 Hz, 2H), 1.80 (t, *J* = 13.2 Hz, 2H), 1.61 (t, *J* = 7.2 Hz, 3H), 1.50 (q, *J* = 9.7 Hz, 1H), 1.35 – 1.29 (m, 8H), 1.16 (s, 9H).
**LC-MS:** m/z 921.42 $[M+1]^+$.

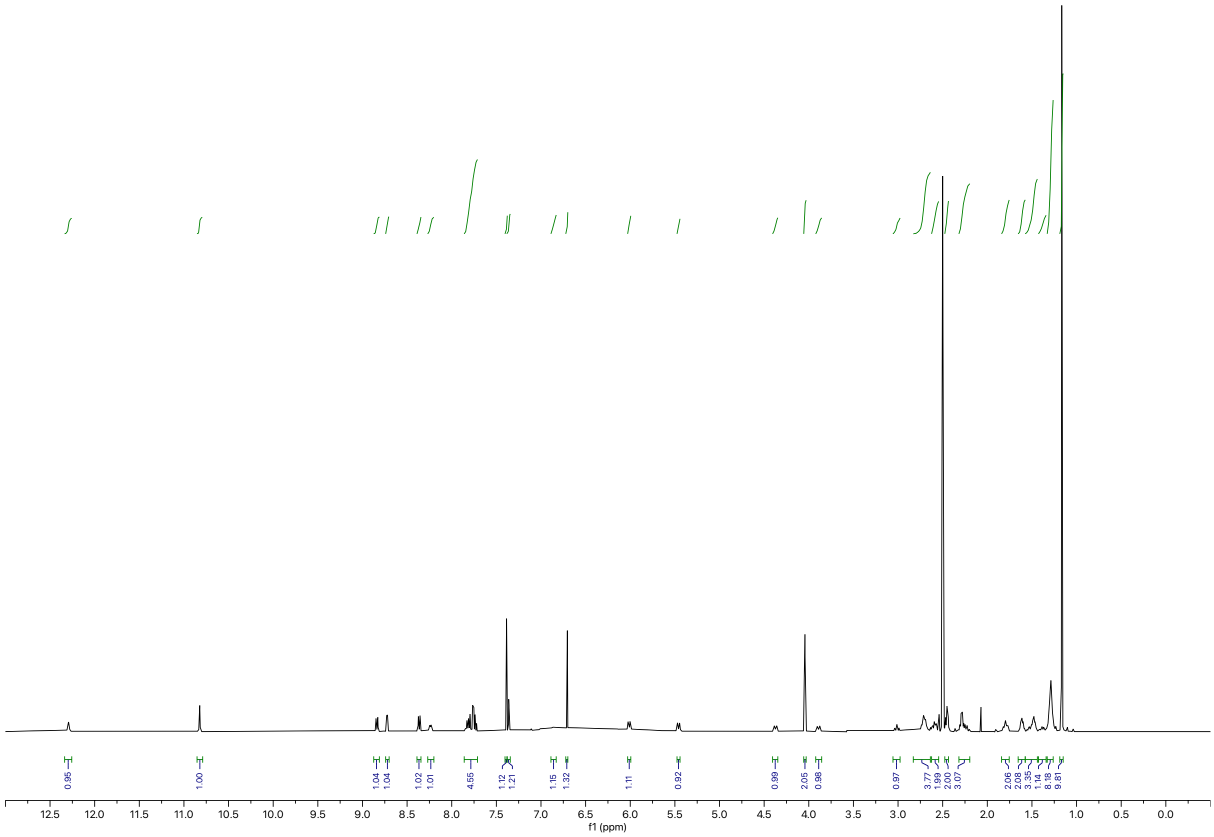

**Synthesis of AT7519 TCIPs**

General Procedure B

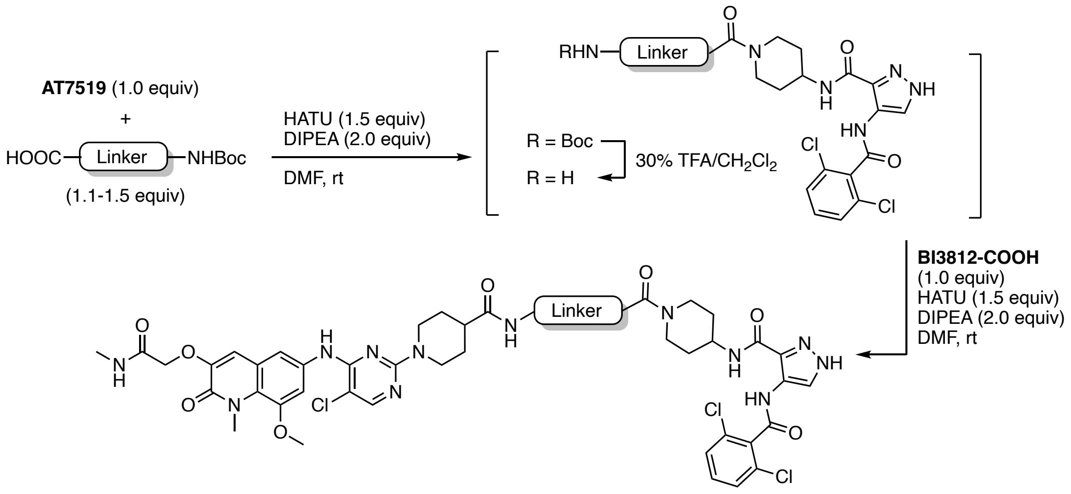


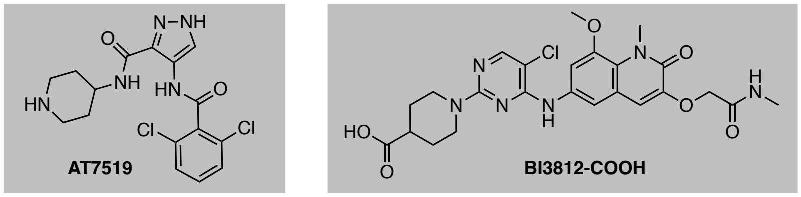


To a solution of **AT7519**[6] (1.0 equiv) and linker (1.2 equiv) in DMF (0.2 M) was added HATU (1.5 equiv) and DIPEA (2.0 equiv). The resulting yellow solution was stirred at ambient temperature for 2 h. Upon full consumption of starting material, as judged by LC-MS and/or TLC analysis, the reaction mixture was diluted with water and EtOAc. After separation of the organic layer, the aqueous phase was extracted with EtOAc (3x) and the combined organic extracts were washed with brine, dried over $Na_2SO_4$, and concentrated under reduced pressure. The resulting intermediate was taken up in 30% v/v TFA in $CH_2Cl_2$ and the resulting solution was stirred at ambient temperature until LC-MS showed quantitative formation of free amine intermediate. All volatiles were removed under reduced pressure. The intermediate was used without further purification, dissolved in DMF (0.2 M), and added to a solution of **BI3812-COOH** (1.0 equiv), HATU (1.5 equiv), and DIPEA (2.0 equiv). The reaction mixture was purified by preparative reverse-phase HPLC to afford the product upon lyophilization.

Synthesis of **BAK-04-139**

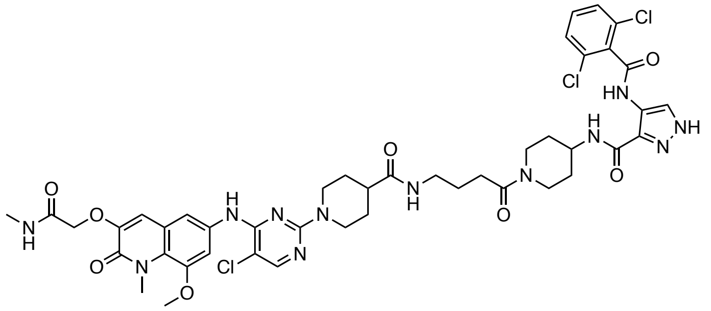

The corresponding compound was prepared following general procedure B using CDK9 inhibitor **AT7519** (3.8 mg, 0.10 mmol, 1.0 equiv) and 4-((*tert*-butoxycarbonyl)amino)butanoic acid as linker (2.4 mg, 0.012 mmol, 1.2 equiv). Purification by reverse-phase HPLC (15-90% MeOH in water) afforded the product (1.5 mg, 10% yield).

**$^1$H NMR** (500 MHz, DMSO-d$_6$) δ = 10.16 (s, 1H), 8.78 (s, 1H), 8.36 (d, *J* = 16.2 Hz, 2H), 8.05 (s, 1H), 7.95 (d, *J* = 6.4 Hz, 1H), 7.81 (t, *J* = 5.7 Hz, 1H), 7.61 – 7.50 (m, 5H), 6.99 (s, 1H), 4.59 – 4.47 (m, 4H), 4.35 (m, 1H), 3.94 (s, 1H), 3.86 (d, *J* = 9.7 Hz, 7H), 3.04 (d, *J* = 8.9 Hz, 4H), 2.88 (t, *J* = 12.3 Hz, 2H), 2.64 (d, *J* = 4.7 Hz, 3H), 2.37 (s, 1H), 2.29 (t, *J* = 7.5 Hz, 2H), 1.71 (d, *J* = 12.8 Hz, 5H), 1.60 (t, *J* = 7.2 Hz, 2H), 1.50 (t, *J* = 13.2 Hz, 3H). **LC-MS:** m/z 981.46 [M+1]$^+$.

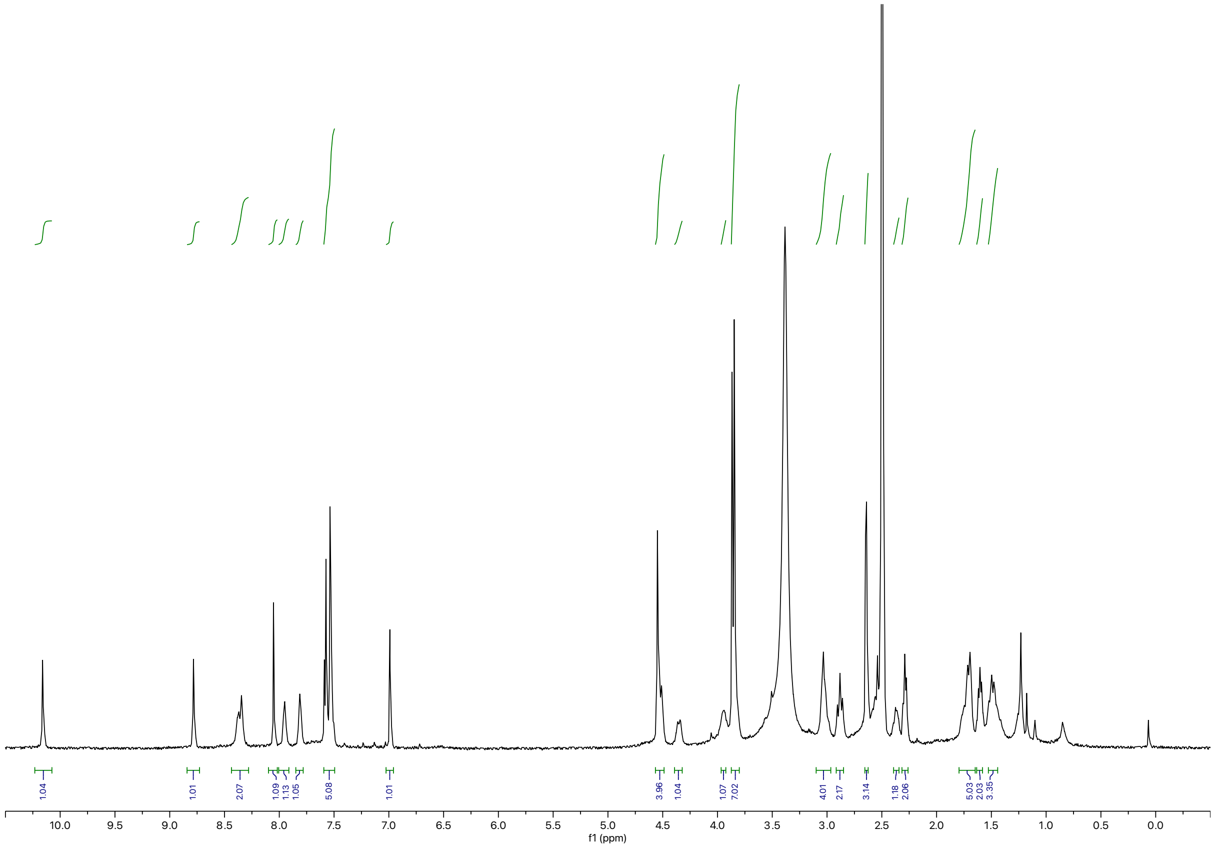

Synthesis of **BAK-04-140**

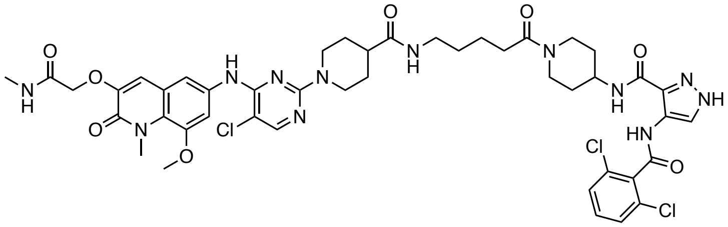

The corresponding compound was prepared following general procedure B using CDK9 inhibitor **AT7519** (3.8 mg, 0.10 mmol, 1.0 equiv) and 5-((*tert*-butoxycarbonyl)amino)pentanoic acid as linker (2.6 mg, 0.012 mmol, 1.2 equiv). Purification by reverse-phase HPLC (15-90% MeOH in water) afforded the product (6.6 mg, 50% yield).

**$^1$H NMR** (500 MHz, DMSO-d$_6$) δ =10.16 (s, 1H), 8.86 (s, 1H), 8.36 (d, *J* = 19.6 Hz, 2H), 8.07 (s, 1H), 7.94 (d, *J* = 5.0 Hz, 1H), 7.79 (t, *J* = 5.6 Hz, 1H), 7.60 – 7.55 (m, 2H), 7.53 (d, *J* = 3.7 Hz, 3H), 7.00 (s, 1H), 4.52 (d, *J* = 30.1 Hz, 4H), 4.35 (d, *J* = 13.1 Hz, 1H), 3.95 (d, *J* = 9.2 Hz, 1H), 3.86 (dd, *J* = 8.3, 2.9 Hz, 7H), 3.03 (q, *J* = 6.8 Hz, 3H), 2.89 (t, *J* = 12.7 Hz, 2H), 2.65 (d, *J* = 4.6 Hz, 3H), 2.38 (dd, *J* = 13.4, 9.4 Hz, 1H), 2.29 (t, *J* = 7.1 Hz, 2H), 1.73 (dd, *J* = 32.7, 12.2 Hz, 4H), 1.58 – 1.33 (m, 8H), 1.23 (m, 2H). **LC-MS:** m/z 995.47 [M+1]$^+$.

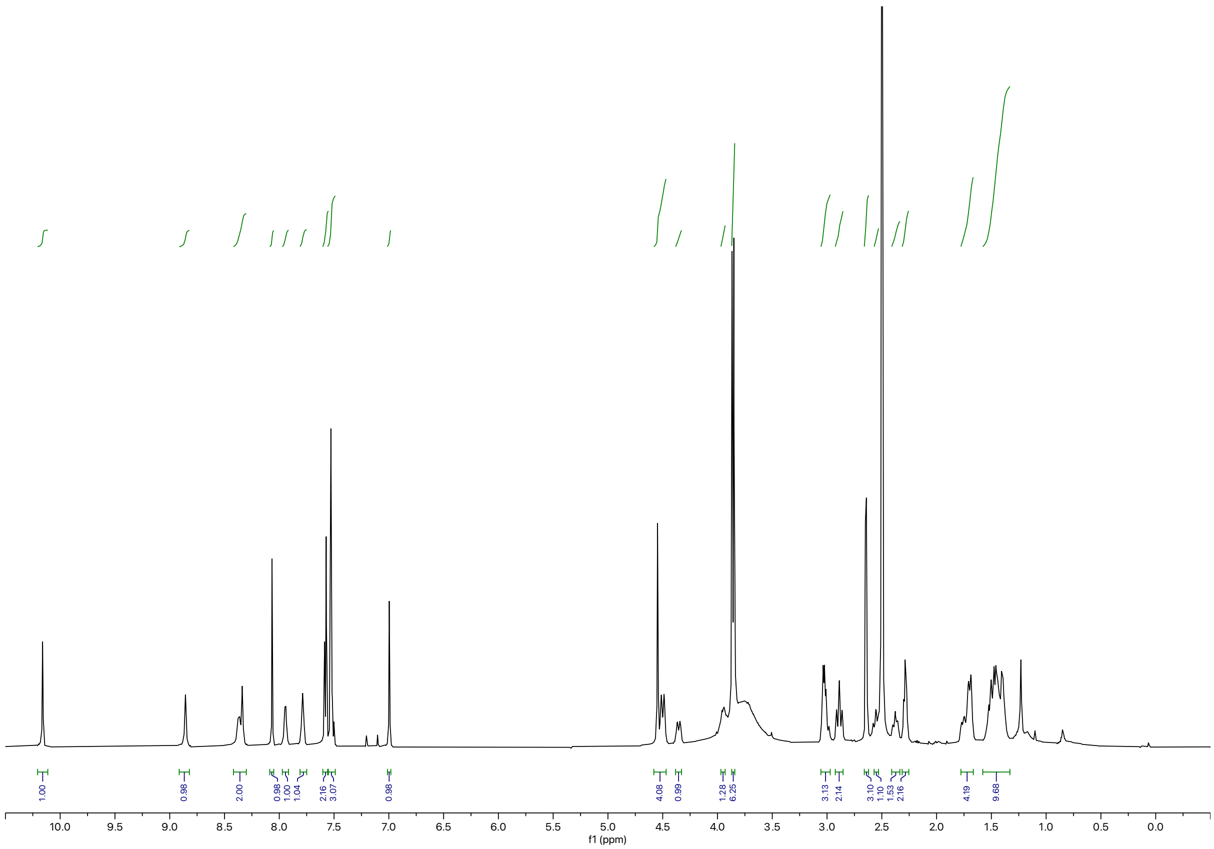

Synthesis of **BAK-04-141**

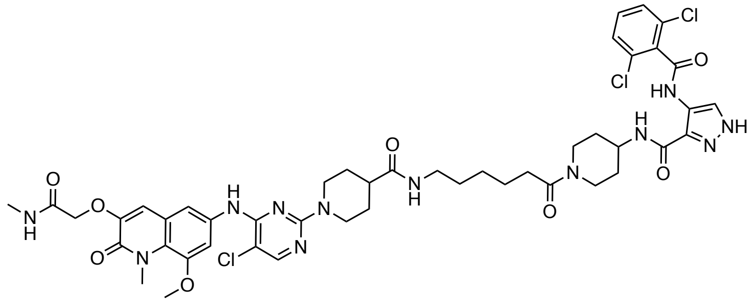

The corresponding compound was prepared following general procedure B using CDK9 inhibitor **AT7519** (3.8 mg, 0.10 mmol, 1.0 equiv) and 6-((*tert*-butoxycarbonyl)amino)hexanoic acid as linker (2.8 mg, 0.012 mmol, 1.2 equiv). Purification by reverse-phase HPLC (15-90% MeOH in water) afforded the product (1.7 mg, 10% yield).

**$^1$H NMR** (500 MHz, DMSO-$d_6$) δ = 10.16 (s, 1H), 8.80 (s, 1H), 8.36 (d, *J* = 16.8 Hz, 2H), 8.06 (s, 1H), 7.94 (d, *J* = 4.9 Hz, 1H), 7.76 (t, *J* = 5.7 Hz, 1H), 7.60 – 7.48 (m, 5H), 6.99 (s, 1H), 4.53 (d, *J* = 20.7 Hz, 4H), 4.35 (d, *J* = 12.9 Hz, 1H), 3.94 (s, 1H), 3.86 (d, *J* = 8.3 Hz, 7H), 3.01 (q, *J* = 7.3 Hz, 3H), 2.88 (t, *J* = 12.4 Hz, 2H), 2.64 (d, *J* = 4.7 Hz, 3H), 2.39 – 2.33 (m, 1H), 2.27 (t, *J* = 7.5 Hz, 2H), 1.76 – 1.67 (m, 4H), 1.46 (dt, *J* = 15.6, 8.8 Hz, 6H), 1.40 – 1.35 (m, 2H), 1.27 – 1.19 (m, 4H). **LC-MS:** m/z 1009.47 $[M+1]^+$.

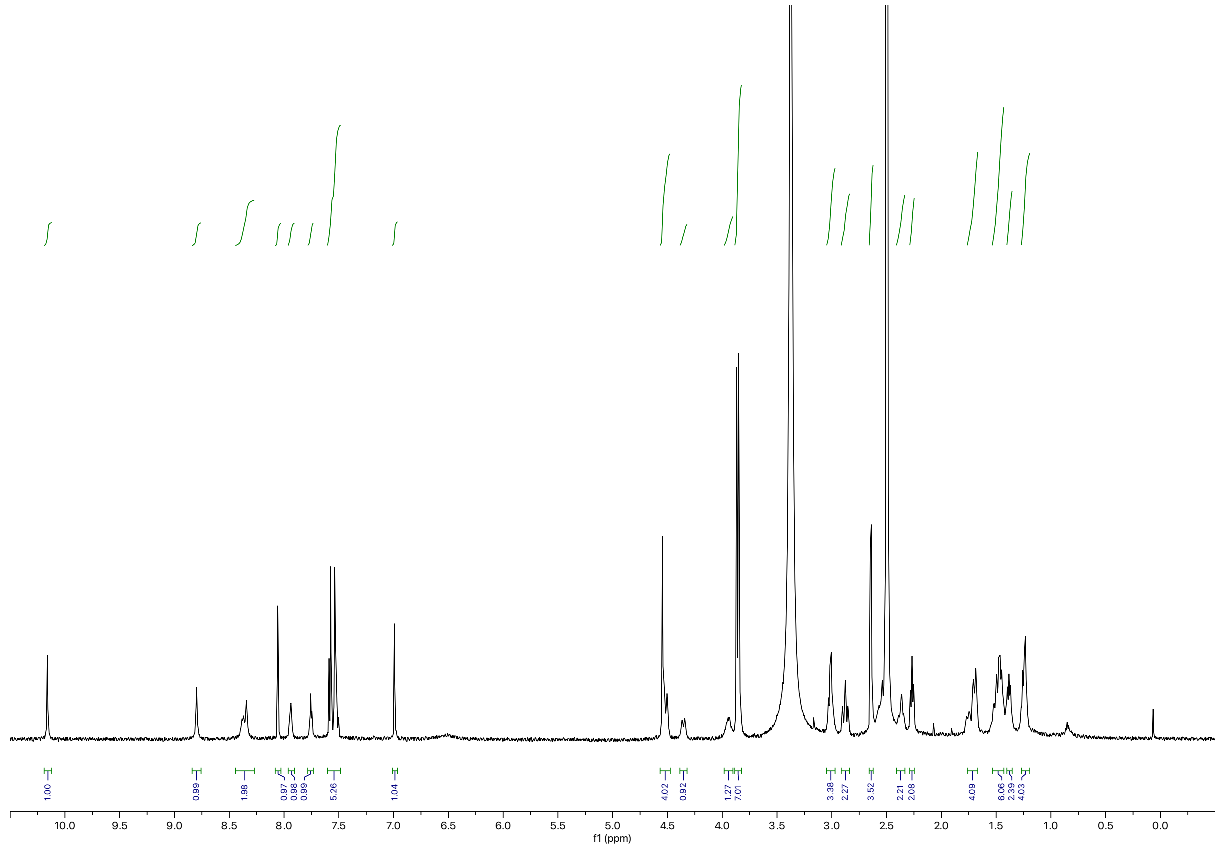

Synthesis of **BAK-04-142**

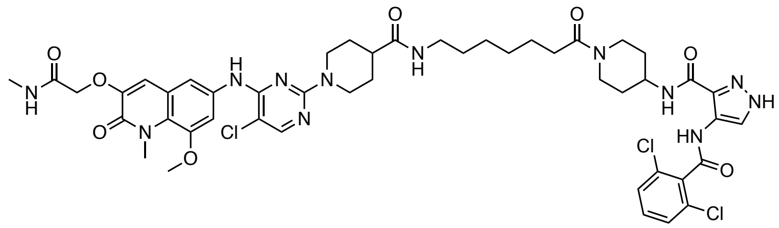

The corresponding compound was prepared following general procedure B using CDK9 inhibitor **AT7519** (3.8 mg, 0.10 mmol, 1.0 equiv) and 7-((*tert*-butoxycarbonyl)amino)heptanoic acid as linker (2.9 mg, 0.012 mmol, 1.2 equiv). Purification by reverse-phase HPLC (15-90% MeOH in water) afforded the product (2.4 mg, 20% yield).

$^1$H NMR (500 MHz, DMSO-$d_6$) δ = 10.16 (s, 1H), 8.78 (s, 1H), 8.42 – 8.28 (m, 2H), 8.05 (s, 1H), 7.96 (d, *J* = 5.0 Hz, 1H), 7.76 (t, *J* = 5.7 Hz, 1H), 7.60 – 7.50 (m, 5H), 6.99 (s, 1H), 4.53 (d, *J* = 19.4 Hz, 4H), 4.36 (d, *J* = 13.0 Hz, 1H), 3.99 – 3.90 (m, 1H), 3.86 (d, *J* = 7.8 Hz, 7H), 3.01 (q, *J* = 7.3 Hz, 3H), 2.87 (t, *J* = 12.4 Hz, 2H), 2.65 (d, *J* = 4.7 Hz, 3H), 2.36 (d, *J* = 11.1 Hz, 1H), 2.27 (t, *J* = 7.5 Hz, 2H), 1.71 (t, *J* = 17.9 Hz, 4H), 1.56 – 1.32 (m, 10H), 1.27 – 1.19 (m, 4H). **LC-MS:** m/z 1023.53 [M+1]$^+$.

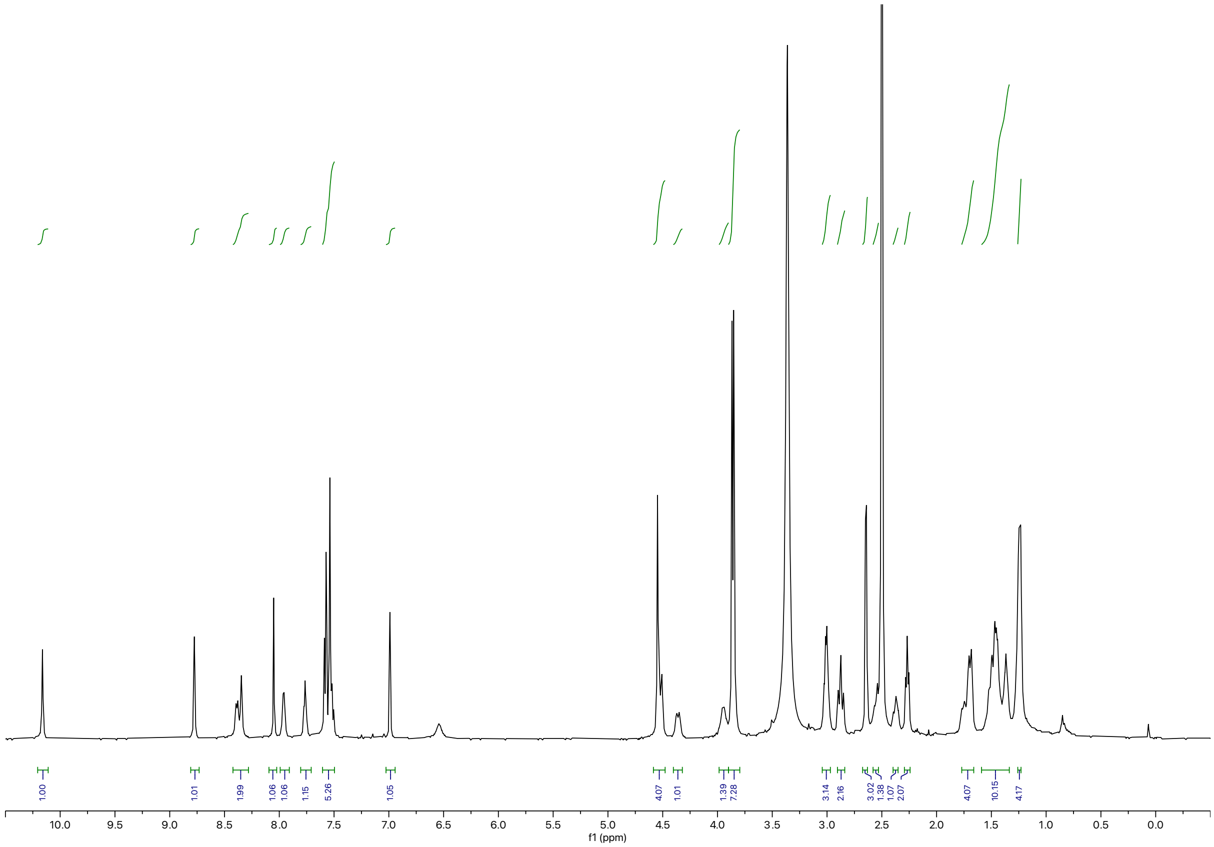

Synthesis of **BAK-04-143**

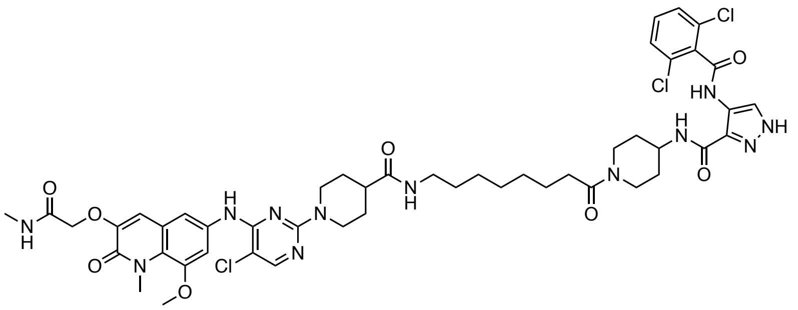

The corresponding compound was prepared following general procedure B using CDK9 inhibitor **AT7519** (3.8 mg, 0.10 mmol, 1.0 equiv) and 8-((*tert*-butoxycarbonyl)amino)octanoic acid as linker (3.1 mg, 0.012 mmol, 1.2 equiv). Purification by reverse-phase HPLC (15-90% MeOH in water) afforded the product (1.2 mg, 8% yield).

**$^1$H NMR** (500 MHz, DMSO-d$_6$) δ = 10.16 (s, 1H), 8.79 (s, 1H), 8.36 (d, *J* = 18.8 Hz, 2H), 8.05 (d, *J* = 1.9 Hz, 1H), 7.94 (d, *J* = 5.0 Hz, 1H), 7.75 (t, *J* = 5.6 Hz, 1H), 7.62 – 7.50 (m, 5H), 6.99 (d, *J* = 2.1 Hz, 1H), 4.52 (d, *J* = 19.9 Hz, 4H), 4.36 (d, *J* = 12.9 Hz, 1H), 3.94 (m, 1H), 3.86 (d, *J* = 8.1 Hz, 7H), 3.01 (q, *J* = 7.4 Hz, 3H), 2.89 (d, *J* = 12.3 Hz, 2H), 2.64 (d, *J* = 4.6 Hz, 3H), 2.39-2.32 (m, 1H), 2.26 (t, *J* = 7.5 Hz, 2H), 1.77 – 1.67 (m, 4H), 1.53 – 1.34 (m, 10H), 1.24 (d, *J* = 5.6 Hz, 6H). **LC-MS:** m/z 1037.57 [M+1]$^+$.

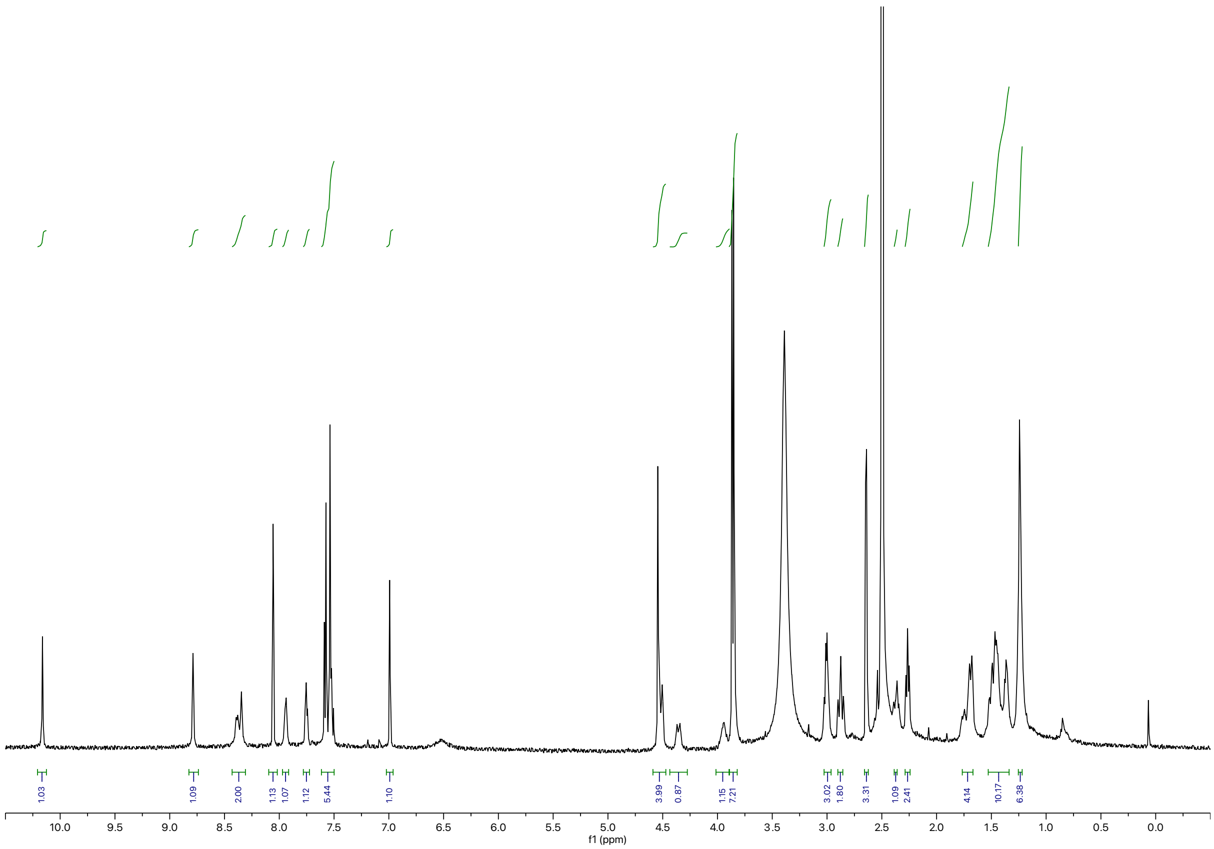

**Synthesis of AZD4573 TCIPs**

General Procedure C

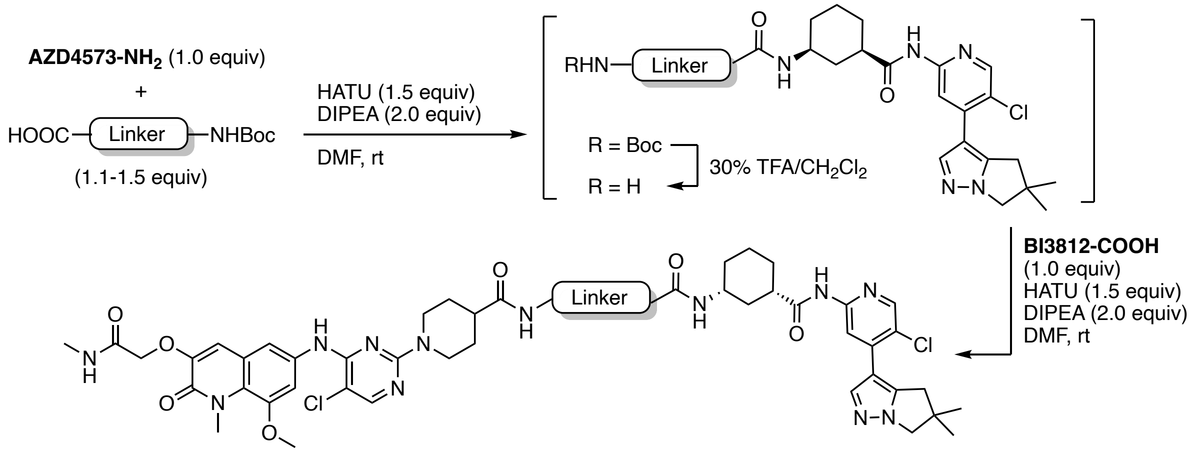


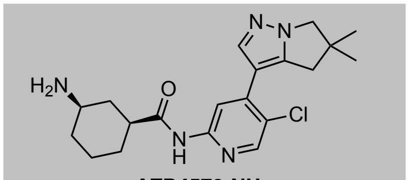

**AZD4573-NH$_2$**

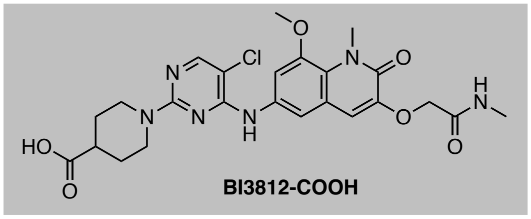

**BI3812-COOH**

To a solution of **AZD4573-NH$_2$**[7] (1.0 equiv) and linker (1.2 equiv) in DMF (0.2 M) was added HATU (1.5 equiv) and DIPEA (2.0 equiv). The resulting yellow solution was stirred at ambient temperature for 2 h. Upon full consumption of starting material, as judged by LC-MS and/or TLC analysis, the reaction mixture was diluted with water and EtOAc. After separation of the organic layer, the aqueous phase was extracted with EtOAc (3x) and the combined organic extracts were washed with brine, dried over $Na_2SO_4$, and concentrated under reduced pressure. The resulting intermediate was taken up in 30% v/v TFA in $CH_2Cl_2$ and the resulting solution was stirred at ambient temperature until LC-MS showed quantitative formation of free amine intermediate. All volatiles were removed under reduced pressure. The intermediate was used without further purification, dissolved in DMF (0.2 M), and added to a solution of **BI3812-COOH** (1.0 equiv), HATU (1.5 equiv), and DIPEA (2.0 equiv). The reaction mixture was purified by preparative reverse-phase HPLC to afford the product upon lyophilization.

Synthesis of **BAK-04-144**

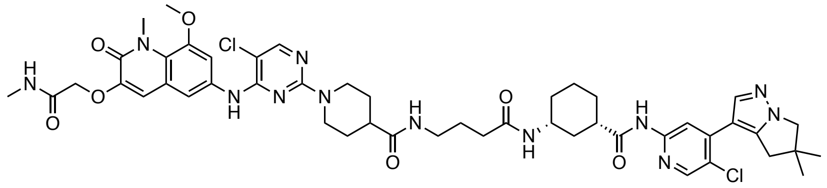

The corresponding compound was prepared following general procedure C using CDK9 inhibitor **AZD4573-NH₂** (3.9 mg, 0.01 mmol, 1.0 equiv) and 4-((*tert*-butoxycarbonyl)amino)butanoic acid as linker (2.4 mg, 0.012 mmol), 1.2 equiv. Purification by reverse-phase HPLC (15-90% MeOH in water) afforded the product (3.7 mg, 30% yield).

**$^1$H NMR** (500 MHz, DMSO-$d_6$) δ = 10.56 (s, 1H), 8.81 (s, 1H), 8.35 (s, 1H), 8.25 (s, 1H), 8.05 (s, 1H), 8.01 (s, 1H), 7.95 (d, *J* = 4.8 Hz, 1H), 7.78 (t, *J* = 5.6 Hz, 1H), 7.73 (d, *J* = 7.9 Hz, 1H), 7.53 (s, 2H), 6.99 (s, 1H), 4.58 – 4.49 (m, 4H), 3.93 (s, 2H), 3.85 (d, *J* = 8.6 Hz, 6H), 3.58 (1H, assigned by HSQC), 3.00 (q, *J* = 6.7 Hz, 2H), 2.87 (s, 4H), 2.64 (d, *J* = 4.6 Hz, 3H), 2.36 (q, *J* = 11.0 Hz, 1H), 2.02 (t, *J* = 7.3 Hz, 2H), 1.88 (d, *J* = 12.2 Hz, 1H), 1.73 (dd, *J* = 32.0, 10.2 Hz, 6H), 1.60 (q, *J* = 7.3 Hz, 2H), 1.48 (d, *J* = 12.0 Hz, 2H), 1.25 (m, 8H, assigned by HSQC), 1.08 (d, *J* = 11.9 Hz, 1H). **LC-MS:** m/z 987.58 $[M+1]^+$.

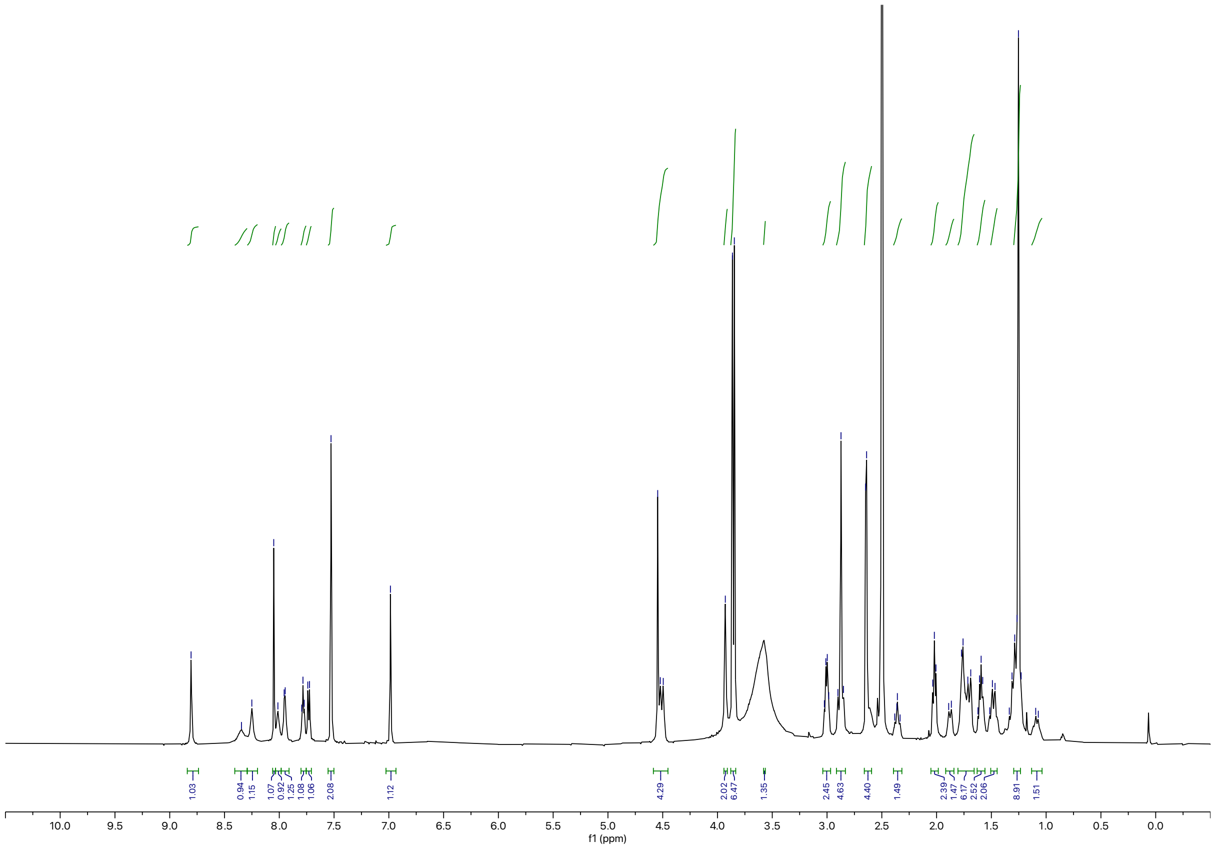

Synthesis of **BAK-04-145**

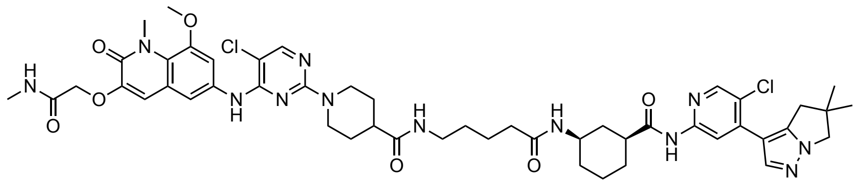

The corresponding compound was prepared following general procedure C using CDK9 inhibitor **AZD4573-$NH_2$** (3.9 mg, 0.01 mmol, 1.0 equiv)) and 5-((*tert*-butoxycarbonyl)amino)pentanoic acid as linker (2.6 mg, 0.012 mmol, 1.2 equiv). Purification by reverse-phase HPLC (15-90% MeOH in water) afforded the product (4.5 mg, 30% yield).

**$^1$H NMR** (500 MHz, DMSO-$d_6$) δ = 10.49 (s, 1H), 8.74 (s, 1H), 8.26 (s, 1H), 8.17 (s, 1H), 7.98 (s, 1H), 7.92 (s, 1H), 7.88 (d, *J* = 4.9 Hz, 1H), 7.70 (t, *J* = 5.7 Hz, 1H), 7.63 (d, *J* = 7.9 Hz, 1H), 7.46 (q, *J* = 2.4 Hz, 2H), 6.92 (s, 1H), 4.50 – 4.41 (m, 4H), 3.86 (s, 2H), 3.79 (d, *J* = 8.1 Hz, 6H), 3.50 (1H, assigned by HSQC), 2.94 (q, *J* = 6.6 Hz, 2H), 2.81 (m, 4H), 2.58 (d, *J* = 4.7 Hz, 3H), 2.54 (m, 1H), 2.34 – 2.25 (m, 1H), 1.95 (t, *J* = 7.3 Hz, 2H), 1.80 (d, *J* = 12.3 Hz, 1H), 1.66 (m, 6H), 1.41 (dq, *J* = 22.7, 8.5 Hz, 4H), 1.32 – 1.15 (10 H, assigned by HSQC), 1.01 (d, *J* = 12.0 Hz, 1H). **LC-MS:** m/z 999.66 [M+1]$^+$.

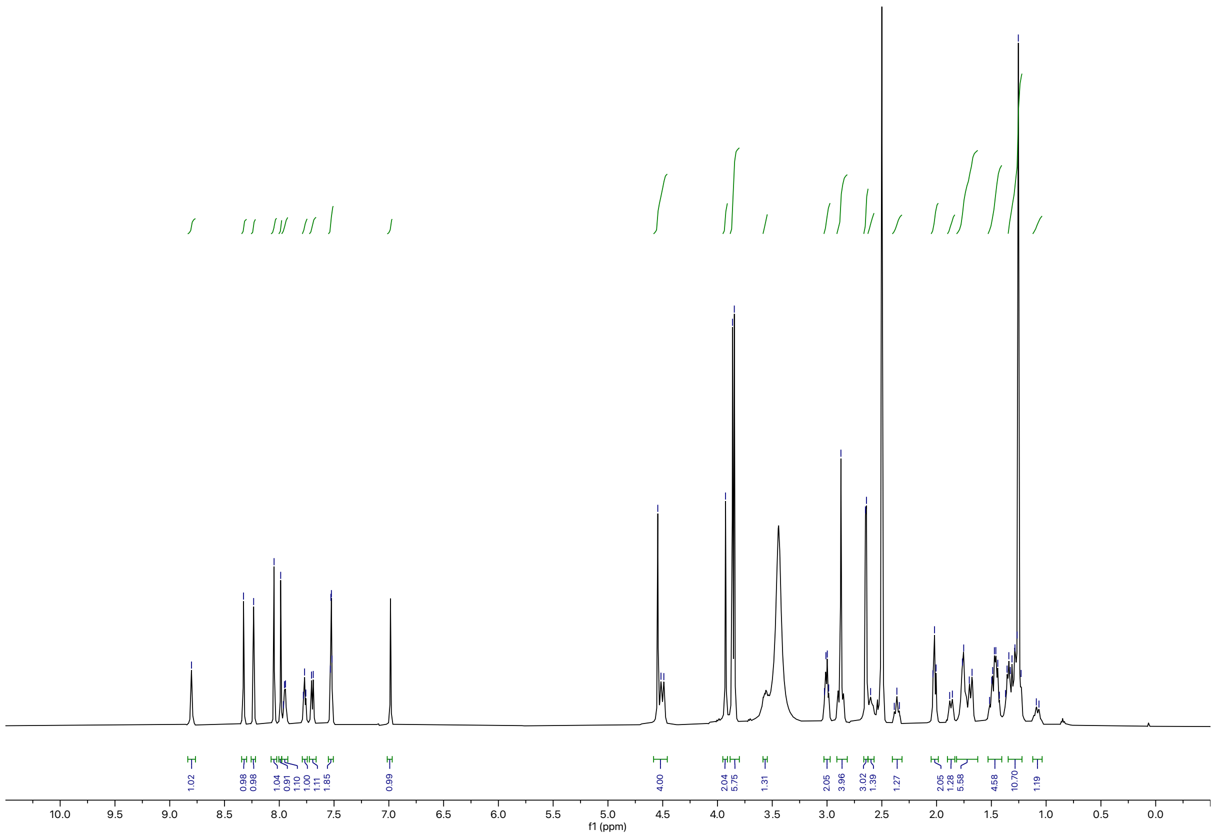

Synthesis of **BAK-04-146**

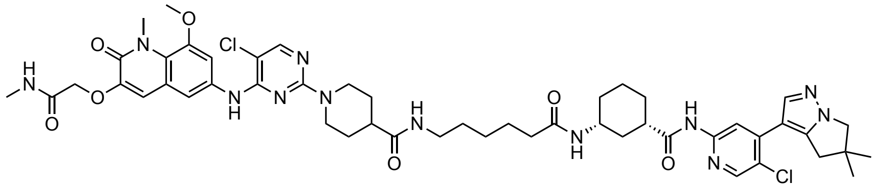

The corresponding compound was prepared following general procedure C using CDK9 inhibitor **AZD4573-NH2** (3.1 mg, 8.0 μmol, 1.0 equiv) and 6-((*tert*-butoxycarbonyl)amino)hexanoic acid as linker (2.2 mg, 9.6 μmol, 1.2 equiv). Purification by reverse-phase HPLC (15-90% MeOH in water) afforded the product (2.8 mg, 30% yield).

**$^1$H NMR** (500 MHz, DMSO-d6) δ = 10.49 (s, 1H), 8.73 (s, 1H), 8.26 (s, 1H), 8.17 (s, 1H), 7.98 (d, *J* = 1.7 Hz, 1H), 7.92 (s, 1H), 7.88 (d, *J* = 4.6 Hz, 1H), 7.68 (t, *J* = 5.5 Hz, 1H), 7.62 (d, *J* = 7.8 Hz, 1H), 7.46 (d, *J* = 1.8 Hz, 2H), 6.92 (s, 1H), 4.46 (d, *J* = 15.1 Hz, 4H), 3.86 (s, 2H), 3.79 (d, *J* = 7.8 Hz, 6H), 3.50 (d, *J* = 4.3 Hz, 1H), 2.93 (q, *J* = 6.5 Hz, 2H), 2.80 (d, *J* = 3.3 Hz, 4H), 2.58 (d, *J* = 4.6 Hz, 3H), 2.53 (s, 1H), 2.28 (d, *J* = 11.9 Hz, 1H), 1.94 (t, *J* = 7.3 Hz, 2H), 1.80 (d, *J* = 12.3 Hz, 1H), 1.65 (dd, *J* = 37.0, 8.6 Hz, 6H), 1.46 – 1.35 (m, 4H), 1.33 – 1.26 (m, 2H), 1.18 (d, *J* = 3.3 Hz, 10H), 1.01 (d, *J* = 11.7 Hz, 1H). **LC-MS:** m/z 1013.70 [M+1]$^+$.

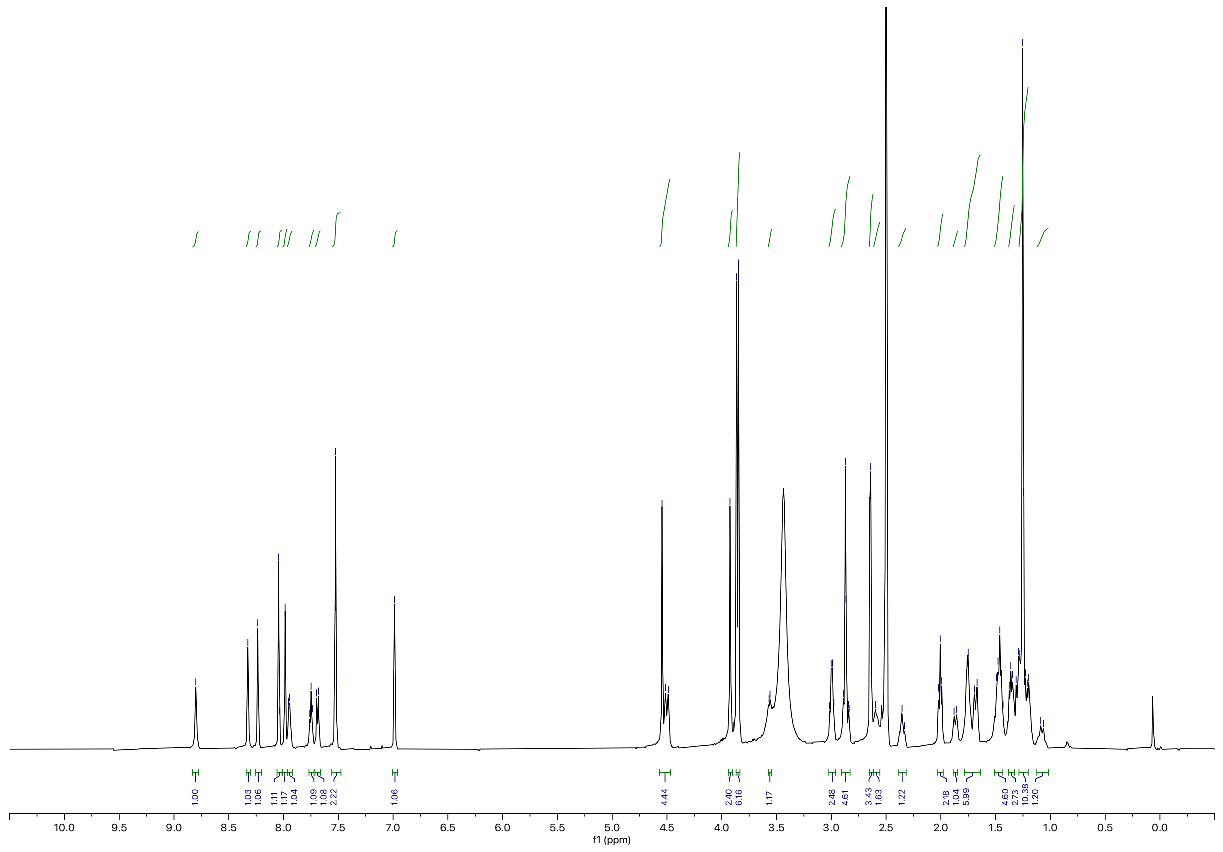

Synthesis of **BAK-04-147**

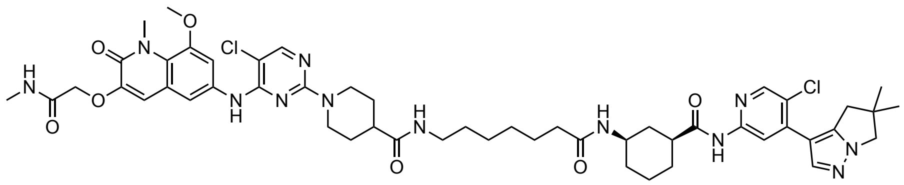


The corresponding compound was prepared following general procedure C using CDK9 inhibitor **AZD4573-NH₂** (3.9 mg, 0.01 mmol, 1.0 equiv) and 7-((*tert*-butoxycarbonyl)amino)heptanoic acid as linker (2.9 mg, 0.012 mmol, 1.2 equiv). Purification by reverse-phase HPLC (15-90% MeOH in water) afforded the product (4.7 mg, 30% yield).

**$^1$H NMR** (500 MHz, DMSO-$d_6$) δ = 10.56 (s, 1H), 8.91 (s, 1H), 8.33 (s, 1H), 8.24 (s, 1H), 8.07 (s, 1H), 7.99 (s, 1H), 7.95 (d, *J* = 4.7 Hz, 1H), 7.75 (t, *J* = 5.6 Hz, 1H), 7.69 (d, *J* = 7.9 Hz, 1H), 7.52 (s, 2H), 7.00 (s, 1H), 4.55 (s, 2H), 4.49 (d, *J* = 13.0 Hz, 2H), 3.93 (s, 2H), 3.86 (d, *J* = 7.2 Hz, 6H), 3.57 (dt, *J* = 7.8, 3.8 Hz, 1H), 2.99 (q, *J* = 6.6 Hz, 2H), 2.88 (d, *J* = 6.8 Hz, 4H), 2.64 (d, *J* = 4.7 Hz, 3H), 2.60 (s, 1H), 2.41 – 2.32 (m, 1H), 2.01 (t, *J* = 7.4 Hz, 2H), 1.87 (d, *J* = 12.7 Hz, 1H), 1.81 – 1.61 (m, 6H), 1.54 – 1.42 (m, 4H), 1.40 – 1.19 (m, 14H), 1.10 (dd, *J* = 21.0, 9.6 Hz, 1H). **LC-MS:** m/z 1027.68 [M+1]$^+$.

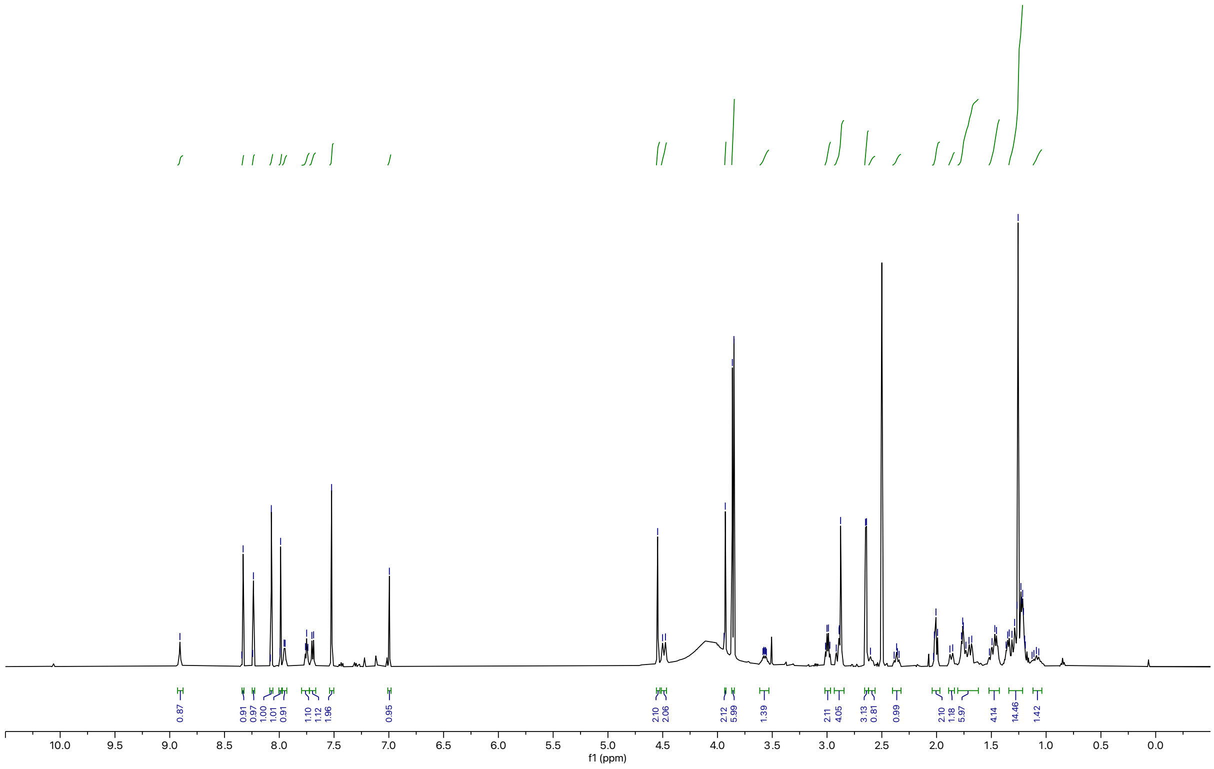

Synthesis of **BAK-04-148**

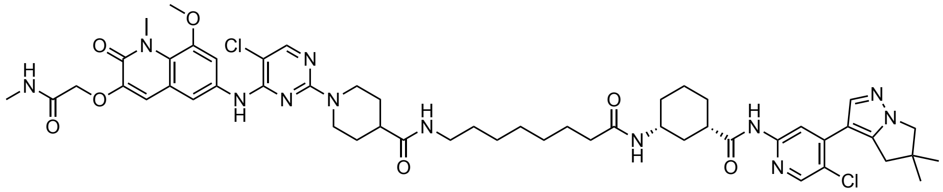

The corresponding compound was prepared following general procedure C using CDK9 inhibitor **AZD4573-NH2** (3.9 mg, 0.010 mmol, 1.2 equiv) and 7-((*tert*-butoxycarbonyl)amino)heptanoic acid as linker (3.1 mg, 0.012 mmol, 1.2 equiv). Purification by reverse-phase HPLC (15-90% MeOH in water) afforded the product (4.2 mg, 30% yield).

**$^1$H NMR** (500 MHz, DMSO-d$_6$) δ = 10.56 (s, 1H), 8.79 (s, 1H), 8.33 (d, *J* = 1.7 Hz, 1H), 8.23 (d, *J* = 1.6 Hz, 1H), 8.05 (d, *J* = 1.4 Hz, 1H), 7.98 (d, *J* = 1.5 Hz, 1H), 7.95 (d, *J* = 4.2 Hz, 1H), 7.74 (t, *J* = 5.5 Hz, 1H), 7.69 (d, *J* = 7.8 Hz, 1H), 7.53 (d, *J* = 1.6 Hz, 2H), 6.99 (d, *J* = 1.4 Hz, 1H), 4.52 (m, 4H), 3.93 (s, 2H), 3.86 (dd, *J* = 7.3, 1.8 Hz, 6H), 3.49 (1H, assigned by HSQC), 2.99 (q, *J* = 6.5 Hz, 2H), 2.87 (d, *J* = 4.2 Hz, 4H), 2.64 (dd, *J* = 6.1, 4.5 Hz, 4H), 2.61 (s, 3H), 2.35 (m, 1H), 2.00 (t, *J* = 7.3 Hz, 1H), 1.86 (d, *J* = 12.1 Hz, 1H), 1.80 – 1.62 (m, 6H), 1.53 – 1.41 (m, 4H), 1.34 (d, *J* = 6.9 Hz, 2H), 1.32 – 1.18 (m, 14H), 1.08 (d, *J* = 11.4 Hz, 1H). **LC-MS:** m/z 1041.51 [M+1]$^+$.

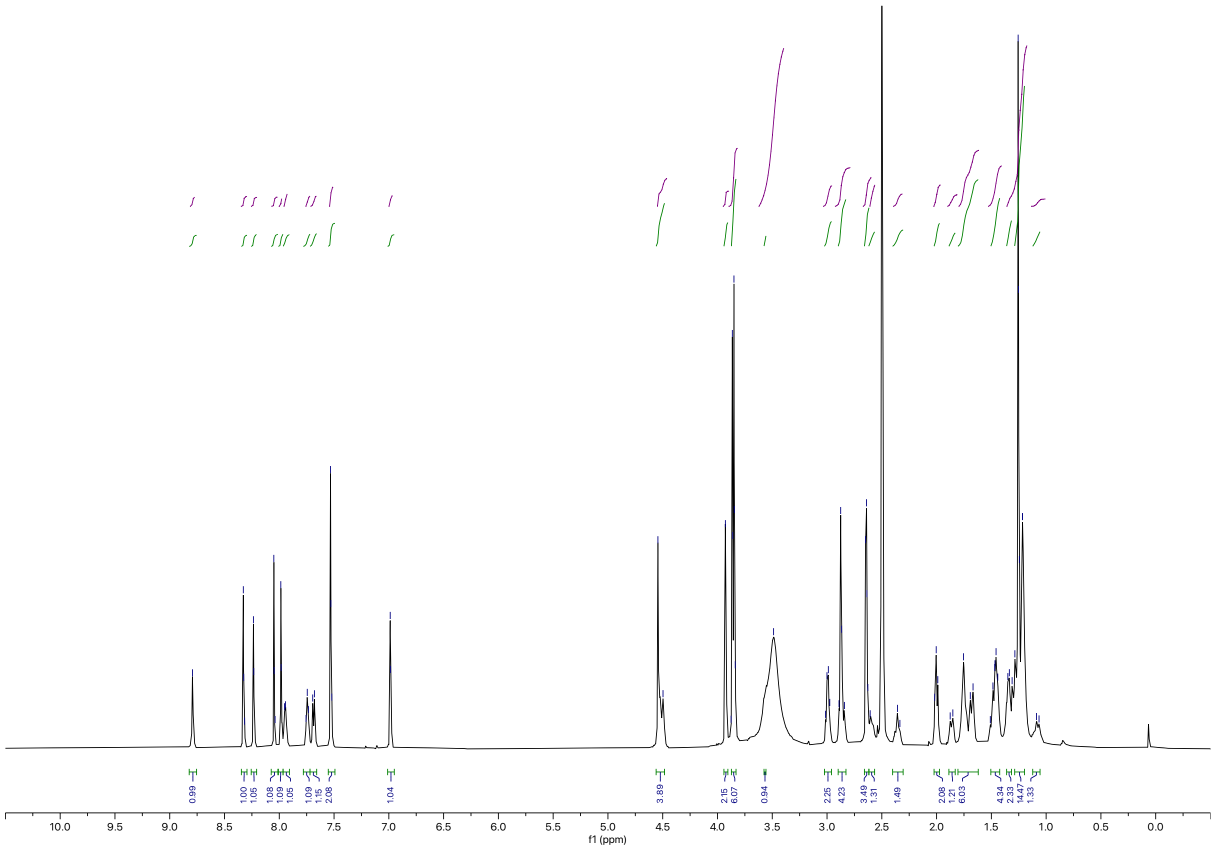

**Synthesis of NVP2 TCIPs**

General Procedure D

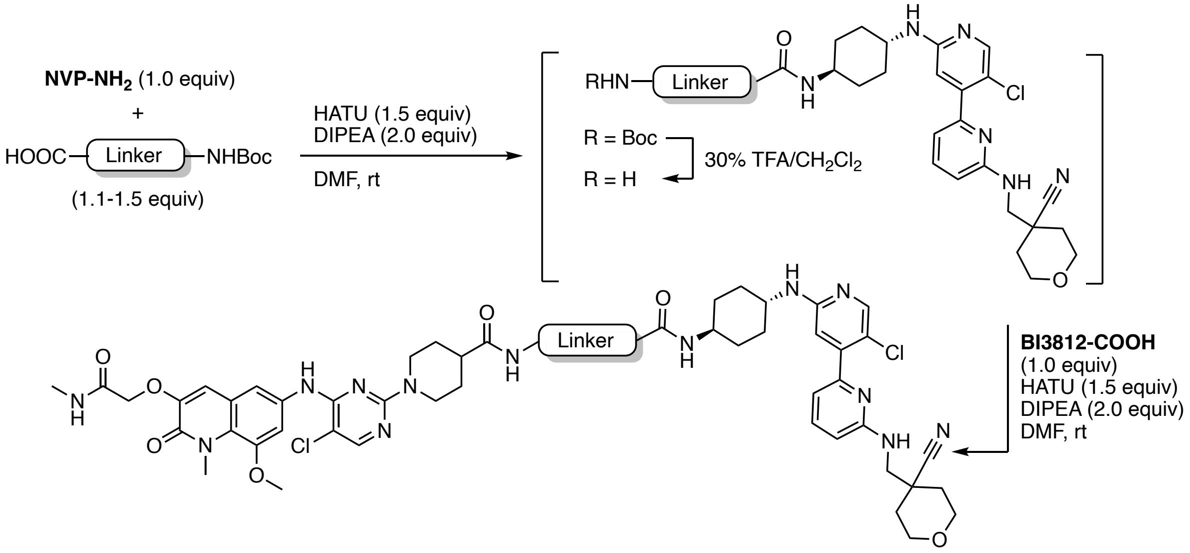


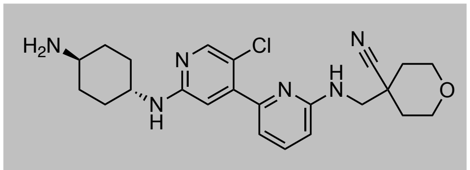

**NVP-NH$_2$**

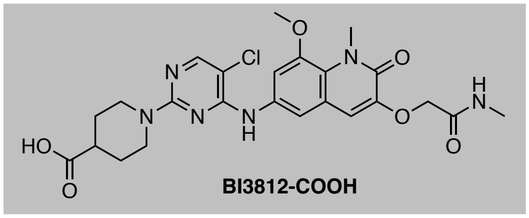

**BI3812-COOH**

To a solution of **NVP-NH$_2$**[8] (1.0 equiv) and linker (1.2 equiv) in DMF (0.2 M) was added HATU (1.5 equiv) and DIPEA (2.0 equiv). The resulting yellow solution was stirred at ambient temperature for 2 h. Upon full consumption of starting material, as judged by LC-MS and/or TLC analysis, the reaction mixture was diluted with water and EtOAc. After separation of the organic layer, the aqueous phase was extracted with EtOAc (3x) and the combined organic extracts were washed with brine, dried over $Na_2SO_4$, and concentrated under reduced pressure. The resulting intermediate was taken up in 30% v/v TFA in $CH_2Cl_2$ and the resulting solution was stirred at ambient temperature until LC-MS showed quantitative formation of free amine intermediate. All volatiles were removed under reduced pressure. The intermediate was used without further purification, dissolved in DMF (0.2 M), and added to a solution of **BI3812-COOH** (1.0 equiv), HATU (1.5 equiv), and DIPEA (2.0 equiv). The reaction mixture was purified by reverse-phase HPLC to afford the product upon lyophilization.

Synthesis of **RCS-03-097**

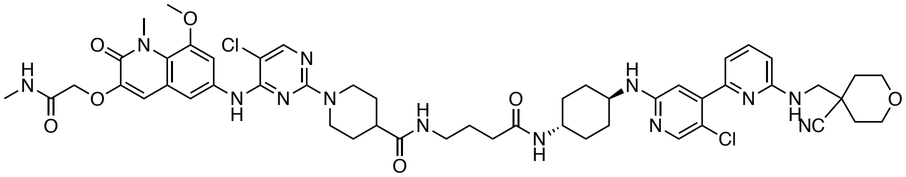

The corresponding compound was prepared following general procedure D using CDK9 inhibitor **NVP-NH₂** (0.013 mg, 0.029 mmol, 1.0 equiv) and 3-((*tert*-butoxycarbonyl)amino)butanoic acid as linker (7.0 mg, 0.035 mmol, 1.2 equiv). Purification by reverse-phase HPLC (15-90% MeOH in water) afforded the product (4.0 mg, 14% yield).

**$^1$H NMR** (500 MHz, DMSO-$d_6$) δ = 9.06 (s, 1H), 8.11 (s, 1H), 8.07 (s, 1H), 7.96 (d, *J* = 4.8 Hz, 1H), 7.81 (t, *J* = 5.7 Hz, 1H), 7.71 (d, *J* = 7.7 Hz, 1H), 7.56 – 7.50 (m, 3H), 7.22 (s, 1H), 7.01 (s, 1H), 6.80 (d, *J* = 7.2 Hz, 1H), 6.75 (s, 1H), 6.72 (d, *J* = 8.5 Hz, 1H), 4.55 (s, 2H), 4.47 (d, *J* = 13.0 Hz, 2H), 3.94 – 3.83 (m, 10H), 3.66 (d, *J* = 5.3 Hz, 2H), 3.60 (d, *J* = 8.7 Hz, 1H), 3.51 (d, *J* = 1.4 Hz, 1H), 3.46 (td, *J* = 12.0, 2.0 Hz, 2H), 3.01 (q, *J* = 6.7 Hz, 2H), 2.93 (t, *J* = 12.6 Hz, 2H), 2.65 (d, *J* = 4.6 Hz, 3H), 2.45 – 2.34 (m, 1H), 2.03 (t, *J* = 7.5 Hz, 2H), 1.96 (s, 2H), 1.88 – 1.45 (m, 12H), 1.26 (d, *J* = 6.5 Hz, 4H). **LC-MS:** m/z 1038.63 [M+1]$^+$.

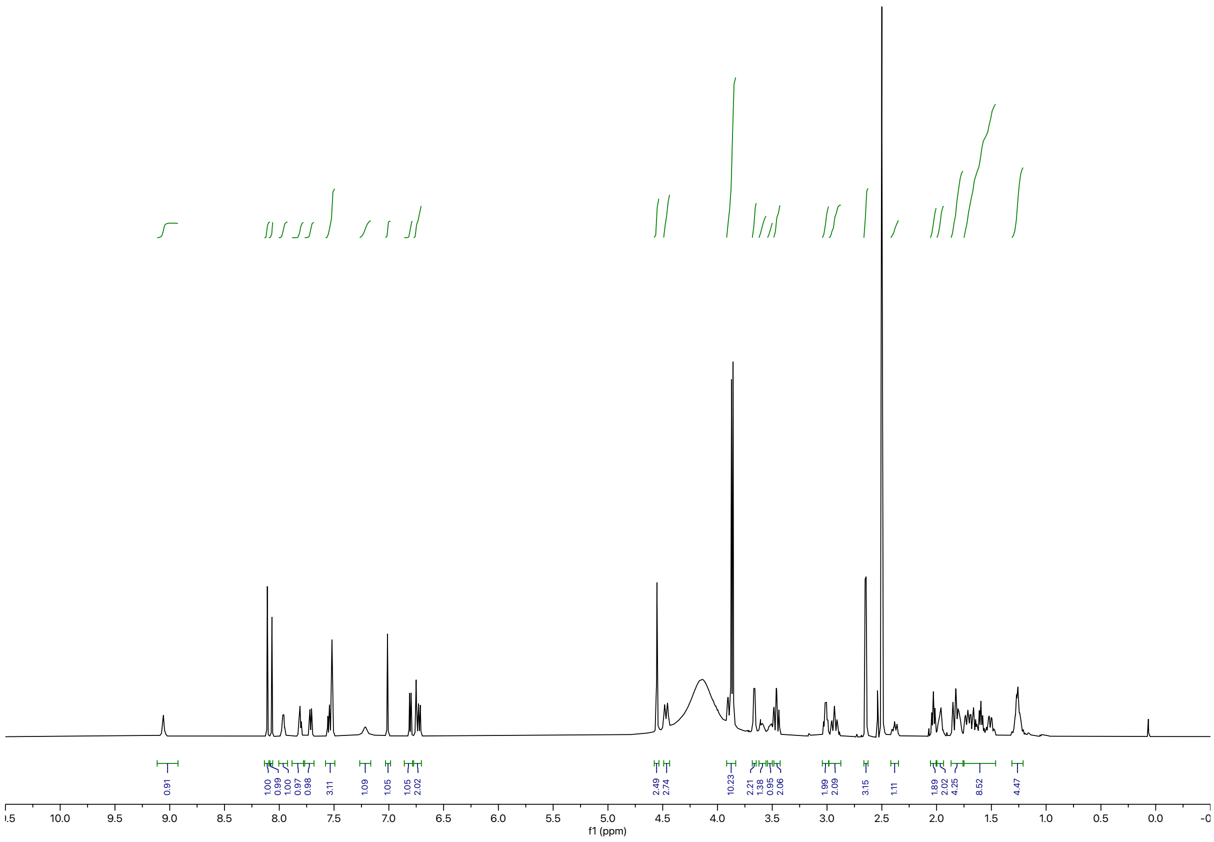

Synthesis of **RCS-03-098**

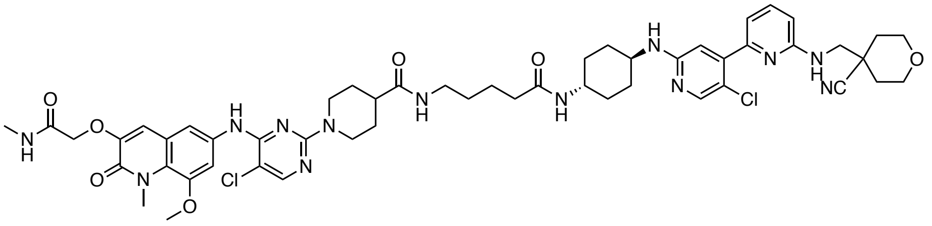

The corresponding compound was prepared following general procedure D using CDK9 inhibitor **NVP-NH₂** (0.014 mg, 0.031 mmol, 1.0 equiv) and 5-((*tert*-butoxycarbonyl)amino)pentanoic acid as linker (7.6 mg, 0.037 mmol, 1.2 equiv). Purification by reverse-phase HPLC (15-90% MeOH in water) afforded the product (4.3 mg, 13% yield).

**$^1$H NMR** (500 MHz, DMSO-$d_6$) δ = 9.10 (s, 1H), 8.11 (s, 1H), 8.07 (d, *J* = 2.2 Hz, 1H), 7.96 (d, *J* = 5.0 Hz, 1H), 7.79 (t, *J* = 5.6 Hz, 1H), 7.68 (d, *J* = 7.8 Hz, 1H), 7.58 – 7.47 (m, 3H), 7.23 (s, 1H), 7.01 (s, 1H), 6.81 (dd, *J* = 7.2, 2.8 Hz, 1H), 6.76 (s, 1H), 6.73 (d, *J* = 8.4 Hz, 1H), 4.55 (s, 2H), 4.45 (d, *J* = 13.1 Hz, 2H), 3.94 – 3.83 (m, 10H), 3.67 (d, *J* = 5.1 Hz, 2H), 3.60 (d, *J* = 9.6 Hz, 1H), 3.47 (tt, *J* = 12.1, 6.2 Hz, 3H), 3.01 (q, *J* = 6.5 Hz, 2H), 2.94 (t, *J* = 12.5 Hz, 2H), 2.65 (d, *J* = 4.6 Hz, 3H), 2.42 – 2.35 (m, 1H), 2.03 (t, *J* = 7.3 Hz, 2H), 1.96 (s, 2H), 1.84 (d, *J* = 13.5 Hz, 2H), 1.80 (s, 2H), 1.75 – 1.62 (m, 4H), 1.48 (dq, *J* = 22.8, 8.9 Hz, 4H), 1.36 (q, *J* = 7.3 Hz, 2H), 1.25 (dd, *J* = 10.8, 7.8 Hz, 4H). **LC-MS:** m/z 1053.64 $[M+1]^+$.

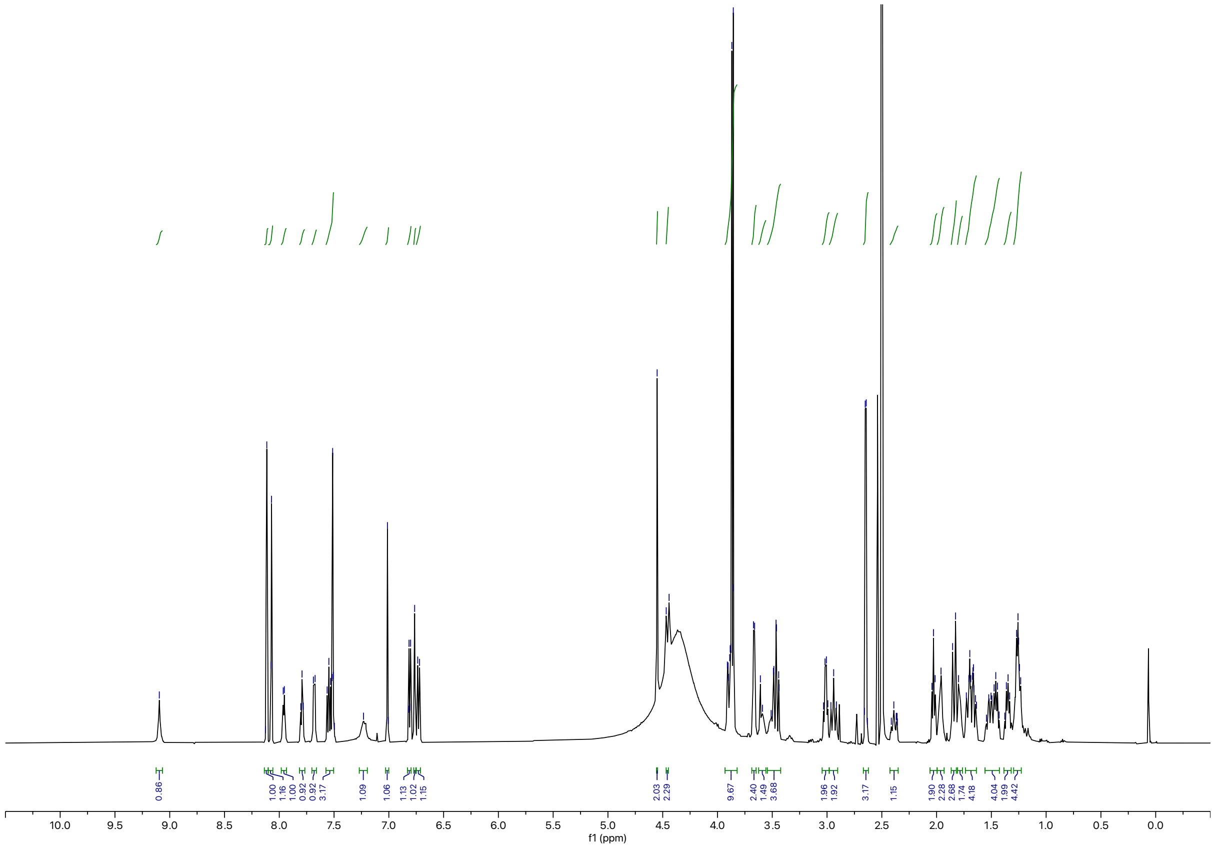

Synthesis of **BAK-04-176**

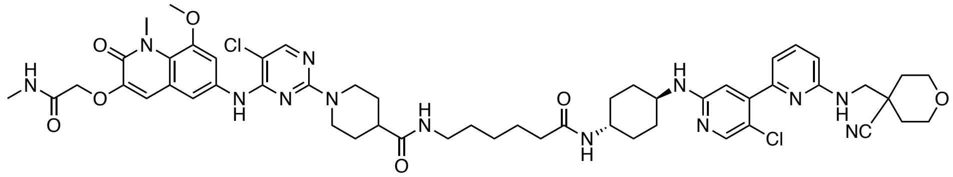

The corresponding compound was prepared following general procedure D using CDK9 inhibitor **NVP-NH₂** (8.8 mg, 0.02 mmol, 1.0 equiv) and 6-((*tert*-butoxycarbonyl)amino)hexanoic acid as linker (5.5 mg, 0.024 mmol, 1.2 equiv). Purification by reverse-phase HPLC (15-90% MeOH in water) afforded the product (6.0 mg, 40% yield). **$^1$H NMR** (500 MHz, DMSO-$d_6$) δ = 8.96 (s, 1H), 8.08 (s, 1H), 8.04 (s, 1H), 7.97 (d, *J* = 4.9 Hz, 1H), 7.77 (t, *J* = 5.6 Hz, 1H), 7.67 (d, *J* = 7.8 Hz, 1H), 7.56 – 7.49 (m, 3H), 7.18 (s, 1H), 7.01 (s, 1H), 6.79 (d, J = 7.2 Hz, 1H), 6.73 – 6.68 (m, 2H), 4.55 (s, 2H), 4.49 (d, *J* = 12.9 Hz, 2H), 3.92 – 3.85 (m, 10H, assigned by HSQC), 3.66 (d, *J* = 5.8 Hz, 2H, assigned by HSQC), 3.62 (br, 1H), 3.51 (br, 1H), 3.46 (td, J = 12.0, 2.0 Hz, 2H), 3.00 (q, *J* = 6.8 Hz, 2H), 2.91 (t, *J* = 12.5 Hz, 2H), 2.65 (d, *J* = 4.7 Hz, 3H), 2.44 – 2.33 (m, 1H), 2.11 – 1.94 (m, 4H), 1.88 – 1.73 (m, 4H), 1.74 – 1.59 (m, 4H), 1.47 (td, *J* = 14.7, 8.3 Hz, 4H), 1.37 (p, *J* = 7.1 Hz, 2H), 1.23 (dt, *J* = 22.3, 8.9 Hz, 6H). **LC-MS:** m/z 1068.66 [M+1]$^+$.

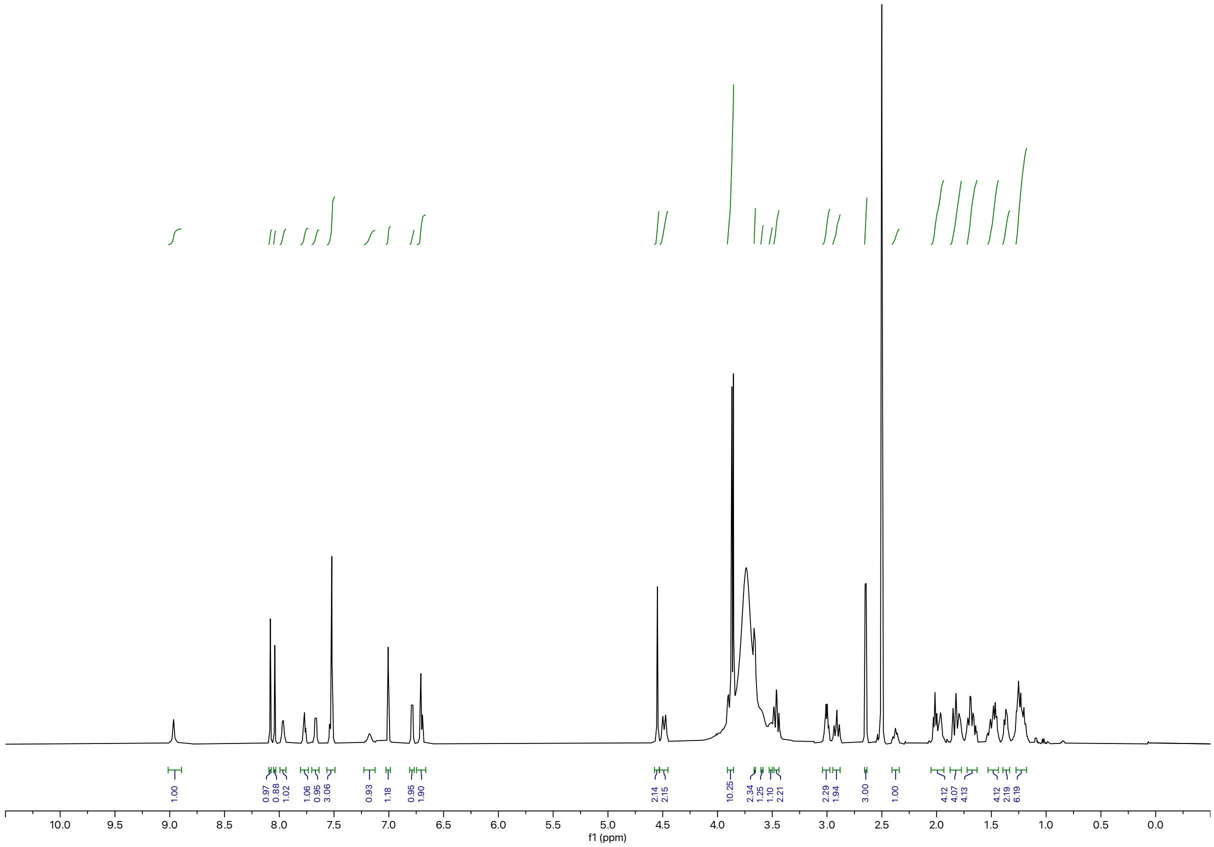

Synthesis of **BAK-04-175**

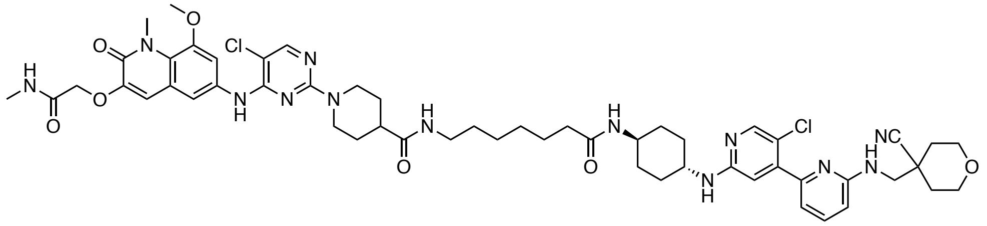

The corresponding compound was prepared following general procedure D using CDK9 inhibitor **NVP-NH₂** (8.8 mg, 0.020 mmol, 1.2 equiv) and 7-((*tert*-butoxycarbonyl)amino)heptanoic acid as linker (5.9 mg, 0.024 mmol, 1.2 equiv). Purification by reverse-phase HPLC (15-90% MeOH in water) afforded the product (4.1 mg, 30% yield).

**$^1$H NMR** (500 MHz, DMSO-d$_6$) δ = 8.95 (s, 1H), 8.08 (s, 1H), 8.04 (s, 1H), 7.98 (s, 1H), 7.77 (t, *J* = 5.6 Hz, 1H), 7.67 (d, *J* = 7.8 Hz, 1H), 7.52 (s, 3H), 7.16 (s, 1H), 7.01 (s, 1H), 6.78 (d, J = 7.2 Hz, 1H), 6.70 (t, *J* = 4.2 Hz, 2H), 4.55 (s, 2H), 4.49 (d, *J* = 13.0 Hz, 2H), 3.86 (d, *J* = 6.7 Hz, 10H), 3.66 (d, *J* = 6.0 Hz, 2H, assigned by HSQC), 3.59 (br, 1H, assigned by HSQC), 3.51 (1H, assigned by HSQC), 3.45 (dd, *J* = 12.1, 2.1 Hz, 2H, assigned by HSQC), 3.00 (q, *J* = 6.6 Hz, 2H), 2.95 – 2.87 (m, 2H), 2.65 (d, *J* = 4.7 Hz, 3H), 2.42 – 2.34 (m, 1H), 2.05 – 1.92 (m, 4H), 1.87 – 1.76 (m, 4H), 1.72 – 1.62 (m, 4H), 1.48 (dt, *J* = 20.1, 6.7 Hz, 4H), 1.40 – 1.33 (m, 2H), 1.28 – 1.22 (m, 8H). **LC-MS:** m/z 1082.70 [M+1]$^+$.

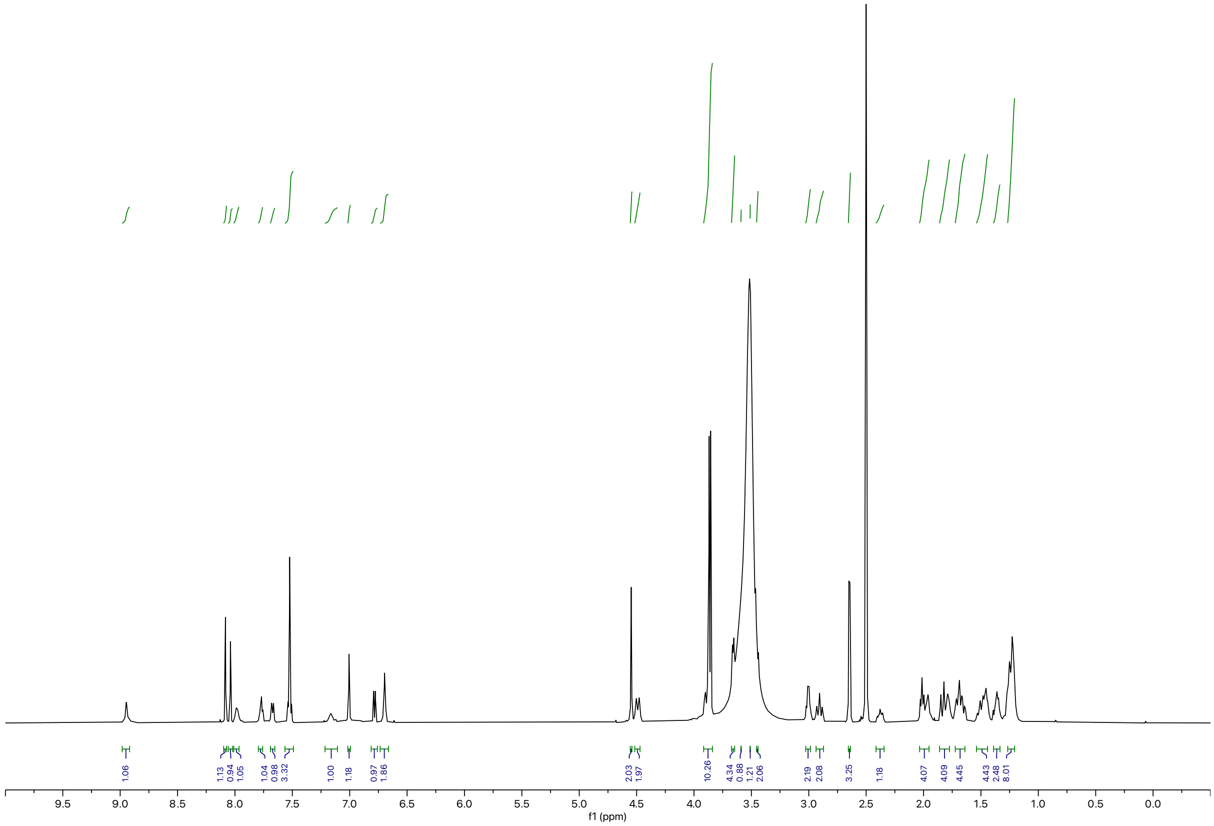

Synthesis of **BAK-04-174**

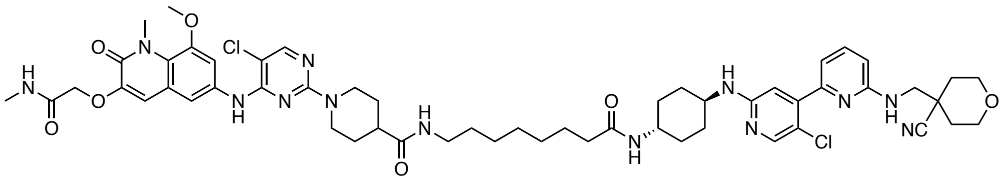

The corresponding compound was prepared following general procedure D using CDK9 inhibitor **NVP-NH2** (8.8 mg, 0.020 mmol, 1.0 equiv) and 8-((*tert*-butoxycarbonyl)amino)octanoic acid as linker (6.2 mg, 0.024 mmol, 1.2 equiv). Purification by reverse-phase HPLC (15-90% MeOH in water) afforded the product (3.1 mg, 20% yield).

**$^1$H NMR** (500 MHz, DMSO-d$_6$) δ = 8.92 (s, 1H), 8.08 (s, 1H), 8.04 (s, 1H), 7.98 (d, *J* = 4.8 Hz, 1H), 7.78 (t, *J* = 5.6 Hz, 1H), 7.67 (d, *J* = 7.8 Hz, 1H), 7.56 – 7.50 (m, 3H), 7.16 (s, 1H), 7.01 (s, 1H), 6.79 (d, *J* = 7.2 Hz, 1H), 6.70 (d, *J* = 8.3 Hz, 2H), 4.56 (s, 2H), 4.50 (d, *J* = 13.0 Hz, 2H), 3.87 (m, 10H), 3.67 (2H, assigned by HSQC) 3.61 (1H, assigned by HSQC), 3.52 (1H, assigned by HSQC) 3.52 – 3.43 (2H, assigned by HSQC), 3.01 (q, *J* = 6.6 Hz, 2H), 2.91 (t, *J* = 12.6 Hz, 2H), 2.65 (d, *J* = 4.7 Hz, 3H), 2.43 – 2.34 (m, 1H), 2.08 – 1.91 (m, 4H), 1.87 – 1.76 (m, 4H), 1.73 – 1.63 (m, 4H), 1.55 – 1.44 (m, 4H), 1.43 – 1.33 (m, 2H), 1.30 – 1.20 (m, 10H). **LC-MS:** m/z 1096.60 [M+1]$^+$.

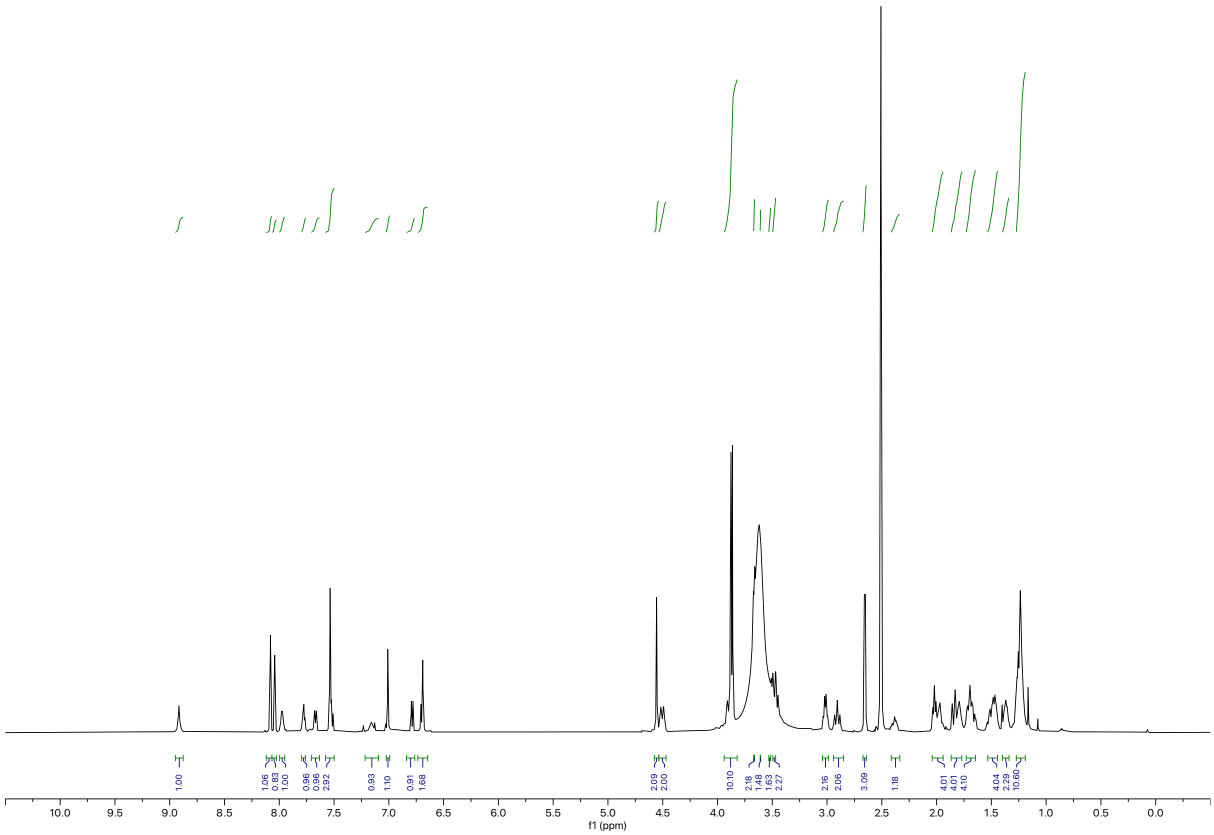

**Synthesis of Atuveciclib (BAY-1143572) TCIPs**

General Procedure E

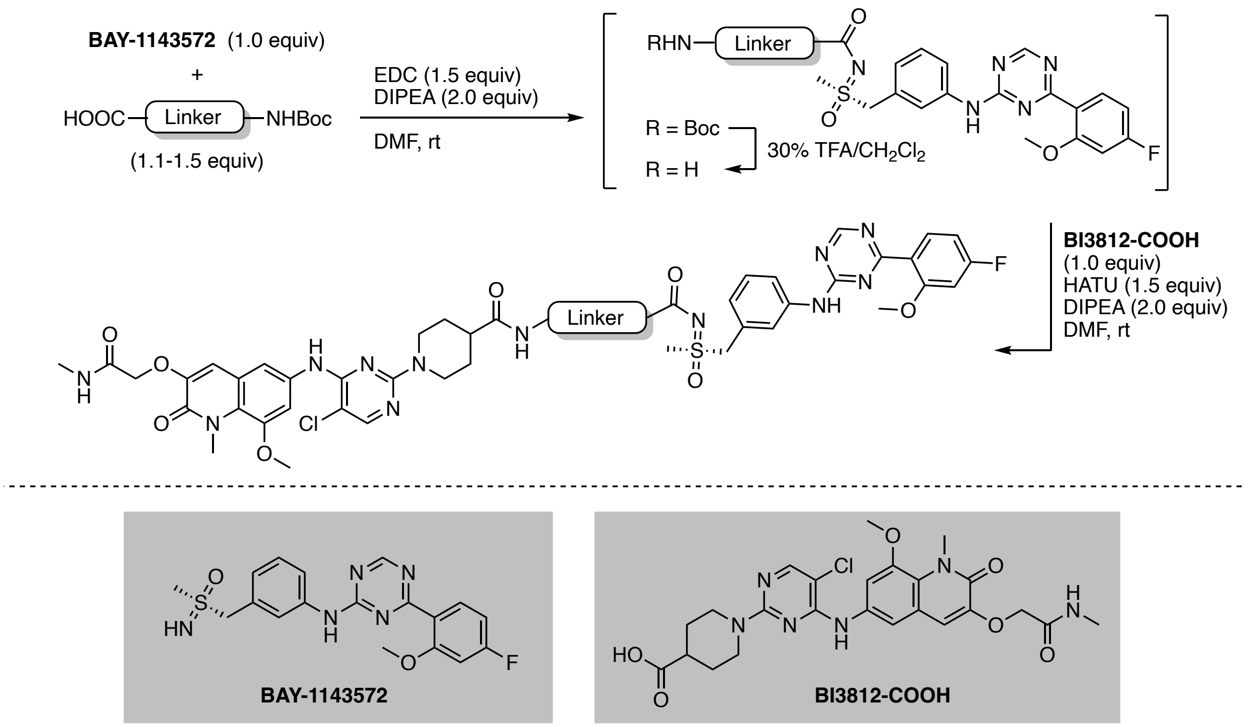


To a solution of **BAY-1143572**[9] (1.0 equiv) and linker (1.2 equiv) in DMF (0.2 M) was added EDC (1.5 equiv) and DIPEA (2.0 equiv). The resulting yellow solution was stirred at ambient temperature for 24 h. Upon full consumption of starting material, as judged by LC-MS and/or TLC analysis, the reaction mixture was diluted with water and EtOAc. After separation of the organic layer, the aqueous phase was extracted with EtOAc (3x) and the combined organic extracts were washed with brine, dried over $Na_2SO_4$, and concentrated under reduced pressure. The resulting intermediate was taken up in 30% v/v TFA in $CH_2Cl_2$ and the resulting solution was stirred at ambient temperature until LC-MS showed quantitative formation of free amine intermediate. All volatiles were removed under reduced pressure. The intermediate was used without further purification, dissolved in DMF (0.2 M), and added to a solution of **BI3812-COOH** (1.0 equiv), HATU (1.5 equiv), and DIPEA (2.0 equiv). The reaction mixture was purified by preparative reverse-phase HPLC to afford the product upon lyophilization.

Synthesis of **BAK-04-105**

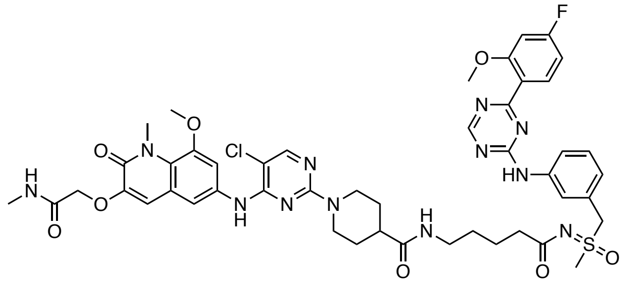

9The corresponding compound was prepared following general procedure E using CDK9 inhibitor **BAY-1143572** (3.9 mg, 0.010 mmol, 1.0 equiv) and 5-((*tert*-butoxycarbonyl)amino)pentanoic acid as linker (2.6 mg, 0.012 mmol, 1.2 equiv). Purification by reverse-phase HPLC (15-90% MeOH in water) afforded the product (2.1 mg, 10% yield). **$^1$H NMR** (500 MHz, DMSO-d$_6$) δ = 10.40 (s, 1H), 8.78 (d, *J* = 8.8 Hz, 2H), 8.05 (s, 1H), 7.94 (d, *J* = 4.9 Hz, 1H), 7.67 (br, 1H), 7.75 (t, *J* = 5.8 Hz, 2H), 7.53 (q, *J* = 2.3 Hz, 2H), 7.38 (t, *J* = 7.9 Hz, 1H), 7.15 – 7.05 (m, 2H), 6.98 (s, 1H), 6.89 (td, *J* = 8.4, 2.4 Hz, 1H), 6.54 (br, 1H). 4.82 (s, 2H), 4.52 (d, *J* = 18.6 Hz, 4H), 3.85 (d, *J* = 8.5 Hz, 9H), 3.14 (s, 3H), 2.98 (q, *J* = 6.5 Hz, 2H), 2.87 (t, *J* = 12.9 Hz, 2H), 2.64 (d, *J* = 4.7 Hz, 3H), 2.43 – 2.32 (m, 1H), 2.16 (dt, *J* = 14.1, 7.3 Hz, 2H), 1.69 (d, *J* = 12.8 Hz, 2H), 1.47 (4H, assigned by HSQC), 1.33 (2H, assigned by HSQC). **LC-MS:** m/z 999.47 [M+1]$^+$.

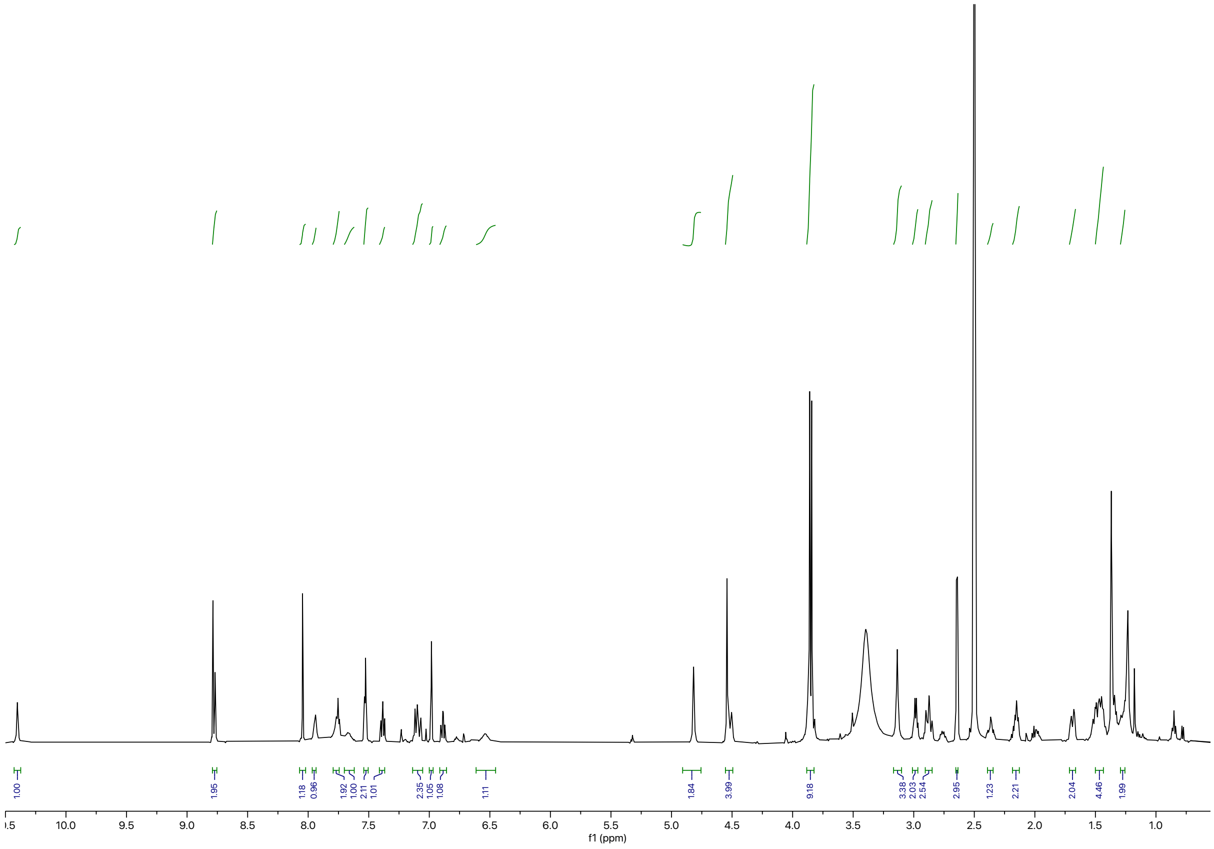

Synthesis of **BAK-04-127**

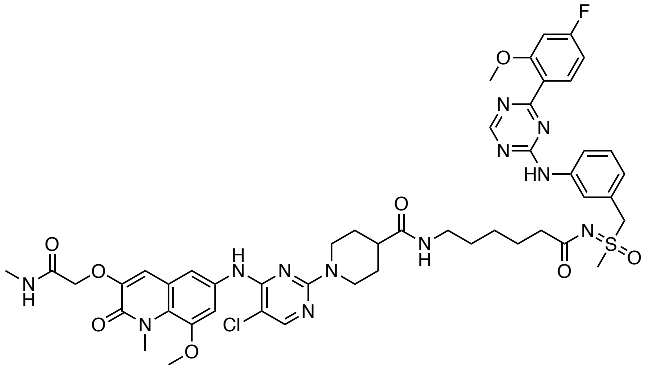

The corresponding compound was prepared following general procedure D using CDK9 inhibitor **BAY-1143572** (3.9 mg, 0.010 mmol, 1.0 equiv) and 6-((*tert*-butoxycarbonyl)amino)hexanoic acid as linker (2.8 mg, 0.012 mmol, 1.2 equiv). Purification by reverse-phase HPLC (15-90% MeOH in water) afforded the product (6.6 mg, 50% yield). **$^1$H NMR** (500 MHz, DMSO-d$_6$) δ = 10.43 (s, 1H), 8.90 (s, 1H), 8.79 (s, 1H), 8.07 (s, 1H), 7.94 (d, *J* = 4.8 Hz, 1H), 7.79 (s, 1H), 7.74 (t, *J* = 5.6 Hz, 1H), 7.52 (t, *J* = 1.7 Hz, 2H), 7.43 – 7.36 (m, 1H), 7.15 – 7.06 (m, 2H), 6.99 (s, 1H), 6.89 (td, *J* = 8.4, 2.4 Hz, 1H), 4.82 (s, 2H), 4.57 – 4.46 (m, 4H), 3.85 (d, *J* = 7.9 Hz, 9H), 3.14 (s, 3H), 2.98 (q, *J* = 6.6 Hz, 2H), 2.89 (t, *J* = 12.5 Hz, 2H), 2.64 (d, *J* = 4.6 Hz, 3H), 2.41 – 2.31 (m, 1H), 2.13 (t, *J* = 7.5 Hz, 2H), 1.69 (d, *J* = 12.8 Hz, 2H), 1.55 – 1.40 (m, 4H), 1.31 (qd, *J* = 14.2, 7.0 Hz, 2H), 1.25 – 1.15 (2H, assigned by HSQC). **LC-MS:** m/z 1013.62 [M+1]$^+$.

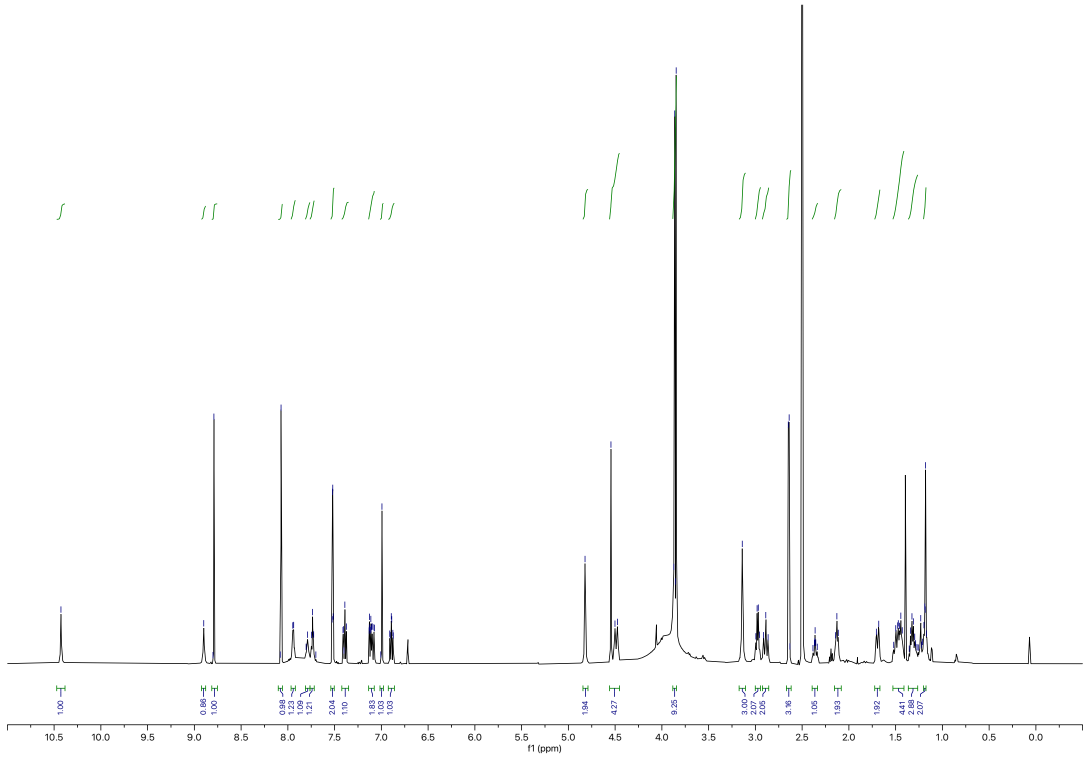

Synthesis of **RCS-03-019**

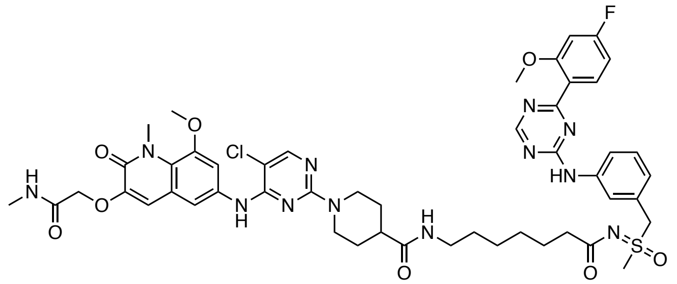

The corresponding compound was prepared following general procedure D using CDK9 inhibitor **BAY-1143572** (6.6 mg, 0.017 mmol, 1.0 equiv) and 7-((*tert*-butoxycarbonyl)amino)heptanoic acid as linker (5.0 mg, 0.020 mmol, 1.2 equiv). Purification by reverse-phase HPLC (15-90% MeOH in water) afforded the product as yellowish powder (0.014 g, 78% yield).

**$^1$H NMR** (500 MHz, DMSO-d$_6$) δ = 10.48 (s, 1H), 9.11 (s, 1H), 8.80 (s, 1H), 8.11 (s, 1H), 7.95 (d, *J* = 4.8 Hz, 1H), 7.80 (s, 1H), 7.75 (t, *J* = 5.6 Hz, 1H), 7.50 (s, 2H), 7.40 (d, *J* = 7.9 Hz, 1H), 7.17 – 7.06 (m, 2H), 7.01 (s, 1H), 6.90 (td, *J* = 8.4, 2.4 Hz, 1H), 4.83 (s, 2H), 4.55 (s, 2H), 4.45 (d, *J* = 13.2 Hz, 2H), 3.85 (d, *J* = 6.6 Hz, 9H), 3.14 (s, 3H), 3.04 – 2.88 (m, 4H), 2.64 (d, *J* = 4.6 Hz, 3H), 2.43 – 2.34 (m, 1H), 2.12 (t, *J* = 7.2 Hz, 2H), 1.71 (d, *J* = 12.7 Hz, 2H), 1.55 – 1.39 (m, 4H), 1.32 (t, *J* = 7.0 Hz, 2H), 1.21 – 1.15 (m, 4H). **$^{19}$F NMR** (471 MHz, DMSO-d$_6$) δ = -74.72. **LC-MS:** m/z 1027.57 [M+1]$^+$.

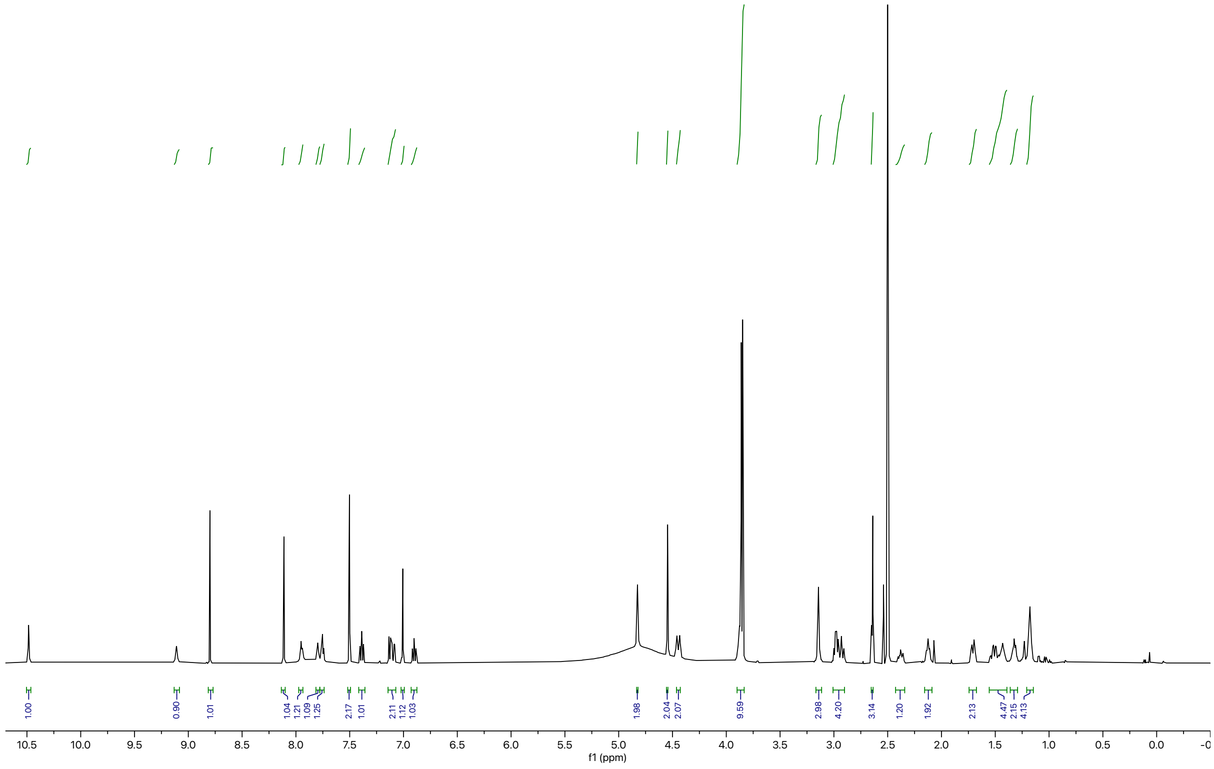

Synthesis of **RCS-03-018**

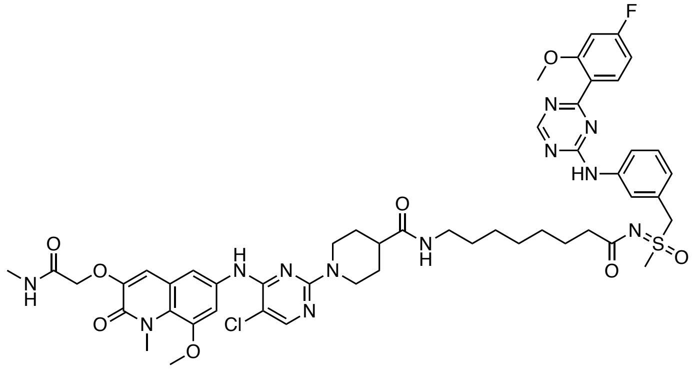


The corresponding compound was prepared following general procedure D using CDK9 inhibitor **BAY-1143572** (3.4 mg, 0.015 mmol, 1.0 equiv) and 8-((*tert*-butoxycarbonyl)amino)octanoic acid as linker (4.7 mg, 0.018 mmol, 1.2 equiv). Purification by reverse-phase HPLC (15-90% MeOH in water) afforded the product (2.4 mg, 15% yield).

**$^{1}$H NMR** (500 MHz, DMSO-d$_6$) δ = 10.44 (s, 1H), 8.96 (s, 1H), 8.80 (s, 1H), 8.09 (s, 1H), 7.98 (d, *J* = 5.0 Hz, 1H), 7.81 (s, 1H), 7.76 (t, *J* = 5.6 Hz, 1H), 7.53 (s, 2H), 7.39 (t, *J* = 7.9 Hz, 1H), 7.16 – 7.07 (m, 2H), 7.01 (s, 1H), 6.90 (td, *J* = 8.4, 2.4 Hz, 1H), 4.83 (s, 2H), 4.55 (s, 2H), 4.49 (d, *J* = 13.1 Hz, 2H), 3.86 (d, *J* = 6.9 Hz, 9H), 3.15 (s, 3H), 2.99 (q, *J* = 6.6 Hz, 2H), 2.91 (t, *J* = 12.6 Hz, 2H), 2.65 (d, *J* = 4.6 Hz, 3H), 2.43 – 2.34 (m, 1H), 2.12 (s, 2H), 1.70 (d, *J* = 12.7 Hz, 2H), 1.57 – 1.39 (m, 4H), 1.32 (d, *J* = 7.1 Hz, 2H), 1.17 (s, 6H). **$^{19}$F NMR** (471 MHz, DMSO-d$_6$) δ = -74.31. **LC-MS:** m/z 1041.42 $[M+1]^+$.

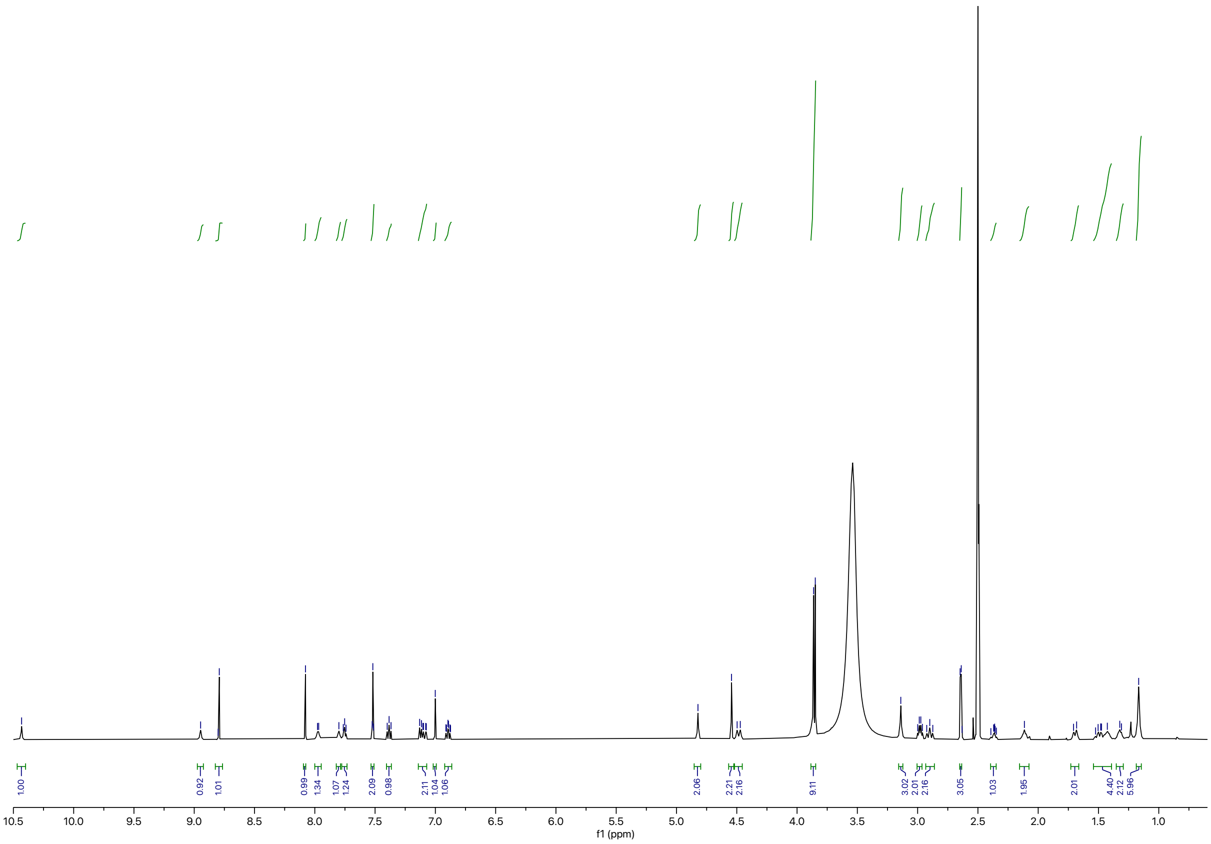
1.00
0.92
1.01
0.99
1.34
1.07
1.24
2.09
0.98
2.11
1.04
1.06
2.06
2.21
2.16
9.11
3.02
2.01
2.16
3.05
1.03
1.95
2.01
4.40
2.12
5.96
f1 (ppm)

Synthesis of **RCS-03-020**

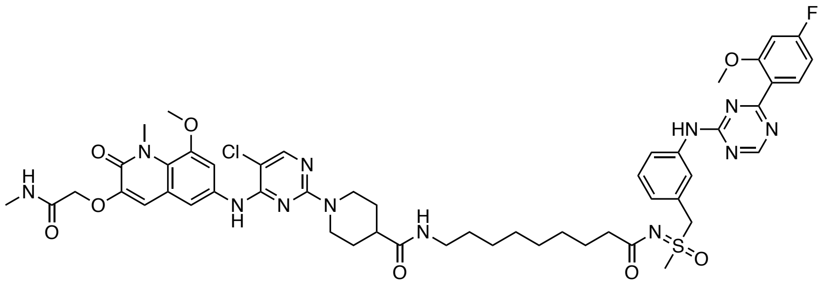

The corresponding compound was prepared following general procedure D using CDK9 inhibitor **BAY-1143572** (6.6 mg, 0.017 mmol, 1.0 equiv) and 9-((*tert*-butoxycarbonyl)amino)nonanoic acid as linker (5.6 mg, 0.020 mmol, 1.2 equiv). Purification by preparative reverse-phase HPLC (15-90% MeOH in water) afforded the product (5.3 mg, 29% yield).

**$^1$H NMR** (500 MHz, DMSO-$d_6$) δ = 10.47 (s, 1H), 9.04 (s, 1H), 8.81 (s, 1H), 8.11 (d, *J* = 1.6 Hz, 1H), 7.95 (d, *J* = 4.9 Hz, 1H), 7.81 (s, 1H), 7.76 (t, *J* = 5.7 Hz, 1H), 7.52 (s, 2H), 7.39 (t, *J* = 7.9 Hz, 1H), 7.16 – 7.08 (m, 2H), 7.01 (s, 1H), 6.91 (td, *J* = 8.4, 2.4 Hz, 1H), 4.83 (s, 2H), 4.55 (s, 2H), 4.47 (d, *J* = 13.0 Hz, 2H), 3.87 (d, *J* = 7.2 Hz, 9H), 3.15 (d, *J* = 1.6 Hz, 3H), 2.99 (q, *J* = 6.6 Hz, 2H), 2.93 (t, *J* = 12.6 Hz, 2H), 2.65 (d, *J* = 4.7 Hz, 4H), 2.42 – 2.34 (m, 1H), 2.12 (s, 2H), 1.71 (d, *J* = 11.8 Hz, 2H), 1.56 – 1.47 (m, 2H), 1.43 (s, 2H), 1.34 (t, *J* = 6.8 Hz, 2H), 1.28 – 1.23 (m, 2H), 1.20 – 1.15 (m, 6H). **$^{19}$F NMR** (471 MHz, DMSO-$d_6$) δ = -74.61. **LC-MS:** m/z 1055.53 $[M+1]^+$.

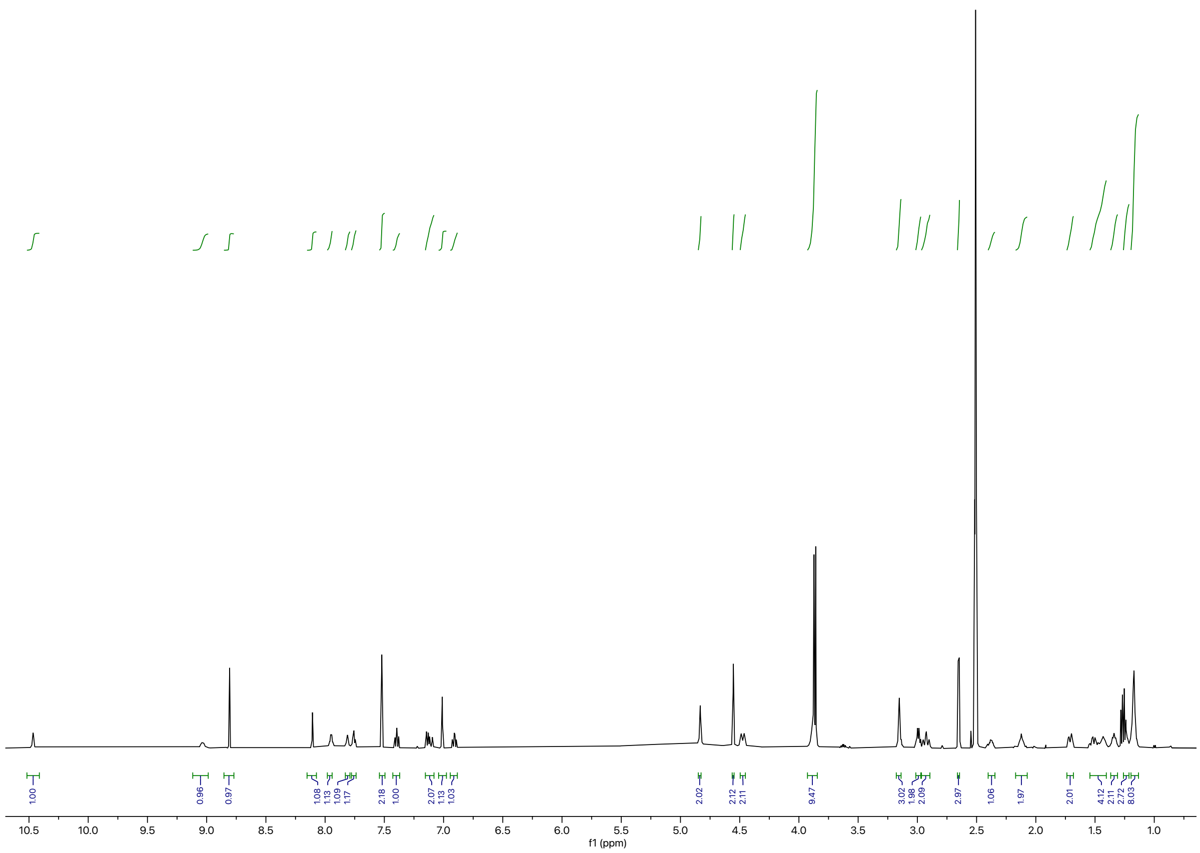

f1 (ppm)

**Synthesis of KB-0742 TCIPs**

General Procedure F

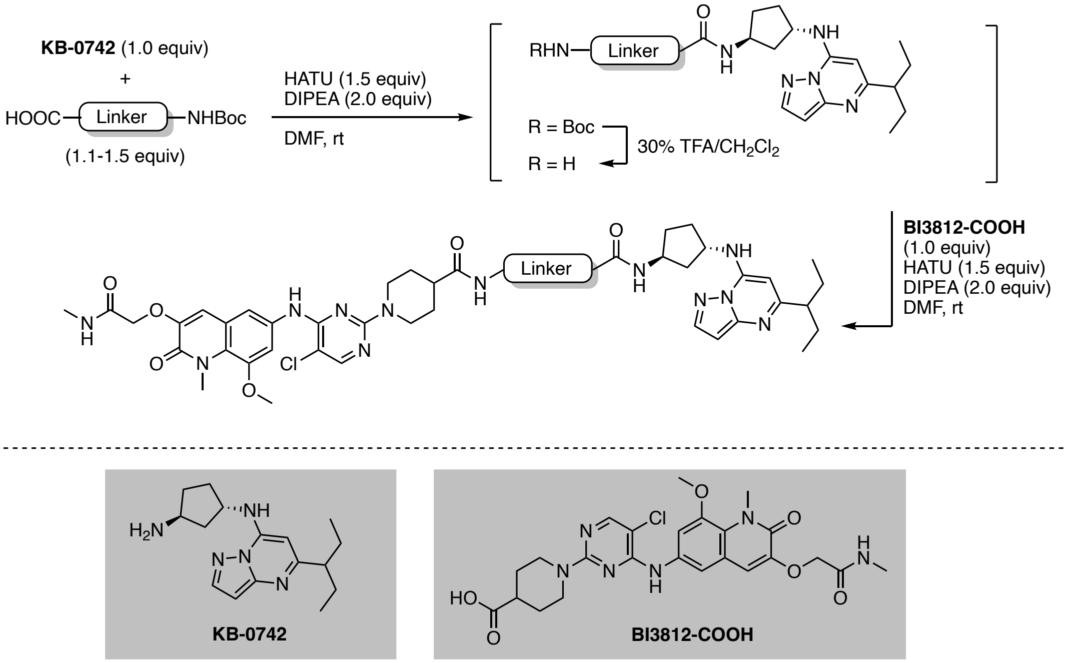


To a solution of **KB-0742**[10] (1.0 equiv) and linker (1.2 equiv) in DMF (0.2 M) was added HATU (1.5 equiv) and DIPEA (2.0 equiv). The resulting yellow solution was stirred at ambient temperature. Upon full consumption of starting material as judged by LC-MS and/or TLC analysis the reaction mixture was diluted with water and EtOAc. After separation of the organic layer, the aqueous phase was extracted with EtOAc (3x) and the combined organic extracts were washed with brine, dried over $Na_2SO_4$, and concentrated under reduced pressure. The resulting intermediate was taken up in 30% v/v TFA in $CH_2Cl_2$ and the resulting solution was stirred at ambient temperature until LC-MS showed quantitative formation of free amine intermediate. All volatiles were removed under reduced pressure. The intermediate was used without further purification, dissolved in DMF (0.2 M), and added to a solution of **BI3812-COOH** (1.0 equiv), HATU (1.5 equiv), and DIPEA (2.0 equiv). The reaction mixture was purified by preparative reverse-phase HPLC to afford the product upon lyophilization.

Synthesis of **RCS-03-010**

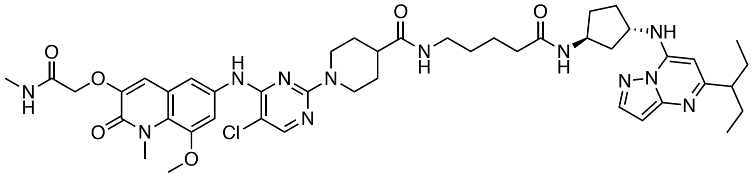

The corresponding compound was prepared following general procedure F using CDK9 inhibitor **KB-0742** (6.8 mg, 0.019 mmol, 1.0 equiv) and 5-((*tert*-butoxycarbonyl)amino)pentanoic acid as linker (5.0 mg, 0.023 mmol, 1.2 equiv). Purification by reverse-phase HPLC (15-90% MeOH in water) afforded the product (0.013 g, 73% yield).

**$^1$H NMR** (500 MHz, DMSO-d$_6$) δ = 9.80 (s, 1H), 8.92 (s, 1H), 8.29 (d, *J* = 2.2 Hz, 1H), 8.07 (s, 1H), 7.97 (d, *J* = 4.9 Hz, 1H), 7.85 (d, *J* = 7.2 Hz, 1H), 7.80 (t, *J* = 5.7 Hz, 1H), 7.53 (q, *J* = 2.3 Hz, 2H), 7.00 (s, 1H), 6.54 (d, *J* = 2.1 Hz, 1H), 6.48 (s, 1H), 4.55 (s, 2H), 4.49 (d, *J* = 12.5 Hz, 3H), 4.25 (h, *J* = 7.2 Hz, 1H), 3.86 (d, *J* = 8.8 Hz, 6H), 3.02 (q, *J* = 6.7 Hz, 2H), 2.90 (t, *J* = 12.6 Hz, 2H), 2.68 – 2.58 (m, 4H), 2.43 – 2.33 (m, 1H), 2.22 – 2.14 (m, 1H), 2.07 (dt, *J* = 21.0, 6.9 Hz, 4H), 1.90-1.83 (m, 1H), 2.12 – 1.64 (m, 7H), 1.55 – 1.42 (m, 5H), 1.39 – 1.31 (m, 2H), 0.82 (t, *J* = 7.4 Hz, 6H). **LC-MS:** m/z 899.41 [M+1]$^+$.

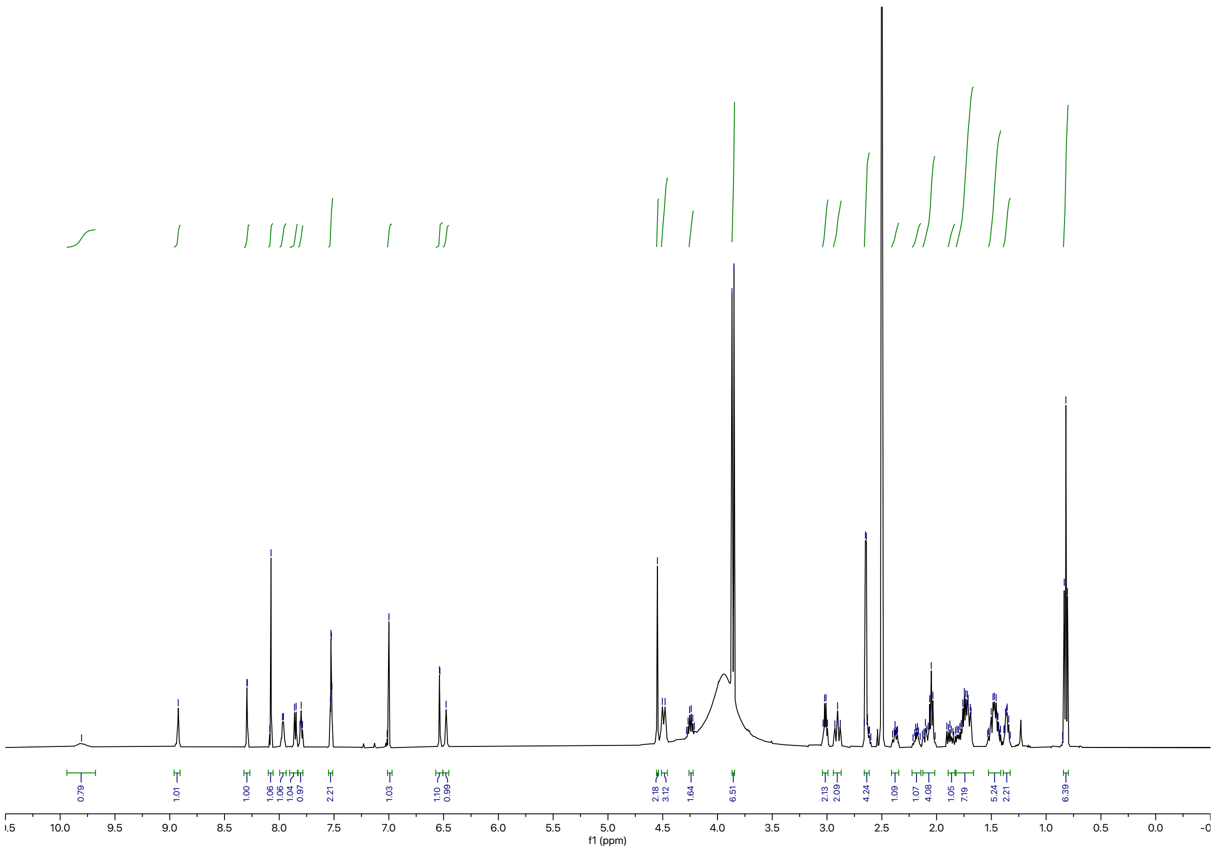

Synthesis of **RCS-03-011**

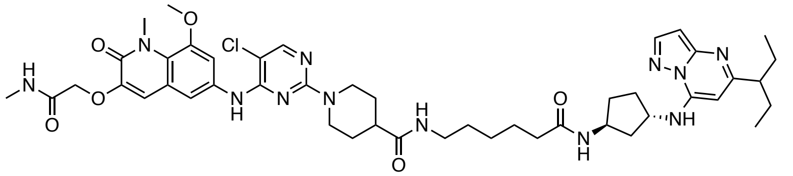

The corresponding compound was prepared following general procedure F using CDK9 inhibitor **KB-0742** (7.0 mg, 0.019 mmol, 1.0 equiv) and 6-((*tert*-butoxycarbonyl)amino)hexanoic acid as linker (5.4 mg, 0.023 mmol, 1.2 equiv). Purification by reverse-phase HPLC (15-90% MeOH in water) afforded the product (0.016 g, 88% yield).

**$^{1}$H NMR** (500 MHz, DMSO-d$_6$) δ = 9.89 (s, 1H), 9.00 (s, 1H), 8.31 (d, *J* = 2.2 Hz, 1H), 8.09 (s, 1H), 7.95 (t, *J* = 4.8 Hz, 1H), 7.83 (d, *J* = 7.2 Hz, 1H), 7.78 (t, *J* = 5.7 Hz, 1H), 7.55 – 7.49 (m, 2H), 7.00 (s, 1H), 6.55 (d, *J* = 2.2 Hz, 1H), 6.50 (s, 1H), 4.55 (s, 2H), 4.47 (d, *J* = 13.6 Hz, 3H), 4.24 (p, *J* = 7.2 Hz, 1H), 3.86 (d, *J* = 8.2 Hz, 6H), 3.01 (q, *J* = 6.7 Hz, 2H), 2.91 (t, *J* = 12.5 Hz, 2H), 2.65 (d, *J* = 4.7 Hz, 4H), 2.41 – 2.34 (m, 1H), 2.19 (ddd, *J* = 12.6, 8.0, 4.3 Hz, 1H), 2.14 – 2.07 (m, 1H), 2.04 (dd, *J* = 9.5, 5.5 Hz, 3H), 1.88 (dt, *J* = 14.3, 7.6 Hz, 1H), 1.83 – 1.78 (m, 1H), 1.77 – 1.66 (m, 6H), 1.44 (dqd, *J* = 45.0, 15.5, 5.6 Hz, 7H), 1.26 – 1.17 (m, 2H), 0.82 (t, *J* = 7.4 Hz, 6H). **LC-MS:** m/z 913.41 [M+1]$^{+}$.

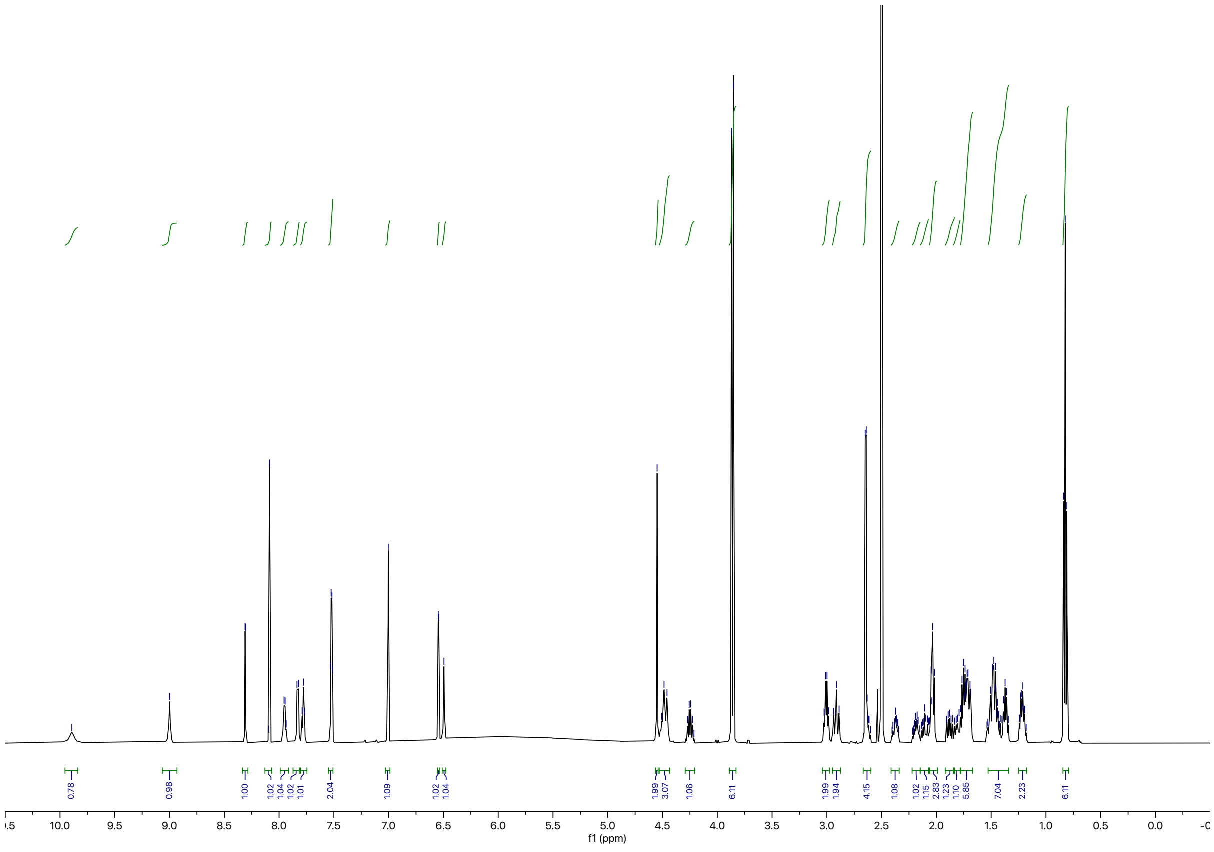

Synthesis of **BAK-04-098**

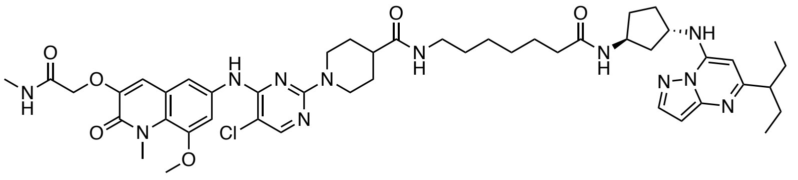

The corresponding compound was prepared following general procedure F using CDK9 inhibitor **KB-0742** (8.6 mg, 0.024 mmol, 1.0 equiv) and 7-((*tert*-butoxycarbonyl)amino)heptanoic acid as linker (7.1 mg, 0.029 mmol, 1.2 equiv). Purification by reverse-phase HPLC (15-90% MeOH in water) afforded the product (6.7 mg, 30% yield).

**$^1$H NMR** (500 MHz, DMSO-$d_6$) δ = 8.81 (s, 1H), 8.21 (s, 1H), 8.06 (s, 1H), 7.98 (s, 1H), 7.86 (d, *J* = 7.2 Hz, 1H), 7.77 (t, *J* = 5.5 Hz, 1H), 7.54 (q, *J* = 2.4 Hz, 2H), 6.99 (s, 1H), 6.47 (s, 1H), 6.34 (s, 1H), 4.53 (d, *J* = 21.3 Hz, 4H), 4.41 (s, 1H), 4.24 (q, *J* = 7.1 Hz, 1H), 3.86 (d, *J* = 8.1 Hz, 6H), 3.00 (q, *J* = 6.6 Hz, 2H), 2.94 – 2.84 (m, 2H), 2.64 (d, *J* = 4.7 Hz, 3H), 2.58 (t, *J* = 7.2 Hz, 1H), 2.41 – 2.33 (m, 1H), 2.21 – 2.13 (m, 1H), 2.04 (q, *J* = 8.5 Hz, 4H), 1.92 – 1.83 (m, 1H), 1.83 – 1.75 (m, 1H), 1.75 – 1.66 (m, 6H), 1.53 – 1.39 (m, 5H), 1.40 – 1.32 (m, 2H), 1.27 – 1.20 (m, 4H), 0.80 (t, *J* = 7.4 Hz, 6H). **LCMS:** m/z 927.61 [M+1]$^+$.

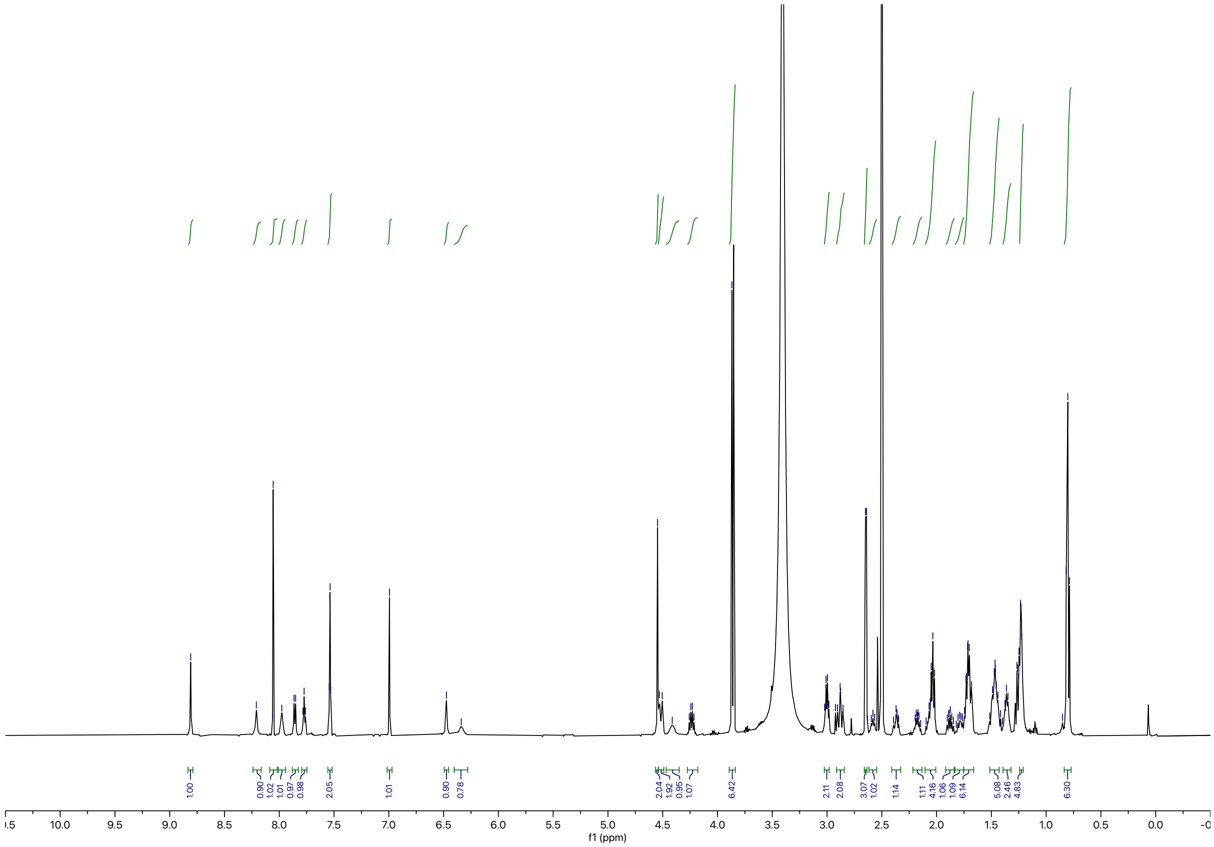

Synthesis of **BAK-04-099**

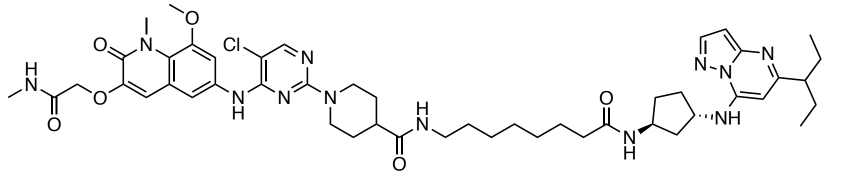

The corresponding compound was prepared following general procedure F using CDK9 inhibitor **KB-0742** (8.3 mg, 0.023 mmol, 1.0 equiv) and 8-((*tert*-butoxycarbonyl)amino)octanoic acid as linker (7.2 mg, 0.028 mmol, 1.2 equiv). Purification by reverse-phase HPLC (15-90% MeOH in water) afforded the product (6.5 mg, 30% yield).

**$^1$H NMR** (500 MHz, DMSO-d$_6$) δ = 8.79 (s, 1H), 8.17 (s, 1H), 8.05 (s, 1H), 7.98 (q, *J* = 4.6 Hz, 1H), 7.83 (d, *J* = 7.2 Hz, 1H), 7.76 (t, *J* = 5.7 Hz, 1H), 7.54 (q, *J* = 2.4 Hz, 2H), 6.99 (s, 1H), 6.44 (d, *J* = 2.2 Hz, 1H), 6.25 (s, 1H), 4.52 (d, *J* = 19.8 Hz, 4H), 4.36 (s, 1H), 4.23 (h, *J* = 7.1 Hz, 1H), 3.86 (d, *J* = 7.5 Hz, 6H), 3.00 (q, *J* = 6.6 Hz, 2H), 2.87 (td, *J* = 13.0, 2.8 Hz, 2H), 2.65 (d, *J* = 4.7 Hz, 3H), 2.57 – 2.52 (m, 2H), 2.41 – 2.32 (m, 1H), 2.18 (dtd, *J* = 12.4, 7.7, 4.3 Hz, 1H), 2.04 (dt, *J* = 14.9, 7.0 Hz, 4H), 1.87 (dt, *J* = 13.7, 7.2 Hz, 1H), 1.78 (dt, *J* = 13.3, 7.5 Hz, 1H), 1.74 – 1.64 (m, 6H), 1.47 (dddd, *J* = 20.5, 16.1, 12.3, 5.8 Hz, 6H), 1.36 (t, *J* = 6.9 Hz, 2H), 1.22 (d, *J* = 7.0 Hz, 6H), 0.79 (t, *J* = 7.4 Hz, 6H). **LC-MS:** m/z 941.71 [M+1]$^+$.

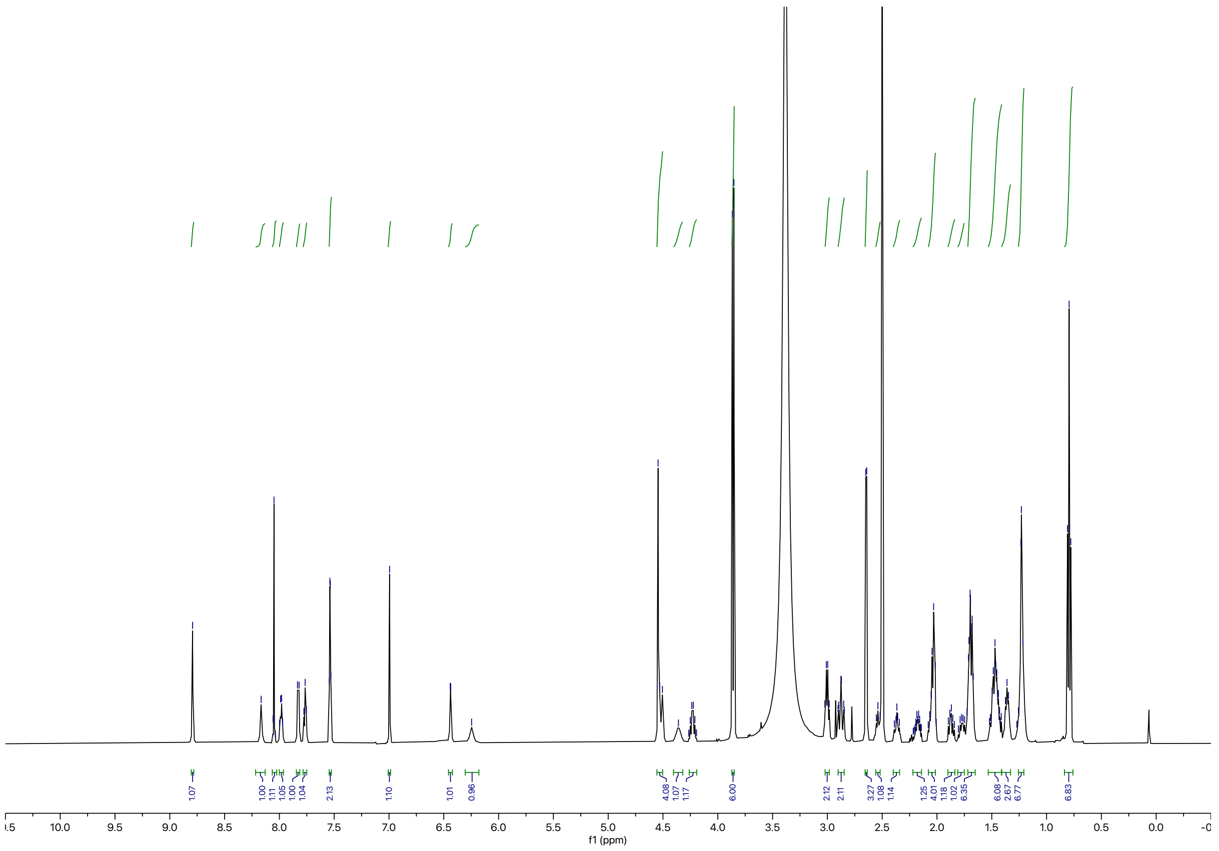

Synthesis of **BAK-04-100**

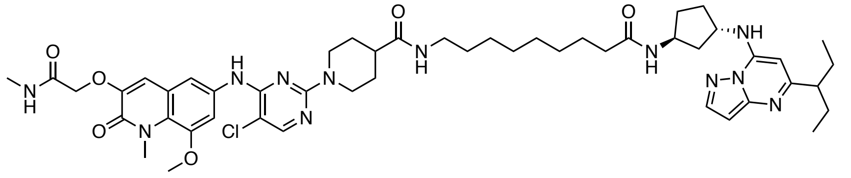

The corresponding compound was prepared following general procedure F using CDK9 inhibitor **KB-0742** (8.3 mg, 0.023 mmol, 1.0 equiv) and 9-((*tert*-butoxycarbonyl)amino)nonanoic acid as linker (7.5 mg, 0.028 mmol, 1.2 equiv). Purification by reverse-phase HPLC (15-90% MeOH in water) afforded the product (8.1 mg, 38% yield).

**$^1$H NMR** (500 MHz, DMSO-$d_6$) δ = 9.39 (s, 1H), 8.82 (s, 1H), 8.23 (d, *J* = 2.3 Hz, 1H), 8.06 (s, 1H), 7.99 (q, *J* = 4.7 Hz, 1H), 7.83 (d, *J* = 7.2 Hz, 1H), 7.76 (t, *J* = 5.6 Hz, 1H), 7.54 (q, *J* = 2.3 Hz, 2H), 7.00 (s, 1H), 6.49 (d, *J* = 2.2 Hz, 1H), 6.36 (s, 1H), 4.52 (m, 4H), 4.43 (d, *J* = 7.7 Hz, 1H), 4.24 (h, *J* = 7.1 Hz, 1H), 3.86 (d, *J* = 7.5 Hz, 6H), 3.00 (q, *J* = 6.6 Hz, 2H), 2.88 (td, *J* = 12.9, 2.7 Hz, 2H), 2.65 (d, *J* = 4.6 Hz, 3H), 2.58 (p, *J* = 7.4 Hz, 1H), 2.41 – 2.32 (m, 1H), 2.18 (dtd, *J* = 12.3, 7.8, 4.2 Hz, 1H), 2.12 – 1.99 (m, 4H), 1.87 (dt, *J* = 13.9, 7.4 Hz, 1H), 1.84 – 1.65 (m, 7H), 1.46 (dqd, *J* = 16.3, 10.1, 4.2 Hz, 5H), 1.36 (t, *J* = 6.9 Hz, 2H), 1.23 (d, *J* = 2.9 Hz, 8H), 0.81 (t, *J* = 7.4 Hz, 6H). **LC-MS:** m/z 955.51 [M+1]$^+$.

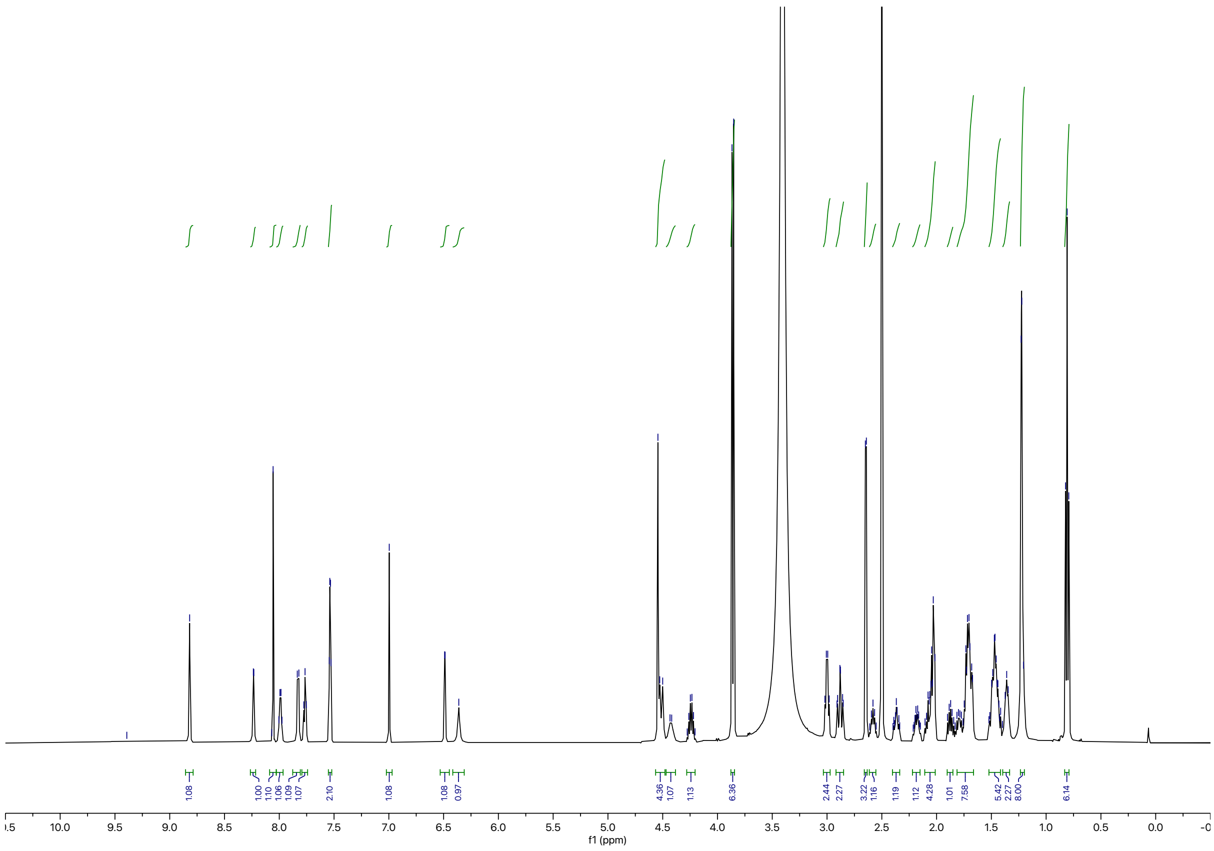

RIGIDFIED DERIVATIVES OF CDK-TCIP1

Synthesis of **RCS-03-207**

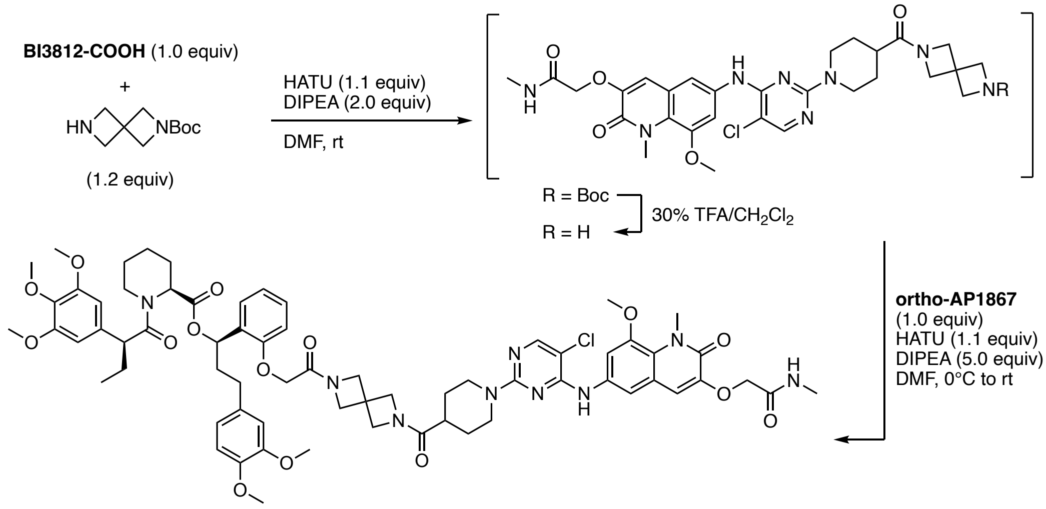


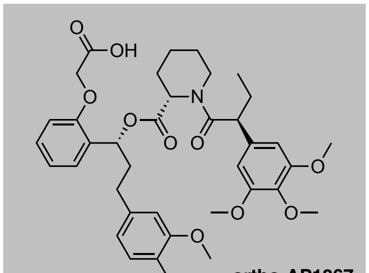

**ortho-AP1867**

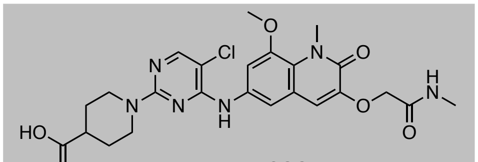

**BI3812-COOH**

To a solution of **BI3812-COOH** (0.015 g, 0.028 mmol, 1.0 equiv) and HATU (0.012 g, 0.031 mmol, 1.1 equiv) in DMF (0.2 mL) was added DIPEA (9.6 μL, 0.056 mmol, 2.0 equiv) and *tert*-butyl 2,6-diazaspiro[3.3]heptane-2-carboxylate (6.7 mg, 0.034 mmol, 1.2 equiv). TLC analysis (10% MeOH in $CH_2Cl_2$) indicated full and clean conversion of starting material after 30 min. The reaction mixture was diluted with EtOAc and the aqueous layer was extracted. The combined organic extracts were washed with brine, dried over $Na_2SO_4$ and concentrated under reduced pressure. The residue was taken up in $CH_2Cl_2$ (0.2 mL), TFA (0.06 mL) was added, and the yellow solution was stirred at 35 °C for 1 h, at which point UPLC-MS analysis indicated clean removal of the Boc protecting group. The mixture was concentrated under reduced pressure and dried under high vacuum to remove excess TFA. Then, a solution of the residue in DMF (0.1 mL) was slowly added to a solution of **ortho-AP1867**[11] (0.019 g, 0.028 mmol, 1.0 equiv), HATU (0.012 g, 0.031 mmol, 1.1 equiv), and DIPEA (0.024 mL, 0.14 mmol, 5.0 equiv) in DMF (0.2 m) at 0 °C. The mixture was allowed to warm to ambient temperature and after stirring for 30 min was directly purified by reverse-phase HPLC (0–40% MeOH in water) to afford **RCS-03-207** as off-white solid upon lyophilization (0.011 g, 30% yield).

**$^1$H NMR** (500 MHz, DMSO-d$_6$) δ = 9.04 (s, 1H), 8.10 (d, *J* = 1.4 Hz, 1H), 7.94 (q, *J* = 4.8 Hz, 1H), 7.57 – 7.46 (m, 2H), 7.18 (td, *J* = 7.8, 2.1 Hz, 1H), 7.00 (q, *J* = 2.9 Hz, 1H), 6.90 – 6.72 (m, 5H), 6.67 – 6.58 (m,

1H), 6.56 (s, 1H), 6.49 (s, 1H), 5.95 (ddd, *J* = 25.1, 8.6, 4.1 Hz, 1H), 5.34 (s, 1H), 4.60 – 4.52 (m, 4H), 4.44 (d, *J* = 13.5 Hz, 2H), 4.37 – 4.23 (m, 4H), 4.16 – 3.92 (m, 8H, assigned by HSQC), 3.89 – 3.85 (m, 6H), 3.70 (dt, *J* = 4.3, 2.6 Hz, 6H), 3.55 (s, 3H), 3.53 (d, *J* = 3.3 Hz, 3H), 3.46 (s, 3H), 2.96 (q, *J* = 11.3 Hz, 2H), 2.90 – 2.80 (m, 1H), 2.64 (d, *J* = 4.7 Hz, 3H), 2.57 (d, *J* = 14.7 Hz, 1H), 2.44 – 2.36 (m, 1H), 2.16 (d, *J* = 12.6 Hz, 1H), 1.99 – 1.84 (m, 3H), 1.70 – 1.35 (m, 10H), 0.81 (t, *J* = 7.3 Hz, 3H). **LC-MS**: m/z 644.05 $[M+1]^{2+}$.

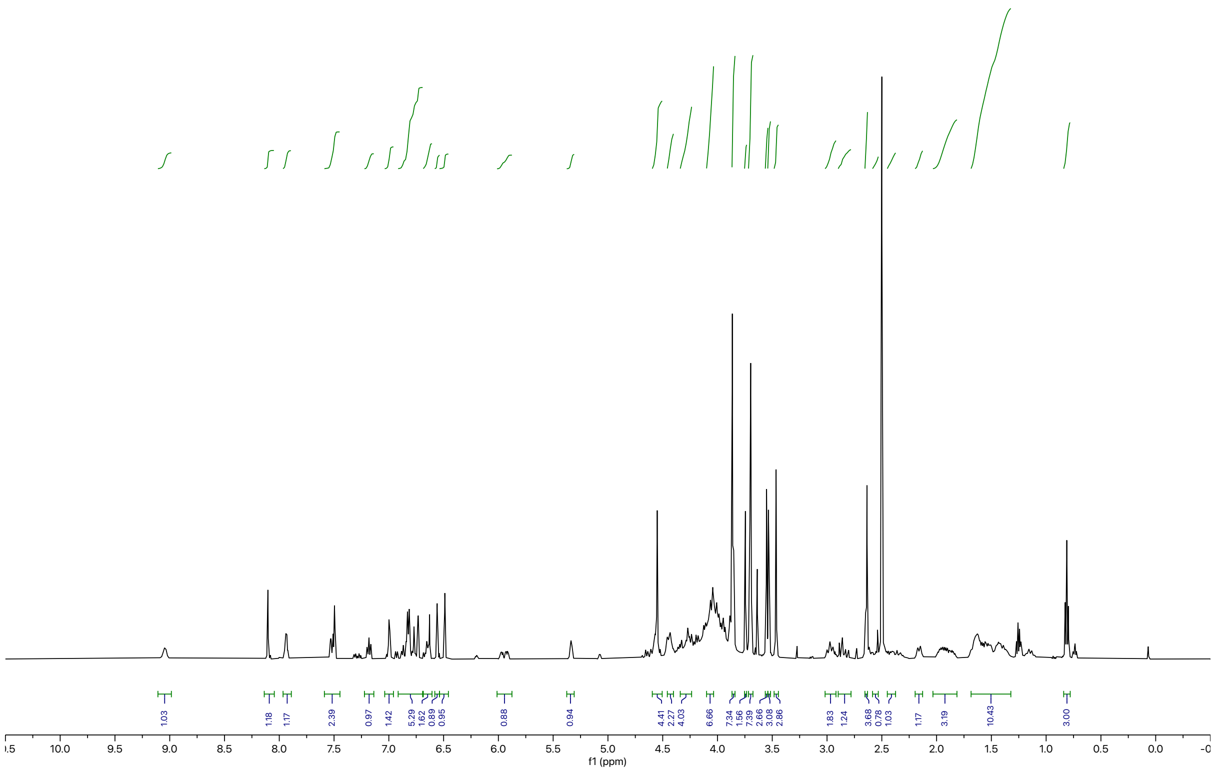

Synthesis of **BAK-04-205**

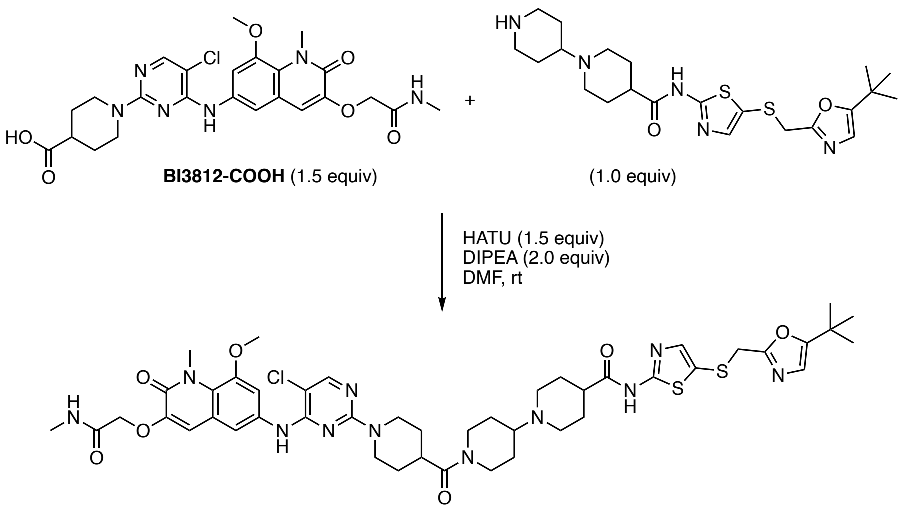


To a solution of **BI23812-COOH** (0.017 g, 0.032 mmol, 1.5 equiv), HATU (0.012 g, 0.032 mmol, 1.5 equiv), and DIPEA (7.5 mL, 0.043 mmol, 2.0 equiv) in DMF (0.2 mL) was added a solution of N-(5-(((5-(*tert*-butyl)oxazol-2-yl)methyl)thio)thiazol-2-yl)-[1,4'-bipiperidine]-4-carboxamide[12] (0.010 g, 0.022 mmol, 1.0 equiv) in DMF (0.2 mL). The resulting brownish solution was stirred at ambient temperature until LC-MS analysis indicated full conversion of starting materials (30 min). The reaction mixture was directly purified by reverse-phase HPLC (15–85% MeOH in water) to afford **BAK-04-205** as off-white solid upon lyophilization (6.0 mg, 30% yield).

**$^1$H NMR** (500 MHz, DMSO-$d_6$) δ = 12.45 (s, 1H), 9.30 (s, 1H), 8.85 (s, 1H), 8.07 (s, 1H), 7.98 (d, *J* = 6.3 Hz, 1H), 7.59 (d, *J* = 2.3 Hz, 1H), 7.51 (d, *J* = 2.3 Hz, 1H), 7.41 (s, 1H), 7.00 (s, 1H), 6.72 (s, 1H), 4.54 (d, *J* = 15.7 Hz, 5H), 4.20 (s, 1H), 4.06 (s, 2H), 3.87 (d, *J* = 7.3 Hz, 6H), 3.58 (m, 1H, assigned by HSQC), 3.46 (m, 2H, assigned by HSQC), 3.13 – 2.93 (m, 7H), 2.77 (t, *J* = 12.1 Hz, 1H), 2.65 (d, *J* = 4.6 Hz, 3H), 2.16 – 2.02 (m, 4H), 1.86 (q, *J* = 13.1 Hz, 2H), 1.67 (d, *J* = 12.7 Hz, 2H), 1.56 – 1.38 (m, 4H), 1.18 (s, 9H). **LC-MS**: m/z 976.46 [M+1]$^+$.

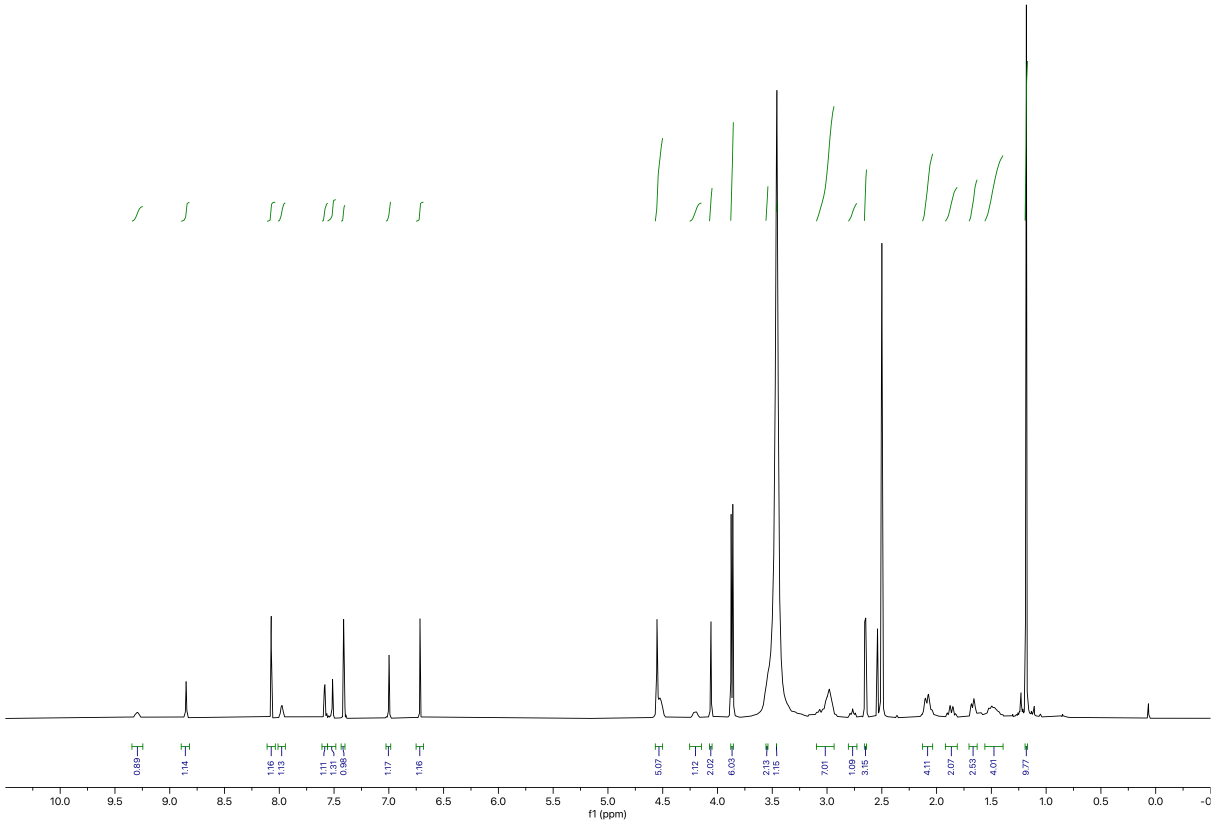
0.89
1.14
1.16
1.13
1.11
1.31
0.98
1.17
1.16
5.07
1.12
2.02
6.03
2.13
1.15
7.01
1.09
3.15
4.11
2.07
2.53
4.01
9.77
10.0
9.5
9.0
8.5
8.0
7.5
7.0
6.5
6.0
5.5
5.0
4.5
4.0
3.5
3.0
2.5
2.0
1.5
1.0
0.5
0.0
f1 (ppm)

Synthesis of **BAK-04-230**

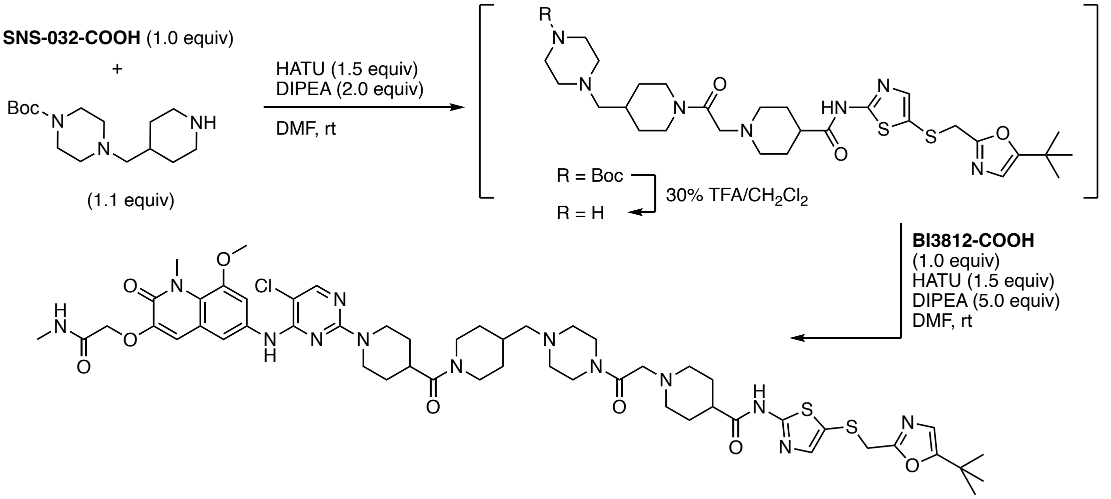


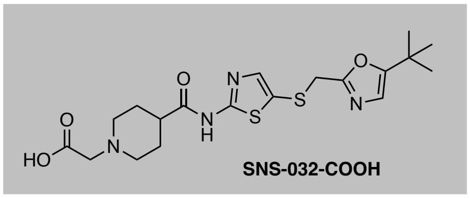


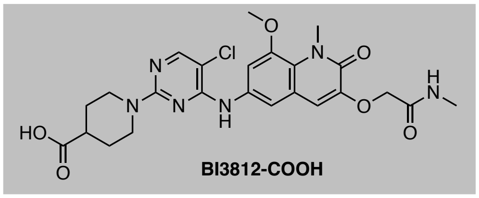


To a solution of **SNS-032-COOH** (4.8 mg, 0.011 mmol, 1.0 equiv) and HATU (6.3 mg, 0.017 mmol, 1.5 equiv) in DMF (0.2 mL) was added DIPEA (3.8 µL, 0.022 mmol, 2.0 equiv) and *tert*-butyl 4-(piperidin-4-ylmethyl)piperazine-1-carboxylate (3.4 mg, 0.011 mmol, 1.1 equiv). TLC analysis (10% MeOH in $CH_2Cl_2$) indicated full and clean conversion of starting material after 15 min. The reaction mixture was diluted with EtOAc and the aqueous layer was extracted. The combined organic extracts were washed with brine, dried over $Na_2SO_4$ and concentrated under reduced pressure. The waxy residue was taken up in $CH_2Cl_2$ (0.2 mL), TFA (0.06 mL) was added, and the yellow solution was stirred at 40 °C for 1 h, at which point LC-MS analysis indicated full removal of the Boc protecting group. The mixture was concentrated under reduced pressure and dried under high vacuum to remove excess TFA. Then, a solution of the residue in DMF (0.1 mL) was slowly added to a solution of **BI3812-COOH** (6.0 mg, 0.011 mmol, 1.0 equiv), HATU (6.5 mg, 0.017 mmol, 1.5 equiv), and DIPEA (9.9 µL, 0.057 mmol, 5.0 equiv) in DMF (0.2 mL) at 0 °C. The mixture was allowed to warm to ambient temperature and after stirring for 30 min was directly purified by reverse-phase HPLC (15–85% MeOH in water) to afford **BAK-04-205** as off-white solid upon lyophilization (4.0 mg, 30% yield).

**$^1$H NMR** (500 MHz, DMSO-$d_6$) δ = 12.42 (s, 1H), 9.75 (s, 1H), 8.94 (s, 1H), 8.10 (s, 1H), 7.96 (d, *J* = 4.9 Hz, 1H), 7.58 (d, *J* = 2.3 Hz, 1H), 7.52 (d, *J* = 2.3 Hz, 1H), 7.42 (s, 1H), 7.01 (s, 1H), 6.73 (s, 1H), 4.56 (s, 2H), 4.51 (d, *J* = 12.5 Hz, 2H), 4.39 (d, *J* = 13.3 Hz, 5H), 4.10 – 4.03 (m, 4H), 3.88 (s, 3H), 3.87 (s, 3H), 3.64 – 3.48 (m, 4H), 3.39 – 3.25 (m, 2H), 3.12 – 2.96 (m, 11H), 2.78 (s, 1H), 2.66 (d, *J* = 4.6 Hz, 3H), 2.61 – 2.56 (m, 1H), 2.16 – 1.91 (m, 5H), 1.88 – 1.74 (m, 2H), 1.66 (s, 2H), 1.59 – 1.43 (m, 2H), 1.19 (s, 9H).

**LC-MS**: m/z 1116.61 $[M+1]^+$.

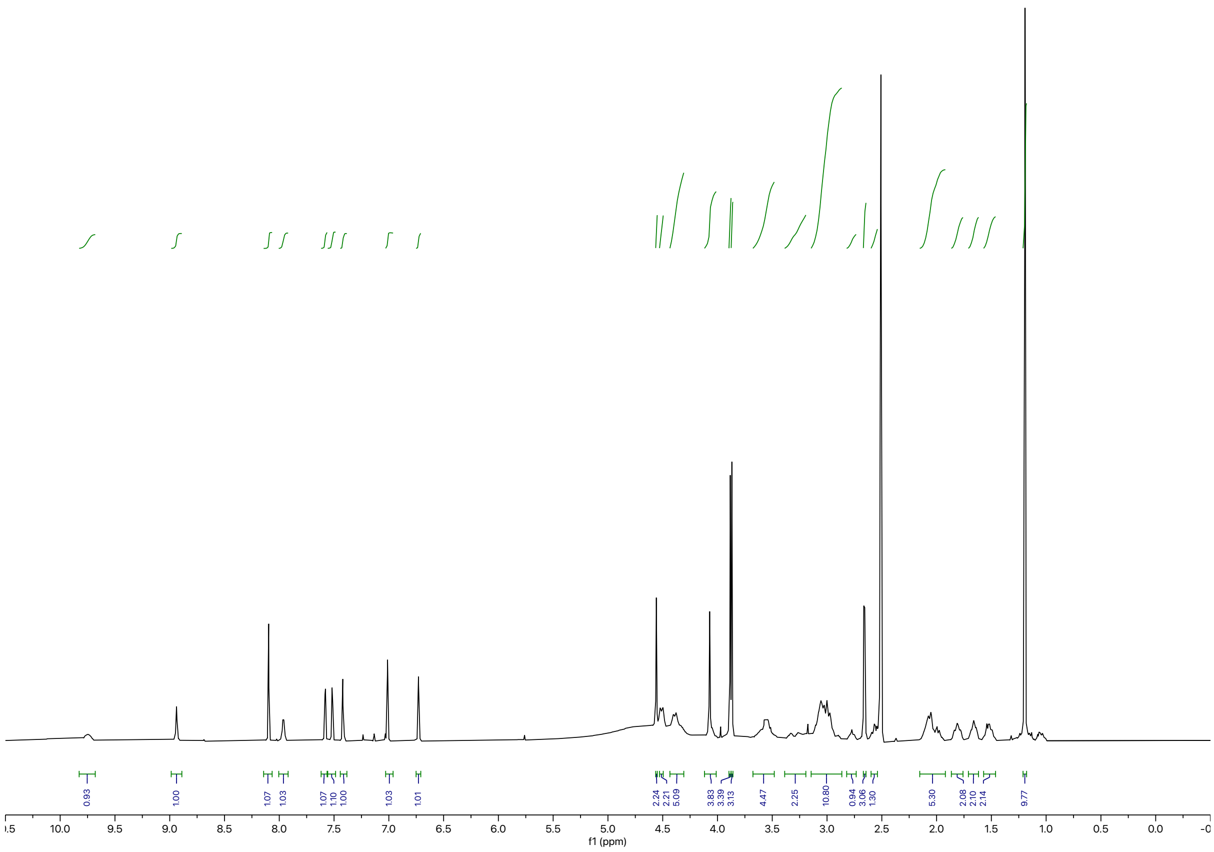

0.93
1.00
1.07
1.03
1.07
1.10
1.00
1.03
1.01
2.24
2.21
5.09
3.83
3.39
3.13
4.47
2.25
10.80
0.94
3.06
1.30
5.30
2.08
2.10
2.14
9.77
10.0 9.5 9.0 8.5 8.0 7.5 7.0 6.5 6.0 5.5 5.0 4.5 4.0 3.5 3.0 2.5 2.0 1.5 1.0 0.5 0.0
f1 (ppm)

Synthesis of **SNS-032-COOH**

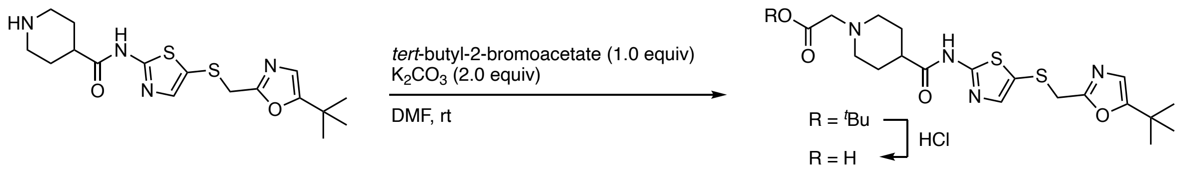


To a solution of **SNS-032** (0.10 g, 0.26 mmol, 1.0 equiv) in DMF (1.3 mL) was added tert-butyl-2-bromoacetate (0.039 mL, 0.26 mmol, 1.0 equiv) and $K_2CO_3$ (0.073 g, 0.53 mmol, 2.0 equiv) and the mixture was stirred at ambient temperature. TLC (10% MeOH in $CH_2Cl_2$) and LC-MS analysis indicated full and clean conversion after 14 h. Water was added and the resulting precipitate was filtered off and dried under high vacuum to afford the intermediate (0.13 g, quantitative yield), which was taken up in 1,4-dioxane (2.0 mL). HCl (4 M in 1,4-dioxane, 0.33 mL, 1.3 mmol, 5.0 equiv) was added and the suspension was stirred at 35 °C for 2 h. All volatiles were removed under reduced pressure to afford **SNS-032-COOH** as off-white solid (0.12 g, quantitative yield).

*$^t$Bu ester intermediate*

**$^1$H NMR** (500 MHz, $CDCl_3$) δ = 7.33 (s, 1H), 6.66 (s, 1H), 4.03 (s, 2H), 3.82 (s, 2H), 3.68 (s, 1H), 3.35 (s, 2H), 2.89 (s, 1H), 2.30 (s, 4H), 1.48 (s, 9H), 1.29 (s, 9H). **LC-MS**: m/z 495.29 $[M+1]^+$.

*SNS-032-COOH*

**$^1$H NMR** (500 MHz, MeOD) δ = 7.33 (s, 1H), 6.67 (s, 1H), 4.11 (s, 2H), 3.99 (s, 2H), 3.75 (s, 2H), 3.17 (p, *J* = 1.6 Hz, 1H), 2.82 (s, 2H), 2.23 – 2.06 (m, 4H), 1.25 (s, 9H). **LC-MS**: m/z 439.27 $[M+1]^+$.

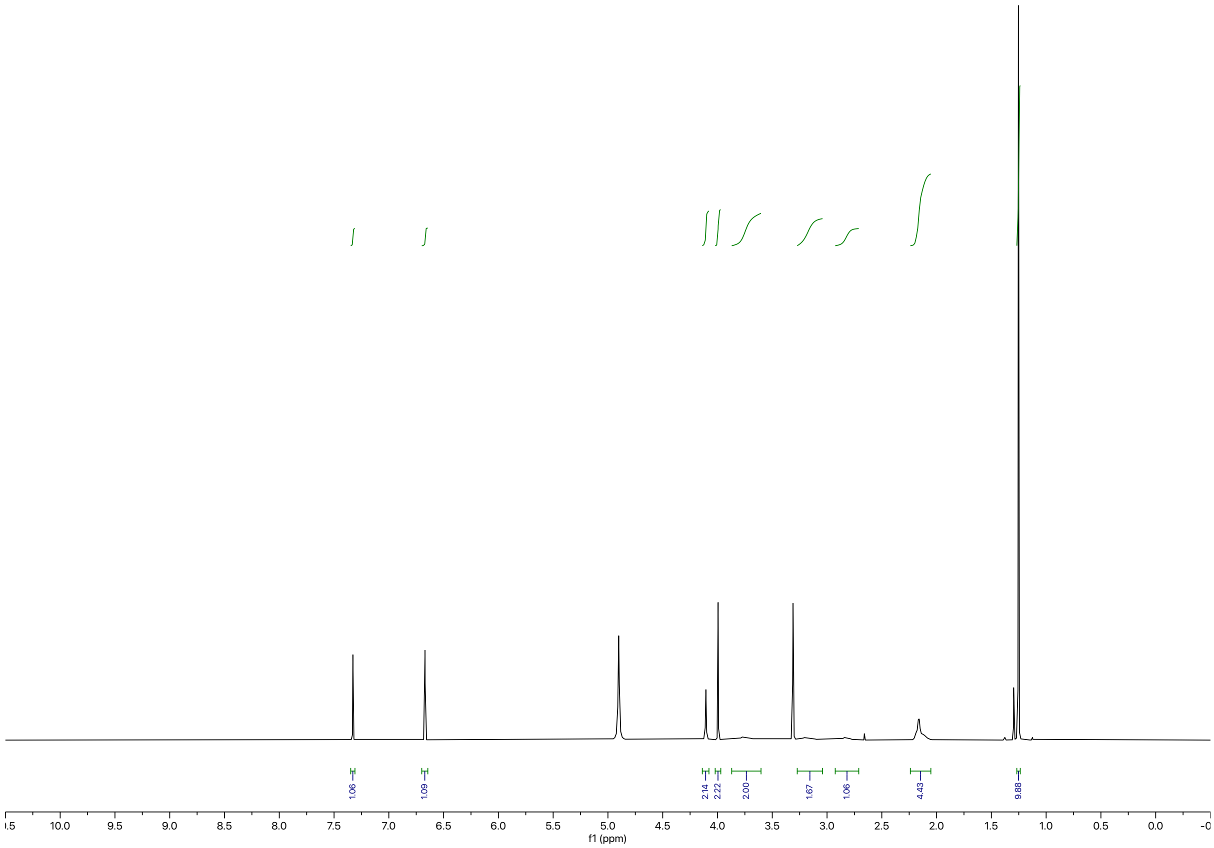
1.06
1.09
2.14
2.22
2.00
1.67
1.06
4.43
9.88
10.0
9.5
9.0
8.5
8.0
7.5
7.0
6.5
6.0
5.5
5.0
4.5
4.0
3.5
3.0
2.5
2.0
1.5
1.0
0.5
0.0
f1 (ppm)

Synthesis of **BAK-04-238**

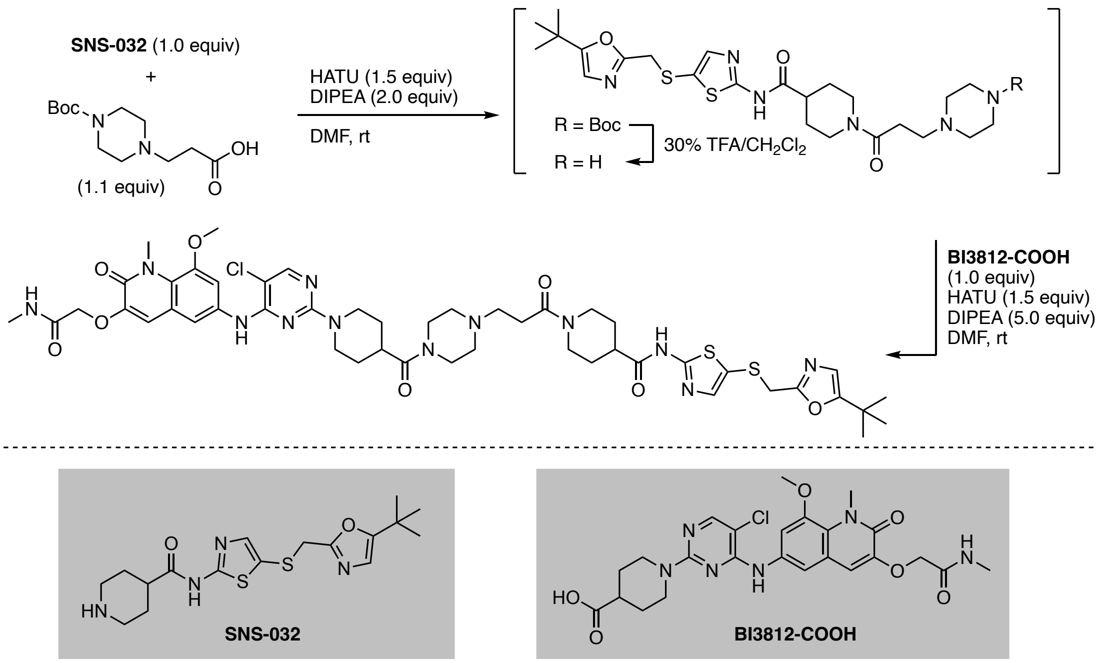


To a solution of 3-(4-(*tert*-butoxycarbonyl)piperazin-1-yl)propanoic acid (4.3 mg, 0.017 mmol, 1.1 equiv), HATU (8.6 mg, 0.023 mmol, 1.5 equiv), and DIPEA (5.2 μL, 0.030 mmol, 2.0 equiv) in DMF (0.2 mL) as added **SNS-032** (5.7 mg, 0.015 mmol, 1.0 equiv) in DMF (0.2 mL). The resulting solution was stirred at ambient temperature until TLC analysis (10% MeOH in $CH_2Cl_2$) indicated full and clean conversion of starting material. The reaction mixture was diluted with EtOAc and the aqueous layer was extracted thoroughly with EtOAc. The combined organic extracts were washed with brine, dried over $Na_2SO_4$ and concentrated under reduced pressure. The waxy residue was taken up in $CH_2Cl_2$ (0.2 mL), TFA (0.06 mL) was added, and the yellow solution was stirred at 40 °C for 1 h, at which point LC-MS analysis indicated full removal of the Boc protecting group. The mixture was concentrated under reduced pressure and dried under high vacuum to remove excess TFA. Then, a solution of the residue in DMF (0.1 mL) was slowly added to a solution of **BI3812-COOH** (8.0 mg, 0.015 mmol, 1.0 equiv), HATU (8.3 mg, 0.022 mmol, 1.5 equiv), and DIPEA (0.013 mL, 0.072 mmol, 5.0 equiv) in DMF (0.2 mL) at 0 °C. The mixture was allowed to warm to ambient temperature and then was directly purified by reverse-phase HPLC (15–85% MeOH in water) to afford **BAK-04-238** as off-white solid upon lyophilization (4.5 mg, 30% yield).

**$^1$H NMR** (500 MHz, DMSO-d$_6$) δ = 12.33 (s, 1H), 9.64 (s, 1H), 8.85 (s, 1H), 8.08 (s, 1H), 7.95 (d, *J* = 4.7 Hz, 1H), 7.60 (d, *J* = 2.3 Hz, 1H), 7.51 (d, *J* = 2.3 Hz, 1H), 7.39 (s, 1H), 6.98 (s, 1H), 6.71 (s, 1H), 4.54 (d, *J* = 21.1 Hz, 4H), 4.46 – 4.31 (m, 2H), 4.29 – 4.18 (m, 1H), 4.05 (s, 2H), 3.96 – 3.78 (m, 7H), 3.51-3.46 (m, 1H, assigned by HSQC), 3.36 (s, 2H), 3.08 (t, *J* = 12.7 Hz, 2H), 2.98 (t, *J* = 11.9 Hz, 5H), 2.87 (dq, *J* = 17.2, 8.2 Hz, 2H), 2.80 – 2.73 (m, 1H), 2.71 – 2.66 (m, 1H), 2.65 (d, *J* = 4.7 Hz, 3H), 1.90 – 1.80 (m, 2H), 1.75 – 1.57 (m, 4H), 1.56 – 1.38 (m, 4H), 1.18 (s, 9H). **LC-MS**: m/z 1033.51 $[M+1]^+$.

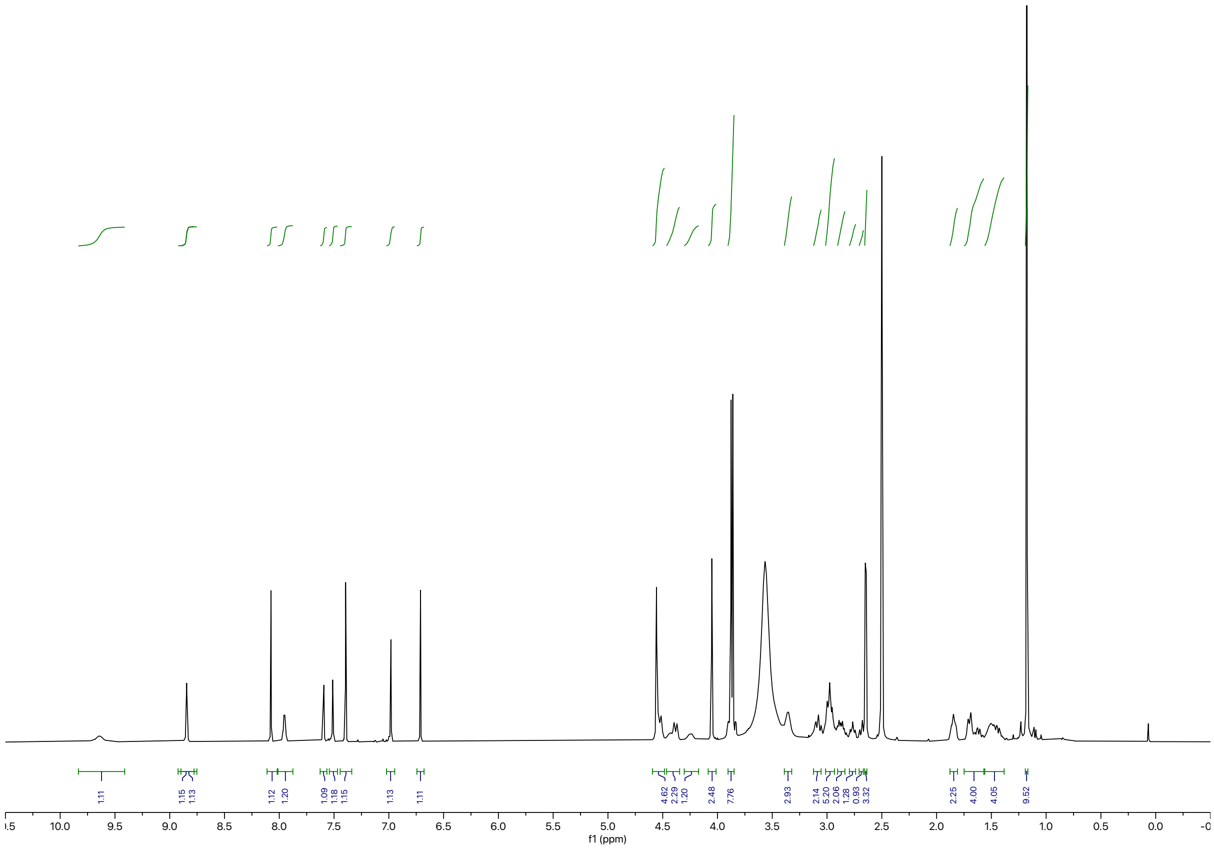
1.11
1.15
1.13
1.12
1.20
1.09
1.18
1.15
1.13
1.11
4.62
2.29
1.20
2.48
7.76
2.93
2.14
5.20
2.06
1.28
0.93
3.32
2.25
4.00
4.05
9.52
10.0
9.5
9.0
8.5
8.0
7.5
7.0
6.5
6.0
5.5
5.0
4.5
4.0
3.5
3.0
2.5
2.0
1.5
1.0
0.5
0.0
f1 (ppm)

Synthesis of **BAK-04-248**

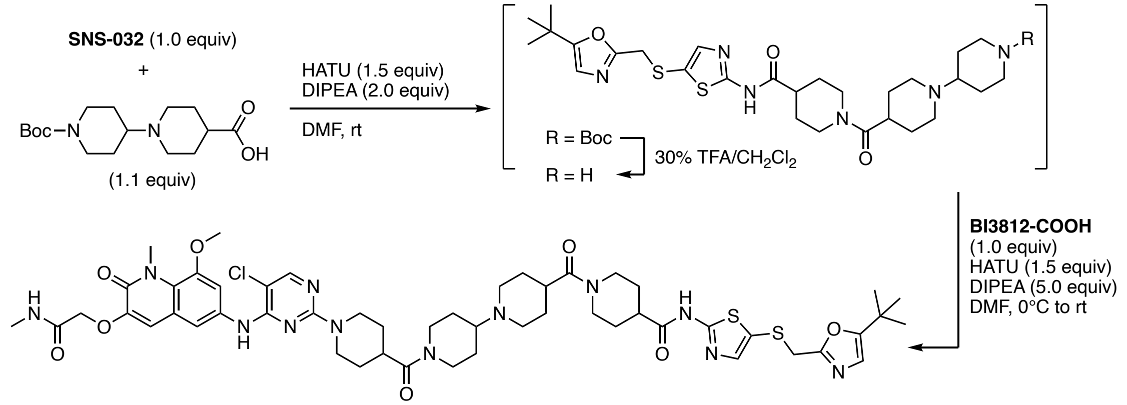


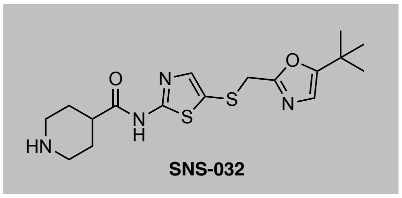


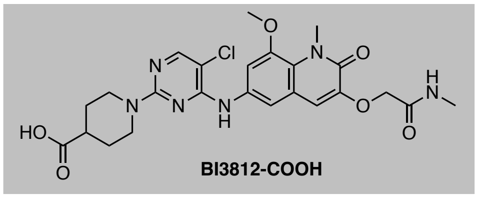


To a solution of 1'-(*tert*-butoxycarbonyl)-[1,4'-bipiperidine]-4-carboxylic acid (5.2 mg, 0.017 mmol, 1.1 equiv), HATU (8.6 mg, 0.023 mmol, 1.5 equiv), and DIPEA (5.2 μL, 0.030 mmol, 2.0 equiv) in DMF (0.2 mL) as added **SNS-032** (5.7 mg, 0.015 mmol, 1.0 equiv) in DMF (0.2 mL). The resulting solution was stirred at ambient temperature until TLC analysis (10% MeOH in $CH_2Cl_2$) indicated full and clean conversion of starting material. The reaction mixture was diluted with EtOAc and the aqueous layer was extracted thoroughly with EtOAc. The combined organic extracts were washed with brine, dried over $Na_2SO_4$ and concentrated under reduced pressure. The resulting residue was taken up in $CH_2Cl_2$ (0.2 mL), TFA (0.06 mL) was added, and the yellow solution was stirred at 35 °C for 3 h, at which point LC-MS analysis indicated full removal of the Boc protecting group. The mixture was concentrated under reduced pressure and dried under high vacuum to remove excess TFA. Then, a solution of the residue in DMF (0.1 mL) was slowly added to a solution of **BI3812-COOH** (7.0 mg, 0.015 mmol, 1.0 equiv), HATU (8.4 mg, 0.022 mmol, 1.5 equiv), and DIPEA (0.013 mL, 0.074 mmol, 5.0 equiv) in DMF (0.2 mL) at 0 °C. The mixture was allowed to warm to ambient temperature and then was directly purified by reverse-phase HPLC (15–85% MeOH in water) to afford **BAK-04-248** as white solid upon lyophilization (3.0 mg, 20% yield).

**$^1$H NMR** (500 MHz, DMSO-d6) δ = 12.32 (s, 1H), 9.22 (s, 1H), 8.83 (s, 1H), 8.07 (s, 1H), 8.01 (t, *J* = 4.6 Hz, 1H), 7.59 (d, *J* = 2.3 Hz, 1H), 7.51 (d, *J* = 2.3 Hz, 1H), 7.39 (s, 1H), 7.00 (s, 1H), 6.71 (s, 1H), 4.58 – 4.47 (m, 5H), 4.38 (d, *J* = 13.1 Hz, 1H), 4.25 – 4.18 (m, 1H), 4.07 – 3.99 (m, 3H), 3.87 (s, 3H), 3.86 (s, 3H), 3.51-3.43 (m, 2H, assigned by HSQC), 3.47-3.41 (m, 1H, assigned by HSQC), 3.13 – 3.03 (m, 5H), 3.02 – 2.90 (m, 4H), 2.79 – 2.72 (m, 1H), 2.65 (d, *J* = 4.6 Hz, 3H), 2.63 – 2.59 (m, 1H), 2.09 – 1.97 (m, 2H), 1.89 – 1.79 (m, 6H), 1.67 (d, *J* = 12.5 Hz, 2H), 1.60 – 1.37 (m, 6H), 1.17 (s, 9H). **LC-MS**: m/z 1087.55 $[M+1]^+$.

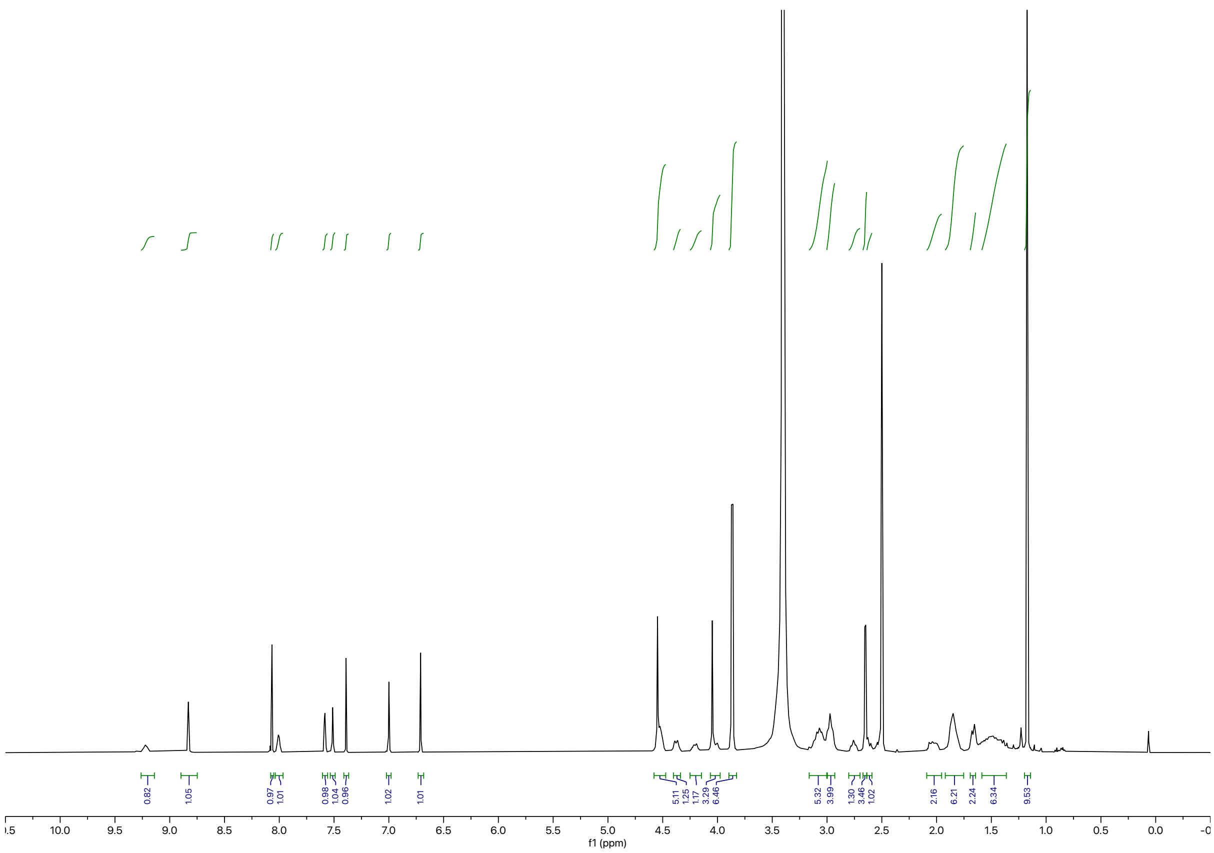

Synthesis of **BAK-04-249**

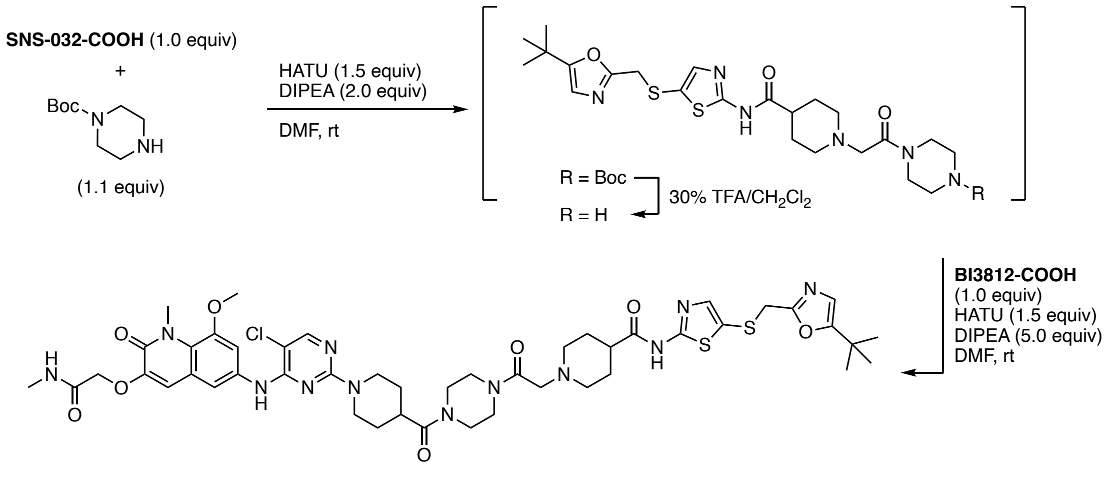


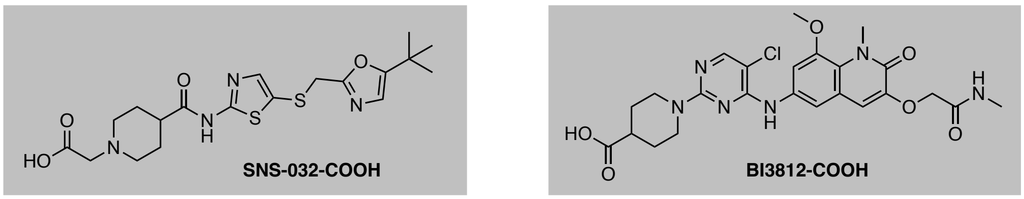


To a solution of **SNS-032-COOH** (4.4 mg, 0.010 mmol, 1.0 equiv), HATU (5.7 mg, 0.015 mmol, 1.5 equiv), and DIPEA (3.5 µL, 0.020 mmol, 2.0 equiv) in DMF (0.2 mL) was added 1'-(*tert*-butoxycarbonyl)-[1,4'-bipiperidine]-4-carboxylic acid (2.0 mg, 0.011 mmol, 1.1 equiv) in DMF (0.2 mL). The resulting solution was stirred at ambient temperature until LC-MS analysis indicated full and clean conversion of starting material. The reaction mixture was diluted with EtOAc and the aqueous layer was extracted thoroughly with EtOAc. The combined organic extracts were washed with brine, dried over $Na_2SO_4$ and concentrated under reduced pressure. The resulting residue was taken up in $CH_2Cl_2$ (0.2 mL), TFA (0.06 mL) was added, and the yellow solution was stirred at 35 °C for 3 h, at which point LC-MS analysis indicated full removal of the Boc protecting group. The mixture was concentrated under reduced pressure and dried under high vacuum to remove excess TFA. Then, a solution of the residue in DMF (0.1 mL) was slowly added to a solution of **BI3812-COOH** (5.0 mg, 0.010 mmol, 1.0 equiv), HATU (5.6 mg, 0.010 mmol, 1.5 equiv), and DIPEA (8.6 µL, 0.049 mmol, 5.0 equiv) in DMF (0.2 mL) at 0 °C. The mixture was allowed to warm to ambient temperature and then was directly purified by reverse-phase HPLC (15–85% MeOH in water) to afford **BAK-04-249** as white solid upon lyophilization (3.0 mg, 30% yield).

**$^1$H NMR** (500 MHz, DMSO-$d_6$) δ = 12.40 (s, 1H), 9.64 (s, 1H), 8.84 (s, 1H), 8.07 (s, 1H), 7.96 (s, 1H), 7.58 (s, 1H), 7.51 (s, 1H), 7.41 (s, 1H), 6.99 (s, 1H), 6.72 (s, 1H), 4.53 (d, *J* = 14.4 Hz, 4H), 4.36 (d, *J* = 32.5 Hz, 2H), 4.06 (s, 2H), 3.87 (s, 3H), 3.86 (s, 3H), 3.67-3.46 (m, 8H, assigned by HSQC), 3.39 – 3.27 (m, 2H), 3.07 – 2.93 (m, 5H), 2.79 – 2.71 (m, 1H), 2.64 (d, *J* = 4.6 Hz, 3H), 2.09 – 1.93 (m, 4H), 1.70 (d, *J* = 12.9 Hz, 2H), 1.57 – 1.47 (m, 2H), 1.18 (s, 9H). **LC-MS**: m/z 1019.43 $[M+1]^+$.

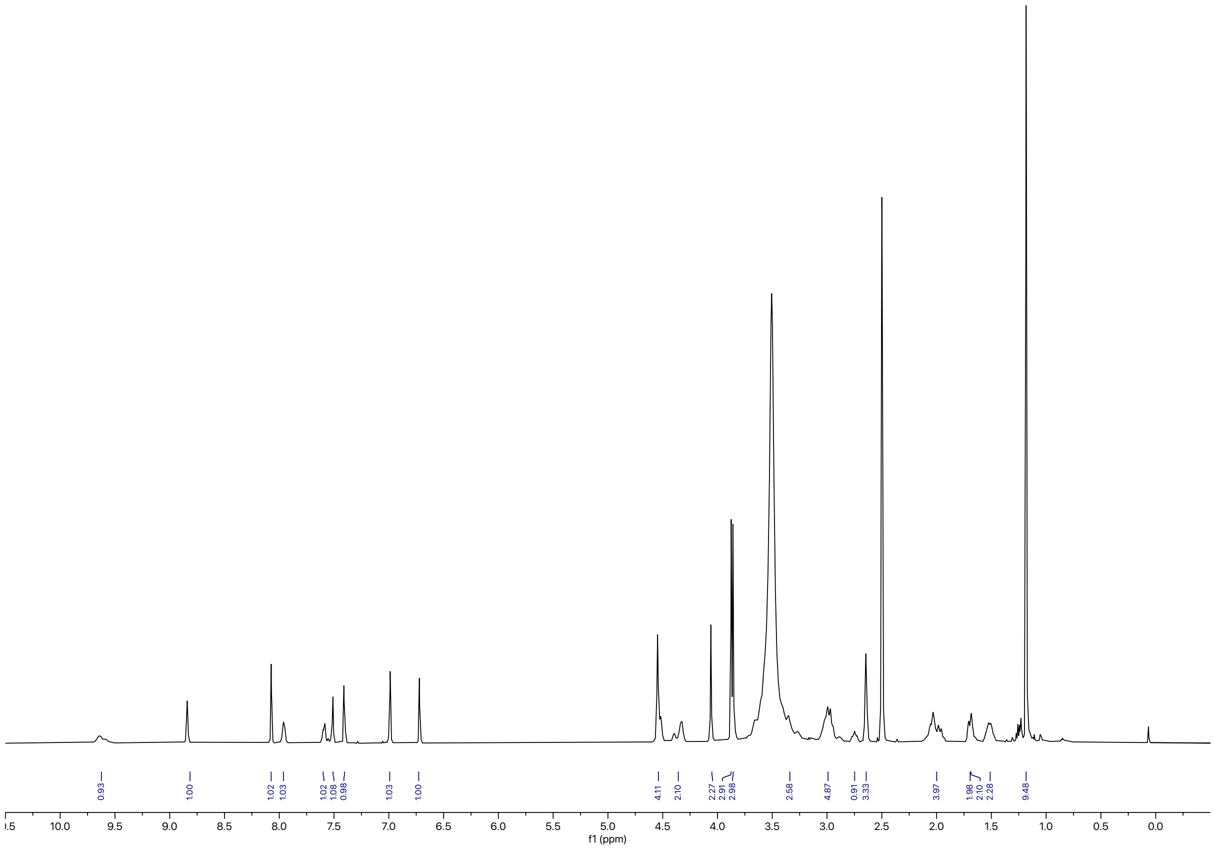
0.93
1.00
1.02
1.03
1.02
1.08
0.98
1.03
1.00
4.11
2.10
2.27
2.91
2.98
2.58
4.87
0.91
3.33
3.97
1.98
2.10
2.28
9.48
10.0
9.5
9.0
8.5
8.0
7.5
7.0
6.5
6.0
5.5
5.0
4.5
4.0
3.5
3.0
2.5
2.0
1.5
1.0
0.5
0.0
f1 (ppm)

Synthesis of **BAK-04-253**

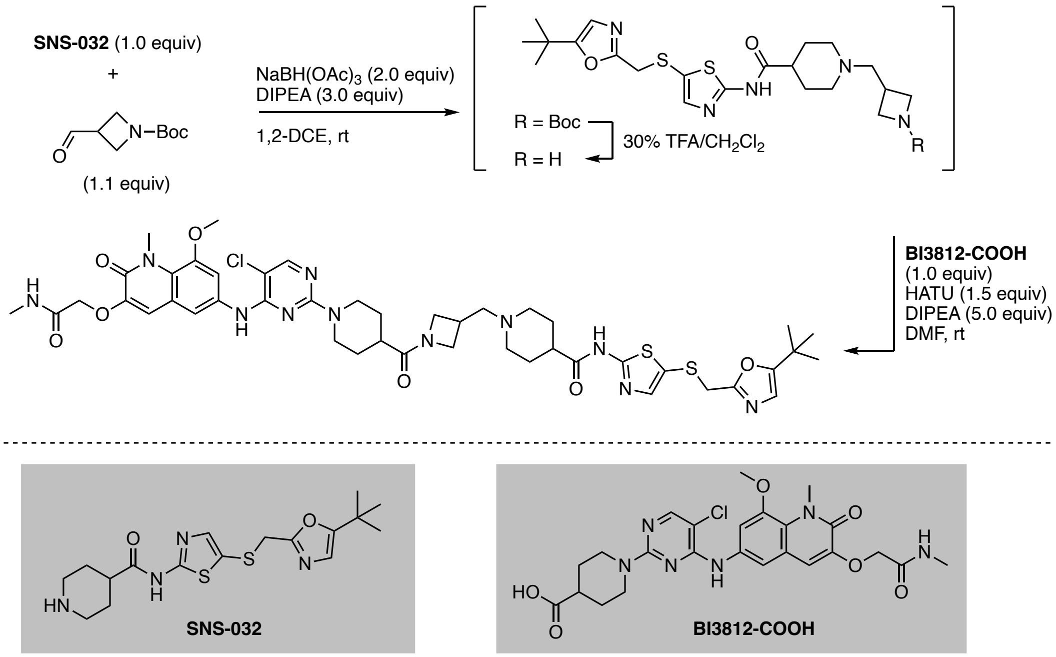


To a solution of **SNS-032** (3.8 mg, 0.011 mmol, 1.0 equiv) and *tert*-butyl 3-formylazetidine-1-carboxylate[13] (2.2 mg, 0.012 mmol, 1.1 equiv) in 1,2-DCE (0.2 mL) were added DIPEA (5.7 μL, 0.033 mmol, 3.0˚equiv) and $NaBH(OAc)_3$ (4.7 mg, 0.022 mmol, 2.0 equiv) and the reaction mixture was stirred for 3 h at ambient temperature. The mixture was diluted with water and $CH_2Cl_2$ and the aqueous layer was extracted with $CH_2Cl_2$. The combined organic extracts were washed with brine, dried over $Na_2SO_4$ and concentrated under reduced pressure. The resulting residue was taken up in $CH_2Cl_2$ (0.2 mL), TFA (0.06 mL) was added, and the yellow solution was stirred at 35 °C for 3 h, at which point LC-MS analysis indicated full removal of the Boc protecting group. The mixture was concentrated under reduced pressure and dried under high vacuum to remove excess TFA. Then, a solution of the residue in DMF (0.1 mL) was slowly added to a solution of **BI3812-COOH** (5.9 mg, 0.011 mmol, 1.0 equiv), HATU (6.3 mg, 0.017 mmol, 1.5 equiv), and DIPEA (9.7 μL, 0.056 mmol, 5.0 equiv) in DMF (0.2 mL) at 0 °C. The mixture was allowed to warm to ambient temperature and then was directly purified by reverse-phase HPLC (15–85% MeOH in water) to afford **BAK-04-253** as white solid upon lyophilization (4.0 mg, 37% yield).

**$^1$H NMR** (500 MHz, DMSO-d$_6$) δ = 12.44 (s, 1H), 9.42 (s, 1H), 8.83 (s, 1H), 8.07 (s, 1H), 7.97 (d, *J* = 5.1 Hz, 1H), 7.59 (d, *J* = 2.3 Hz, 1H), 7.51 (d, *J* = 2.3 Hz, 1H), 7.41 (s, 1H), 6.98 (s, 1H), 6.72 (s, 1H), 4.56 (s, 2H), 4.55 – 4.49 (m, 2H), 4.38 (t, *J* = 8.7 Hz, 1H), 4.06 (s, 2H), 4.03 – 3.96 (m, 2H), 3.87 (s, 3H), 3.86 (s, 3H), 3.66 (dd, *J* = 9.9, 5.6 Hz, 1H), 3.54-3.37 (m, 4H, assigned by HSQC), 3.09 – 3.02 (m, 1H), 3.00 – 2.91 (m, 4H), 2.79 – 2.70 (m, 1H), 2.66 (d, *J* = 4.7 Hz, 3H), 2.54-2.45 (m, 1H, assigned by HSQC), 2.05 (d, *J* = 13.7 Hz, 2H), 1.83 (q, *J* = 13.2 Hz, 2H), 1.73 – 1.63 (m, 2H), 1.51 – 1.41 (m, 2H), 1.18 (s, 9H). **LC-MS**: m/z 962.46 [M+1]$^+$.

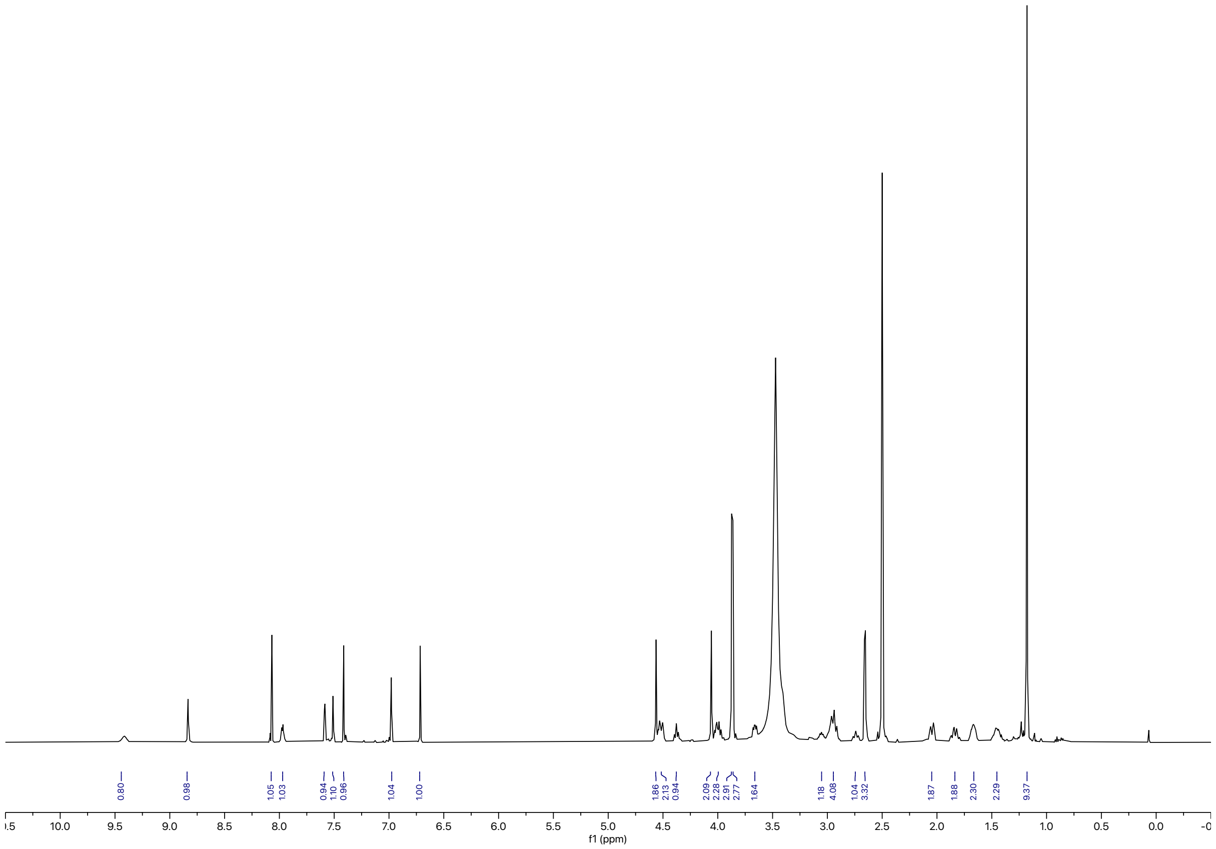
0.80
0.98
1.05
1.03
0.94
1.10
0.96
1.04
1.00
1.86
2.13
0.94
2.09
2.28
2.91
2.77
1.64
1.18
4.08
1.04
3.32
1.87
1.88
2.30
2.29
9.37
10.0
9.5
9.0
8.5
8.0
7.5
7.0
6.5
6.0
5.5
5.0
4.5
4.0
3.5
3.0
2.5
2.0
1.5
1.0
0.5
0.0
f1 (ppm)

Synthesis of **BAK-04-263**

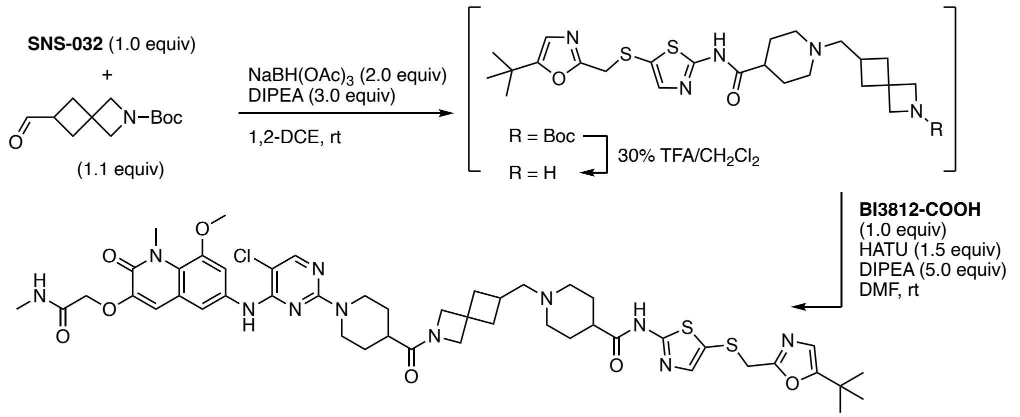


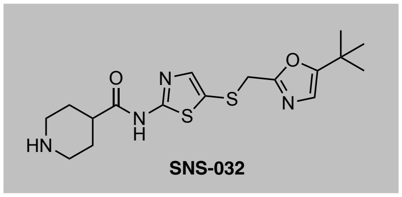


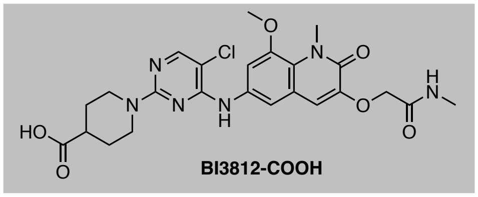


To a solution of **SNS-032** (6.5 mg, 0.017 mmol, 1.0 equiv) and *tert*-butyl 6-formyl-2-azaspiro[3.3]heptane-2-carboxylate[14] (4.6 mg, 0.021 mmol, 1.1 equiv) in 1,2-DCE (0.2 mL) were added DIPEA (8.9 µL, 0.051 mmol, 3.0˚equiv) and $NaBH(OAc)_3$ (7.2 mg, 0.034 mmol, 2.0 equiv) and the reaction mixture was stirred for 3 h at ambient temperature, at which point LC-MS analysis indicated full conversion. The mixture was diluted with water and $CH_2Cl_2$ and the aqueous layer was extracted with $CH_2Cl_2$. The combined organic extracts were washed with brine, dried over $Na_2SO_4$ and concentrated under reduced pressure. The resulting residue was taken up in $CH_2Cl_2$ (0.2 mL), TFA (0.06 mL) was added, and the yellow solution was stirred at 35 °C for 3 h, at which point LC-MS analysis indicated full removal of the Boc protecting group. The mixture was concentrated under reduced pressure and dried under high vacuum to remove excess TFA. Then, a solution of the residue in DMF (0.1 mL) was slowly added to a solution of **BI3812-COOH** (9.0 mg, 0.017 mmol, 1.0 equiv), HATU (9.7 mg, 0.025 mmol, 1.5 equiv), and DIPEA (0.015 mL, 0.085 mmol, 5.0 equiv) in DMF (0.2 mL) at 0 °C. The mixture was allowed to warm to ambient temperature and then was directly purified by reverse-phase HPLC (15–85% MeOH in water) to afford **BAK-04-263** as white solid upon lyophilization (5.2 mg, 31% yield).

**$^1$H NMR** (500 MHz, DMSO-d$_6$) δ = 12.22 (s, 1H), 8.77 (s, 1H), 8.05 (s, 1H), 7.94 (q, *J* = 4.7 Hz, 1H), 7.57 (t, *J* = 1.7 Hz, 1H), 7.52 (d, *J* = 2.3 Hz, 1H), 7.38 (s, 1H), 6.97 (s, 1H), 6.71 (s, 1H), 4.55 (s, 2H), 4.51 (d, *J* = 13.1 Hz, 2H), 4.23 (s, 1H), 4.09 (s, 1H), 4.05 (s, 2H), 3.87 (s, 3H), 3.86 (s, 3H), 3.84 (s, 1H), 3.71 (s, 1H), 2.95 – 2.87 (m, 2H), 2.86 – 2.77 (m, 2H), 2.65 (d, *J* = 4.6 Hz, 3H), 2.48 – 2.42 (m, 2H), 2.39 – 2.18 (m, 5H), 1.94 – 1.69 (m, 6H), 1.68 – 1.53 (m, 4H), 1.42 (q, *J* = 12.4 Hz, 2H), 1.17 (s, 9H). **LC-MS**: m/z 1002.52 $[M+1]^+$.

Synthesis of **BAK-04-232 (CDK-TCIP2)**

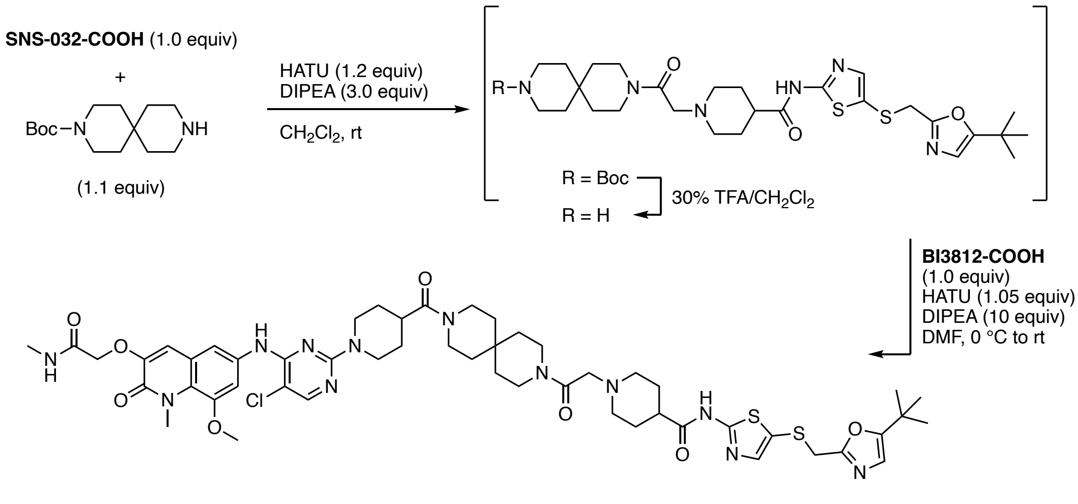


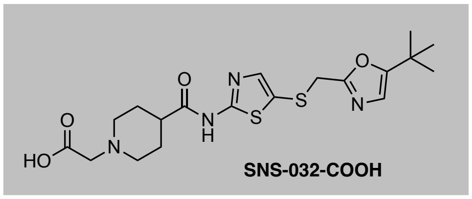


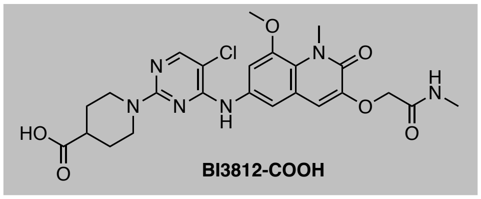


To a solution of **SNS-032-COOH** (0.16 g, 0.37 mmol, 1.0 equiv), HATU (0.17 g, 0.44 mmol, 1.2 equiv), and DIPEA (0.19 mL, 1.1 mmol, 3.0 equiv) in $CH_2Cl_2$ (1.5 mL) was slowly added *tert*-butyl 3,9-diazaspiro[5.5]undecane-9-carboxylate (0.10 g, 0.40 mmol, 1.1 quiv) and the reaction mixture was stirred for 30 min. Water was added and the aqueous layer was extracted with $CH_2Cl_2$ and after concentration of the organic extracts the crude material was purified by flash column chromatography on silica (0-10% MeOH in $CH_2Cl_2$) to afford the intermediate in quantitative yield. The intermediate was taken up in $CH_2Cl_2$ (2.0 mL), TFA (0.6 mL) was added, and the yellow solution was stirred at 35 °C for 1 h, at which point LC-MS analysis indicated full removal of the Boc protecting group. The mixture was concentrated under reduced pressure and dried under high vacuum to remove excess TFA. Then, a solution of the residue in DMF (1.0 mL) was slowly added to a solution of **BI3812-COOH** (0.19 g, 0.37 mmol, 1.0 equiv), HATU (0.15 g, 0.38 mmol, 1.05 equiv), and DIPEA (0.64 mL, 3.7 mmol, 10 equiv) in DMF (1.0 mL) at 0 °C. The mixture was allowed to warm to ambient temperature and then water (10 mL) was added and the precipitate was filtered off, redissolved and the organic layer washed with sat. aq. $NaHCO_3$ and water, dried and concentrated under reduced pressure. Purification by flash column chromatography on silica (0-30% MeOH in CH2Cl2) afforded **BAK-04-232** as yellow solid (0.19 g, 47% yield from SNS-032-COOH).

**$^1$H NMR** (500 MHz, DMSO-$d_6$) δ = 12.23 (s, 1H), 8.77 (s, 1H), 8.06 (s, 1H), 7.93 (d, *J* = 4.9 Hz, 1H), 7.56 (d, *J* = 2.3 Hz, 1H), 7.53 (d, *J* = 2.3 Hz, 1H), 7.38 (s, 1H), 6.99 (s, 1H), 6.71 (s, 1H), 4.56 – 4.47 (m, 4H), 4.05 (s, 2H), 3.87 (s, 3H), 3.86 (s, 3H), 3.54 – 3.48 (m, 4H), 3.47 – 3.40 (m, 4H), 3.11 (s, 2H), 2.94 (q, *J* = 13.8 Hz, 3H), 2.84 (d, *J* = 10.9 Hz, 2H), 2.65 (d, *J* = 4.7 Hz, 3H), 2.44 (t, *J* = 11.5 Hz, 1H), 2.01 (t, *J* = 11.5

Hz, 2H), 1.76 (d, *J* = 12.1 Hz, 2H), 1.69 – 1.55 (m, 4H), 1.53 – 1.45 (m, 6H), 1.38 (s, 4H), 1.17 (s, 9H). **LC-MS**: m/z 1087.60 [M+1]+.

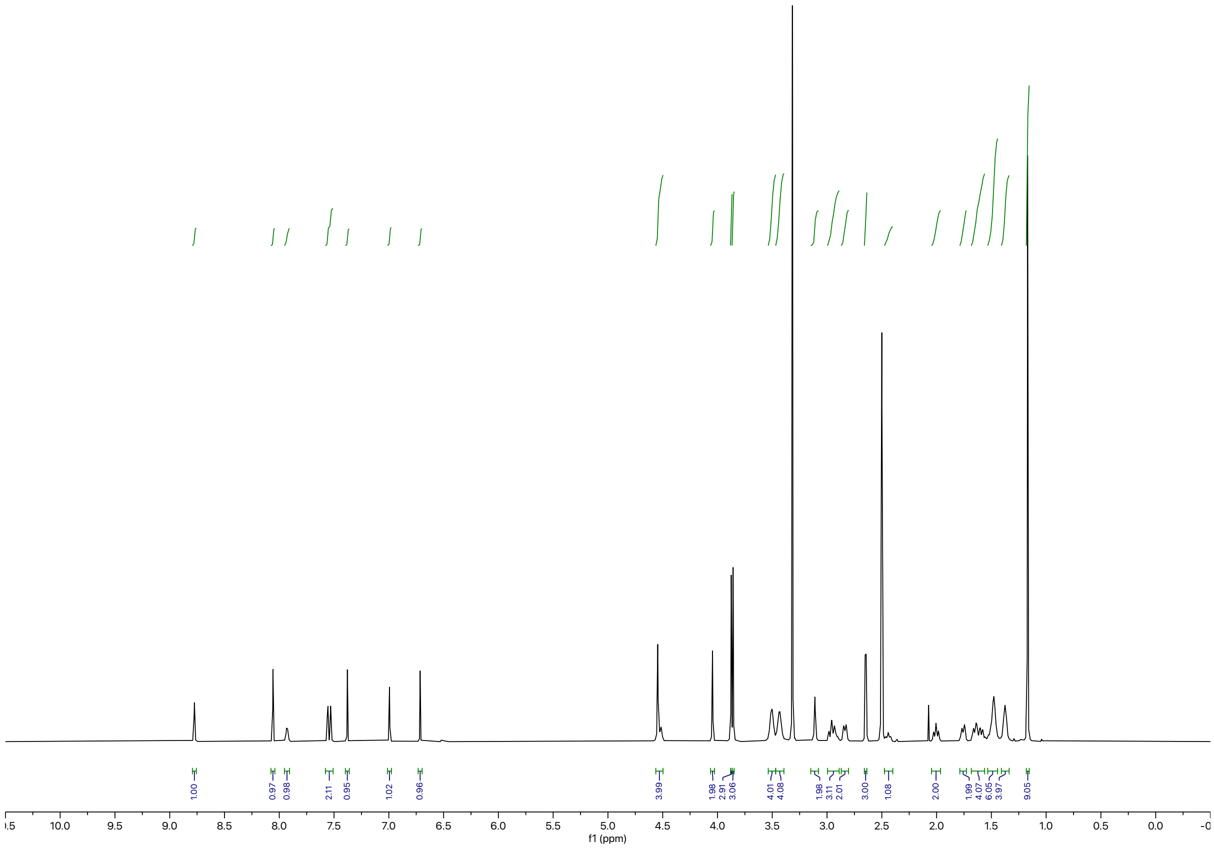

Synthesis of **BAK-04-198**

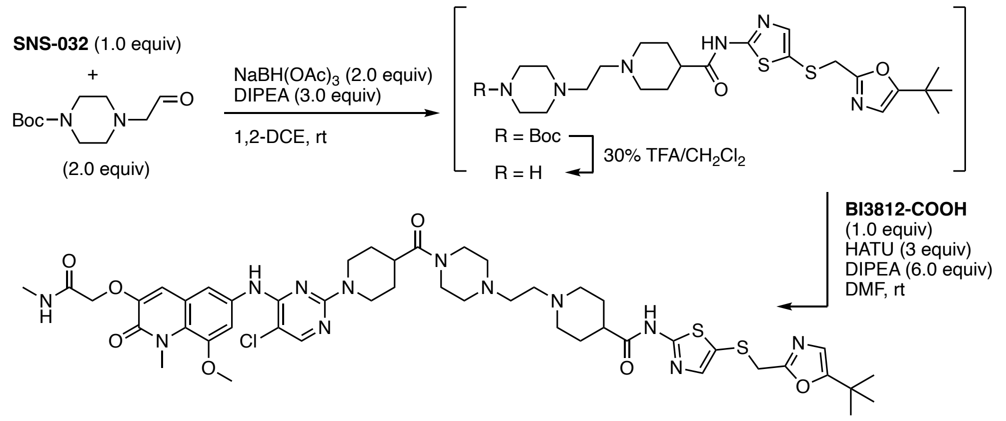


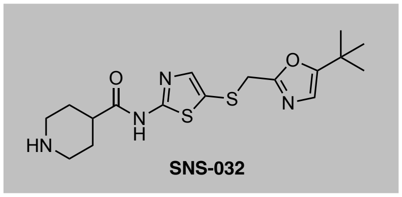
SNS-032

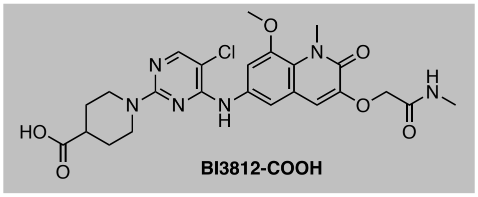
BI3812-COOH

To a solution of *tert*-butyl 4-(2-oxoethyl)piperazine-1-carboxylate[15] (0.10 g, 0.44 mmol, 2.0 equiv) in 1,2-DCE (2.0 mL) was added **SNS-032** (0.083 g, 0.22 mmol, 1.0 equiv) and DIPEA (0.11 mL, 0.044 mmol, 4.0˚equiv). After stirring at ambient temperature for 10 min, $NaBH(OAc)_3$ (0.093 g, 0.044 mmol, 2.0 equiv) was added and the mixture stirred for 2 h. Water was added and the aqueous layer was extracted with $CH_2Cl_2$. The combined organic extracts were washed with water and brine, dried over $Na_2SO_4$ and concentrated under reduced pressure. The crude material was purified by flash column chromatography on silica (0-20% MeOH in $CH_2Cl_2$) to afford the intermediate (0.10 g, 77% yield). Boc deprotection was achieved in 30% TFA in $CH_2Cl_2$ (0.5 mL) at 35 °C, residual TFA was removed under high vacuum. Then, a solution of the deprotection product in DMF (2.0 mL) was slowly added to a solution of **BI3812-COOH** (0.090 g, 0.17 mmol, 1.0 equiv), HATU (0.19 g, 0.51 mmol, 3.0 equiv), and DIPEA (0.18 mL, 1.0 mmol, 6.0 equiv) in DMF (2.0 mL) at 0 °C. The mixture was allowed to warm to ambient temperature and after 3 h the reaction was stopped by addition of water (10 mL). The resulting precipitate was filtered off and purified by reverse-phase HPLC (25–85% MeOH in water) to afford **BAK-04-198** as white solid upon lyophilization (0.047 g, 28% yield from SNS-032).

**$^1$H NMR** (500 MHz, MeOD) δ = 8.06 (t, *J* = 1.4 Hz, 1H), 7.51 – 7.45 (m, 1H), 7.37 (d, *J* = 2.5 Hz, 1H), 7.32 – 7.30 (m, 1H), 7.13 – 7.03 (m, 1H), 6.66 (s, 1H), 4.51 (dd, *J* = 10.1, 4.9 Hz, 2H), 4.36 (d, *J* = 12.9 Hz, 2H), 4.06 – 3.90 (m, 8H), 3.9 – 3.67 (m, 6H), 3.59 – 3.43 (m, 2H), 3.23 (q, *J* = 14.0 Hz, 6H), 3.16 – 2.92 (m, 5H), 2.84 (t, *J* = 1.9 Hz, 4H), 2.15 (dtd, *J* = 21.9, 10.9, 3.8 Hz, 4H), 1.86 (d, *J* = 13.2 Hz, 2H), 1.76 (td, *J* = f13.4, 6.5 Hz, 2H), 1.24 (s, 9H). **LC-MS**: m/z 1005.57 $[M+1]^+$.

Synthesis of **BAK-04-338 (CDK-TCIP3)**

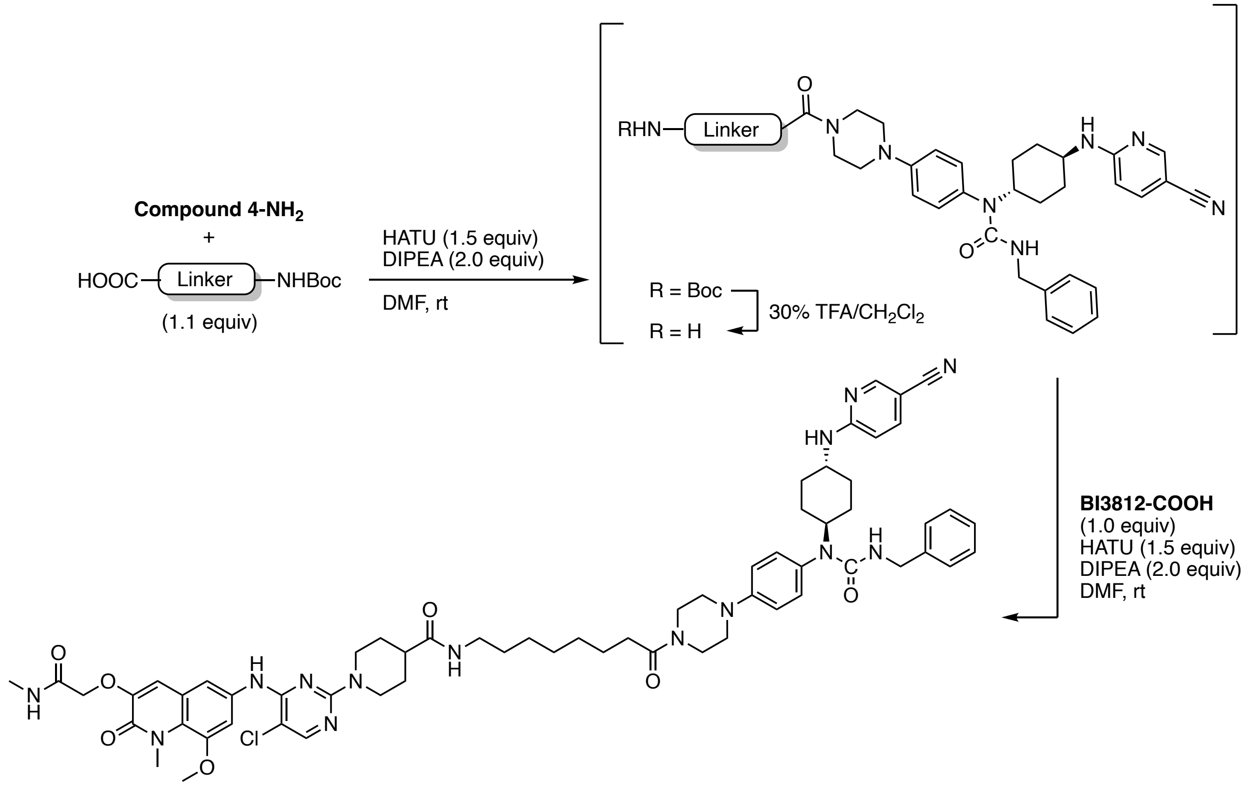


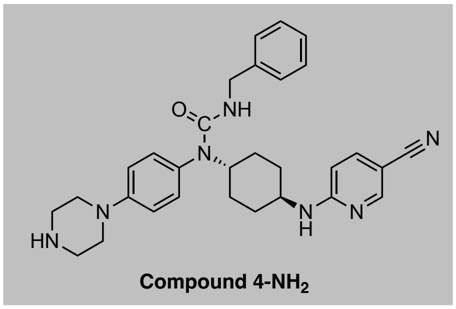


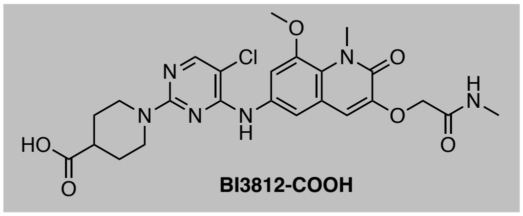


To a solution of **Compound 4-NH$_2$**[16] (9.0 mg, 0.015 mmol, 1.0 equiv), HATU (8.4 mg, 0.022 mmol, 1.5 equiv), and DIPEA (5.6 μL, 0.030 mmol, 2.0 equiv) in DMF (0.2 M) was slowly added 8-((*tert*-butoxycarbonyl)amino)octanoic acid (4.0 mg, 0.017 mmol, 1.1 equiv) as linker. The reaction mixture was stirred for 30 min. Water was added, and the aqueous layer was extracted with EtOAc and after concentration of the organic extracts, the crude material was purified by flash column chromatography on silica (0-10% MeOH in $CH_2Cl_2$) to afford the intermediate in quantitative yield. The intermediate was taken up in $CH_2Cl_2$ (2.0 mL), TFA (0.6 mL) was added, and the yellow solution was stirred at 35 °C for 1 h, at which point LC-MS analysis indicated full removal of the Boc protecting group. The mixture was concentrated under reduced pressure and dried under high vacuum to remove excess TFA. Then, a solution of the residue in DMF (0.1 M) was slowly added to a solution of **BI3812-COOH** (9.6 mg, 0.015 mmol, 1.0 equiv), HATU (8.6 mg, 0.023 mmol, 1.5 equiv), and DIPEA (5.6 μL, 0.030 mmol, 2.0 equiv)

in DMF (0.1 M) at 0 °C. The mixture was allowed to warm to ambient temperature and then was directly purified by reverse-phase HPLC (15–85% MeOH in water) to afford **BAK-04-338** as a white solid upon lyophilization (6.3 mg, 29% yield).

**$^1$H NMR** (500 MHz, DMSO-$d_6$) δ = 8.97 (s, 1H), 8.29 (d, *J* = 2.3 Hz, 1H), 8.09 (s, 1H), 7.94 (d, *J* = 4.9 Hz, 1H), 7.77 (t, *J* = 5.7 Hz, 1H), 7.60 (dd, *J* = 9.0, 2.4 Hz, 1H), 7.53 (q, *J* = 2.3 Hz, 3H), 7.26 (t, *J* = 7.5 Hz, 2H), 7.20 – 7.13 (m, 3H), 7.04 – 6.95 (m, 5H), 6.47 (d, *J* = 8.9 Hz, 1H), 5.57 (t, *J* = 6.1 Hz, 1H), 4.55 (s, 2H), 4.48 (d, *J* = 13.0 Hz, 2H), 4.30 – 4.21 (m, 1H), 4.14 (d, *J* = 5.9 Hz, 2H), 3.86 (d, *J* = 7.7 Hz, 6H), 3.57 (t, *J* = 5.2 Hz, 4H), 3.46 (br, 1H), 3.21 – 3.10 (m, 4H), 3.01 (q, *J* = 6.6 Hz, 2H), 2.91 (t, *J* = 12.4 Hz, 2H), 2.65 (d, *J* = 4.7 Hz, 3H), 2.42 – 2.28 (m, 3H), 1.90 (d, *J* = 11.9 Hz, 2H), 1.76 (d, *J* = 11.8 Hz, 2H), 1.70 (d, *J* = 12.7 Hz, 2H), 1.50 (dt, *J* = 12.6, 7.1 Hz, 4H), 1.43 – 1.33 (m, 2H), 1.24 (d, *J* = 13.5 Hz, 8H), 1.09 (d, *J* = 12.5 Hz, 2H). **LC-MS**: m/z 1163.34 $[M+1]^+$.

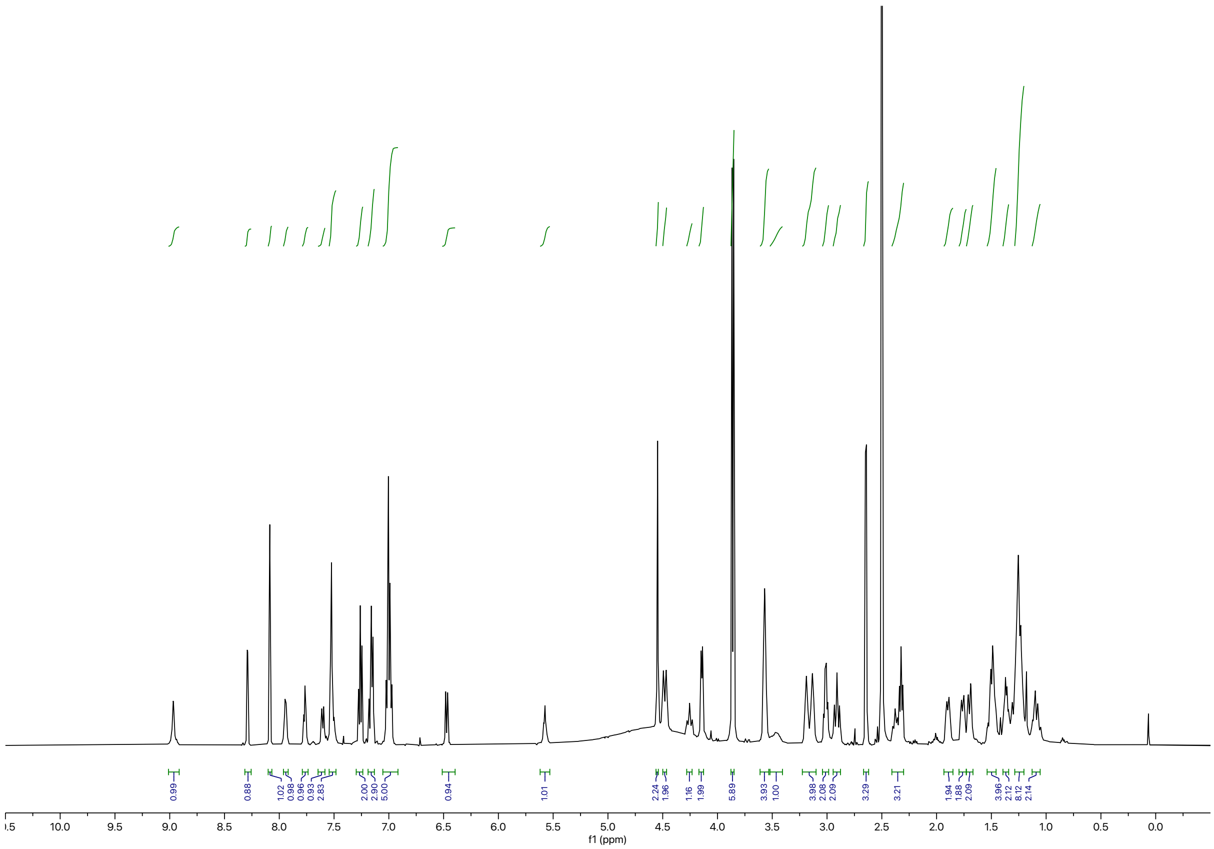

### 2. Full Scans of Western Blots

**Full scan of Fig 2F**

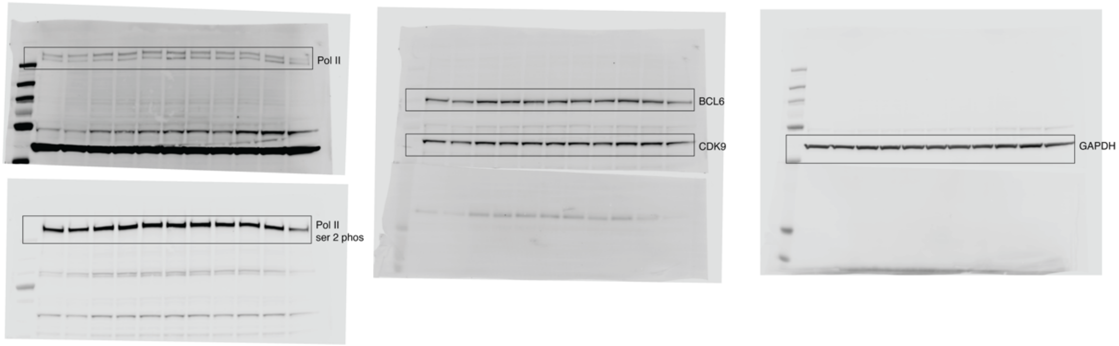


**Full scan of Fig 3D**

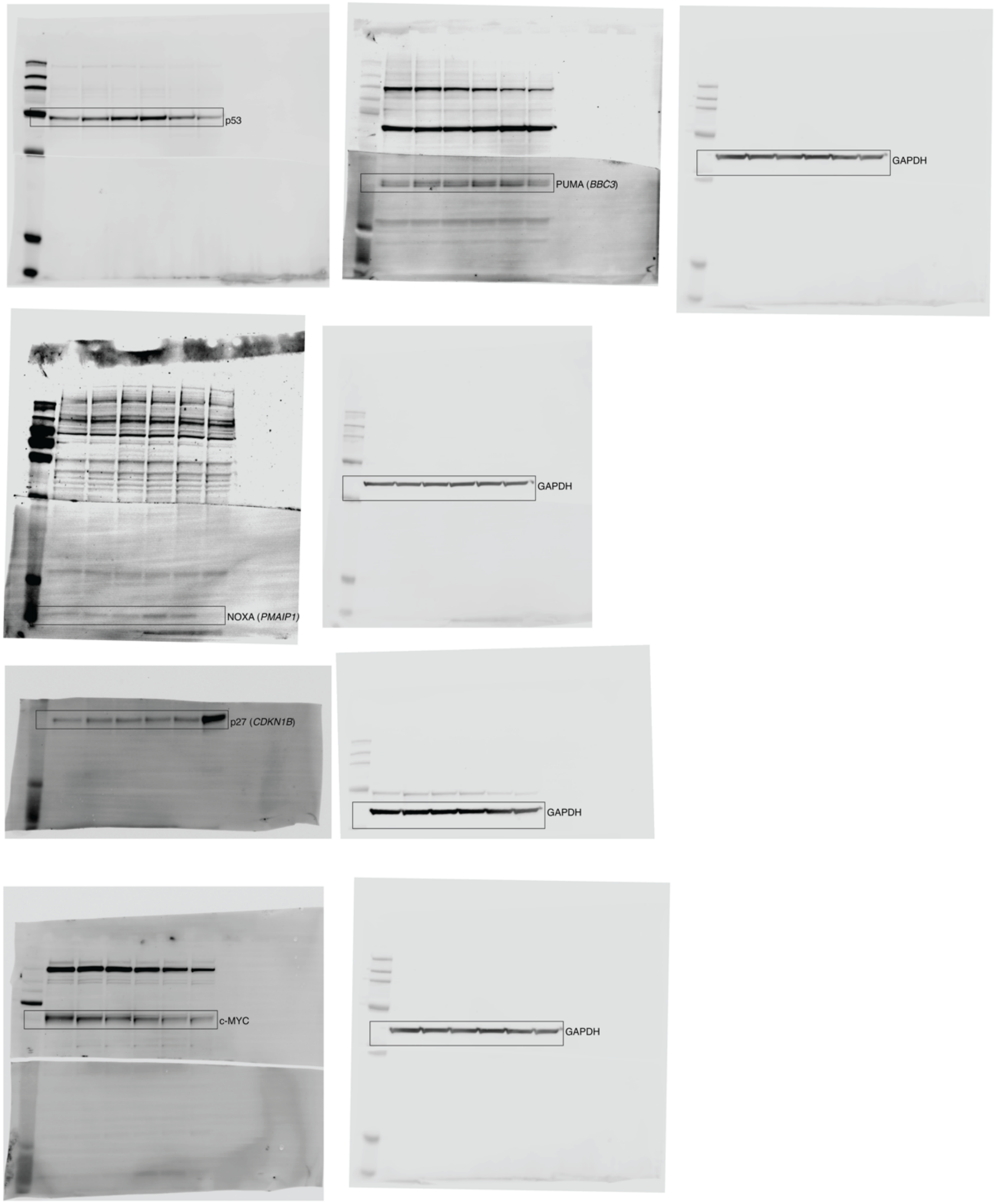

**Full scan of Fig 3E**

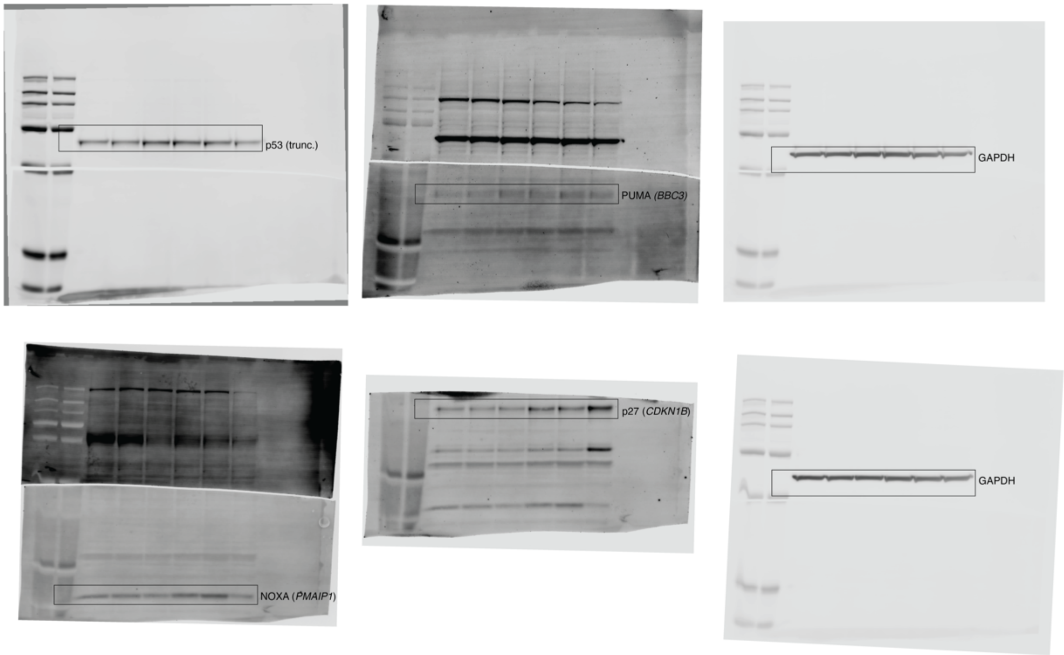

### 3. Flow Cytometry Gating Strategy

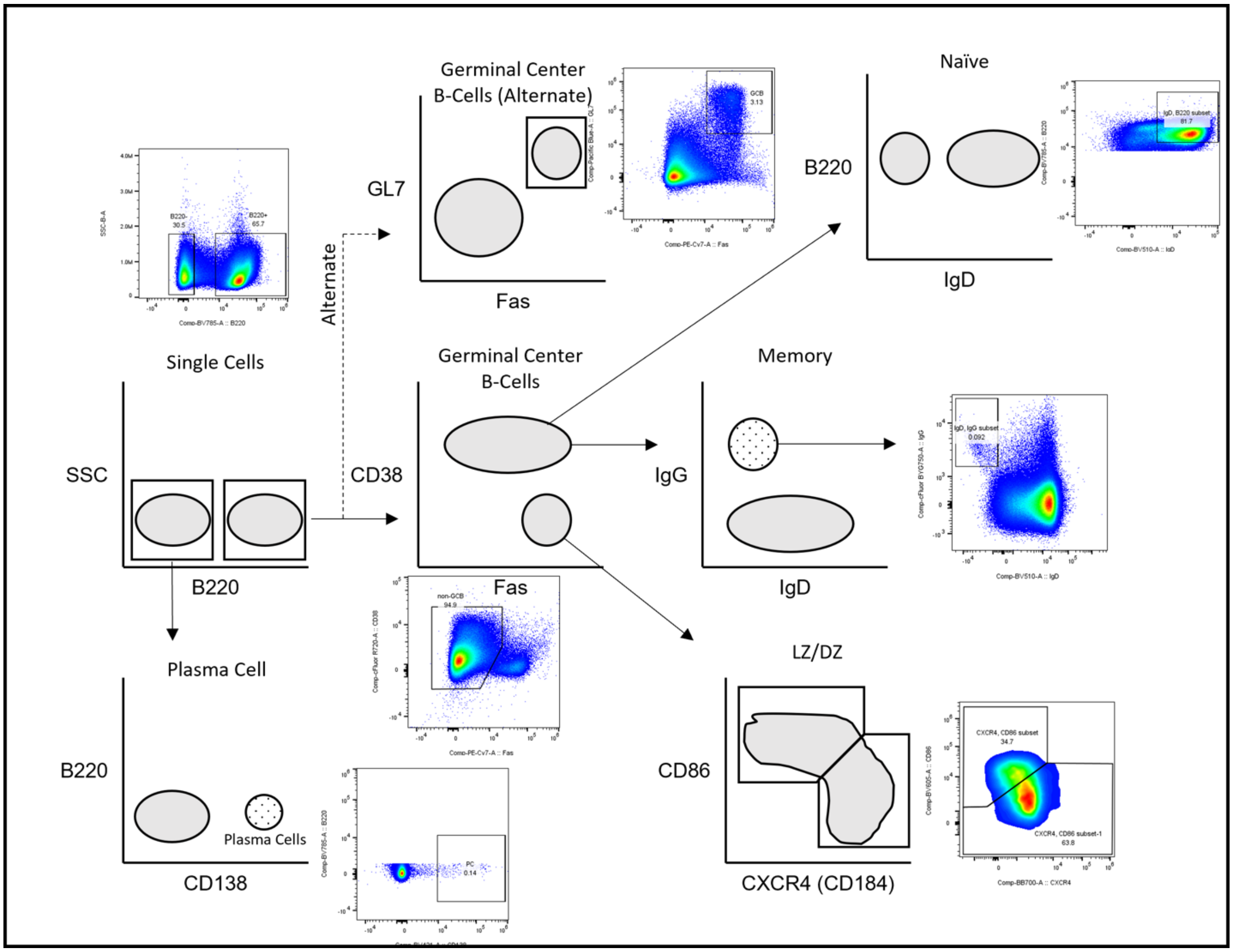
